# Widespread structural variations at human chromosome ends

**DOI:** 10.64898/2026.09.03.748401

**Authors:** Kar-Tong Tan, Ryan Jun Xiang Ong, Brandon Bing Rui Kee, Russell Ker Han Yap, Alicia Jun Ting Ng, Gihan Kaushalya Rajapaksha, Max Garrity-Janger, Qiyu Lin, Jerome Chong Rui, Cin Thet Kyi, Human Pangenome Reference Consortium, Matthew Meyerson, Heng Li

## Abstract

The highly repetitive regions of the human genome were long underrepresented from reference assemblies, limiting study of their biological function. Long-read sequencing and improved assembly algorithms have since resolved many of these regions, from centromeres to ribosomal DNA arrays, revealing structural variation increasingly linked to human disease. However, the subtelomeres, the repeat-rich regions adjacent to the telomeres at each chromosome end, have remained poorly characterized. Here we present a collection of complete subtelomeric sequences spanning all non-acrocentric chromosome arms, derived from 860 haploid assemblies across six ancestry groups. We find that while subtelomeres are mosaics of blocks shared between chromosome arms, individual arms diverge extensively, such that most non-acrocentric autosomal arms (54%, 21 of 39) carry multiple haplotypes differing by up to 100-200 kb. These blocks are broadly conserved across the great apes. In humans, their diversity is associated with chromosome arm rather than ancestry, suggesting that cross-arm paralogy block duplications predate human population divergence, although some haplotypes show ancestry-specific enrichment. Remarkably, these divergent haplotypes differ in gene content, driving gene copy-number variation between individuals among olfactory receptors and other genes. This study also revealed rare subtelomeric recombination. We further show that our subtelomere data set enables the accurate measurement of telomere length at individual chromosome ends from long-read data. Together, these assemblies reveal an unappreciated scale of variation at human chromosome ends and provide a resource for studying the roles of this variation in disease, telomere biology and genome evolution across diverse populations.

## Introduction

The past several years have witnessed remarkable progress in the completeness and resolution of the human genome. The completion of the first truly gapless human reference genome by the Telomere-to-Telomere (T2T) consortium revealed millions of previously inaccessible base pairs, including centromeric satellite arrays, ribosomal DNA arrays on the acrocentric chromosomes, and large segmental duplications that had resisted assembly for decades^1,2^. Recently, large-scale pangenomics efforts, notably by the Human Pangenome Reference Consortium (HPRC), the Human Genome Structural Variation Consortium (HGSVC), and other consortia, have generated hundreds of high-quality haplotype-resolved assemblies, providing an unprecedented view of human genomic diversity^3–8^. These advances, enabled by long-read sequencing^9,10^ and new assembly algorithms^11–17^, now yield near-complete, often telomere-to-telomere, diploid assemblies and allow characterization of previously intractable regions, revealing extensive variation invisible to earlier approaches^18–21^. Despite this rapid progress, the precise ways in which subtelomeric regions vary across human populations and how these influence human diseases remain largely unclear.

Human subtelomeric regions are operationally defined as the 500 kb immediately proximal to the telomeric repeat arrays at each chromosome end^22–24^, and occupy a structurally and functionally distinct niche in the genome. Rather than being uniformly organised, they form a mosaic patchwork of paralogy blocks, highly-conserved regions shared across at least two chromosome arms^23,25,26^, reflecting extensive interchromosomal exchange^25^. This exchange renders them hotspots for segmental duplication and genomic rearrangement, driving accelerated evolution relative to the rest of the genome. In humans, subtelomeres are enriched for rapidly evolving and biologically important gene families. These include the olfactory receptors, whose copy-number and positional variation has been characterised at individual loci^27–29^, and the WASH actin regulators, which are essential in animals yet vary in copy number across human chromosome ends^30^. As immediate neighbours of the telomere, subtelomeres may also influence telomere length in a chromosome-arm-specific manner through telomere-associated repeat (TAR) elements^31^. Subtelomeric rearrangements are also a recognised cause of idiopathic intellectual disability^32,33^. Beyond humans, subtelomeres in other organisms drive some of the most dramatic examples of this adaptability. In *Plasmodium falciparum* and *Trypanosoma brucei*, they house the *var* and *VSG* surface-antigen gene families^34,35^, whose continual recombination lets the parasite switch its molecular disguise faster than the host immune system can respond. Collectively, these properties place subtelomeres at the intersection of genome stability, gene regulation, and disease.

Despite their biological importance, comprehensive sequence-level characterisation of subtelomeric variation across the human population has remained out of reach. Initial restriction-fragment mapping in the 1990s established that human terminal repeats are polymorphic and structurally variable^36,37^, but covered limited chromosome ends without resolving their underlying sequence. Subsequent clone-based sequence assemblies produced a reference atlas of subtelomeric sequence^22,23,38^, and defined the paralogy blocks that structure these regions^23,25,26^, but yielded only a single composite reference per chromosome end. Optical mapping has surveyed subtelomeric structures across individuals^24,39^, but its resolution (∼5–20 kb marker spacing) cannot resolve the complex repetitive architecture of these regions, nor does it yield the nucleotide-level sequences needed to assess sequence divergence, gene content and functionality, or precise arm-specific read mapping^40^. The T2T CHM13 reference and HG002 diploid assembly provided the first complete single-nucleotide-resolution subtelomeric sequences^1,2^, but only one or two haplotypes per arm, far too few to capture population-level variation, rare structural forms, or differences across ancestral backgrounds.

Here we demonstrate that recent advances in genome assembly enable the generation of high-quality subtelomeric sequences at population scale. By systematically extracting and characterizing subtelomeric assemblies from 860 haploid assemblies across multiple initiatives spanning six major ancestry groups (African, AFR; Admixed American, AMR; East Asian, EAS; European, EUR; Middle Eastern and North African, MENA; and South Asian, SAS) (**Figure 1a**), we show that assembly completeness has improved substantially across successive data releases. These sequences reveal extensive haplotypic diversity, with 21 of 39 non-acrocentric autosomal arms (54%) harboring multiple distinct haplotypes that differ structurally by up to 100-200 kb, show ancestry-associated frequency differences, and, in several arms, vary in olfactory receptor gene copy number. This work establishes a population-scale map of human subtelomeric diversity, enabling these complex regions to be integrated into studies of genome evolution, gene regulation, and disease.

**Figure 1.**
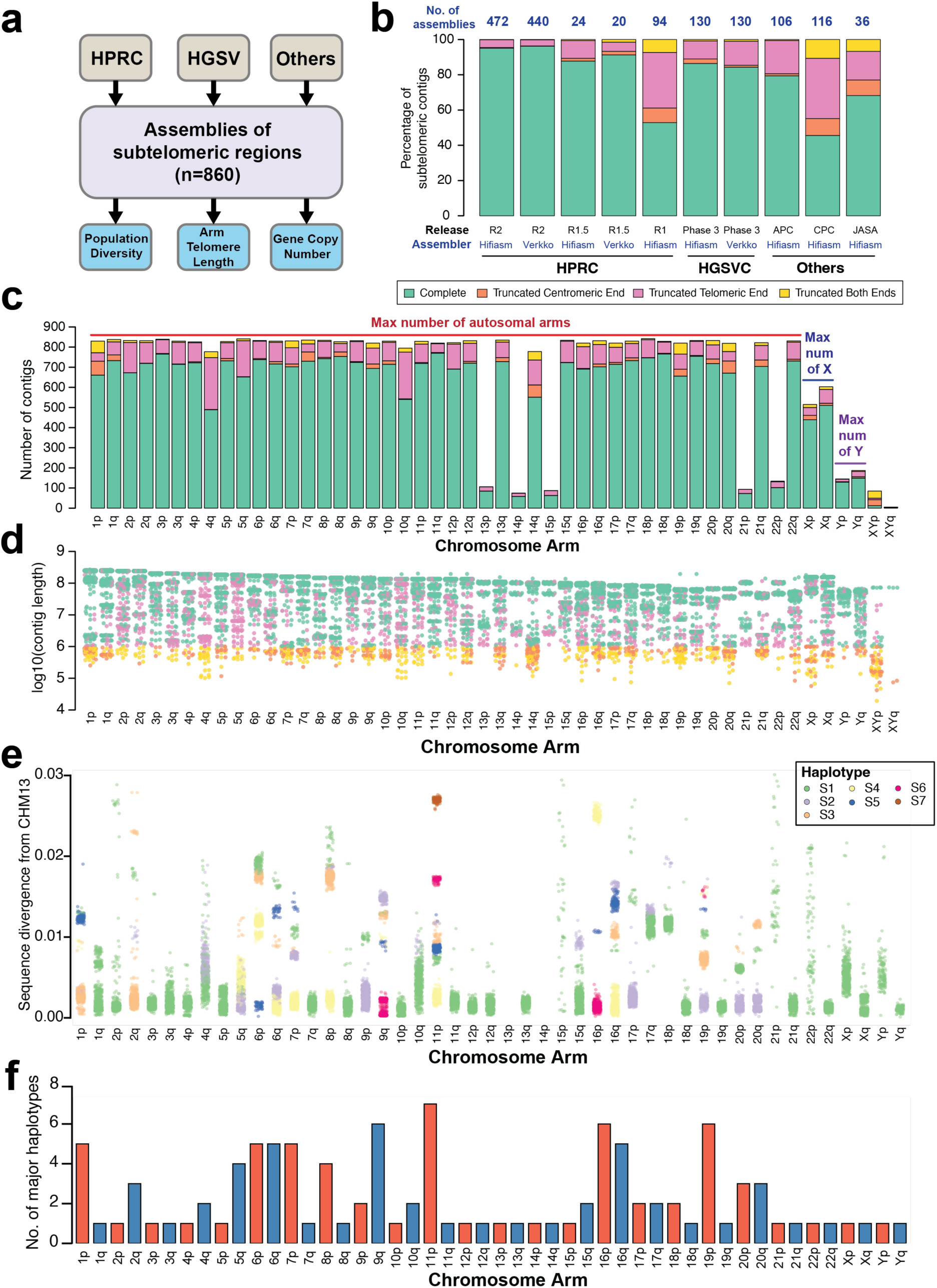
A high-quality pangenome collection of 860 subtelomeric assemblies reveals extensive haplotypic diversity across chromosome arms. (**a**) Overview of the study design, including sources of genome assemblies used for extraction of subtelomeric contigs and analyses performed. (**b**) Completeness of subtelomeric assemblies extracted from different studies. Contigs were classified as complete if they contained telomeric sequences at one end and spanned at least 1000 kb. Contigs lacking telomeric sequences were classified as truncated at the telomeric end; contigs shorter than 1000 kb from the telomeric end were classified as truncated at the centromeric end; contigs meeting both criteria were classified as truncated at both ends. (**c**) Number of subtelomeric contigs assigned to each chromosome arm, with bars coloured by completeness status. (**d**) Length of subtelomeric contigs per chromosome arm, coloured by completeness status. (**e**) Sequence divergence between the terminal 500 kb of each subtelomeric contig and the CHM13 reference genome, shown per chromosome arm. Each dot represents a single contig, coloured by subtelomeric haplotype (S1–S7). (**f**) Number of subtelomeric haplotypes (major only) observed per chromosome arm.

## Results

### Generating a high-quality reference pangenome of subtelomeric regions

We first collected genome assemblies (n = 1670) from five different consortia spanning diverse ancestries^3–8^, including the HPRC2 assemblies^41^ (**Figure 1a, Supplementary Table S1-S2, Methods**). These included multiple assemblies of the same haplotype generated by different assemblers, corresponding to 860 unique haplotypes. We then assigned each contig to its chromosome arm, requiring consistent assignment to the same arm across at least four of five reference sequences (**Methods**). Inspecting telomeric edges unexpectedly revealed repeat artifacts (mostly <1 kb) distal to long telomeric repeat tracts (often >2 kb) across consortia and assemblers (**Supplementary Fig. S1-S6**). As these telomeric tracts were too long to be interstitial (**Supplementary Fig. S1-S6**), the artifacts at the ends of these tracts were likely sequencing or assembly errors and were trimmed away (**Supplementary Note 1**, **Methods**).

We next assessed the quality and completeness of these subtelomeric sequences across different assemblies. Notably, while we found that subtelomeric sequences extracted from earlier genome assemblies (e.g. HPRC1) were often incomplete (52% complete), with many subtelomeric sequences missing telomeric repeat sequences at the telomeric ends (**Figure 1b**), subtelomeric sequences generated by HPRC2 were mostly complete, with >95% of subtelomeric sequences deemed to be complete with >1000 kb, ensuring the 500 kb subtelomeric region and all potential variation was fully captured, and terminated in telomeric repeats at the telomeric end. As some samples were sequenced and assembled by multiple assemblers and consortia, we curated a non-overlapping collection of subtelomeric sequences by prioritizing assemblies with higher fraction of T-to-T and arm-unique assemblies (**Methods**), comprising 860 subtelomeric assemblies of each chromosome arm (total = 35,150; mean = 40.9 per haploid assembly). We also compared contigs generated by different assemblers, and found that they were similar (**Extended Data Fig. 1** and **Supplementary Table S3**).

We then assessed the completeness of subtelomeric sequences for each chromosome arm (**Figure 1c**). Notably, only 12% of maximum expected contigs could be reliably assigned to the five acrocentric chromosome arms (i.e. 13p, 14p, 15p, 21p, and 22p). In contrast, we recovered complete assemblies for 57-90% of the non-acrocentric autosomal arms (mean = 82%). By accounting for the shared pseudoautosomal regions (PAR1 and PAR2)^42^ when mapping contigs to each sex chromosome (**Methods, Supplementary Fig. S7**), we recovered subtelomeric contigs for 91% of the sex chromosome arms, 81% of which were complete (**Figure 1c**). Consistent with this, contigs classified as complete were typically far longer than the 1,000 kb threshold, with some spanning entire chromosomes (**Figure 1d**). Thus, we generated a large, high- quality collection of subtelomeric sequences for the non-acrocentric arms across diverse ancestries.

### Subtelomeres show major haplotypic variation across chromosome arms

To assess how these subtelomeric sequences differ within each chromosome arm, we compared them to the CHM13 reference genome and calculated their sequence divergence (**Figure 1e, Methods**). We excluded contigs with truncated telomeric ends to ensure that only contigs with high coverage of the subtelomere were used (retained contigs per arm: min = 490, max = 776, mean = 717.5). While some arms showed low divergence (e.g., 1p, 2q, 9q, 10p), others diverged highly across all contigs. For instance, all 17q and 18p subtelomeric sequences showed significant divergence (>0.005) from the reference (**Figure 1e**), suggesting that most 17q and 18p subtelomeres in the human population deviate substantially from the CHM13 reference. Notably, we also observed distinct clusters of points which likely represent different haplotypes (**Figure 1e**). Since subtelomeres contain paralogy blocks^23–25^, highly conserved regions shared across multiple arms, we mapped the reference paralogy blocks from Stong *et al*.^23^ to the CHM13 subtelomeres (**Supplementary Fig. S8**), revealing arm-specific block arrangements that differed from prior literature^22,23,25,26^. Mapping these blocks to all pangenome subtelomeres, then filtering and clustering, revealed at least two major haplotypes in 21/39 non-acrocentric autosomal arms (**Figures 1f and 2**).

**Figure 2.**
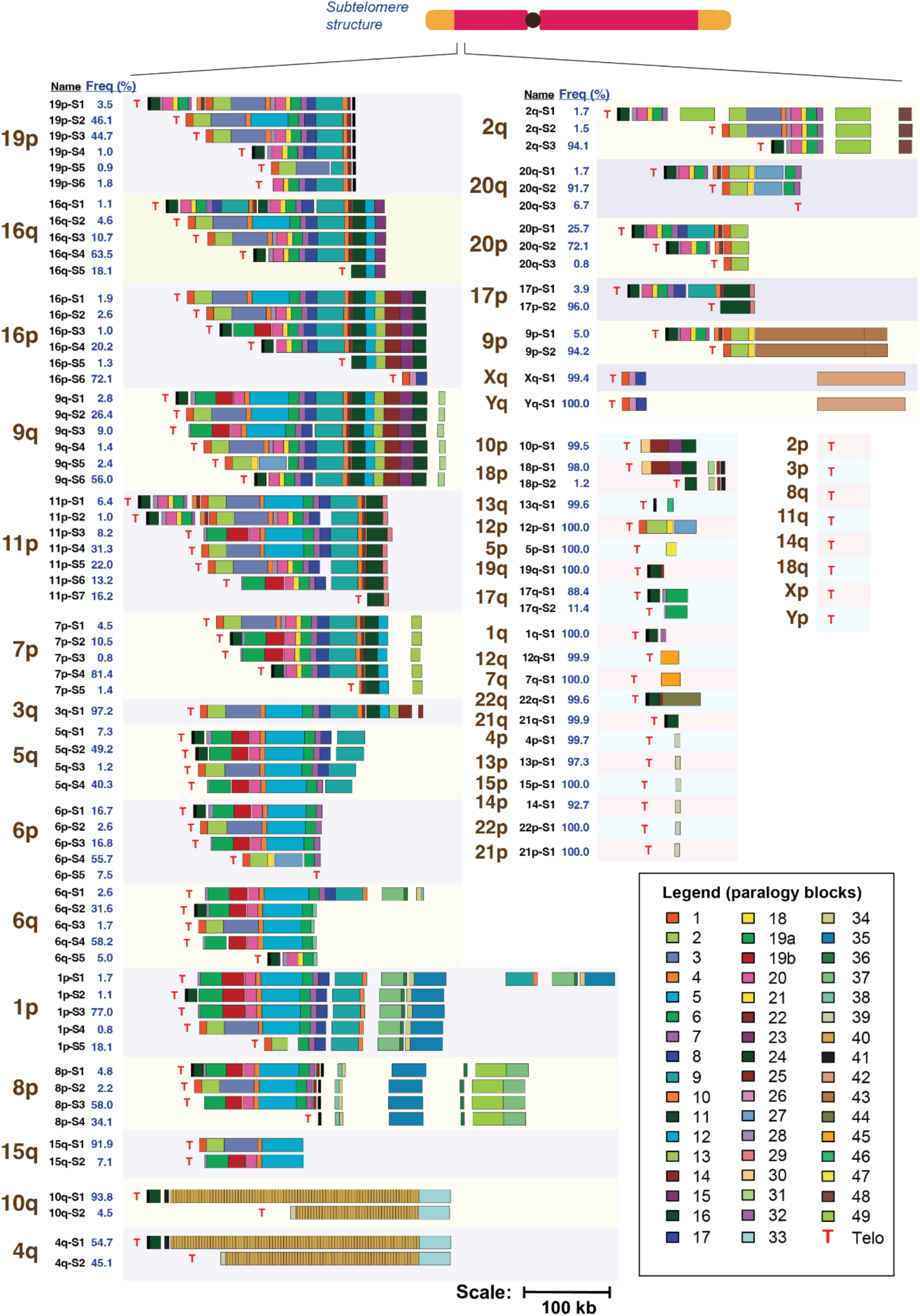
Large-scale structural variation defines distinct subtelomeric haplotypes across chromosome arms. Schematic depicting the major subtelomeric haplotypes identified for each chromosome arm. Haplotypes were mapped and compared using shared paralogy blocks (numbered 1-49, of which 19 is split into 19a and 19b, giving 50 blocks in total; see legend.), which reveal structural differences in composition and size across arms. Haplotypes are named S1, S2, etc. in descending order of size, with frequency across all assemblies indicated as a percentage. While some arms exhibit high structural variability with multiple distinct haplotypes, others contain a single haplotype with no paralogy blocks detected. "T" denotes telomeric sequences. Scale bar: 100 kb.

We next examined the structure of these paralogy blocks and how they differ across arms and haplotypes. Blocks were pervasive across the collection, present in 83% (40/48) of chromosome arms and in the majority of contigs within each (all arms with blocks had them in ≥50% of contigs and 73% in ≥80%), ranging from 7q (highest, 90%) to 14p (lowest, 53%) (**Supplementary Figs. S9-S48, Supplementary Table S4**). Because block arrangements varied by thousands of bases within the same arm, we clustered contigs into major haplotypes differing by at least two paralogy blocks of ≥3 kb (**Figure 2, Methods, Supplementary Table S5**). This resolved three or more major haplotypes in 29% (14/48) of arms, with 11p the most variable at seven haplotypes and three others (9q, 16p, 19p) with six. Notably, many of these haplotypes differed by up to 100-200 kb (**Figure 2**). Together, these results show that paralogy blocks serve as effective markers for resolving large-scale subtelomeric structural variation into distinct haplotypes across the population.

We next sought to understand where sequence variation occurs and how haplotypes are distributed within each arm. All-to-all comparisons between representative contigs of each major haplotype (**Supplementary Figs. S49-S54**) showed that differences between haplotypes of the same arm lay largely at the telomeric end, while centromeric ends were more likely to be homologous. Haplotypes were also unevenly distributed (**Figure 2**): in 71% (15/21) of multi-haplotype arms a single haplotype accounted for more than 50% of contigs, and even among the 14 arms with three or more haplotypes, 57% (8/14) were dominated by one (e.g., 11p-S4, the most prevalent haplotype in the most diverse arm, 11p, at 31%). A second round of clustering to capture finer-scale variation (**Supplementary Figs. S9-S48, Methods**) subdivided 43% (43/100) of major haplotypes into minor haplotypes, 18% (18/100) with two or more; their prevalence ranged widely, from 35% (28/81) present in at least 1% of the population to 38% (31/81) represented by a single contig. This distribution of major and minor haplotypes underscores the need for a large collection of subtelomeric sequences to both capture rare haplotypes and represent the full range of variation across arms.

### Subtelomeric structural variations differ by ancestry groups

The structural diversity captured across chromosome arms raises the question of whether subtelomeric haplotypes are randomly distributed across human populations or reflect underlying ancestry. To address this, we analysed haplotype frequencies across six superpopulations spanning 431 individuals from 29 subpopulations worldwide **(Figure 3a)**. Analysis of the 21 chromosome arms showing large-scale structural variations revealed ancestry-associated variation, with ten arm-ancestry combinations showing significant haplotype distributions **(Figure 3b, Extended Data Fig. 2, Supplementary Table S6-S7, Methods)**. AFR samples showed the most consistent pattern with significant haplotype distributions across seven arms, the strongest of which were 5q (q = 0.0002, Cramer’s V Effect Size = 0.516) and 16q (q = 0.0003, V = 0.578), where haplotypes 5q-S4 and 16q-S3 predominated at 81% and 33% respectively compared to markedly lower frequencies globally (5q-S4: 41%, 16q-S3: 11%). Population-specific differential distributions were also observed in non-African superpopulations, including strongly differentiated haplotype distributions in SAS on 2q (q = 0.0006, V = 0.342) and in EAS on 11p (q = 0.0002, V = 0.307) and 19p (q = 0.0003, V = 0.376). In contrast, MENA, AMR and EUR samples showed no significant differential distributions across any arm examined, though smaller sample sizes may limit power to detect modest effects. Together, these results show that subtelomeric haplotype distributions are substantially shaped by ancestral background, with AFR samples showing the broadest differentiation, consistent with the greater genetic diversity of African populations^43^.

**Figure 3.**
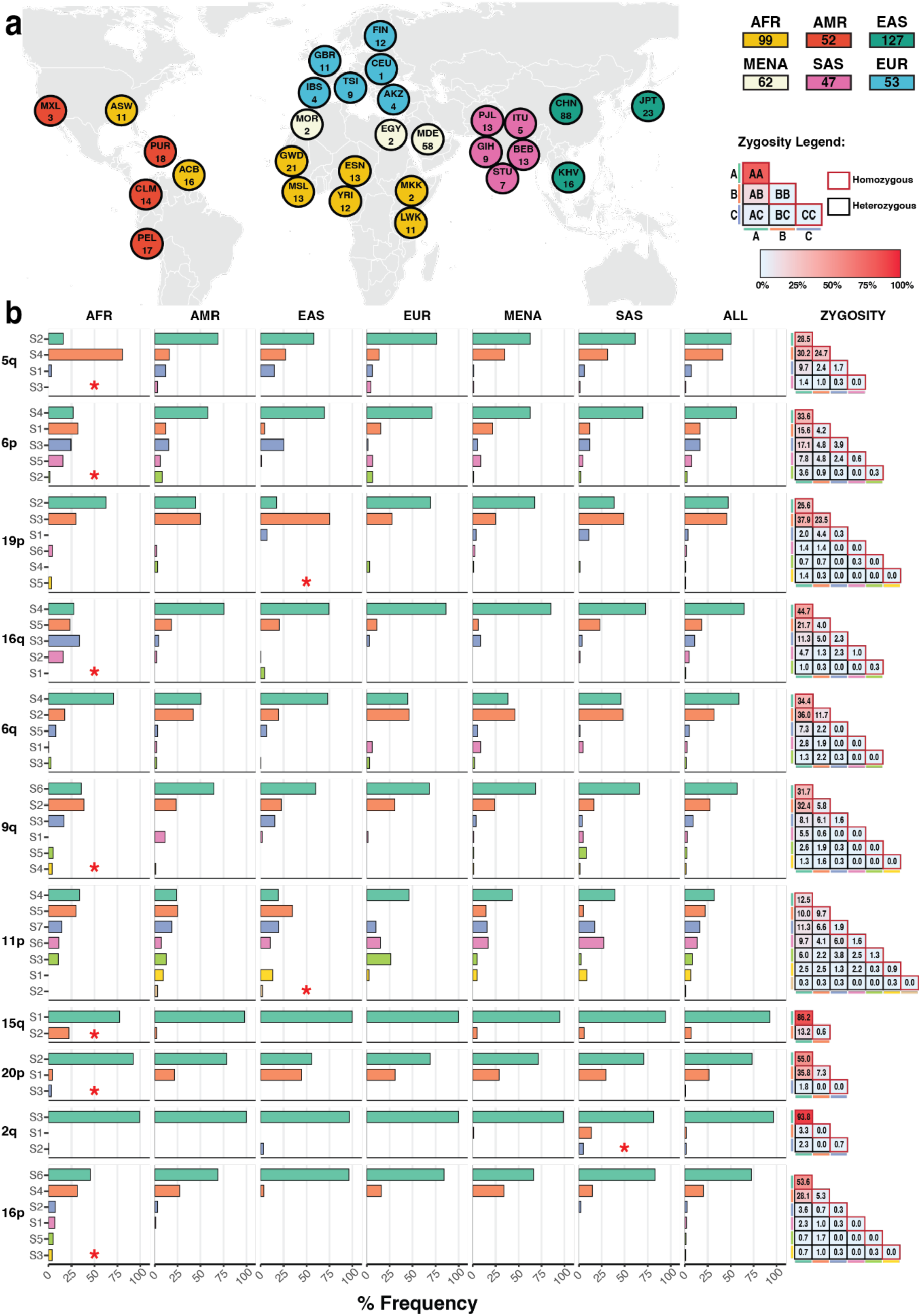
Subtelomeric haplotype frequencies show ancestry-associated variation across chromosome arms. (**a**) Geographic distribution of sampled populations grouped by major ancestry category: African (AFR), Admixed American (AMR), East Asian (EAS), European (EUR), Middle Eastern and North African (MENA), and South Asian (SAS), with sample sizes indicated. (**b**) Frequency of major subtelomeric haplotypes per chromosome arm across ancestry groups and the global population. The asterisk denotes a statistically significant differential haplotype distribution (Fisher’s exact test with Monte Carlo simulation, B = 10,000; Benjamini–Hochberg-adjusted q < 0.05) and a bias-corrected Cramér’s V > 0.3, indicating a moderate effect size. The zygosity matrix (right) shows the frequency of haplotype combinations in the global population for each arm.

To characterize how these haplotypes combine within individuals, we examined the diploid zygosity of subtelomeric haplotype combinations across the population (**Figure 3b, Supplementary Table S8**). Across most arms, the most common haplotype was frequently observed in the homozygous state, while less frequent haplotypes were rarely observed as homozygotes and instead occurred predominantly in heterozygous combination with the dominant haplotype. However, 5q and 19p were notable exceptions to this pattern. On both arms, the dominant haplotype was only marginally more frequent than the second (50% vs 41% for 5q, 47% vs 46% for 19p). On 19p, S2/S2 and S3/S3 haplotype combinations were observed at 26% and 24% respectively, while on 5q, S2/S2 and S4/S4 were observed at 28% and 25% respectively. Altogether, these findings reveal that subtelomeric structural variation is a population-structured feature of the human genome, with ancestry shaping haplotype identity and frequency.

### Cross-arm paralogy block duplications predate human population divergence

Given that paralogy blocks recur across multiple chromosome arms (**Figure 2**), we next assessed whether a given block is homogeneous across the arms it occupies, or whether its properties differ by arm. Using the HPRC2 cohort, we quantified per-individual paralogy block copy number and arm distribution (**Supplementary Fig. S55a,b**, median range: 2 - 94). Copy number and arm specificity varied considerably: Block 40 had the highest genome-wide copy number (median CN: 94) but was largely restricted to 4q and 10q, whereas Block 2 appeared across 23 arms despite lower copy number (median CN: 17). Arm-stratified length analysis showed that the same block can differ in length across arms: Block 2 ranged from 18kb to 26kb (**Supplementary Fig. S56**), while Block 40 maintained a uniform length (3.3kb) across both 4q and 10q (**Supplementary Fig. S57**). Copy number and length therefore differ between arms for the same block, indicating that paralogy blocks are not homogeneous units but are instead structured by the arm they occupy.

To resolve whether this arm structure extends to the sequences themselves, we moved beyond copy number and length to direct sequence comparison. Prior FISH-based studies showed that the same paralogy blocks can occur on different arms, but how similar these sequences are between arms was previously unknown^23–25^. To assess if paralogy blocks have similar sequences across arms, haplotypes or ancestry, we compared, for each paralogy block, sequence divergence across the arms, haplotypes, and ancestries^44^ (**Methods**). Clustering blocks by sequence divergence showed that most blocks cluster by arm identity rather than by haplotype or ancestry, indicating divergence is driven more by arm than by ancestry (**Supplementary Fig. S58-S106**). This clustering varied by block: Block 15 showed strong arm-level stratification with minimal ancestry signal (**Figure 4a and 4b**), while Block 20, present across more arms, showed poor stratification at both levels, suggesting greater sequence homogeneity and more frequent inter-arm exchange (**Figure 4c,d**). Haplotypes within the same arm were often poorly separated, indicating high intra-arm sequence similarity regardless of haplotype (**Supplementary Fig. S58-S106**). To quantify sequence divergence within block types, we used a variance-partitioning model with ancestry and haplotype as explanatory variables (**Figure 4e**). Haplotype explained more of the divergence than ancestry. Because haplotypes are nested within arms, this indicates that divergence is structured by arm and haplotype lineage rather than by ancestry, consistent with the clustering above (**Figure 4a and 4c**). Thus, the same haplotype lineages are shared across ancestries rather than being ancestry-specific, indicating that most cross-arm block duplications are ancient, predating human population divergence.

**Figure 4.**
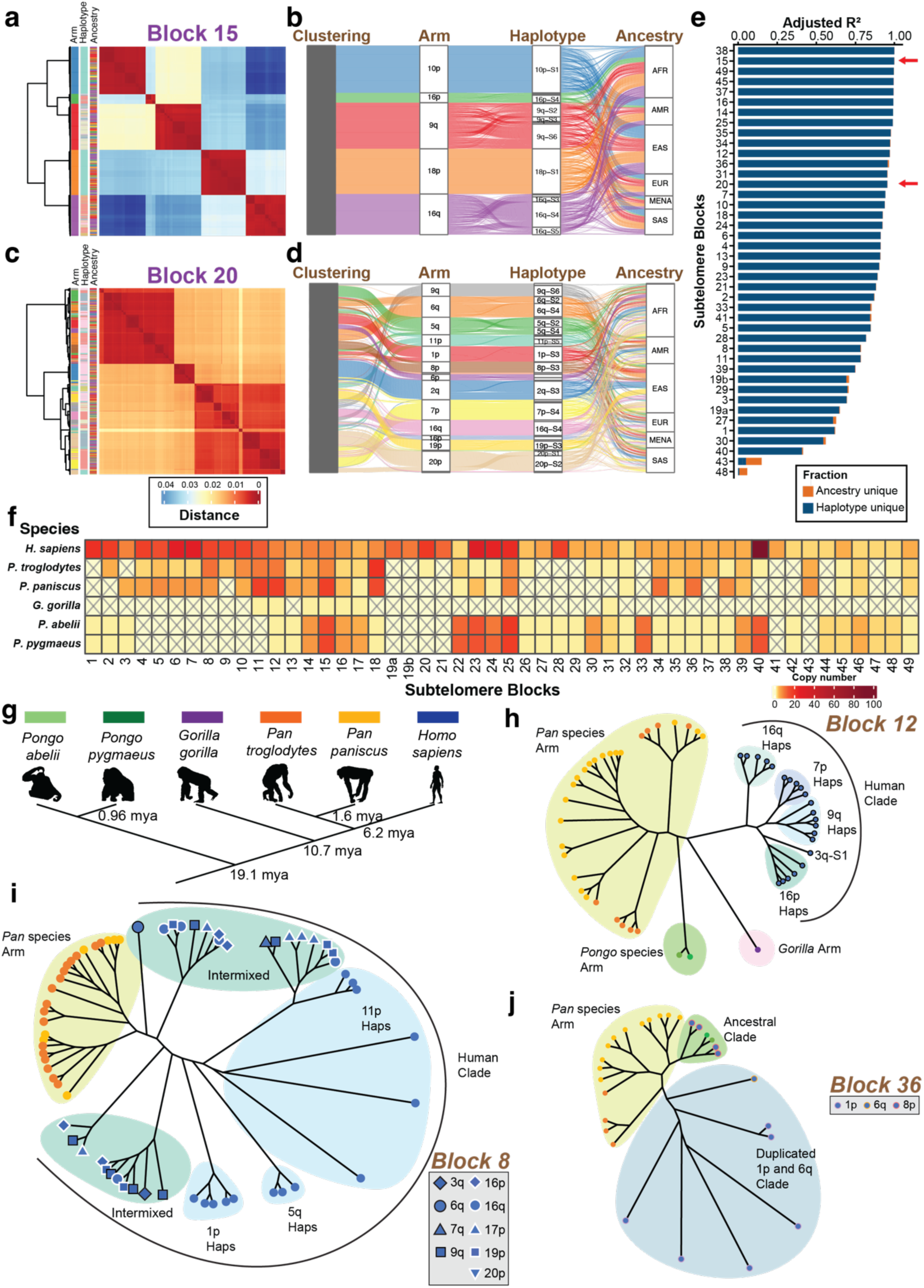
Paralogy block duplications predate human population divergence and show arm-specific, cross-species conservation patterns. (**a**) Heatmap of pairwise sequence similarity for a random subsample of 1,000 Block 15 sequences, with hierarchical clustering and annotation bars indicating chromosome arm, haplotype, and population ancestry. (**b**) Alluvial diagram showing how Block 15 sequence-similarity clusters map to chromosome arm, haplotype, and population ancestry (AMR, EAS, SAS, MENA, EUR, AFR). (**c–d**) As in (a–b) but for a random subsample of 1,000 Block 20 sequences, which shows less differentiated haplotype structure compared to Block 15. (**e**) Adjusted R² values from a variance partitioning algorithm for each subtelomeric block, showing the proportion of sequence variation explained by haplotype identity alone (blue) versus population ancestry alone (orange), ranked by total explained variance. (**f**) Copy number of all 50 subtelomeric blocks across six great ape species plotted on a log scale. Colour intensity reflects copy number; crosses indicate blocks absent in a given species. (**g**) Phylogeny of the six primate species examined, with estimated divergence times in millions of years ago (mya). from Mao *et al*. (2024). Species silhouettes are from PhyloPic, except *P. abelii*. (**h–j**) Maximum-likelihood unrooted phylogenetic trees for Blocks 12, 8, and 36, coloured by chromosome arm of origin (*Pan*, *Pongo*, *Gorilla* arms) and species (colours as in g); human sequences are representative sequences per haplotype. (**h**) Block 12 human clade separates into 16q, 7p, 9q, 3q-S1, and 16p haplotype subclades. (**i**) Block 8 human clade separates into 11p, 1p, and 5q haplotype subclades, alongside two intermixed clades containing multiple arms (3q, 6q, 7q, 9q, 16p, 16q, 17p, 19p, 20p). (**j**) Block 36 human clade separates into an ancestral clade and a duplicated 1p/6q clade.

### Paralogy blocks are broadly conserved across great apes with lineage-specific expansions

Given that paralogy block diversity is most associated with chromosome arm, rather than ancestry, we next assessed whether this pattern has deep evolutionary origins by tracing block dynamics and ancestral states across great apes. Blocks 2, 3, 5, and 20 were previously reported in some great apes, but no comprehensive survey exists^25^. Mapping blocks to recent great ape genome assemblies^45^ identified orthologues in all five species (**Figure 4f**, **Supplementary Fig. S107, Supplementary Table S9**). *Pan troglodytes* retained the most paralogy blocks from human subtelomeres (45/50); *Gorilla gorilla* retained the fewest (6/50). Most blocks were shared across species other than *Gorilla gorilla*; 19a, 19b, and 41 were present in humans only, suggesting that these three blocks arose in the human lineage. Block conservation was higher between the two *Pongo* species than between the two *Pan* species despite similar divergence times (0.96 vs. 1.6 mya; **Figure 4f–g**). Subterminal heterochromatin caps^25^, present in *Pan* but absent in *Pongo*, are associated with ectopic recombination between non-homologous chromosome ends and may drive this accelerated block turnover. The same mechanism may explain *Gorilla gorilla*’s low count of human-similar paralogy blocks despite its more recent divergence from humans (10.7 vs. 19.1 mya; **Figure 4g**). Together, these results indicate that most subtelomeric paralogy blocks predate human speciation, being present across multiple great-ape lineages. At the same time, the lineage-specific losses and turnover, and the few human-specific blocks (19a, 19b, 41), show that this block repertoire is not strictly conserved but continues to turn over within lineages.

We built phylogenetic trees for all paralogy blocks to assess their evolutionary histories (**Supplementary Fig. S108-S119**), revealing three broad categories (**Supplementary Table S10**). (i) Some blocks show clean arm stratification: Block 12’s topology matches the species tree (**Figure 4g**), with human copies clustering by arm and *Pan troglodytes* and *Pan paniscus* copies formed a shared clade (**Figure 4h**). This suggests the block had a post-speciation expansion with limited interchromosomal exchange. (ii) Block 8 also forms a human-specific clade (**Figure 4i**) but without arm clustering (e.g., 19p-S1, 19p-S3, 16q-S3, and 7p-S1 share a clade), indicating post-speciation expansion followed by frequent interchromosomal exchange that homogenized its sequence across arms. (iii) Block 36 displayed a bipartite topology separated by a long branch (**Figure 4j**): a basal clade comprised of human 8p and *Pongo* species’ 7p (syntenic to 8p)^46^, representing the ancestral form predating the divergence of *Pongo/*human-*Pan* split. The derived clade, restricted to human and *Pan* species spanning multiple arms, suggests a lineage-specific expansion after the human-*Pan*/*Pongo* split. Blocks 34-38 follow the same pattern, with ancestral clades anchored at 8p and derived clades at 1p (secondary expansion to 6p) (**Supplementary Fig. S116-S117)**, suggesting a single coordinated duplication in the human-*Pan* lineage. Together, these patterns show markedly distinct evolutionary trajectories across blocks, with interchromosomal exchange frequency varying drastically.

### Subtelomeric gene content differs between haplotypes

Subtelomeric regions are known to harbour multiple genes, including olfactory and immune-related gene families, located both within paralogy blocks and in-between these blocks^23,25^, concordant with our analysis (**Supplementary Fig. S120, Supplementary Table S11).** We next assessed whether haplotype identity influences gene content, by searching for regions with (1) valid open reading frames (ORFs), defined by Liftoff^47^ as those beginning with a start codon, ending with a stop codon, and free of premature in-frame stop codons, and (2) high sequence and coverage identity (**Methods**). We transferred CHM13 gene annotations to each contig and found that gene content varied substantially between haplotypes on the same arm (**Figure 5a, Extended Data Fig. 3-4, Supplementary Fig. S121-S122, Supplementary Table S11-S51**). For each haplotype, we quantified the percentage of contigs carrying a copy of the gene. Olfactory receptor gene presence was strongly haplotype-dependent: 98% of contigs carrying the 5q-S2 haplotype contained at least one complete copy of *OR4F3*, unlike other 5q haplotypes that lacked complete copies of *OR4F3* (5q-S1: 11%, 5q-S3: 0%, and 5q-S4: 38%). Similarly, *OR4F17* presence on 6p was enriched in 6p-S2 haplotypes relative to other 6p haplotypes (6p-S2: 68%, 6p-S1: 0%, S3-4: 0%). These results suggest that olfactory gene content differs between subtelomeric haplotypes.

**Figure 5.**
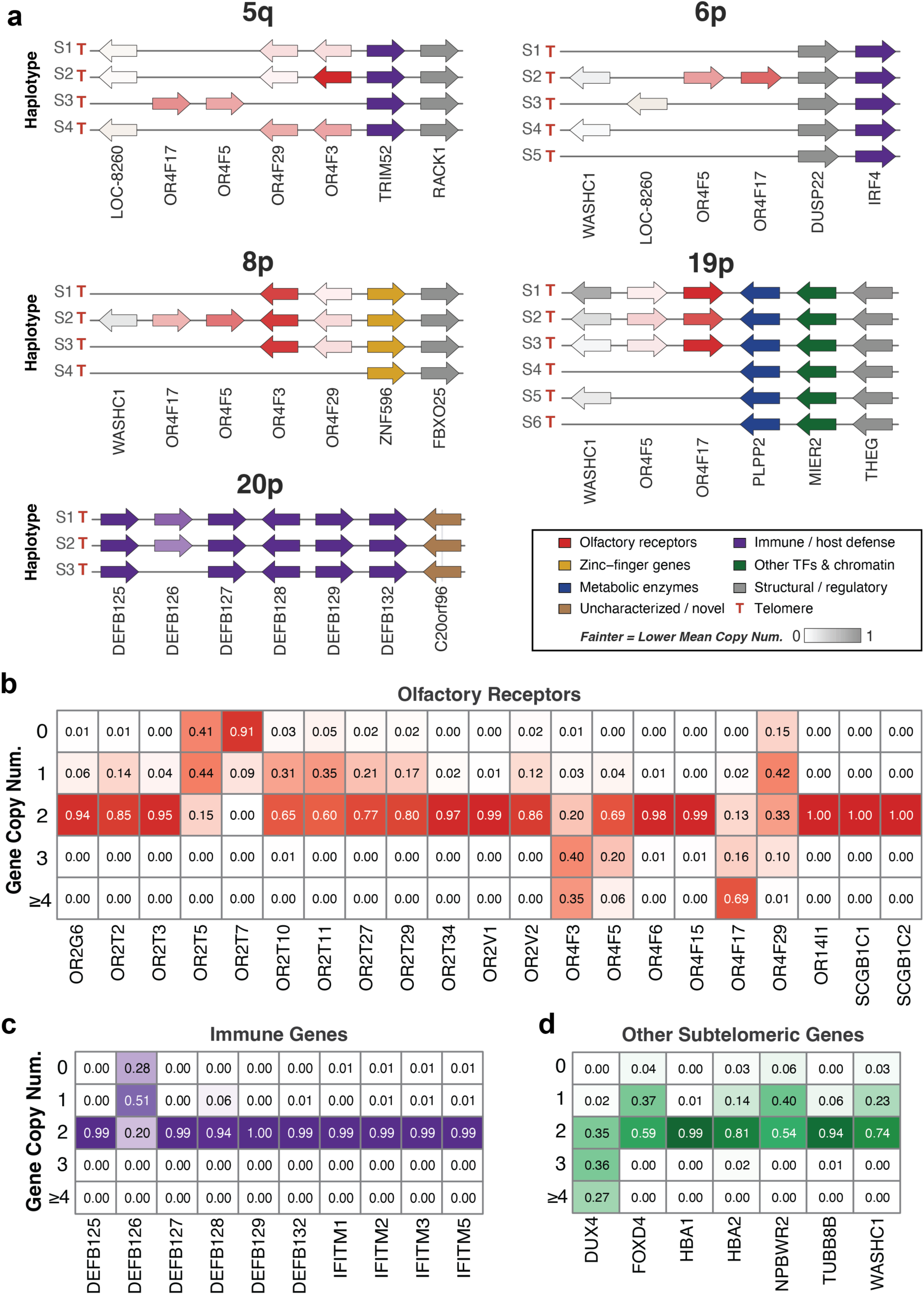
Structurally distinct subtelomeric haplotypes underlie gene copy-number variation in the human population. (**a**) Gene-arrow diagrams across haplotypes (rows, labelled S1–Sn; the number of haplotypes varies by arm) of five representative chromosome arms (5q, 6p, 8p, 19p, 20p), showing that haplotypes differ in gene content and arrangement. Genes are represented as arrows ordered along the panel from telomeric ("T") to centromeric, with arrow orientation indicating the direction of transcription. Arrows are coloured by functional category (see legend), with colour intensity indicating the mean copy number across all instances of the corresponding haplotype in the pangenome collection, from absent (white) to one copy (full colour). LOC112268260 is abbreviated to LOC-8260. (**b–d**) Heatmaps showing the frequency of each gene copy number (0, 1, 2, 3, ≥4) across all individuals in the cohort with complete diploid assemblies of the corresponding chromosome arm, for (**b**) olfactory receptor genes, (**c**) immune genes, and (**d**) other subtelomeric genes.

Beyond olfactory genes, immune gene content was also haplotype-dependent. On 20p, contigs assigned the S3 haplotype lacked fully intact open reading frames of *DEFB126*, a gene for which a common germline variant has been reported to be associated with impaired sperm function^48,49^, while S1 and S2 haplotypes retained the full-length *DEFB126* ORF at high frequency (**Figure 5a**, 20p-S1: 60%, 20p-S2: 42%, 20p-S3: 0%). There were also non-immune genes such as *HBA2*, known to undergo copy number variation in the context of α-thalassemia^50^, whose presence varied between subtelomere haplotypes (**Extended Data Fig. 3,** 16p-S1: 46%, 16p-S2: 67%, 16p-S3: 100%, 16p-S4: 93%, 16p-S5: 100%, 16p-S6: 94%), highlighting that haplotype-associated gene differences extend beyond canonical subtelomeric gene families reported in previous studies. *WASHC1*, a gene located within Block 2^23^ and involved in endosomal sorting^51,52^, is duplicated across multiple arms (e.g. 8p, 9p and 6q), but gene copies vary both between arms and between haplotypes within the same arm (**Figure 5a**, **Extended Data Fig. 3-4, Supplementary Table S11**, 8p-S2: 19%, 8p-S1/S3-4: 0%). These examples indicate that haplotype-linked differences in gene content are a general feature of subtelomeres, affecting genes across many functions rather than a few specialized families.

### Subtelomeric genes show extensive copy number variation at the population level

We next assessed whether the differences in gene content between haplotypes produce population-level variation in diploid gene copy number. Analysing gene copy number in each diploid genome, we found substantial variation across subtelomeric gene families between individuals (**Figure 5b-d, Supplementary Fig. S121-S122, Supplementary Table S52**). For olfactory receptor genes, the population displayed a broad range of copy number states: 92% of individuals carried zero copies of *OR2T7*, while 69% of individuals carried four or more copies of *OR4F17* (**Figure 5b**), illustrating that subtelomeric olfactory receptor content spans from near-complete gene absence to multi-copy amplification across the population.

We also assessed the gene copy number for subtelomeric immune genes in our cohort and found that the main gene with copy number variation is *DEFB126* (**Figure 5c**). As previously reported, this gene has multiple disease-associated variants that disrupts the ORF^48,49^. We found that a majority of the population (51%, n = 167/327) carried only a single functional copy of *DEFB126*, consistent with previously reported literature^49^. Inspection of *DEFB126* identified two known frameshift variants within the coding sequence: a 4bp deletion (19%, rs11467497) and a 2bp deletion (54%, rs140685149) (**Supplementary Fig. S123**), which disrupts DEFB126 protein expression^48,49^. Altogether, these results demonstrate the value of a high-quality collection of subtelomeric sequences for characterizing gene content variation in structurally complex genomic regions.

Among other subtelomeric genes (**Figure 5d**), *DUX4,* a gene involved in facioscapulohumeral muscular dystrophy^53^, exhibited notably high copy number, with 27% of sequenced individuals carrying four or more copies of *DUX4* with valid ORFs. Similarly, predicted copy number for *FOXD4,* a gene involved in neuronal differentiation^54^, differs within the population, with 41% of the sequenced individuals containing either 0 or 1 copy of the gene (n = 131/321). Closer inspection of *FOXD4* shows two main variants that results in a disrupted open-reading frame (ORF): a 1bp frameshift insertion (3.0%, rs532235124) and a C/G SNP that results in the loss of the stop codon (19%, rs79220013) (**Supplementary Fig. S124**). For *HBA2*, 14% of the sequenced population carried only a single copy, attributable largely to a large structural deletion (∼3.8kb) in the subtelomeric region that spans the entire gene (**Supplementary Fig. S125**). *NPBWR2, a gene* involved in neuroendocrine regulation^55^, also showed reduced copy number, with 46% of individuals carrying one or zero copy, driven primarily by variation in the 3’ UTR region (25%) (**Supplementary Fig. S126**). *WASHC1* copy number did not exceed 2 copies in the population, despite functional copies spanning 14 arms (**Supplementary Fig. S127**), suggesting a selective constraint limiting its expansion. Given these findings, it shows that subtelomeric gene copy number is highly variable across the population.

### Structural resolution of D4Z4 locus uncovers interchromosomal exchange between 4q and 10q

The subtelomeric 4q and 10q regions harbour the medically important D4Z4 repeat arrays (Block 40)^23^ which is a highly repetitive region that still lacks complete reference sequences. Contraction of the D4Z4 arrays has been associated with the expression of the *DUX4* gene which results in facioscapulohumeral muscular dystrophy (FSHD)^56^. Given the importance of this locus, we wanted to assess whether our collection might reveal unexpected features of these complex loci. As 9% of the 4q and 10q contigs are not fully assigned (**Figure 1c**), we first performed a rescue procedure to capture all 4q and 10q sequences (**Supplementary Table S53, Methods**). Examination of this locus identified 2 major haplotypes for each of the chromosome arms, S1 and S2, corresponding to the established A and B type alleles (**Figure 2**, **Figure 6a**)^57^. Across all D4Z4 repeats we unexpectedly found three clusters, Block 40α (4q-S1), 40β (10q-S1), and 40γ (4q-S2 and 10q-S2), with 40α and 40γ corresponding to the B^−^X^+^ 4q-type repeat and 40β to the B^+^X^−^ 10q-type repeat (**Figure 6b, Methods**)^57^. Further investigation of D4Z4 repeats also found that repeat number and type varied across haplotypes, with 4q-S1 haplotypes showing the broadest range and greatest diversity (**Extended Data Fig. 5a and 5b, Supplementary Tables S54-S55**).

**Figure 6.**
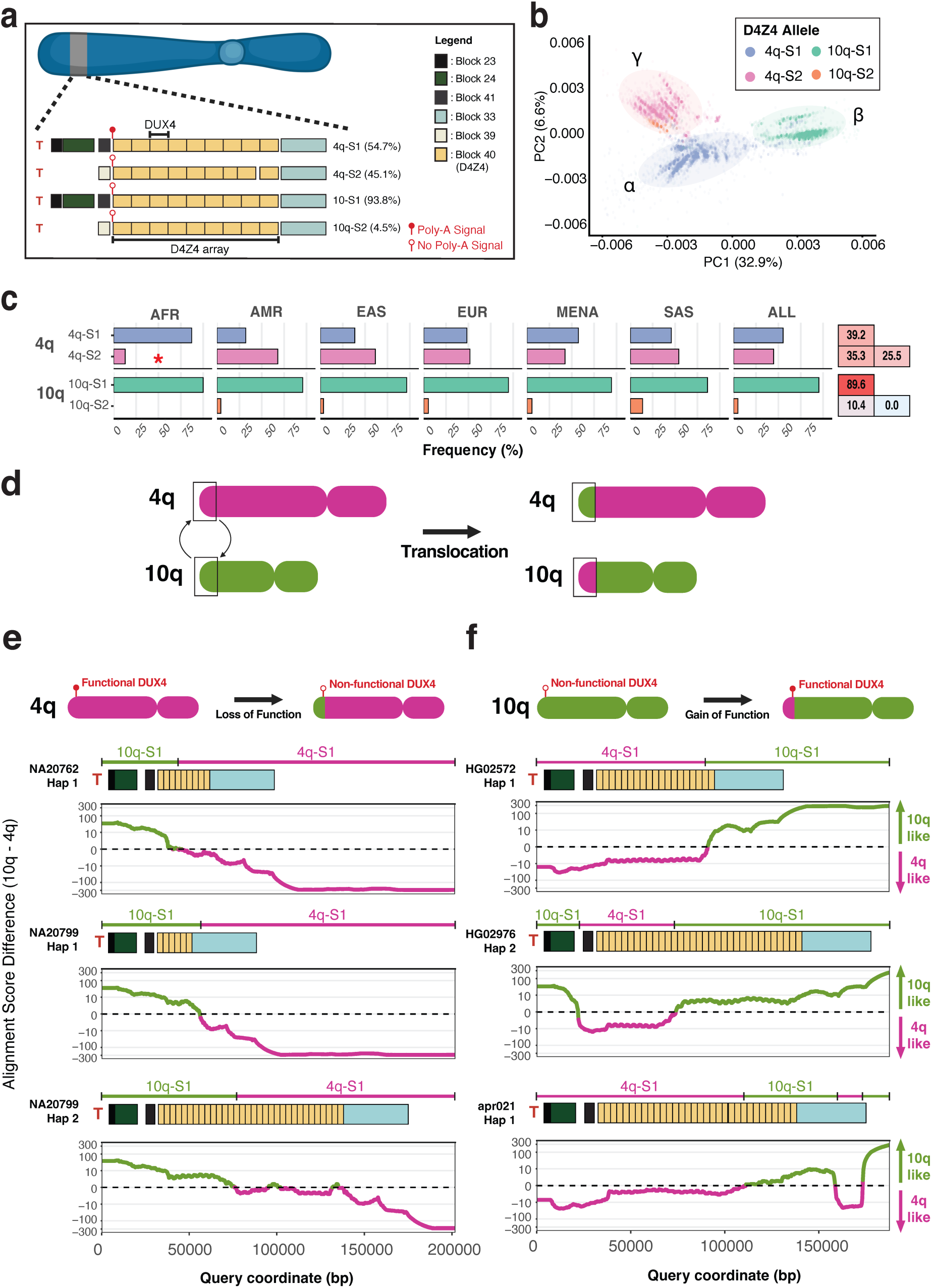
Subtelomeric exchange results in 4q *DUX4* functional loss and 10q risk-allele acquisition. (**a**) Schematic of 4q and 10q subtelomeric paralogy block structures. All 4q and 10q subtelomeres contain a D4Z4 tandem repeat array (paralogy Block 40 in yellow). Each D4Z4 unit contains a copy of the *DUX4* gene; stable DUX4 expression requires a polyadenylation signal (PAS) carried only by permissive haplotypes. Facioscapulohumeral muscular dystrophy (FSHD) arises when contraction of the array occurs on a permissive haplotype, allowing stable *DUX4* expression in muscle. (**b**) Principal coordinate analysis (PCoA) of individual D4Z4 units enables segregation into three distinct clusters (α,β,γ). (**c**) Population distribution of the four subtelomeric haplotypes (4q-S1, 4q-S2, 10q-S1, 10q-S2) across the six superpopulations. Statistical significance was assessed using a Monte Carlo simulation of Fisher’s exact test with Benjamini-Hochberg correction for multiple testing. The asterisk denotes a statistically significant differential haplotype distribution (BH-adjusted q < 0.05, bias-corrected Cramér’s V > 0.3), in the AFR superpopulation. Zygosity is shown in the accompanying matrix (right). (**d**) Schematic depicting a translocation between the subtelomeres of 4q and 10q. (**e**) Translocation of 10q-S1 sequences onto 4q-S1 subtelomeres. The distal exchange removes the PAS that stabilises the DUX4 transcript, rendering *DUX4* non-functional and the 4q-S1 allele non-permissive. Alignment score difference plots for three representative haplotypes show, along the subtelomeric region, whether each position aligns preferentially to a 10q reference (positive, green) or a 4q reference (negative, magenta). (**f**) Translocation of 4q-S1 sequences onto 10q subtelomeres. The distal exchange introduces a PAS, stabilising the DUX4 transcript and rendering the normally non-permissive 10q-S1 allele functional and permissive. Alignment score difference plots are interpreted as in (**e**).

Beyond the composition of the array itself, the pathogenic potential of a contracted D4Z4 array is determined by specific sequence elements at its terminal end. FSHD arises only when a functional polyadenylation signal (PAS, ATTAAA) within a 260bp sequence called pLAM stabilises the *DUX4* transcript^56^. Notably, the 4q-S1 allele carries this functional PAS while 10q-S1 carries an inactive variant (ATCAAA), explaining why contractions on 10q-S1 do not cause FSHD^56^. Resolution of the terminal D4Z4 region across our 4q-S1 assemblies revealed three structural configurations (**Extended Data Fig. 5c, Supplementary Table S55**): the established 4q-S1-S (92%) and 4q-S1-L (4.4%) alleles^58^, and a recently described intermediate, 4q-S1-M (1.5%), which has not yet been linked to FSHD^59^. Critically, the distribution of these terminal configurations, and the overall frequency of disease-susceptible 4q-S1 haplotypes, were both structured by ancestry: 4q-S1-M was restricted to AFR and MENA superpopulations (**Extended Data Fig. 5d, Supplementary Table S55**), and AFR samples carried 4q-S1 at substantially higher frequency (82%, V=0.395) than the global average (53%) (**Figure 6c, Supplementary Tables S56-S58**), underscoring the need for ancestrally diverse cohorts in FSHD research.

Interestingly, our assemblies also revealed rare interchromosomal exchange between 4q and 10q subtelomeres (**Figure 6d**, **Methods**). Among 4q-S1 contigs, 7/275 (2.5%) showed evidence of 10q-S1-derived sequence displacing the functional PAS at their terminal ends, rendering them unable to support pathogenic *DUX4* expression despite their chromosomal origin on 4q (**Figure 6e, Extended Data Fig. 5e**). Conversely, 3/514 (0.58%) 10q-S1 contigs carried 4q-S1-derived sequence introducing a functional PAS onto an otherwise non-susceptible allele (**Figure 6f**). As an orthogonal check against assembly artefacts, we independently confirmed a detected translocation using HiFi long reads and the corresponding Hifiasm assembly of the same sample (**Extended Data Fig. 5f,g**). Together, these findings demonstrate that interchromosomal exchange occurs at low but detectable frequency in the general population, with the potential to both abolish and confer the ability to support pathogenic *DUX4* expression depending on the direction of exchange.

### Subtelomere pangenome reference improves arm-specific telomeric read mapping accuracy

High sequence similarity between subtelomeric arms makes it hard to assign long-read sequences spanning telomeric/subtelomeric regions to the correct arm. We next assessed if mapping to large diverse collection of subtelomeric sequences (i.e. a pangenome reference) could improve mapping accuracy. We simulated reads from the terminal 50 kb of donor assemblies and mapped them to (1) CHM13, (2) the donor’s own assembly, and (3) the pangenome reference (**Figure 7a, Methods**). Mapping accuracy was lowest against CHM13 (69%), higher against the pangenome (94%) and highest against the self-assembly (99%) (**Figure 7b, Supplementary Table S59**). Read accuracy, depth, and technology had minimal effect on mapping accuracy (**Figure 7b, Supplementary Fig. S128a**). Longer read length improved telomeric read mapping accuracy overall (e.g. CHM13: 49-63%; **Figure 7b, Supplementary Table S59**).

**Figure 7.**
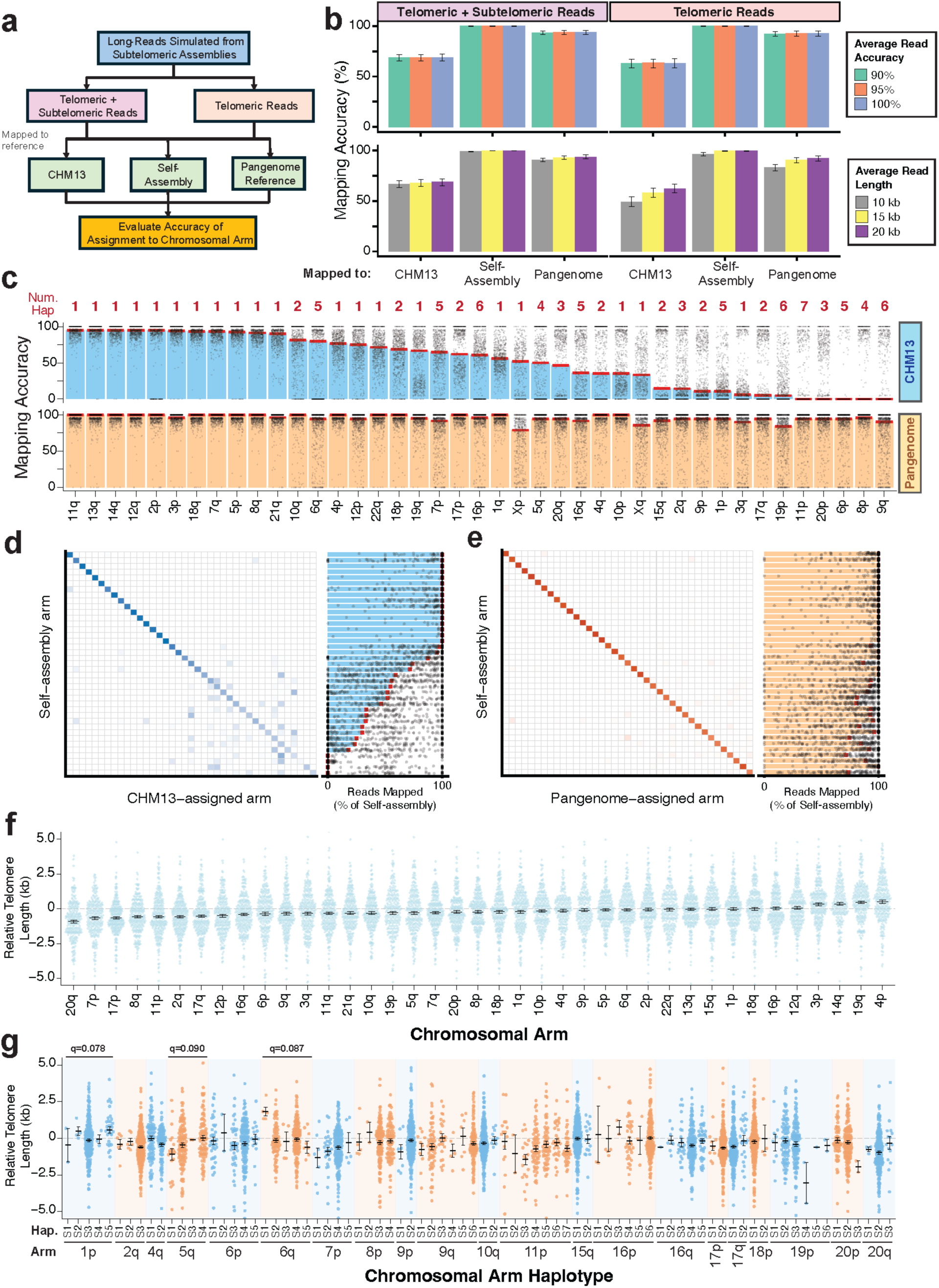
A pangenome subtelomere reference facilitates chromosome arm-specific telomere length assessment. (**a**) Schematic of the simulation pipeline. Reads are simulated from the subtelomere assemblies in our cohort using PBSIM3, then partitioned into (1) all subtelomeric reads and (2) subtelomeric reads containing telomeric repeats. Each subset is mapped to CHM13, the pangenome reference, and the source assembly. (**b**) Barplot of mapping accuracy for all subtelomeric and telomeric-only reads across references, stratified by simulated read accuracy (90%, 95%, 100%; *top*) and average read length (10kb, 15kb, 20kb; *bottom*). (**c**) Per-chromosomal-arm mapping accuracy of simulated telomeric reads (15kb, 95% read accuracy) mapped to CHM13 (top, blue) and the pangenome reference (bottom, orange). Each point represents one sample. The number of haplotypes per arm is indicated above each panel (Num Hap). The red line indicates the median mapping accuracy across all samples for that arm. (**d**) Heatmap showing the comparison between chromosome arms assigned by mapping to the self-assembly reference and arms assigned by mapping to the CHM13 reference. Color intensity reflects the percentage of reads from each self-assembly arm (rows) that were assigned to each corresponding CHM13 arm (columns), normalized within each row. The adjacent bar plot shows, for each arm, the percentage of reads assigned to the same arm by both methods (i.e., diagonal concordance), with bars indicating the median across samples and points showing individual sample values. (**e**) as in (**d**), but comparing arm assignment between the self-assembly reference and the pangenome reference. (**f**) Relative telomere length (kb), normalized to the per-sample average telomere length, shown per chromosome arm. Box-and-whiskers represent the SEM across samples. (**g**) Relative telomere length (kb) per chromosome arm stratified by subtelomeric haplotype group. Chromosome arms are displayed in alternating blue and orange for visual distinction. Box-and-whiskers represent the SEM. Significant differences in telomere length across haplotype groups within each arm were assessed by Kruskal-Wallis test with Benjamini-Hochberg correction; arms with q-values annotated (1p: q = 0.078, 5q: q = 0.090, 6q: q = 0.087).

Given this improvement, we assessed whether mapping accuracy varies by chromosome arm using these simulated long-reads. CHM13 accuracy was non-uniform across arms (**Figure 7c**), with minimal effect from ancestry (**Supplementary Fig. S129**). Reads from 11q and 13q mapped to CHM13 with 95% median accuracy, while 8p and 9q reads mapped with 0% median accuracy (**Supplementary Table S60**). These reads were misassigned to other arms sharing the same subtelomeric blocks (8p reads misassigned to 6q; 9q reads misassigned to 11p and 3q; **Figure 2, Supplementary Fig. S130–132**). Pangenome mapping accuracy stayed consistently high across arms (91-100%) except at acrocentric arms and sex chromosomes (**Supplementary Fig. S128b, Supplementary Table S60**). These results track haplotype diversity: poorly mapped CHM13 arms had multiple subtelomere haplotypes (8p: 4; 9q: 6), while high-accuracy arms had only one (**Figure 7c, Supplementary Fig. S128c**).

We next assessed if real telomeric long-reads extracted from the HPRC2 long-read dataset can be accurately assigned to chromosome arms using the pangenome reference, and whether they can reveal arm- or haplotype-specific differences in telomere length. To do this, we mapped the HPRC2 long reads to the pangenome reference, to the CHM13 reference, and to each sample’s own genome assembly (i.e. self-assembly), which served as ground truth. Benchmarking against the self-assembly, the pangenome assigned telomere-containing reads to the correct arm more reliably than CHM13 (**Figure 7d,e, Supplementary Tables S61–63**). Concordance was consistently high for the pangenome (median 100%, mean 87.2%) but far more variable for CHM13 (median 91.7%, mean 66.6%), where the low mean reflects a subset of arms that mapped at near-zero accuracy. We then calculated relative telomere length (normalized to per-sample average) for each chromosome arm^40^, based on mapping against the self-assembly. Relative lengths formed a broad, continuous distribution, from -875bp (shortest, 20q) to +534bp (longest, 4p) **(Figure 7f**). Arms 20q, 17p, and 11p trended shorter; 9p, 4p, and 14q trended longer. As a quality check, the number of reads was evenly distributed across pangenome arms, indicating no arm-specific bias in the assignment of telomeric long-reads (**Supplementary Fig. S133**). We then tested whether telomeric repeat length associates with subtelomeric haplotype, comparing relative telomere length across major haplotypes within each arm (Kruskal-Wallis, **Figure 7g, Supplementary Table S64**). Three arms showed suggestive differences at a relaxed FDR < 0.1 threshold: 1p (q = 0.078, nominal p = 0.004), 5q (q = 0.090, nominal p = 0.013), and 6q (q = 0.087, nominal p = 0.009). All others were non-significant (q > 0.1, **Supplementary Table S64**). These marginal associations suggest, rather than establish, that subtelomeric haplotypes defined in this study influence telomere length at a subset of arms. Together, these results establish the pangenome as a reliable reference for arm-resolved telomere length measurement, and point to subtelomeric haplotype as a potential factor influencing telomere length at specific chromosome ends, a hypothesis that will require confirmation in larger, better-powered cohorts.

## Discussion

Our study reveals that human subtelomeres are not merely repetitive chromosomal appendages but a reservoir of extensive, ancestry-structured haplotypic variation with broad functional and technical consequences. Across 21 of the 39 non-acrocentric chromosome arms, we find major haplotypic differences, exceeding 100 kb in some cases, that segregate with ancestry, reshape local gene content, and confound the assignment of telomeric reads to their correct arms. This scale of variation, resolved here at complete-sequence resolution across 860 haplotypes, indicates that a single reference genome cannot capture subtelomeric diversity, and that this diversity is biologically consequential rather than incidental. In this light, the subtelomere emerges as an underappreciated axis of human genomic variation—one that links population history, gene-family evolution, and the accurate measurement of telomere length.

A key discovery of this study is the unprecedented level of structural heterogeneity characterized at the sequence level across a large fraction of human subtelomeric regions. While subtelomeric structural variation was documented well before sequence-resolved assembly, from early restriction and FISH-based mapping of polymorphic terminal regions^29,36,37^ to more recent optical mapping^24,39^, these approaches captured only sparse markers across a handful of individuals, resolving neither nucleotide-level variation nor complex structural rearrangements at the scale of population diversity. Here, leveraging previously defined paralogy blocks as high-resolution sequence markers across a large panel of 860 haplotypes, we generated a comprehensive structural map that reveals each haplotype at single-nucleotide resolution. This approach uncovered massive structural variations, with haplotypes on several chromosome arms differing by over 100 kb in sequence content. This scale and mode of variation are distinct from the tandem-repeat enrichment previously described at chromosome ends^60,61^, which was assessed across the terminal 5 Mbp and reflects variation in tandem repeat-unit number within individual loci, rather than the arrangement of paralogous blocks we resolve within the most distal ∼500 kb. By bridging the gap between low-resolution physical maps and base-level assemblies, our work provides an unmatched look into the true architectural diversity of human subtelomeres.

A striking finding from our study is the remarkable gene-content variability of human subtelomeric regions, driven primarily by olfactory receptor gene families whose copy number ranged from zero to more than four (**Figure 5b**), unlike the two copies typical of standard autosomal genes. Although copy-number and locational polymorphism of subtelomeric olfactory receptor blocks has long been recognized^27–29^, earlier low-resolution approaches could not reliably distinguish intact receptors from pseudogenized or truncated copies. By resolving the complete sequences of these genes, our data identify which copies retain intact ORFs, reframing copy-number variation as functional gene dosage rather than raw locus count. Full-length resolution further reveals two structural architectures: (1) localized haplotype-specific deletions or gene inactivation reduce copy numbers, while (2) inter-chromosomal duplications across highly homologous paralogy blocks drive expansions, distributing genes like *OR4F3*, *OR4F5*, and *OR4F17* across multiple distinct chromosome ends. These variants differ markedly across populations, suggesting that ancestry-specific haplotype frequencies shape olfactory receptor copy number and, potentially, odorant sensitivity. Ultimately, just as single-nucleotide variants that alter or disrupt olfactory receptor function shape odour perception^62^, our findings show that subtelomeric structural variation is a previously overlooked source of human olfactory copy-number diversity that may similarly influence smell across populations^63^.

Beyond olfactory genes, we observed considerable variations in other subtelomeric genes such as *FOXD4*, *DEFB126* and *DUX4*. In particular we were able to capture a rare interchromosomal translocation between the 4q and 10q subtelomeres that transfers a medically relevant risk locus onto an otherwise non-susceptible chromosomal background, a finding independently corroborated by recent studies^64,65^. Having complete sequences also allows for better quantification of variant frequency across the population, enabling the detection of haplotype-associated differences in gene copy number, and ORF validity across clinically relevant subtelomeric loci that would be challenging to resolve with conventional approaches.

Subtelomeres have long been characterized as among the most plastic regions of the genome, acting as hotspots for interchromosomal recombination and segmental duplication^25^. Our analysis of paralogy blocks in this large cohort offers a window into how these structures originated and evolved. (1) First, the block sequences themselves are ancient, with most also present in the great apes and only a smaller subset (e.g. 19a and 19b) restricted to humans, indicating that they largely predate the human lineage. (2) Later, these blocks spread to multiple arms, as our phylogenetic tree shows that blocks on different arms are more similar to one another within humans than each is to its counterpart in the other great apes, indicating that they were duplicated across arms within the human lineage. Notably, this aligns with recent T2T ape assemblies demonstrating lineage-specific, mosaic segmental duplications at acrocentric and subtelomeric compartments^45,66^. (3) This spread nonetheless predates the divergence of human populations, as the diversity of these cross-arm copies tracks chromosome arm rather than population ancestry in our data. (4) Further interchromosomal recombination^25^ then shuffled these blocks into the new arm haplotypes we observed in this study. (5) Sequence exchange between copies further shaped these blocks. We find that this exchange is not global: even when a block is present on many arms, our data show that sequence is transferred preferentially among a specific subset of those arms, presumably through gene conversion^67^, so that some copies were homogenized while others remained distinct. This arm-structured exchange is consistent with concurrent work identifying discrete subtelomeric sequence communities across the pangenome^68^, which further links such structure to three-dimensional proximity of chromosome ends at meiosis. Together, these findings reveal subtelomere evolution as a layered process, in which ancient blocks spread across arms within the human lineage and were then reshaped by recombination and by structured, arm-specific sequence exchange at chromosome ends.

Our study also provides a foundational resource for the telomere biology community. Resolving telomere length at individual chromosome ends is of broad biomedical interest^40,69–74^, but long-read approaches have been hampered by reference bias, where aligning to a single reference genome fails to assign terminal reads to the correct chromosome arm. Building on our earlier use of a draft HPRC1 pangenome^3^ to show that human telomere length is chromosome end-specific^40^, we trace this assignment difficulty directly to the haplotypic divergence we uncovered here: arms with high subtelomeric diversity showed lower mapping accuracy, which the pangenome references released here resolve. Profiling telomeric repeat lengths across haplotypes, we found preliminary evidence that specific subtelomeric haplotypes, most notably on 1p, 5q, and 6q, may associate with arm-level telomere length, suggesting that they could act in cis to influence length at their respective chromosome ends. These findings are limited by two factors: our randomly sheared long-read sequences cannot confirm capture of the terminal telomeric repeats, unlike enrichment methods targeting the true chromosome-end overhang^40,69,70,72^, and the lymphoblastoid cell lines used here are prone to immortalization-associated telomere abnormalities^75^. Nonetheless, they provide a framework for validation in larger primary-tissue cohorts and point to a possible link between subtelomeric haplotype architecture and chromosome-end maintenance.

Several limitations should be noted. First, clustering based on paralogy block structure struggles to fully capture interblock differences at high resolution; for example, on 1q, distinct clusters of points are apparent that are not resolved by the block structure alone. Second, paralogy block-based analysis may not fully represent subtelomeric variation, exemplified by the fact that some genes fall outside of annotated block boundaries. Third, differences in assembly quality across contributing consortia may introduce bias; we mitigated this by restricting analysis to complete subtelomeric assemblies throughout but this may underrepresent haplotypes that are intrinsically harder to assemble, including complex or ancestry-specific forms. Finally, gene prediction using Liftoff is reference-biased, as gene validity is assessed solely against CHM13 sequence and its HPRC-derived annotation, and our stringent filtering for intact ORFs and high alignment coverage favours specificity over sensitivity, so genuine genes absent from or divergent in the reference may be missed.

In conclusion, our work reveals widespread structural variation at human chromosome ends and provides comprehensive maps of these long-overlooked regions. We anticipate these maps will serve as a springboard for connecting subtelomeric variation to telomere biology, gene regulation, and disease susceptibility across diverse human populations.

## Methods

### Reference genomes

Six reference quality genomes were used as part of this study. These were GRCh38, CHM13, HG002-maternal, HG002-paternal, CN1-maternal, and CN1-paternal^1,2,76,77^.

### Assessment of genome assemblies

We compiled high-quality diploid human genome assemblies from five major long-read sequencing initiatives: the Human Pangenome Reference Consortium (HPRC-Y2) (n = 486), the Human Genome Structural Variation Consortium Phase 3^5^ (HGSVC3) (n = 130), the Chinese Pangenome Consortium^6^ (CPC) (n = 116), the Japan–Saudi Genome Project^8^ (JaSaPaGe) (n = 36), and the UAE-based Arab Pangenome Reference^7^ (UPR) (n = 106).

All assemblies were generated using long read sequencing technologies and haplotype resolved assembly pipelines provided by each consortium. These datasets span diverse global populations and were downloaded from their publicly released data freezes. All assemblies were processed uniformly using our subtelomere extraction and analysis workflow to ensure consistency across cohorts.

### Identification and extraction of contig corresponding to each chromosome arm

As contig sequences from genome assemblies extracted from these consortia were unassigned, we developed a custom pipeline for identification, extraction, and assignment of these contigs to chromosome arms.

To assign contigs to chromosome arms, we mapped each assembly to five human reference genomes (CHM13, HG002-maternal, HG002-paternal, CN1-maternal, and CN1-paternal)^1,2,76,77^ with minimap2 (v.2.28-r1209)^78^ with parameters: -x asm5. For every assembly-reference mapping, we noted the contig with the largest overlap with the terminal 1,000 kb region of each chromosome arm. If the same contig was chosen for at least four out of the five reference genomes, then that contig was assigned to that chromosome arm.

To ensure consistent, comparable subtelomere contigs, we extracted sequences from the assembly based on the mapped coordinates, removed sequencing artifacts on the telomeric edge, trimmed the telomeric repeats to 2kb, and adjusted the subtelomeric region on the centromeric edge to 1,000kb.

### Assessment of contig and assembly quality

To evaluate the quality of the assemblies, we developed a classification pipeline to categorize contigs by how reliably they can be assigned to specific chromosome ends. Each contig was assigned a quality "flag" based on its relationship to the reference chromosomes and sample identity:

- **Telomere-to-Telomere** (T-to-T): Contigs were flagged as "T-to-T" if they were assigned to both the p- and q-arms of the same chromosome.
- **Duplicated**: Contigs were flagged as "duplicated" in instances where a single contig yielded assignments across multiple distinct chromosomes, suggesting that these contigs could not be uniquely assigned to a specific chromosome end.
- **Arm-Unique**: Contigs were classified as “arm-unique” if it was assigned to only a single chromosome end.

The classification was performed using a custom R script utilizing the dplyr (v.1.2.1)^79^ library to group data by sample and chromosome ID, ensuring that structural features were assessed on a per-sample basis.

### Selection of a non-overlapping collection of subtelomeric sequences

Where a sample had multiple assemblies, we selected a single representative by ranking assemblies on three criteria in order: (1) most T2T contigs; (2) most arm-unique contigs; and (3) fewest duplicated contigs. Assemblies were ranked first by T2T count, ties broken by arm-unique count, and remaining ties by fewest duplicated contigs; the top-ranked assembly was retained. Processing was performed in R using dplyr^79^ and tidyr (v.1.3.2)^80^, and results were exported as tab-separated (TSV) files for downstream analysis.

### Sex Chromosome Assignment and Validation

Contigs assigned to chrX or chrY were aligned to CHM13 chrX/chrY references using minimap2^78^ with parameters -x asm5, retaining primary alignments covering ≥20% of contig length. Alignment fraction (summed nmatch [number of matching bases in alignment] / contig length) was used with reported biological sex and chromosome length filters (chrX < 160 Mb, chrY < 70 Mb) to assign each contig: chrX fraction >0.5 for females, and a >0.05 fraction difference between chrX and chrY for males. For each unassigned contig, we recomputed an alignment score (nmatch / contig length, using only the top-scoring alignment) against chrX and chrY separately, and assigned the contig to the higher-scoring chromosome if the two scores differed by more than 0.2.

Assignments were cross-checked across both haplotypes per individual, using the resolved haplotype to reassign ambiguous contigs in the other (alignment fraction ≥ 0.98) where possible; samples with both haplotypes unresolved were excluded. Assignments were validated against four marker genes with known arm-specific localisation: *XIST* (NCBI Gene ID: 7503, Xq), *KAL1* (NCBI Gene ID: 3730, Xp), *SRY* (NCBI Gene ID: 6736, Yp), and *CDY1* (NCBI Gene ID: 9085, Yq), requiring ≥ 95% alignment coverage of gene length.

### Comparison of hifiasm and Verkko Assemblies

Assembled contigs from matched Verkko^17^ and hifiasm^13^ samples were evaluated for concordance in their assembled sequences. As both assemblers output two haplotypes, and the correspondence between the hifiasm and Verkko haplotypes is not known *a priori*, haplotype matching was performed computationally to determine the correct pairing of haplotypes generated by hifiasm and Verkko for each sample, and contigs assigned to each chromosome arm.

Specifically, each hifiasm-generated haplotype contig assigned to a given chromosomal arm was aligned to both Verkko-generated haplotype contigs assigned to the same chromosomal arm using minimap2 (version 2.28-r1209; -x asm5, --secondary=no, and -c)^78^. The alignments were concatenated and gap-compressed identity per unique contig-contig mapping was calculated.

The gap-compressed identity is defined as^81^:

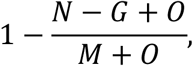

where **N** is the edit distance, **M** is the total number of aligned columns, **G** is the total number of gap bases, and **O** is the number of gap opens, such that each contiguous gap is counted as a single difference irrespective of its length.

Each hifiasm contig was then assigned to the Verkko haplotype contig with the highest gap-compressed identity. The number of assembled contigs per chromosomal arm was counted per sample based on these assignments; contigs that could not be assigned to any chromosomal arm were flagged as unassigned.

### Sequence divergence from CHM13 reference

To quantify sequence divergence between subtelomeric assemblies and the CHM13 reference, k-mer based distance estimation was performed using Mash (v.2.3)^44^. For each chromosome arm, a reference sketch using the terminal 500kb sequences of CHM13 (-k 21 -s 10,000), and individual sketches for every contig with an assigned chromosome arm were generated. Pairwise Mash distances between each contig and its corresponding CHM13 arm were then computed, yielding an estimated sequence divergence for each contig.

Each contig’s Mash distance was subsequently annotated with its major haplotype assignment (S1-S7) obtained from the paralogy block clustering results for the corresponding arm. Contigs belonging to arms with only a single detected haplotype were assigned to a single haplotype (S1). Contigs that were assigned to rare major haplotypes (n < 5 contigs) were excluded from the analysis.

### Grouping of subtelomeric contigs

Highly-conserved regions of paralogy blocks were identified in the subtelomeric contigs of each chromosome arm. Paralogy block reference sequences from Stong *et al*.^23^ were mapped to the subtelomere sequences using minimap2 (v.2.28-r1209; -x asm5)^78^. To identify the sequence of paralogy blocks in each subtelomere sample, all alignments that passed the following filters were extracted from the mapping results: primary alignment, more than 1,000 matching and total bases, greater than 40% of the length of the reference block sequence.

The block sequence of each subtelomere sample was converted to a string. Each character of the string encoded a single paralogy block and its strand. All such block sequence strings for a single chromosome arm were subjected to multiple sequence alignment by mafft (v.7.149b)^82^ with parameters: --text --op 0. Hamming distance was calculated between the block sequence strings in each multiple sequence alignment. Distance between two paralogy blocks of a different type or any paralogy block and a gap was counted as one.

In the first round of grouping, the resultant distance matrix was used to perform agglomerative clustering using sklearn (v.1.6.1)^83^ with complete linkage and a distance threshold of three, with blocks smaller than 1kbp being ignored. This grouped all sequences within a chromosome arm into “major” clusters. A second round of grouping was then performed within major clusters using the same distance matrix, agglomerative clustering, complete linkage, a distance threshold of one, and with all blocks considered. This further grouped sequences into “minor” clusters. The minor cluster with the highest population frequency was designated as the major haplotype for that cluster.

During clustering of block sequences from chromosome arms 4q and 10q, chains of Block 40 were treated as a single block to account for variation caused by the D4Z4 tandem repeat array. Within each sub-cluster, all subtelomere contigs were mapped against each other using minimap2 (v.2.28-r1209)^78^ with parameters: -x asm5 -X -c. Contigs were ranked by the average alignment score of all of their highest-scoring mappings to every other contig. The contig with the highest average alignment score was assigned as the representative for the entire sub-cluster.

All major haplotypes were ordered according to the length of the entire sequence of paralogy blocks, from the start of the first block to the end of the last block (e.g. S1 is the longest, S2 is the second-longest). Sub-clusters under that major haplotype were designated as minor haplotypes and also ordered by length (e.g. S1.m1 is the longest).

### Subtelomere haplotype frequency and zygosity

Subtelomeric haplotypes were analysed across a total of 431 samples categorised into six different superpopulations (AFR, AMR, EAS, EUR, MENA, SAS). Population metadata was obtained from the respective consortia from which the assemblies were derived^5–8^. Sample locations were visualised on a world map using the maps (v.3.4.3)^84^ and mapdata (v.2.3.1)^85^ packages in R, with subpopulations plotted at approximate geographic coordinates.

Contigs were sorted into major haplotype labels for each chromosome arm. Contigs not assignable to a major cluster were labelled unassigned and excluded from downstream analyses, as were samples with unknown ancestry metadata. For each chromosome arm with more than one major haplotype (n = 21 arms), haplotype frequency was calculated within each superpopulation and across all samples, as the proportion of haplotype assignments per group.

Fisher’s exact tests with Monte Carlo simulation (10,000 permutations, seed = 42) were used to test for differential haplotype distribution between ancestries, comparing each superpopulation against all others combined for each arm, using the fisher.test function in R. P-values were corrected using Benjamini-Hochberg within each arm, across the six superpopulation tests performed. Bias-corrected Cramér’s V was computed for each arm-ancestry combination using the cramerV function in the rcompanion package (v.2.5.2)^86^ to quantify effect size. Arm-ancestry combinations were considered to show strong ancestry-associated enrichment at BH-adjusted p < 0.05 and corrected Cramér’s V > 0.3 (moderate effect). Zygosity was determined per chromosome arm for each diploid individual by comparing haplotype assignments across both haplotypes. Individuals with only one haplotype present or assignable were excluded.

### Population-level subtelomere block analysis

Average subtelomeric block copy number was calculated per individual across the HPRC-Y2 cohort by summing block occurrences across both haplotypes. For each block, pairwise distances between 1,000 subsampled sequences using SeqKit (v.2.10.0)^87^ were computed using Mash (v.2.3)^44^ with parameters: -s 10,000 -k 21, used for hierarchical clustering with ComplexHeatmap (v.2.18.0)^88^ and variance partitioning by ancestry and subtelomere haplotype using vegan (v.2.7-3)^89^. Inter- and intra-arm Mash distances were compared to assess block similarity within versus between chromosome arms. Alluvial plots depicting relationships between clustering, chromosome arm, subtelomere haplotype, and ancestry were generated using the ggalluvial^90^ package in R (v.0.12.6).

### Subtelomere block analysis and construction of phylogenetic tree in great apes

The terminal 500kb of each chromosome arm was extracted from ape assemblies^45^ for five species: western gorilla (*Gorilla gorilla,* GCA_029281585.2), bonobo (*Pan paniscus,* GCF_029289425.2), chimpanzee (*Pan troglodytes,* GCA_028858775.2), Sumatran orangutan (*Pongo abelii,* GCA_028885655.2), and Bornean orangutan (*Pongo pygmaeus,* GCA_028885625.2). To account for inter-species sequence divergence, Stong *et al*. 2014^23^ subtelomeric reference blocks were aligned to these terminal 500kb sequences using minimap2 (v.2.28-r1209)^78^ with parameters: -ax asm20, retaining primary alignments with nmatch/block length > 0.4. Block presence per species was visualised as a binary heatmap and clustered into four row-wise clusters using ComplexHeatmap (v.2.18.0). Copy number per block per species was computed as the number of primary alignments per block.

Blocks from representative human samples and apes were concatenated. Multiple sequence alignment was performed using mafft (v.7.149b)^82^ with parameters: --auto. Each block tree was plotted using iqtree3 (v.3.1.2)^91^ with parameters: -m TEST -B 1,000 and visualised using ggtree (v.4.0.4)^92^.

### Subtelomeric gene analysis

Terminal 700 kb intervals for each chromosome were defined from CHM13v2.0. CHM13 annotation, taken from HPRC consortium (https://github.com/human-pangenomics/hprc_intermediate_assembly/tree/main/data_tables/annotation) were intersected against these intervals using bedtools intersect (v.2.13.0)^93^ with parameters: -a, to generate a list of overlapping genes. This list was then used to subset the annotation with agat_sp_filter_feature_from_keep_list.pl from AGAT (v.1.7.0)^94^. Subtelomeric gene annotations were lifted from the annotation onto the assemblies and Stong et al.’s reference block using Liftoff (v.1.6.3)^47^ with parameters: -copies -polish and -sc 0.97. Only protein-coding genes with valid ORFs > 0, coverage > 0.95 and sequence identity > 0.95 were retained for analysis. As subtelomeric boundaries differ between individuals, only genes present in at least 50% of the chromosome arm’s haplotype for all haplotypes are retained. The gene is then ordered from telomeric end to centromeric end based on the haplotype which contains the largest proportion of genes. If none exists, all genes are plotted.

For each gene, we first identified all chromosome arms on which it was detected. We then defined a gene-specific cohort pool comprising only individuals with complete subtelomeric contigs for both haplotypes across all arms where that gene was observed. Copy number was calculated independently within each gene’s sample pool using Liftoff annotations meeting the quality thresholds described above and normalised against the pool size for that gene. This per-gene pooling strategy ensures that copy number estimates reflect only individuals with sufficient assembly completeness at the relevant loci, and accounts for the fact that different genes may be assessable in different subsets of individuals. Ggsankey (v.0.0.9999)^95^ was utilised to visualise the distribution of contigs across the different copy number categories as well as the reasons why Liftoff did not detect the gene.

### Rescue of 4q and 10q contigs

Assemblies with contigs unassignable to 4q or 10q due to high sequence similarity underwent a rescue procedure to recover contigs of sufficient length and quality for haplotype assessment. Contigs with truncated D4Z4 arrays were filtered out by dividing them into 300 bp chunks and aligning to 4q-S1, 4q-S2, 10q-S1, and 10q-S2 reference haplotypes using minimap2 (v.2.28-r1209)^78^ with parameters: -x asm5. Contigs were called truncated if ≥80% of chunks at either end showed similar alignment scores across all four references, and excluded from rescue.

Untruncated contigs were assigned to 4q or 10q based on their best scoring reference alignment (nmatch > 250,000bp). Telomere length of the rescued contigs was estimated using estimateTelSize^96^ from Teltools (https://github.com/ktan8/teltools) and were trimmed to retain 2kb of telomeric sequence and a maximum length of 500kb, consistent with other chromosome arms. Haplotypes were assigned through alignment to Stong *et al*.^23^ paralogy block references: S1 (Block 23, Block 24, Block 41, Block 33) and S2 (Block 39, Block 33). Contigs with duplicated blocks other than Block 40 were excluded. To verify arm assignments, the sequence distal to the last paralogy block was aligned to CHM13. All rescued contigs aligned concordantly with their new assigned arm. The rescue procedure recovered 28 and 24 contigs for 4q and 10q, respectively.

### Analysis of 4q and 10q subtelomeres

Assemblies containing more than two separate D4Z4 arrays at distinct loci were excluded from downstream analysis. D4Z4 repeats were first classified by locus based on the presence or absence of the BlnI site (CCTAGG) and XapI site (RAATTY), with BlnI-positive/XapI-negative assigned to 10q, XapI-positive/BlnI-negative assigned to 4q, and arrays containing both or neither sites classified as hybrid. To further characterise D4Z4 repeats, Mash (v.2.3)^44^ sketches were computed for each assembly with parameters: -s 10,000 -k 21, and pairwise Mash distances were used to construct a distance matrix for principal coordinates analysis (PCoA). *DUX4* alleles were classified by extracting the 3’ end of the terminal paralogy Block 40 from each assembly, performing multiple sequence alignment with ClustalO (v.1.2.4)^97^ against *DUX4S* (Ensembl ENST00000569241.5) and *DUX4L* (GenBank MF693913.1) references, and confirming presence of the PAS sequence (ATTAAA) within the pLAM region.

Interchromosomal translocation between 4q and 10q was assessed by dividing contigs into 300 bp windows and aligning each independently to 4q and 10q pangenome reference haplotypes using minimap2 (v.2.28-r1209)^78^ with parameters: -x asm5. Alignment score difference (S1_10q_ – S1_4q_) was calculated per window using the highest-scoring primary alignment, smoothed with a 50-window rolling mean, and plotted along contig coordinates. Contigs showing an abrupt transition in this signal were interpreted as carrying a translocation junction.

### Subtelomere read simulation and mapping

To assess the accuracy of mapping of telomeric and subtelomeric reads to chromosome arms, long-read sequences were simulated from subtelomeric reference sequences. Specifically, sample-specific subtelomeric reference sequences were generated from assembled subtelomeric contigs assigned to each chromosome arm. Simulated long-read datasets were then generated from these references using PBSIM3 (v.3.0.5)^98^ under multiple sequencing scenarios.

a. ONT high-quality reads: simulated using the QSHMM-ONT-HQ model at 15x depth, with a mean read length of 20 kb and read accuracies of 90%, 95%, and 100%;
b. ONT reads at varying depth: simulated using the QSHMM-ONT-HQ model at 5x, 10x, and 15x depth, with a mean read length of 20 kb and 95% accuracy;
c. ONT reads at varying read length: simulated using the QSHMM-ONT-HQ model at 15x depth, with mean read lengths of 10 kb, 15 kb, and 20 kb at 95% accuracy;
d. PacBio CLR reads: simulated using the QSHMM-RSII model at 15x depth, a mean read length of 20 kb, and 95% accuracy. Simulated CLR reads were subsequently collapsed into circular consensus sequences using the CCS software^99^ (Pacific Biosciences).

Simulated subtelomeric reads containing telomeric sequence were isolated by retaining reads overlapping regions containing at least 1 kb of telomeric (TTAGGG)n repeats. Two read sets were therefore analysed for each simulation condition: (1) all simulated subtelomeric reads and (2) a telomeric only subset.

A subtelomeric pangenome reference was constructed by concatenating 50kb subtelomeric sequences from samples passing quality-control criteria (see Methods above). Because the pangenome reference was derived from the same cohort as the simulated reads, a leave-one-out strategy was employed, whereby each donor’s subtelomeric sequence was excluded from the pangenome prior to alignment of that donor’s simulated reads.

Simulated reads were mapped to three reference datasets: CHM13^1^, the subtelomeric pangenome reference, and the donor’s own assembly, using minimap2 (v.2.28-r1209)^78^ with parameters: -ax map-ont for ONT reads and -ax map-hifi for PacBio HiFi reads. Mapping accuracy was defined as the proportion of reads assigned to the correct chromosome arm out of all simulated reads. The true arm of origin for each read was known from its simulation source coordinates. Reads were assigned to chromosome arms using reference annotations, and a read was considered correctly mapped if its assigned arm matched its arm of origin.

### Telomere length analysis

For each individual in HPRC2 (n = 128), telomere length data were obtained using Teltools^96^. Reads were retained only if classified as either left-telomeric or right-telomeric, excluding non-telomeric and intra-telomeric reads. A sample-wide mean telomere length was calculated from this filtered read set to serve as a normalization baseline^40^. Chromosomal assignments for each read were obtained through alignment to its matched reference genome assembly using minimap2 (v.2.28-r1209)^78^ with parameters: -ax map-hifi preset for PacBio HiFi reads. Only primary alignments (PAF tag: *tp:A:P*) were retained to assign each read to a single chromosomal location. Reads were stratified at two levels: by chromosome arm, and by chromosome arm further partitioned into the major haplotypes defined in Figure 2.

At each level, groups with fewer than five supporting reads were excluded to avoid unreliable estimates. For each remaining group, a relative mean telomere length was computed as the difference between the group-specific mean and the sample-wide mean, normalizing for inter-sample variation in overall telomere length. Acrocentric short arms (13p, 14p, 15p, 21p, and 22p), which lack canonical telomeric sequence and are not reliably assembled, were excluded from both stratifications. For the haplotype-level stratification, relative mean telomere lengths were compared within major haplotype groups using the Kruskal-Wallis H-test from scipy (v.1.15.2)^100^, a non-parametric method suitable for comparing more than two independent groups without assuming normality of the underlying distributions. P-values were adjusted for multiple testing using the Benjamini-Hochberg procedure.

Separately, to assess evenness of coverage across chromosome arms using the pangenome-mapping approach for the real long-reads, left- and right-telomeric flanks were aligned using minimap2^78^ with parameters: -cx map-hifi against a 20 kb subtelomeric pangenome reference panel for each chromosome arm, comprising 2 kb telomeric sequence and 18 kb subtelomeric flank sequence. The sample’s own reference sequences were excluded from the reference panel to avoid self-mapping bias. Alignments with mapping quality ≥30, identity ≥90%, and query coverage ≥20% were retained. For each chromosome arm, best-scoring alignments meeting these criteria were counted, including flanks with equally scoring best alignments to multiple arms. For each sample, each arm’s fraction was calculated as its flank count divided by the sample’s total mapped flanks.

Chromosome arm identity for each read was first defined by mapping to sample-specific self-assembled contigs. Reads were then separately mapped to the CHM13 and pangenome references, and concordance with the self-assembly-based arm was determined for each method. Per-sample, per-arm accuracy and read-assignment confusion matrices were computed and visualized in R.

## Acknowledgements

We thank all contributing consortia for sharing assemblies, and all members of the K.T.T lab for feedback and discussion. Work in the K.T.T. lab was supported by the National Medical Research Council, Singapore (OFYIRG24jul-0039); the Ministry of Education, Singapore (Tier 1 grant A-8002917-00-00 and Academic Research Tier 3 grant MOE-MOET32021-0004); the National Research Foundation, Singapore (NRF-CRP32-2024-0005); and internal grants from the National University of Singapore (ODPRT startup grant and NUS Medicine grant A-0010093-00-00). Computational work was performed in part using resources provided by the National Supercomputing Centre (NSCC) Singapore and the Bioinformatics Core Facility, Yong Loo Lin School of Medicine, National University of Singapore. M.M. was supported by an NIH Outstanding Investigator Award (R35CA197568) and the American Cancer Society Research Professorship.

## Author contributions

K.T.T. conceived and initiated the study. K.T.T., R.J.X.O., B.B.R.K., R.K.H.Y., M.M., and H.L. designed the study and analyses. K.T.T., R.J.X.O., B.B.R.K., R.K.H.Y., A.J.T.N., Q.L., G.K.R., J.C.R., and C.T.K. collected and curated the data. R.K.H.Y. developed the haplotype definition algorithm. R.J.X.O. performed the 4q/10q and ancestry analyses. B.B.R.K. performed the evolutionary analyses. B.B.R.K., A.J.T.N., M.G.J. and G.K.R. performed the telomere length and mapping analyses. B.B.R.K., R.J.X.O., and J.C.R. performed the gene analyses. K.T.T., R.J.X.O., B.B.R.K., R.K.H.Y., and A.J.T.N. wrote the initial draft of the manuscript. M.M. and H.L. reviewed and revised the manuscript. K.T.T., M.M., and H.L. jointly supervised the work. All authors read and approved the final manuscript.

## Data Availability

Pangenome reference sequences of subtelomeres are available at Zenodo (https://zenodo.org/records/21807156).

## Code Availability

The software pipeline used in this analysis is available at https://github.com/ktanlab/subtelomere/

## Ethics Statement

This study analyzed only publicly available, de-identified human genome assemblies generated and released by existing consortia (including HPRC and HGSVC) under their own ethical approvals and data-use agreements. As no new samples or identifiable data were collected, no additional ethical approval was required.

## Materials & Correspondence

Correspondence to Kar-Tong Tan or Heng Li

---

**Extended Data Figure 1.**
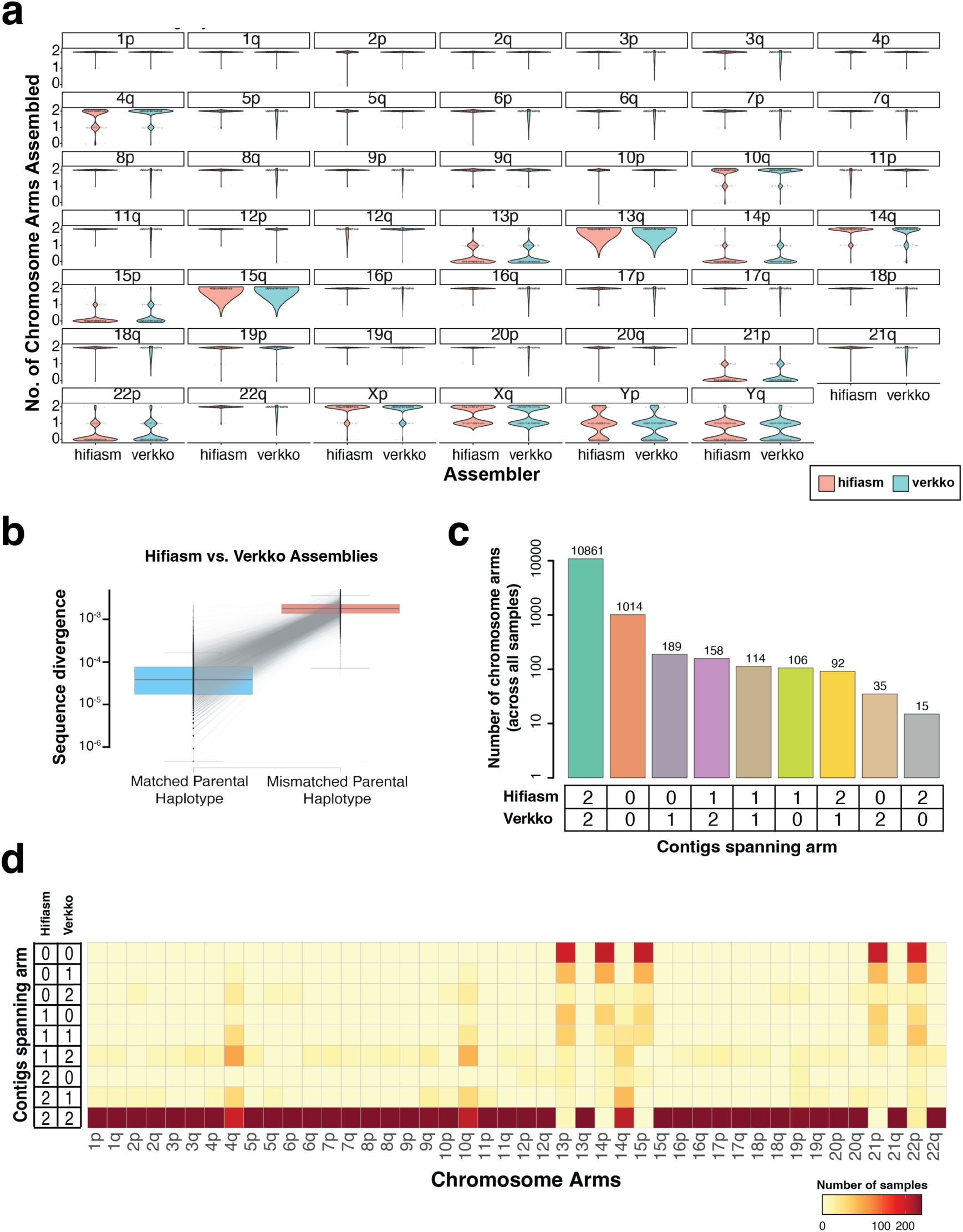
Concordance between hifiasm and Verkko assemblies in HPRC2. (**a**) Violin plot of assembled contigs for each chromosomal arm for both hifiasm (red) and Verkko (blue). (**b**) Boxplot of sequence divergence score calculated using the gap-compressed identity between matched parental haplotype vs mismatched parental haplotype. Dotted red line indicates 0. (**c**) Barplot of the number of chromosomal arms per (hifiasm, verkko) contig count combination, aggregated across all samples. (**d**) Heatmap of sample counts per chromosomal arm (x-axis) for each hifiasm–verkko contig count combination (y-axis), where each row represents a (hifiasm, verkko) pair ranging from (0,0) to (2,2). Colour scaled on a square root transformation.

**Extended Data Figure 2.**
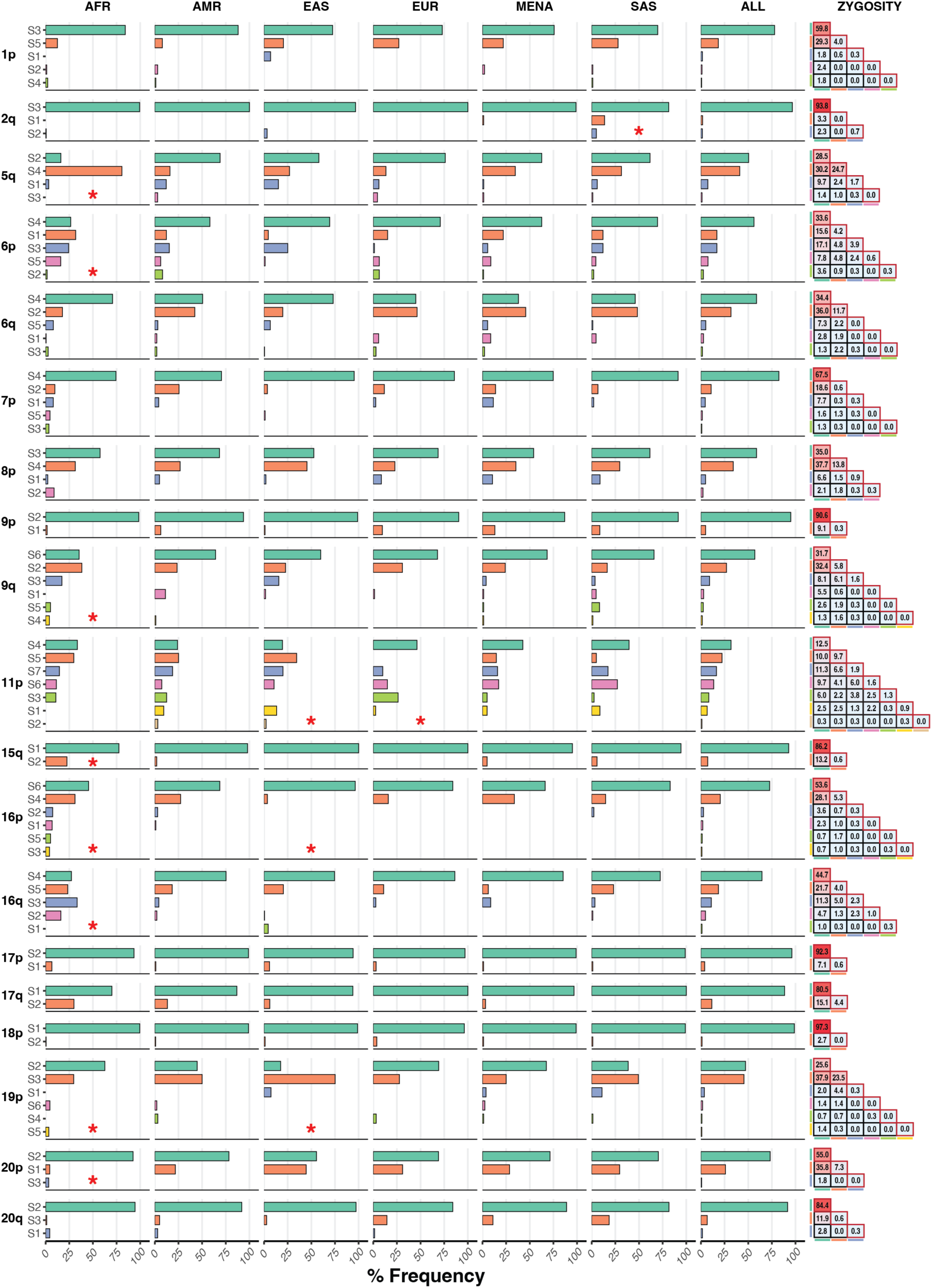
Subtelomeric haplotype frequencies for all chromosomal arms exhibiting large structural variations. Panel interpretation follows that of Figure 3.

**Extended Data Figure 3.**
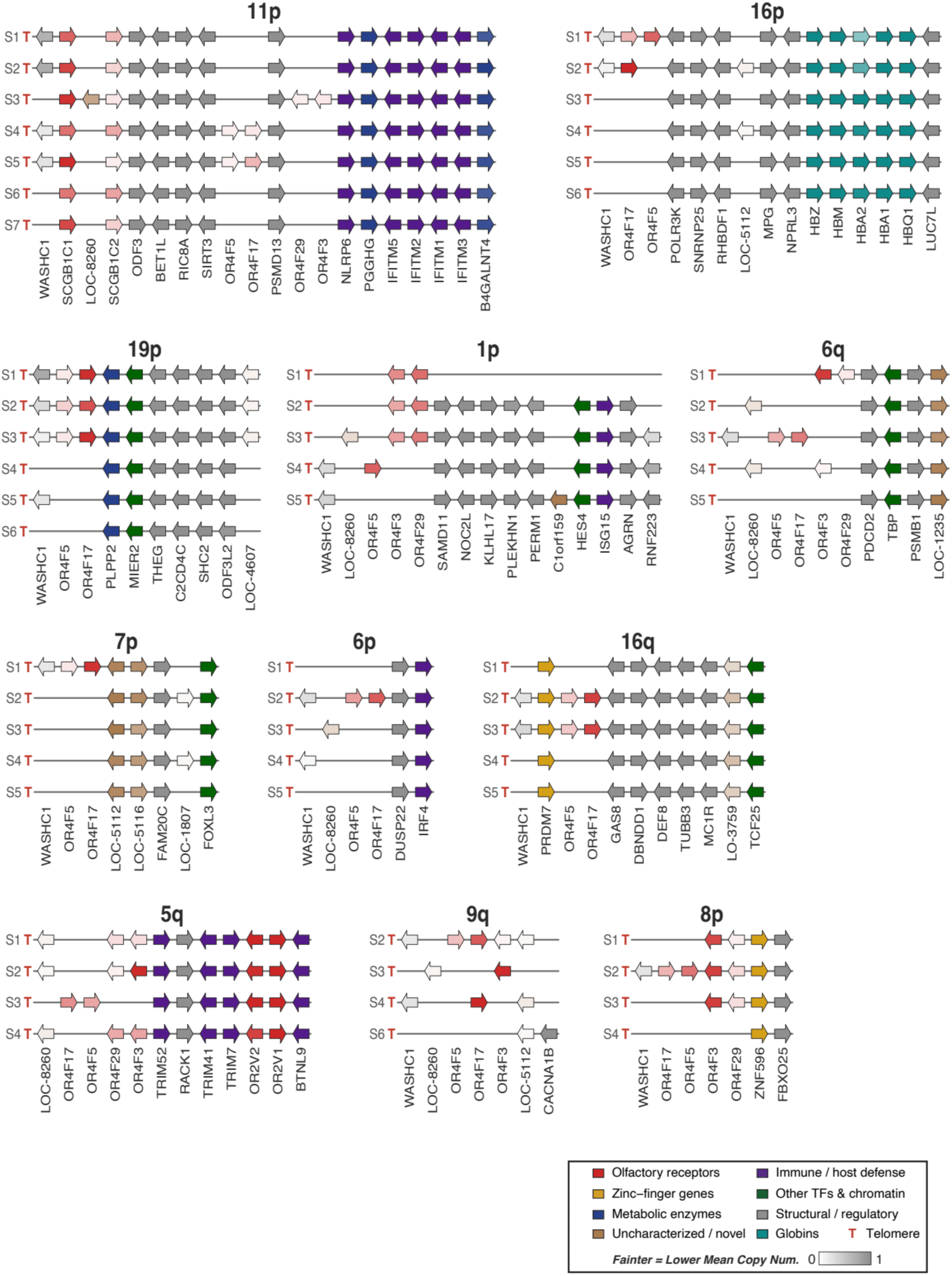
Subtelomeric gene content and copy number differ across haplotypes and across population (I). Gene-arrow diagrams across haplotypes (S1-S7, rows) for chromosome arms (11p, 16p, 19p, 1p, 6q, 7p, 6p, 16q, 5q, 9q, 8p). Genes are represented as arrows ordered along the panel from telomeric ("T") to centromeric, with arrow orientation indicating the direction of transcription. Arrows are coloured by functional category, with colour intensity indicating the mean copy number of across all instances of the corresponding haplotype in the pangenome collection, from absent (white) to one copy (full colour). Gene names beginning with "LOC" followed by a long numeric identifier (NCBI Gene IDs) have been abbreviated to "LOC-" followed by the last four digits for clarity: LOC112268260 (LOC-8260), LOC124900641 (LOC-0641), LOC124900642 (LOC-0642), LOC124901235 (LOC-1235), LOC105375112 (LOC-5112), LOC105375116 (LOC-5116), LOC124901807 (LOC-1807), LOC124901794 (LOC-1794), LOC124902106 (LOC-2106), LOC112268317 (LOC-8317), LOC105379417 (LOC-9417), LOC124903759 (LOC-3759), LOC124904094 (LOC-4094), LOC124904095 (LOC-4095), LOC124904607 (LOC-4607), LOC101929937 (LOC-9937).

**Extended Data Figure 4.**
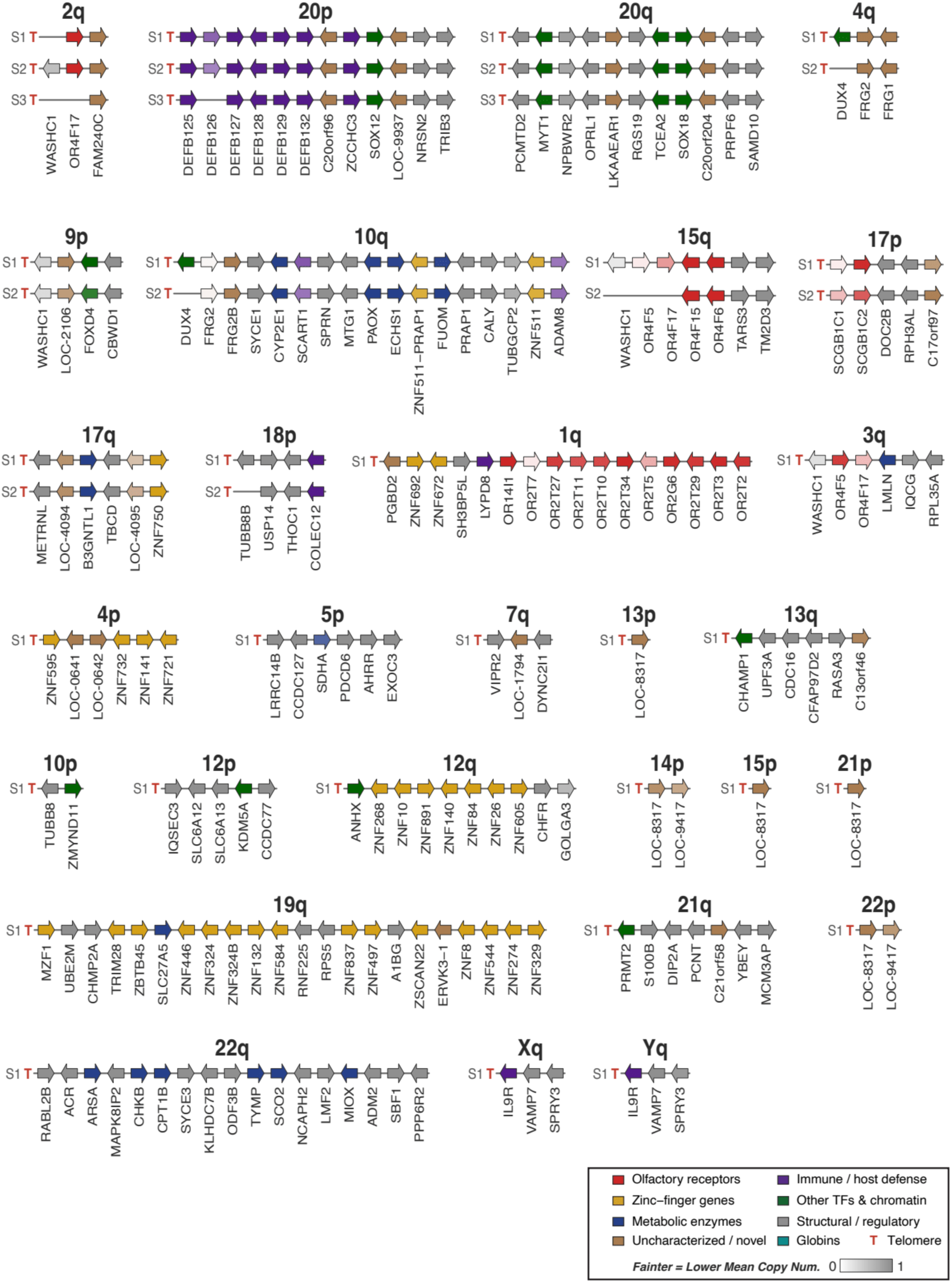
Subtelomeric gene content and copy number differ across haplotypes and across population (II). Gene-arrow diagrams across haplotypes (S1-S7, rows) for all chromosome arms (2q, 20p, 20q, 4q, 9p, 10q, 15q, 17p, 17q, 18p, 1q, 3q, 4p, 5p, 7q, 13p, 13q, 10p, 12p, 12q, 14p, 15p, 21p, 19q, 21q, 22p, 22q, Xq, Yq). Panel interpretation follows that of Extended Data Figure 3.

**Extended Data Figure 5.**
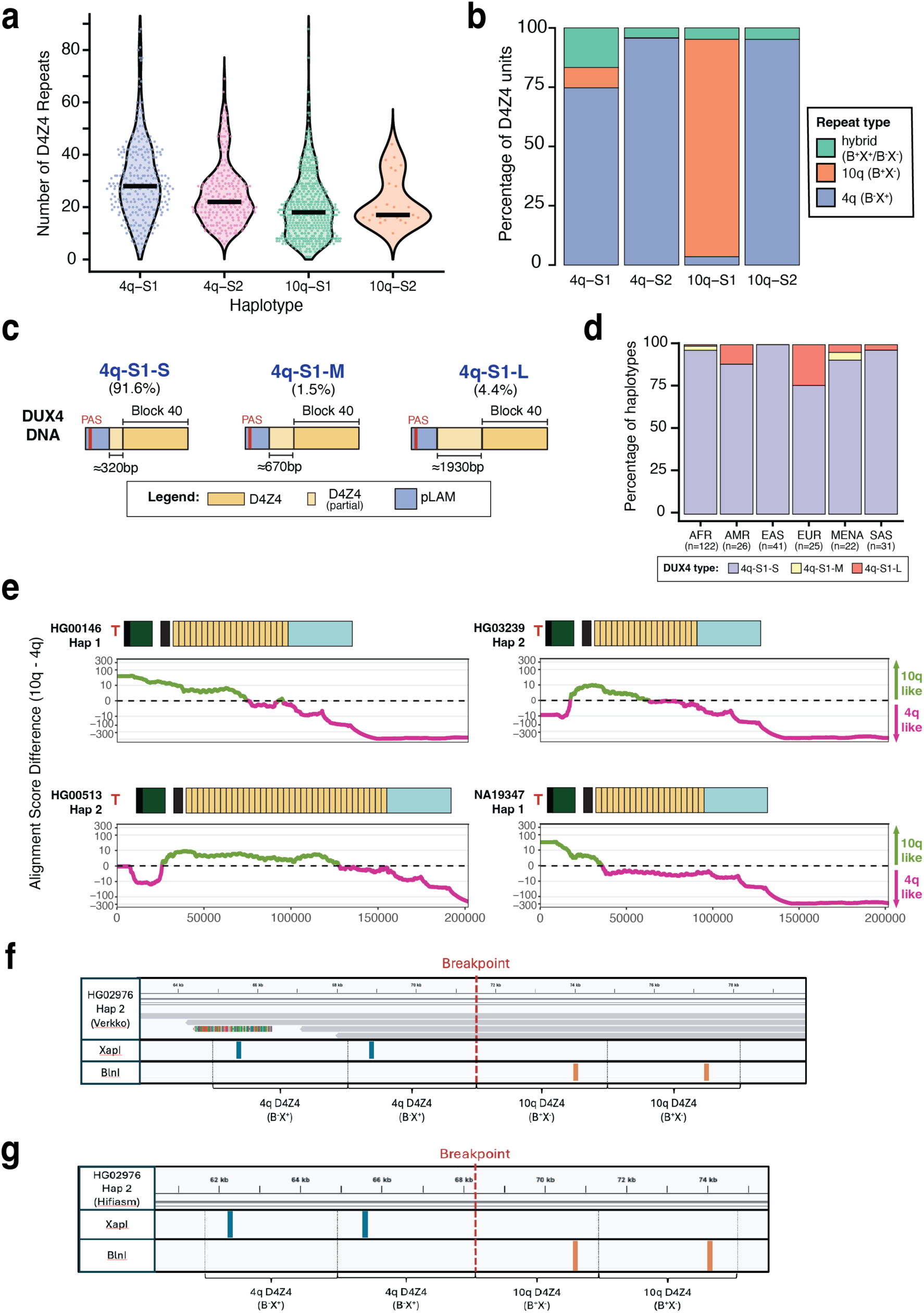
D4Z4 repeat array characteristics, DUX4 structural variation, and validation of translocation. **(a)** Distribution of D4Z4 repeat unit copy number per contig for the 4q-S1, 4q-S2, 10q-S1, and 10q-S2 haplotype clusters; horizontal bars indicate medians. **(b)** Proportional composition of D4Z4 repeat unit types, 4q-type (B-X+), 10q-type (B+X-), and hybrid (B+X+/B-X-), within each cluster. **(c)** Schematic of the distal D4Z4 locus encoding DUX4 on the permissive 4q-S1 allele, showing three structural variants: short (4q-S1-S, 92%), intermediate (4q-S1-M, 1.5%), and long (4q-S1-L, 4.4%). **(d)** Ancestry-stratified frequency of DUX4 structural configurations across six superpopulations (AFR, AMR, EAS, EUR, MENA, SAS); the 4q-S1-L variant, previously considered European-specific, is present in other ancestries at lower frequency. **(e)** Alignment score difference (10q-like minus 4q-like) along four additional 4q assemblies carrying a 10q-S1 translocation, following the format of Fig. 6f. **(f,g)** Validation of the translocation breakpoint in HG02976 Hap2 by restriction-site profiling (XapI, blue; BlnI, orange) across the D4Z4 array, distinguishing 4q-type (XapI-sensitive, BlnI-resistant) from 10q-type (BlnI-sensitive, XapI-resistant) units. The dashed red line marks the inferred translocation breakpoint. **(f)** Verkko assembly, with HiFi reads spanning the transition confirming it is not an assembly artefact. **(g)** Independent hifiasm assembly of the same sample showing a concordant breakpoint, providing assembler-independent validation.

