## Supplementary Information for "Widespread structural variations at human chromosome ends"

### SUPPLEMENTARY NOTES

#### Supplementary Note 1: Cleaning Subtelomere Contigs

Inspecting telomeric edges unexpectedly revealed repeat artifacts (mostly <1 kb) distal to long telomeric repeat tracts (often >2 kb). As the telomeric tracts were too long to be interstitial (**Supplementary Fig. S1-S6**), we suspected that the repeat artifacts were caused by assembly or sequencing errors. First, ultra-long Nanopore reads are known to be more erroneous. In assemblies incorporating both ultra-long Nanopore reads and HiFi reads, these errors could have been incorporated into the chromosomal ends when there were few supporting HiFi reads, creating the repeat artifacts. Second, long reads from other parts of the genome can have short telomeric repeats, which could have been falsely assembled into the long telomeric repeats at the chromosomal ends and resulted in repeat artifacts. As these artifacts were likely to be sequencing or assembly errors, we chose to trim them away (**Methods**).

### SUPPLEMENTARY FIGURES

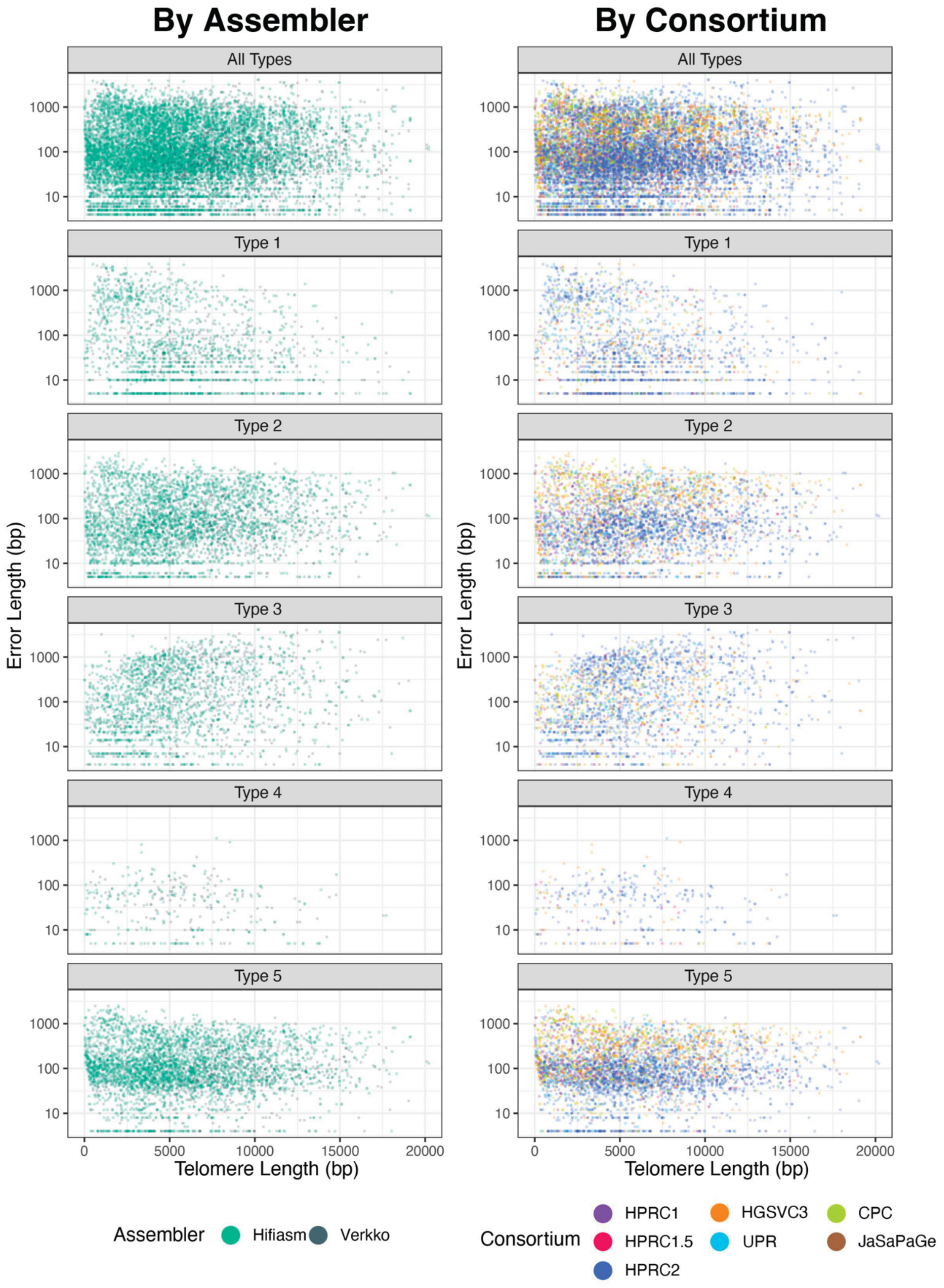


**Supplementary Fig. S1. Five types of edge repeat errors occur across contigs with different telomere length.** Each type of repeat error is represented by a different set of motifs. Points are coloured by either assembler or the consortium that sequence came from. No visible stratification by telomere length, assembler or consortium was observed.

**Type 1: TTAAAGTT, TTAAGTTA, AAGTTAA, AAGTTA, AAGTT, AAGTA, GTTAA**

```
> NA18747#1#HPRCY2-hifiasm_JBHDWE010000009.1:109566500-
110666499/rc#chr1:q-arm (1-7918)

TAATACAGTTAACTTTAAAGTTAAAGTTCAAGTTAAAGTTAAAGTTAAAGTTAAAGTTAAGTTAAAGTTACGTTTAAAGTTAAAGAGTTAAAGTTACAAGTTAAAGT
TAACGTTAAAGTTAAAGTTAAAGTTAACAGTTAAAGTTAAAGTTACAAGTTACAAGTTAAGTTAAAGTTAAAGTTAAAGTTAAGTTAACGTTAAAGTTAAGTGTAAG
TTAAAGTTAAAGTAAACGTTAAAGTTAAGTTACAGTTAAAGTTAAAGTAAAGTTAAAGTTAACGTTAAAGTTAAAGTTAAAGTTACAGTTAAGTTAAAGTTAAAGTT
AAGTTAAAGTTAAAGTTAAAGTTAAATTAAAGTTAAAGTTAAGTTAAGTTAAAGTTAAAGTTAAGTTAAAGTTACAGTTACGTTAAAGTTAACGTTAAAGTTAAAGT
TAAAGTTAAAGTTAAAGTTAAAGTTAAAGTTTAAGTTAAAGTTAAAGTTAAAGTTAAGTTAAGTTAAAGTTAAAGTTAACGTTAAAGTTAAAGTTAAGTAAAGTTAA
AGTTAAAGTTAAAGTTAAAGTTAAGTTAAAGTTAAAGTTCAAAGTCTAAAGTTAAAGTTAAAGTTAAGTTAAAGTTAAAGTTAAAGTTAAAGTTAAAGTTACAGTTA
AAGTTAAAGTTAAAGTTAACAGTTAAAAGTTACAGTTCAAGTTAAGTTAAAGTTAAAGTTAAAGTTAAAGTTAAGTTAAAGTTAAAGTTAACGTTAAAGTTAAAGTT
AAAGTTAAAGTTAAAGTTAAAGTTAAAGTTAAAGTTAAGTACAGTTAAAGTTAAAGTTAAAGTTAAAGTTAAAGTTAAAGTTAAAGTTAAGTTAAAGTTAAAGTTAA
AGTTAAAGTTAAAGTTAAAAGTTAAAGTTAAAGTTAAAGTTAAAGTAAAGTTAAGTTAAAGTTAAAGTTAAAGTTAAAGTTAAAGTTAAAGTTAAAGTTAAAGTTAA
AGTTACAGTTAAAGTTAAAGTTAAAGTTAAAGTTCAAGTTAAAGTTAAAGTTAAAGTTAAAGTTAAAGTTAAAGTTAAAGTTAAAGTTAAGTTAAAGTTAAAGTTAA
AGTTAAAGTTAAATTAAAGTTAAAGTTCAAGTTAAAGTAAAGTTAAAGTTAAAGTTAAAGTTAACGTTAAAGTTAAAGTTAAAGTTAAAGTTAAGTTAAAGTTAAAG
TTAAAGTTAAAGTTAAAGTTAAAGTTAAAGTTAAAGTTAAAGTTAAAGTTAAAGTTAAAGTTAAAGTTAAAGTTAAAGTTAAGTTAAAGTTAACAGTTACAAGTTAA
AGTTAAAGTTAAAGTTAAAGTTAAAGTTAAAGTTAAAGTTCAAGTTAAAGTTAAAGTTAAGTAAAAGTTAACGTTAAAGTTAAAGTTAAAGTTAAAGTTAAGTTAAA
GTTAAAGTTAAAGTAAAGTTAAAGTTAAGTAAAGTTAACGTTAAAAGTTAAAGTTAACGTTAAAGTTAAAGTTAAAGTTAAAGTTACAAGTTAAAGTTAAAGTTAAA
GTTAAAGTTAAAGTTAAAGTTAAAGTTAAGTTAAAGTTAAAGTTAAAGTTAAAGTACAGTTCCAAGTTAAAGTTAAAGTTAAAGTTAAAGTTAAAGTTAAGTTAAAG
TTAAAGTTAAAGTTAAAGTTAAAGTTAAAGTTAAAGTTAAAGTTAAAGTTAAAGTTAAAGTTAAAGTTAAACGTTAAAGTTAAAGTTAAAGTTAAAGTTAAAGTTAA
AGTTAAAGTTAAAGTTAAAGTTAAAGTTAAAGTTAAAGTTAAAGTTAAAGTTAACAGTTAAAGTTAAAGTTAAAGTTAAAGTCAAAGTTACAAGTTACAAGTTAACG
TTAAAGTTACAAGTTAAAGTTAAAGTTAAAGTTAAAGTTAAAGTTAACGTTAAAGTTAAAGTTAAAGTTAAAGTTAAAGTTAAGTTAACGTAAAGTTCACAGTTAAC
GTTAAAGTTAAAGTTAAAGTTAAAGTTAAGTTAAAGTTAAAGTTAAAGTTAAAGTTAAAGTTAAAGTTAAAGTTAACGTTAAAGTTAAAGTTAAAGTTAAAGTTAAC
GTTAAAGTTAAAGTTAAAGTTAAAGTTAAAGTTAAAGTTAAAGTAAAGTTAAAGTTAAAGTTAAAGTTAAAGTTACAAGTTAAAGTTACAAGTTA
ACGTTAAAGTTAAAGTTAAAGTTAAAGTTAAAGTTAAAGTTAAAGTTAAAGTTAAAGTTAAAGTTACAGTTAAAGTTAAAGTTAAAGTTAACGTTAAAGTTAAAGTT
AAAGTTAAAGTTAAAGTTAACGTTACAGTTAAAGTTAAAGTTAAAGTTAAAGTTAAGTTAAAGTTACAAGTTAAGTTAAAGTTAAAGTAAAGTTAAAGTACAGTTAA
AGTTAAAGTTAAAGTTAAAGTTAACGTTAAAGTTAAAGTTAAAGTTAAAGTTAAAGTTAAAGTTAAAGTTAAAGTTACAGTTAAAGTTAAAGTTAAAGTTAAAGTTA
AAGTTAAAGTTAAAGTTAAAGTTAAAGTTAAAGTTAAAGTTAAAGTTAAAGTTAAAGTTAAAGTTAAGTTAAAGTTAAAGTTAAAGTTAAAGTTCAAGTTCAAGTTA
AAGTTAAAGTTAAAGTTAAAGTTAAAGTTAAAGTTAAAGTTAAAGTTAAAGTTAAAGTTAAAGTTAAGTTAAAGTTAAAGTTAACGTTAAAGTTAAAGTTAAAGTTC
AAAGTTAAAGTAAAGTTAAAGTTAAAGTTAAAGTTCAAGTTAAAGTTAAAGTTAAAGTTAAAGTTAAAGTTAAAGTTACAGTTAAGTTAAAGTTAAAGTTAAGTTCA
AAGTTAAAGTTAAAGTTAAAGTTAAAGTTAAAGTTAAAGTTAAAGTTAAAGTTAAGTTAAAGTTAAAGTTAAAGTTAAAGTTAAAGTTAAAGTTAAAGTTAAAGTTA
ACGTTAAAGTTAAGTTAAAGTTAAAGTTAAAGTTAAAGTTACAGTTAAAGTTAAAAGTTAAAGTTAAAGAAAGTTAAAGTTAAAGTTACAGTTAAAGTTAAAGTTAA
GTTCAAGTTAAAGTTCAAGTTAAAGTAAAGTTAAAGTTAAAGTTAAAGTTAAAGTTAAAGTTACAGTTAAAGTTAACGTTAAAGTTAAAAGTTAAAGTTAAAGTTAA
AGTTAAAGTTAAAGTTAAAGTTAAAGTTAAAGTTAAAGTTAAAGTTAAAGTTAAAGTTAAAGTTAAAGTTAAAGTTAAAGTTAACGTTAAAGTTAAAGTTAAAGTTA
AAGTTAAAGTACAGTTAAAGTTCAAGTAAGTTAAAGTTAAAGTTAAGTTAAAGTTAAAGTTAAAGTTAAACGTTAAAGTTACAGTTACAGTAAAGTTAAAGTTCAAG
TTAAAGTTAAAGTTAAAGTTAACGTTAACGTTAAAGTTAAAGTTACAAGTTAAAGTTAAAGTTACAAGTTCAAGTTAAAAGTTAAAGTTAAGTTACAGTTAAAGTTA
AAGTTAAAGTTAAAGTTAAAGTTCAAGTTAAAGTTAACGTACCGTTAACGTACAAAGTTAAAGTTAAAGTTAAAGTTACAGTTAAAGTTAAAGTTAAAGTTAAAGTT
AAAGTTAACGTCTAAAGTTAACCGTTAAAGTTAAGTTAAAGTTAAGTTAACGTTAACGTTTAAAGTTAAAGTTAAAGTTACACGTTAAAGTTAAAGTAAGTTAACGT
TAAAGTTAAAAGTTAAAGTTAAAGTAAAGTTAAAGTTAAAGTTAAAGTTAAAGTTAAAGTTACAAGTTAAAGTTAAAGTTAAAGTTAAAGTTAAAGTTAAAAGTTAA
AGTTAAAGTTCAAGTTAACGTTAAAGTTAACGTAAAGTTAAAGTTAAAGTAAAGTTTAAAAGTTAAAGTTAAAGTTAAAAGTTAAGTTAGTTAGTAAAGTTAAGTTA
AAGTTAAAGTTCCAGTTAAAGTTAAAGTTTAAAGTTAAAGTTAAAGTTAACGTTAAAGTTAAAGTTCAAGTTAACGTTACAGTTAAAGTTAAAGTTACGTAAAGTTC
AAGTTCAAAGTTCACAGTTACAGTTCACAGTTCCAGTCACCCTTAACGTTAAAGTTCAAGTTAAAGTTCAAGTTAAGTAAGTTAAAGTTAAAGTTACACGTAAAGTT
ACAGTTAAAGTAAAGTTACAAGTTAACGTTAAAGTTAACGTAATACAGTTCAAGTTAAAGTTACCGTTAACGTTAAAGTTCAAGTTAACGTTACCAGTACCAGTTCA
AGTTACAAGTTACACCCGTTAACGTTAAGTTACGTTCAGTCCTAACGTTCAAGTTAACGTTAAAAGTTTAATAACGTTAACGTTCAAAGTTAAAAGTTCCCAGTTAC
ACGTAACACGTTCACGATCCCGTTAACTAACTACACTTAAATTAAAGTTCACGTTACCGTAACAGTAACGAATCCTAACCCTAACCCTAACCCTAACCCTAACCCTA
ACCCTAACCCTAACCCTAACCCTAACCCTAACCCTAACCCTAACCTAACCCTAACCCTACCCTAACCCTAACCCTAACCCTAACCCTAACCCTAACCCTAACCCTAA
CCCTAACCCTAACCCTAACCCTAACCCTAACCCTAACCCTAACCCTAACCCTAACCCTAACCCTAACCCTAACCCTAACCCTAACCCTAACCCTAACCCTAACCCTA
ACCCTAACCCTAACCCTAACCCTAACCCTAACCCTAACCCTAACCCTAACCCTAACCCTAACCCTAACCCTAACCCTAACCCTAACCCTAACCCTAACCCTAACCCT
AACCCTAACCCTAACCCTAACCCTAACCCTAACCCTAACCCTAACCCTAACCCTAACCCTAACCCTAACCCTAACCCTAACCCTAACCCTAACCCTAACCCTAACCC
TAACCCTAACCCTAACCCTAACCCTAACCCTAACCCTAACCCTAACCCTAACCCTAACCCTAACCCTAACCCTAACCCTAACCCTAACCCTAACCCTAACCCTAACC
CTAACCCTAACCCTAACCCTAACCCTAACCCTAACCCTAACCCTAACCCTAACCCTAACCCTAACCCTAACCCTAACCCTAACCCTAACCCTAACCCTAACCCTAAC
CCTAACCTAACCCTAACCCTAACCCTAACCCTAACCCTAACCCTAACCCTAACCCTAACCCTAACCCTAACCCTAACCCTAACCCTAACCCTAACCCTAACCCTAAC
CCTAACCCTAACCCTAACCCTAACCCTAACCCTAACCCTAACCCTAACCCTAACCCTAACCCTAACCCTAACCCTAACCCTAACCCTAACCCTAACCTAACCCTAAC
CCTAACCCTAACCCTAACCCTAACCCTAACCCTAACCCTAACCCTAACCCTAACCCTAACCCTAACCCTAACCCTAACCCTAACCCTAACCCTAACCCTAACCCTAA
CCCTAACCCTAACCCTAACCCTAACCCTAACCCTAACCCTAACCCTAACCCTAACCCTAACCCTAACCCTAACCCTAACCCTAACCCTAACCCTAACCCTAACCCTA
ACCCTAACCCTAACCCTAACCCTAACCCTAACCCTAACCTAACCCTAACCCTAACCCTAACCCTAACCCTAACCCTAACCCTAACCCTAACCCTAACCCTAACCCTA
ACCCTAACCCTAACCCTAACCCTAACCCTAACCCTAACCCTAACCTAACCCTAACCCTAACCCTAACCCTAACCCTAACCCTAACCCTAACCCTAACCCTAACCCTA
ACCCTAACCCTAACCCTAACCCTAACCCTAACCCTAACCCTAACCCTAACCCTAACCCTAACCCTAACCCTAACCCTAACCCTAACCCTAACCCTAACCCTAACCCT
AACCCTAACCCTAACCCTAACCCTAACCCTAACCCTAACCCTAACCCTAACCCTAACCCTAACCCTAACCCTAACCCTAACCCTAACCCTAACCCTAACCCTAACCC
TAACCCTAACCCTAACCCTAACCCTAACCCTAACCTAACCCTAACCCTAACCCTAACCCTAACCCTAACCCTAACCCTAACCCTAACCCTAACCCTAACCCTAACCC
TAACCCTAACCCTAACCCTAACCCTAACCCTAACCCTAACCCTAACCCTAACCCTAACCCTAACCCTAACCCTAACCCTAACCCTAACCCTAACCCTAACCCTAACC
CTAACCCTAACCCTAACCCTAACCCTAACCCTAACCCTAACCCTAACCCTAACCCTAACCCTAACCCTAACCCTAACCCTAACCTAACCCTAACCCTAACCCTAACC
CTAACCCTAACCCTAACCCTAACCCTAACCCTAACCCTAACCCTAACCCTAACCCTAACCCTAACCCTAACCCTAACCCTAACCCTAACCCTAACCCTAACCCTAAC
CCTAACCCTAACCCTAACCCTAACCCTAACCCTAACCTAACCCTAACCCTAACCCTAACCCTAACCCTAACCCTAACCCTAACCCTAACCCTAACCCTAACCCTAAC
CCTAACCTAACCCTAACCCTAACCCTACCCTAACCCTAACCCTAACCCTAACCCTAACCCTAACCCTAACCCTAACCCTAACCCTAACCCTAACCCTAACCCTAAC
CCTAACCCTAACCCTAACCCTAACCCTAACCCTAACCCTAACCCTAACCCTAACCCTAACCCTAACCCTAACCCTAACCCTAACCCTAACCCTAACCCTAACCCTAA
CCCTAACCCTAACCCTAACCCTAACCCTAACCCTAACCCCTAACCTAACCCTAACCCTAACCCTAACCCTAACCCTAACCCTAACCCTAACCCTAACCCTAACCCTA
ACCCTAACCCTAACCCTAACCCTAACCCTAACCCTAACCCTAACCCTAACCCTAACCCTAACCCTAACCCTAACCCTAACCCTAACCCTAACCCTAACCCTAACCCT
AACCCTAACCCTAACCCTAACCCTAACCCTAACCCCTAACCCTAACCCTAACCCTAACCCTAACCCTAACCCTAACCCTAACCCTAACCCTAACCCTAACCCTAACC
CTAACCCTAACCCTAACCCTAACCCTAACCCTAACCCTAACCCCTAACCCTAACCCTAACCCCTAACCCTAACCCTAACCCTAACCCTAACCCTAACCCTAACCCTA
ACCCTAACCCTAACCTAACCCTAACCCTAACCCTAACCCTAACCCTAACCCTAACCCTAACCCTAACCCTAACCCTAACCCTAACCCTAACCCTAACCCTAACCCTA
ACCCTAACCCTAACCCTAACCCTAACCCTAACCCTAACCCTAACCCTAACCCTAACCCTAACCCTAACCCTAACCCTAACCCTAACCCTAACCCTAACCCTAACCCT
AACCCTAACCCTAACCCTAACCCTAACCCTAACCCTAACCCTACCCTAGCCCTAGCCCTAGCCCTAGCCCTAGCCCTAGCCCTAGCCCTAGCCCTAGCCCT
AGCCCTAGCCCTAGCCCTAGCCCTAGCCCTAGCCCTAACCCCTAACCCCTAACCCCTAACCCCTAACCCTAACCCTAACCCTAACCCTAACCCTAACCCTAACCCTA
ACCCTAACCCTAACCCTAACCCTAACCCTAACCCTAACCCTAACCCTAACCCTAACCCTAACCCTAACCCCTAACCCTAACCCTAACCCTAACCCTAACCC
CAACCCCAACCCCAACCCCAACCCCTAACCCTAACCCTAACCCTAACCCTAACCCTAACCCTAACCCTAACCCTAACCCTAAAACCCTAACCCTAACCCTAACCCTA
ACCCTAACCCTAACCCTAACCCTAACCCTAACCCTAAACCCTAACCCTAAACCCTAACCCTAACCCTAACCCTAACCCTAACCCTAACCCTAACCCCTAACCCTAAC
CCTAACCCTAACCCTAACCCTAACCCTAACCCCTAACCCTAACCCTAACCCTAACCCAACCCTAACCCTAACCCTAACCCTAACCCTAACCCTAACCCTAACCCTAA
```

**Supplementary Fig. S2. Type 1 repeat errors and telomeric repeats in telomeric sequence.** Motifs are highlighted by whether they are suspected repeat errors or true telomeric repeats. Motifs that overlap are still highlighted.

**Type 2:** CCTTCCCT, CCTTCC, CCCCT, CCCTC, CCTCC, ACCCC, ACCCA

```
> NA20509#2#HGSVC3-hifiasm_h2tg000071-1-138897669:1-1100000/nc#chr9:p-arm (1-5775)
```

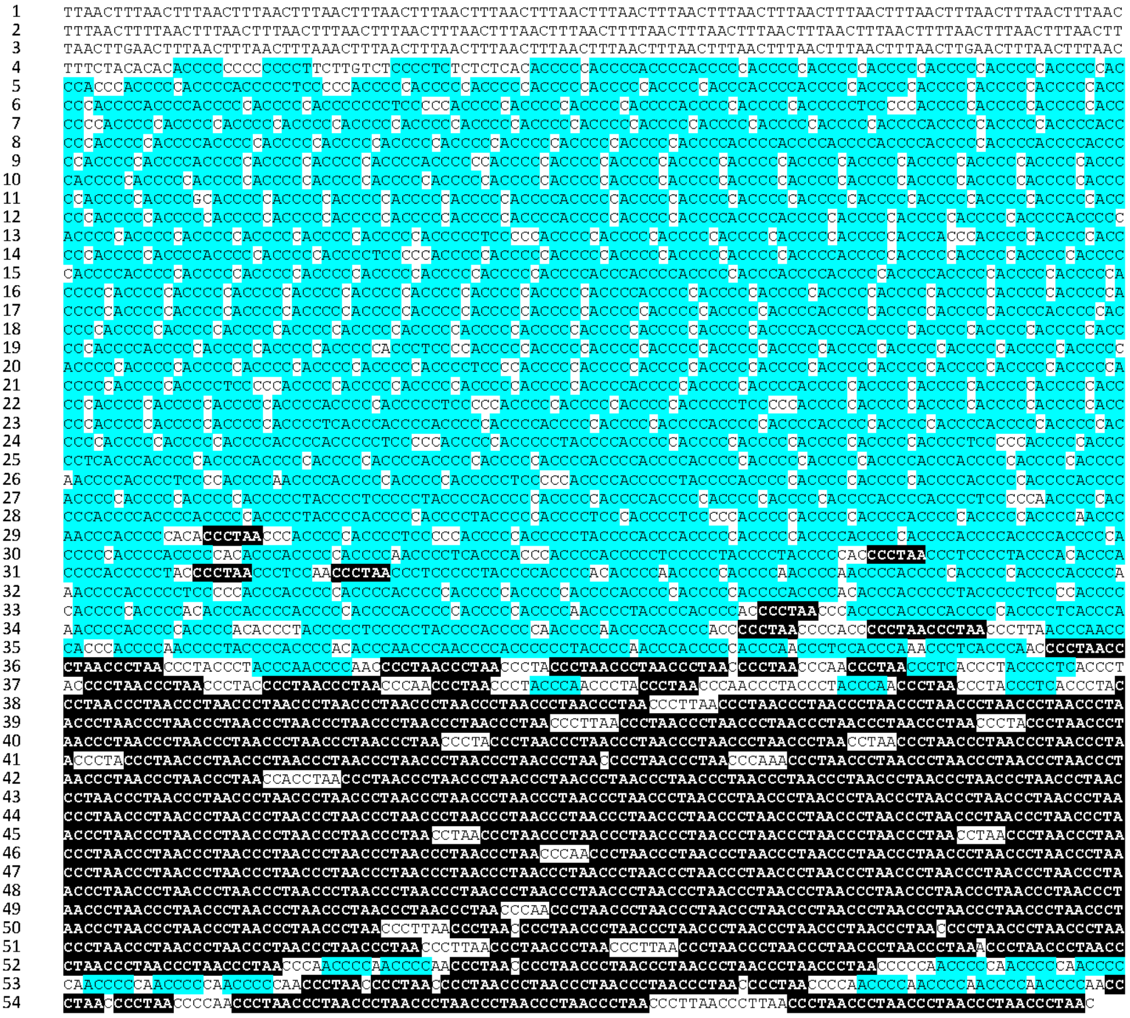

**Supplementary Fig. S3. Type 2 repeat errors and telomeric repeats in telomeric sequence.** Motifs are highlighted by whether they are suspected repeat errors or true telomeric repeats. Motifs that overlap are still highlighted.

**Type 3:** AACTTTA, AACGTTA, AATTTAA, AACTTA, AACTTT, TACTTA, ACTT

> NA18508#1#HPRCY2-hifiasm_JBHIJG010000004.1:1-1100000/nc#chr7:p-arm (1-7918)

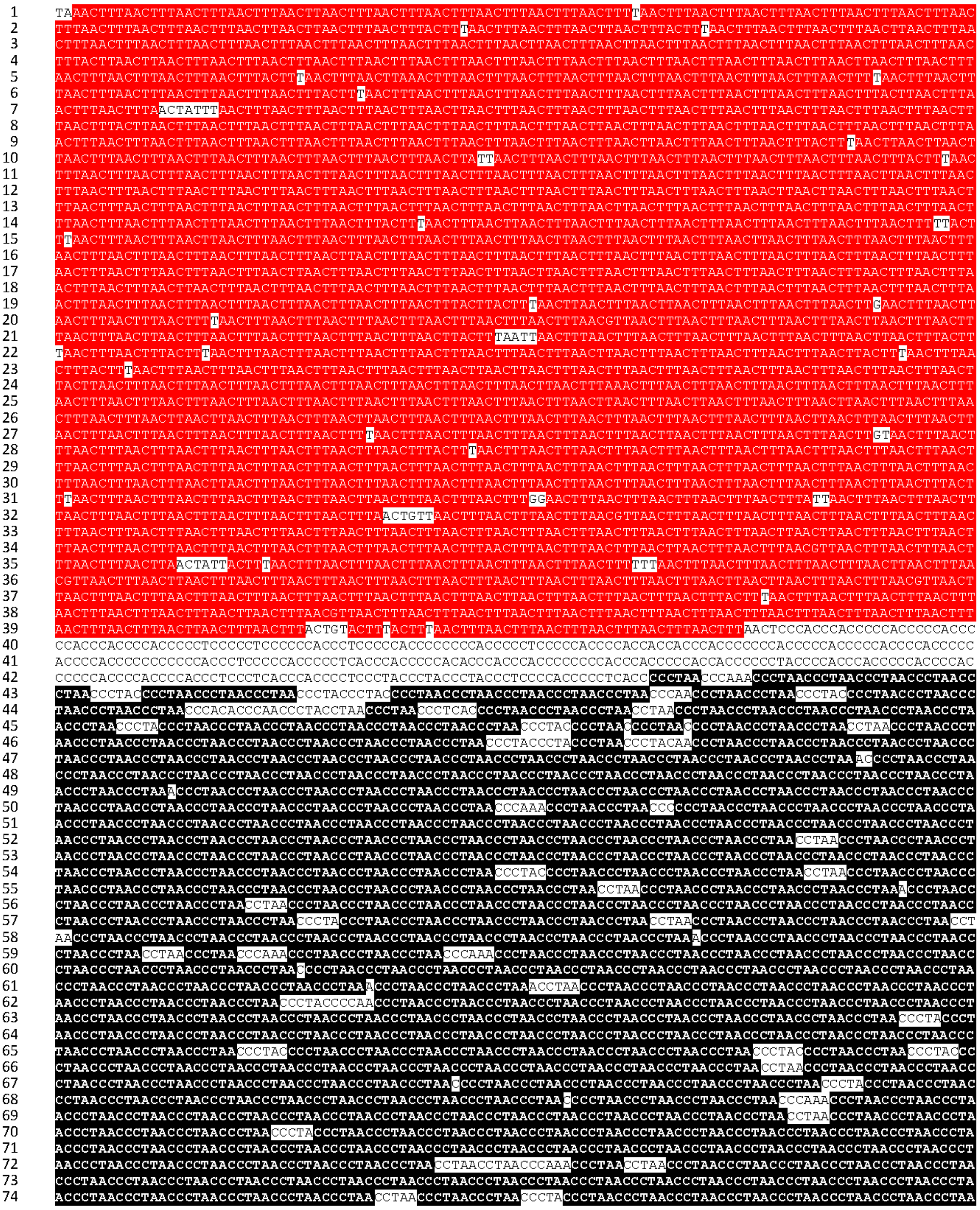

**Supplementary Fig. S4. Type 3 repeat errors and telomeric repeats in telomeric sequence.** Motifs are highlighted by whether they are suspected repeat errors or true telomeric repeats. Motifs that overlap are still highlighted.

**Type 4: GGGGGG, GGGGG, GGTGG, TGGGG**

`> HG03248#2#HGSVC3-hifiasm_h2tg0000081-1-58595512:57495513-58595512/rc#chr17:q-arm (1-7383)`

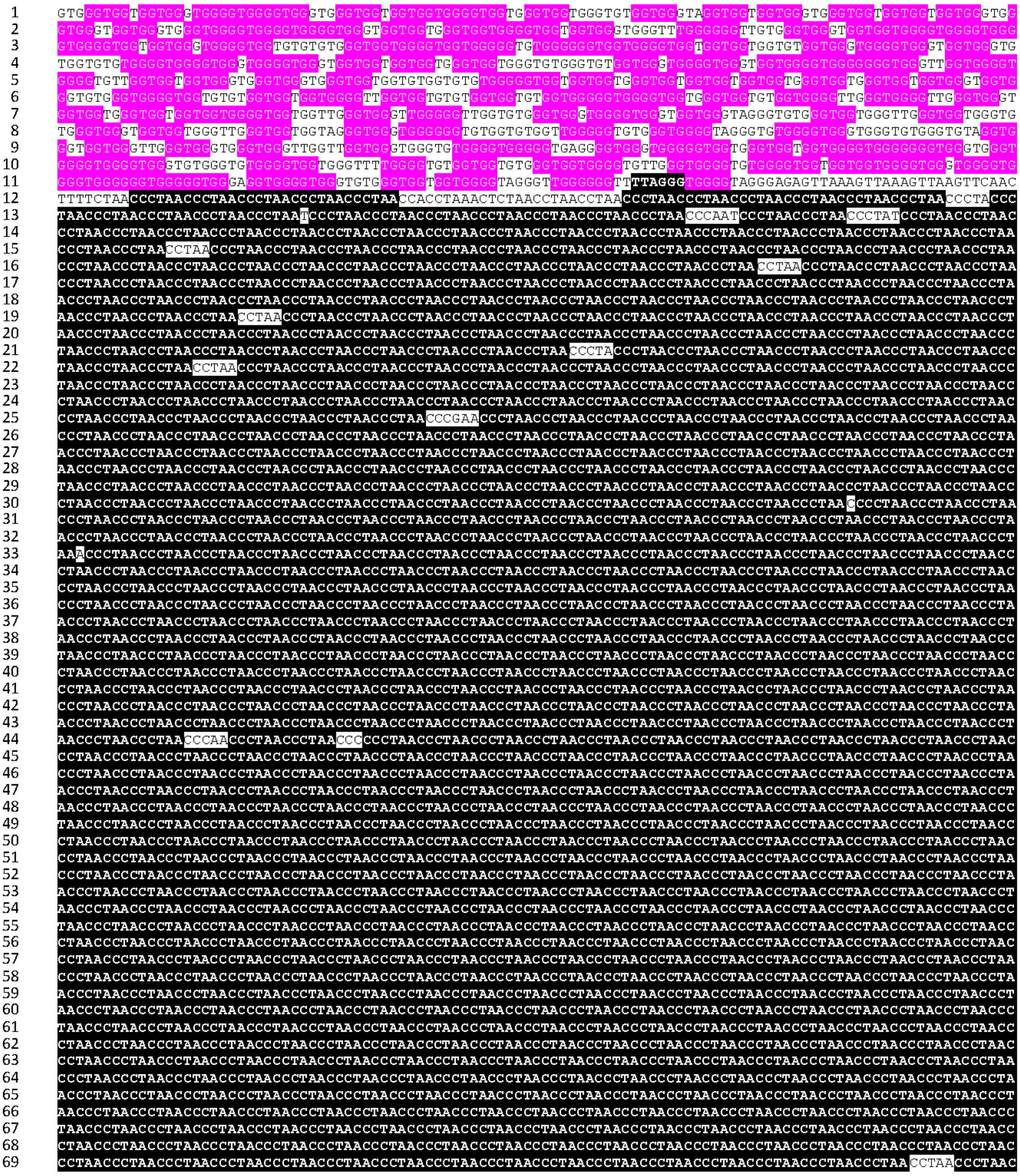

**Supplementary Fig. S5. Type 4 repeat errors and telomeric repeats in telomeric sequence.** Motifs are highlighted by whether they are suspected repeat errors or true telomeric repeats. Motifs that overlap are still highlighted.

**Type 5:** AACC, CCTA, ACCC, CTAACC, AACCCT

```
> HG00096#2#HGSVC3-hifiasm_h2tg000037l-1-24459145:1-
1100000/nc#chr17:p-arm (1-5175)
```

```
TTTAACTTCTACCCTAACCTTTAACCTTTAACTTAAATTTAACCTTCTAACCTTTAAATCTTAAACTTAACCTTTAACCTTTAACCTTTAACTTTAAATTTAACTTT
ACTTAACTTTAACTTTAACTTTACTAACTTTAACTTTAATTTAACTTTAACTTTAACTACTTAAAATTTACATTAACTTAACCTTTAACCTTTAACCCTTAACCCTA
ACCTACTTAACCCTTAACCTAATTTCACTTAACCCTTAAACCCTTAACTTCCTAACCTAACCCTAACCCTTTACTCCTTAACCCTTTAACTTCTAATCTAAACCCTA
ACCCTAACCCTAACCCAACCCACCTAACCCTAACCCTAACCTCTAACCCTAACCCAACCCACCCAACCCTACCCACCCCAACCCAACCCACCCCCAACCCCACCCCA
ACCACCCACCCACCACCCCACCCCACCCAACCCCCACCCAACCACCCCACCCACCCACACCCAACCCCAACCCAACCCCAACCCAACCACCCACCCCACCCACCCCA
CCACCCAACCCCACCCCACCCCACCCCCAACCCAACCCCAACCCACACCCAACCCAACCCCACCCACCCCAACCACCACCCACCCACCCCCAACCCACCCACCCCAC
CCCACCCCACCCACCCCAACCCCACCACCCACACCCCTACCCCAACCCACCCCAACCACCCCAACCCCACACCCTAACCCCCACACCCCACCACCCACACCACCCCA
CCCCACCCCAACCCCACCCACCCCAACCCACCACCCCACCCCAACCCACCCCACCCCACCACCCACCCACCCACACCCCACACCCACCACCCAACCCACACCCAACC
CCAACCCAACCCCCAACCCCAACCACCCACACCCAACCCACCCAACCCACCCCAACCCACCCACCCAACCCAACCCCAACCCAACCCCACACCCACCCCAACCCACC
CACACCCCTCACCCCAACCCACCCACCCACCCCAACCCCACCCCACCCCAACCCCAACCCCAACCCACCCCACCACCCACCACCCAACCCAACCCCACACCCACACC
CACCCCACCACCCCCACCCCACCCCACCCACACCACCCCACCCACACCCCACCCACCCCACCCCACCCCCACCCACACCCACACCCACACCCAACCCCCACCCAACC
CCAACCCCACCACCCACCCCCAACCCCACCCACCCCAACCCAACCCACCCACCCCACCCACCCCACACCCCAACCCACCCCACACCCACCCCAACCCCAACCACCCC
CAACCCCACCCCACCCACCCCACCCACCCCAACCCACCACCCACCACCCCCAACCCCACCCCACCCAACCCCACCCCAACCCAACCCCAACCCAACCCCACCACCCA
ACCCAACCCAACCCACCCCACCCAACCCCACCCACCCCACCACCCACCCACCCACCCCCAAACCCCACCCCCACCCCACCCAACCCAACCCCAACCCACCCACCCCT
ACCCCACCACCCACCACCACCCAACCCACCCAACCCACCCCACCCCAACCCACCCCACCACCCACACCCAACCCAACCCACCCACCCCCACCCACCCCAACCCACAC
CCACCCCACCCCACCCCTACCACCCACCCCAACCCACCCCAACCCCAACCCACCCACCCACCACCCCACCCCACCCCAACCCCACCCCACCCACCCCACCCCA
CCCACCCCACCCCACCCCAACCCACCCACCCCCACCCACCCACCCACCACCCACCCCACCCCACCCCCACCCCCACCACACCCCACCCCACCCCACCCACCCACCCA
CCCCACCCACCCAACCCCACCCACCCACCCAACCCCACCCCACCCCACCCCAACCCCACCCCAACCCACCCACCCACCCACCCCCACCCACCCACCCCACCCCACCC
CACCCCACCCCACCCAACCCACCCACCCCACACCCACCCCCACCCCACCCCCACCACCCCCACCCACACCCACCCCCCACCCACCCACCCCACCCAACCCCACCCCC
ACCCCCCACCCACCCACCCCAACCCACCACCCCAACCCAACCCACCCACCCACCCAACCACCCACCACCCACCCCACCCACCACCCACCCCACCCAACCCCACCCCA
CCCAACCCACCCACCCCACCCCACCACCCCACCCACCCACCCAACCCACCCACCCACCCACCCCAACCCACCCCACCCACCCACCCCACCCCACCCCAACCCC
AACCCCACCCACCCCCACCCCCACCCCCACCCCCACCCACCCCACCCCACCCCACCCCACACCCACCCACCCACCCCACCACCCAACCCAACCCACCCCACCCACCA
CCCCCACCCCACCACCACCCCACCCACCCCACCACCCACCCCACCCCCACCCCACCCACCCACCCACCCCAACCCACCCCCACCCCACCCCACCCCAACCCACCCCA
CCCCCACCCCCACCCCACCCCACCCACCCCACCCCACCCACCCCCACCCCCACCCCACCCCCACCCACCACCCCACCCCACCCCCACCCCACCCCCACCCCCACCCC
ACCCCACCCCACCCCCACACCCCACCCACCCCAACCCCACCCACCCCCACCCCACCCCAACCCCACCCACCCAACCCCAACCCCAACCCAACCCACACCCAACCCCA
CCCACCCCAACCCCACCCCAACCCAACCCCACCCAACCCCCACCCCAACCCCAACCCCACCCAACCCCACCCAACCCACCCACCCACCCACCACCCCCACCCCACCA
CCCAACCCCAACCCCACCACCCAACCCCCACCCCAACCCCCACCCCACCCCACCCCAACCCCAACCCCACCCCACCCCAACCCCAACCCCAACCCCCACCCCCACCC
CACCCAACCCCAACCCCCACCCCAACCCCAACCCCAACCCCCCCACCCAACCCAACCCTAACCCCAACCCCACCCCCCCAACCCCAACCCACACCCAACCCTAACCC
TAACCCACCCAACCCACCAACCCCAACCCCAACCCAACCCCAACCCCACCCAACCCAACCCCAACCCAACCCCTAACCCAACCCAACCCCTAACCCCCAACCCAACC
CAACCCAACCCCAACCCAACCCCAACCCCAACCCAACCCTAACCCAACCCAACCCTAACCCAAACCCTAACCCCTAACCCTAACCCTAACCCTACCCAACCCTAACC
CTAACCCCTAACCCTAACCCTAACCCTAACCCTAACCCTACCCTAACCCCTAACCCTAACCCAACCAACCCACCCACACCCAACCCAACCCTACCCACCCAACCCCT
AACCCACCCTAACCCAACCCACACCCCTAACCCAACCCTAACCCTAACCCAAACCCTAACCCCAACCCTAACCCTAACCCTAACCCTCAACCCTAACCCTAACCCTA
ACCCTAACCCTAACCCAACCCTAAACCCTAACCCTAACCCTAACCCTAACCCTAACCCTAACCCTAACCCTAACCCTAACCCTAACCCTAACCCTAACCCTAACCCT
AACCCTAACCCTAACCCTAACCCTAACCCTAAACCCTAAACCCTAAACCCTAAACCCTAACCCTAACCCTAACCCTAACCCTAACCCTAACCCTAACCCTAACCCTA
ACCCTAACCCTAACCCTAACACCCTAACACCCTAACACCCTAACACCCTAACACCCTAACCCTAACCCTCACCCTCACCCTCACCCTCACCCTCACCCTCACCCTCA
CCCTCACCCTCACCCTCACCCTAACCCAACCCTAACCCTAACCCTCACCCTCACCCTCACCCTCACCCTCACCCTCACCCTAACCCCAACCCCAACCCCCAACCCC
AACCCCAACCCCAACCCCAACCCCAACCCTAACCCTAACCCTAACCCTAACCCTAACCCTAACCCCTAACCCCTAACCCCTAACCCCTAACCCTAACCCTAACCCTA
ACCCTAACCCTAACCCTAACCCTAACCCTAACCCTAACCCCTAACCCCTAACCCCTAACCCCTAACCCCTAACCCTAACCCTAACCCTAACCCTAACCCTAACCCTA
ACCCTAACCCTAACCCTAACCCTAACCCTAACCCTAACCCTAACCCCTAACCCTAACCCCTAACCCCTAACCCTAACCCTAACCCTAACCCTAACCCTAACCCTAAC
CTAACCCTAACCCTAACCCTAACCCTAACCCTAACCCTAACCCTAACCCCCTAACCCTAACCCCCTAACCCCCTAACCCTAACCCTAACCCTAACCCTAACCCTAAC
CCTAACCCTAACCCTAACCCTAACCCTAACCCTAACCCTAACCCTAACCCTAACCCTAACCCCCTAACCCTAACCCCCTAACCCCCTAACCCTAACCCTAA
CCCTAACCCTAACCCTAACCCTAACCCTAACCCTAACCCTAACCCTAACCCTAACCCTAACCCCCTAACCCCCTAACCCTAACCCTAACCCTAACCCTAAC
CCTAACCCTAACCCTAACCCTAACCCTAACCCTAACCCTAACCCTAACCCTAACCCCCAACCCCAACCCCAACCCCAACCCCAACCCCAACCCCAACCCCAACCCCA
ACCCTAACCCTAACCCTAACCCTAACCCTAACCCTAACCCTAACCCTAACCCTAACCCCTAACCCCCTAACCCCCTAACCCCCTAACCCTAACCCTAACCC
TAACCCTAACCCTAACCCTAACCCTAACCCTAACCCTAACCCTAACCCTAACCTACCCCCTACCCCTACCCCTACCCCTACCCCTACCCCTAACCCTAACCCTAACCCTAAC
CCTAACCCTAACCCTAACCCTAACCCTAACCCTAACCCTAACCCTAACCCTAACCCTAACCCTAACCCTAACCCTAACCCTAACCCTAACCCTAACCCTAACCCTAA
CCCTAACCCTAACCCTAACCCTAACCCTAACCCTAACCCCTAACCCCTAACCCCTAACCCCTAACCCTAACCCTAACCCTAACCCTAACCCTAACCCTAACCCTAACCCCTAA
CCCTTAACCCTAACCCTAACCCTAACCCTAACCCTAACCCTAACCCTACCCTAACCCTAACCCTAACCCTAACCCTAACCCTAACCCTAACCCTAACCCTACCCTAA
CCTAACCCTAACCCTAACCCTAACCCTAACCCTAACCCT
```

**Supplementary Fig. S6. Type 5 repeat errors and telomeric repeats in telomeric sequence.** Motifs are highlighted by whether they are suspected repeat errors or true telomeric repeats. Motifs that overlap are still highlighted.

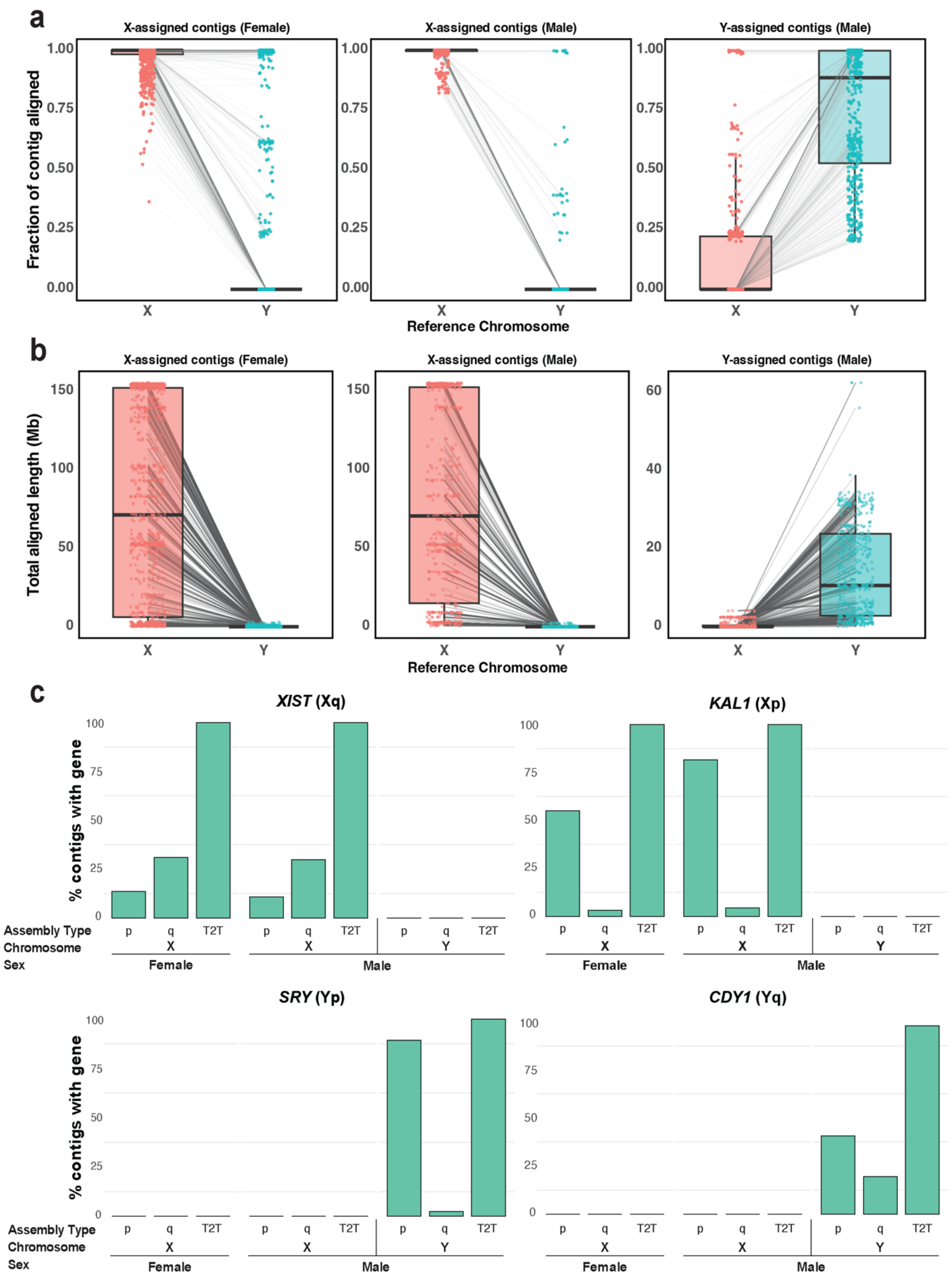


**Supplementary Fig. S7. Assignment of contigs to sex chromosomes.** (**a**) Alignment fraction and (**b**) alignment length of contigs to ChrX (red) and ChrY (blue). Each point represents a single contig. (**c**) Percentage of contigs assigned to each sex chromosome that contain genes specific to ChrX (top), and ChrY(bottom).

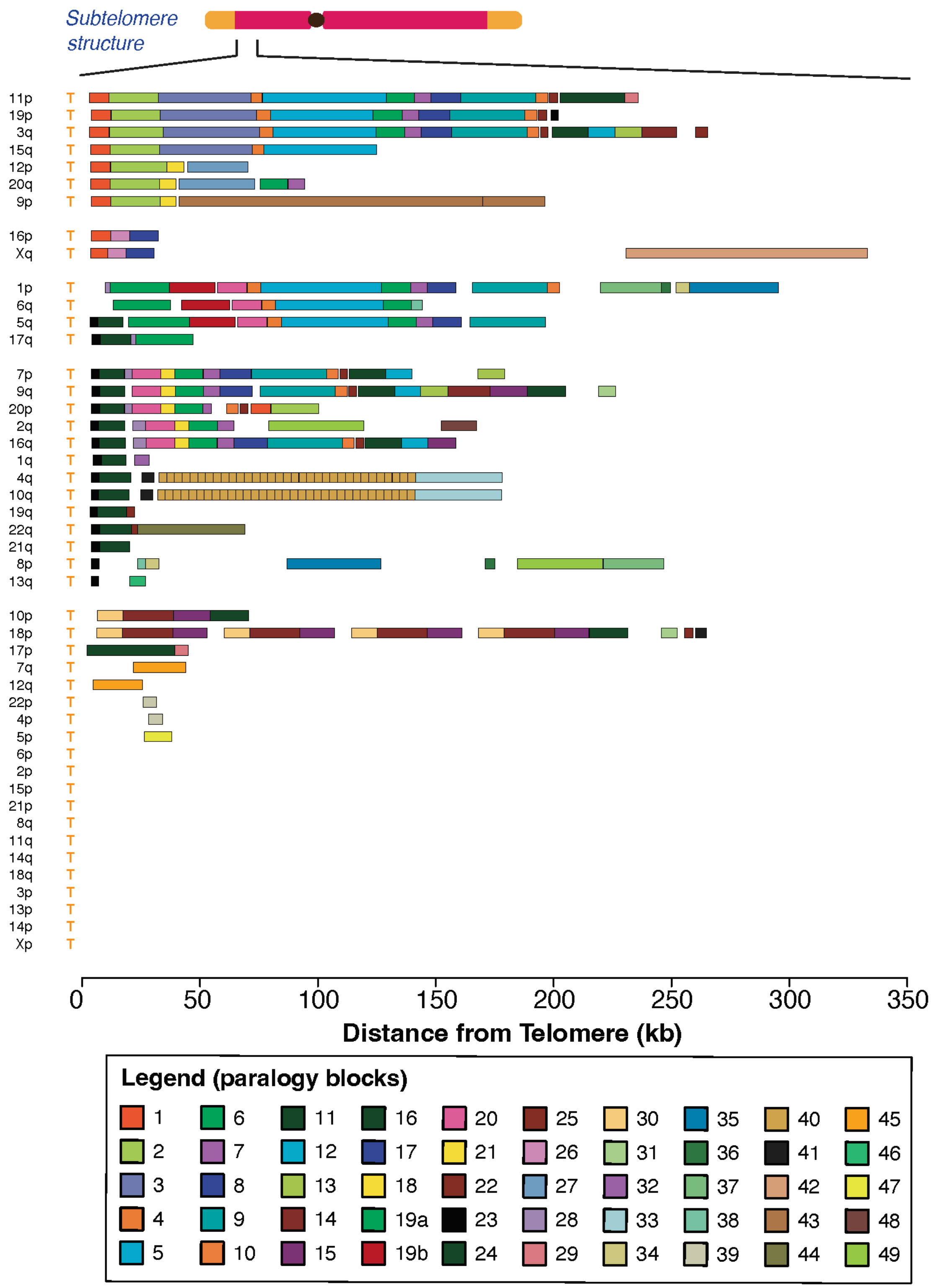


**Supplementary Fig. S8. Paralogy blocks found through mapping to subtelomeric sequences of CHM13.** Reference sequences from previous literature were mapped to CHM13 via minimap2, and the resultant paralogy block sequences were sorted visually.

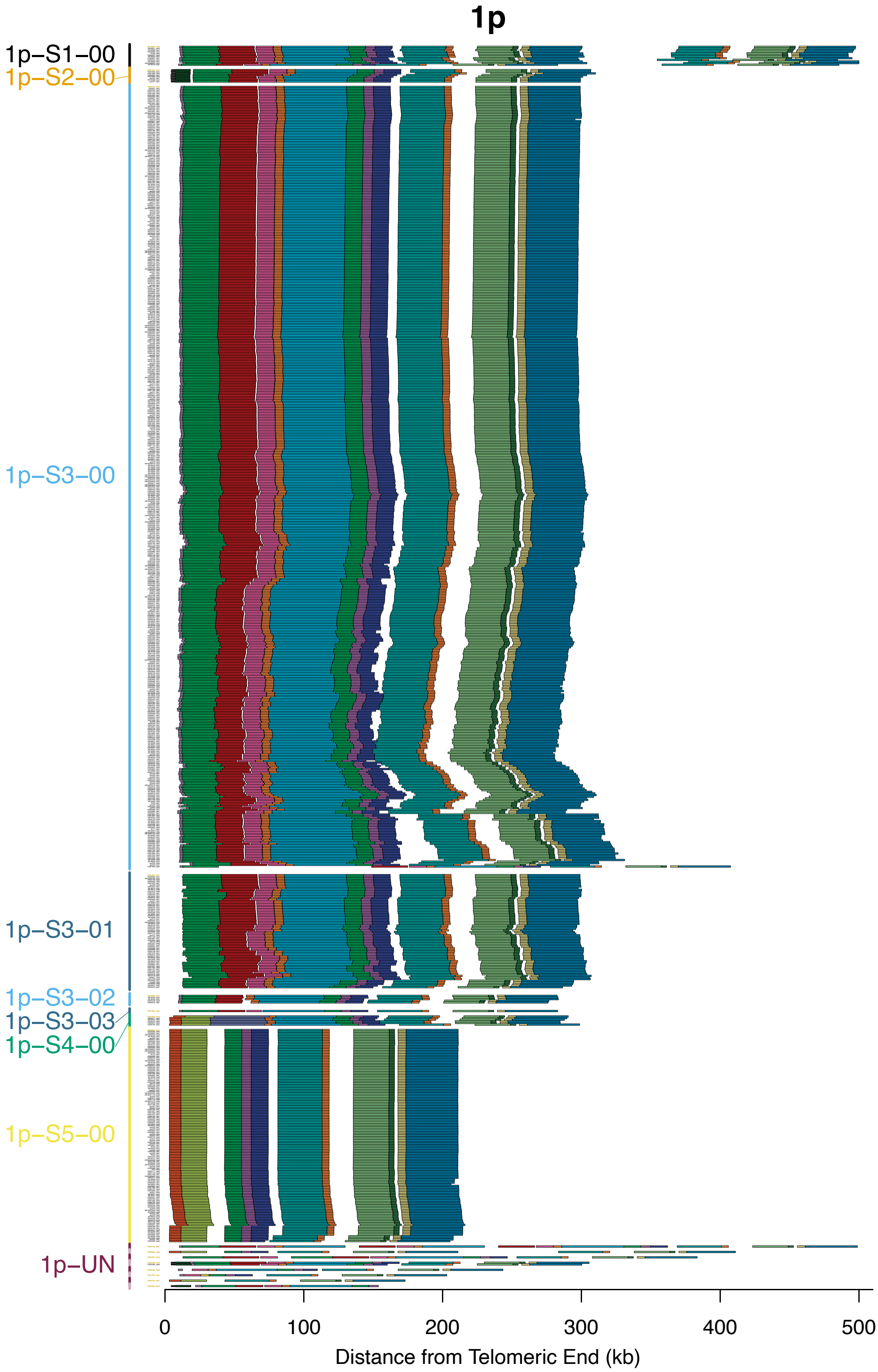


**Supplementary Fig. S9. Subtelomere contigs in 1p grouped based on paralogy block sequence.** Contigs differing by two or fewer paralogy blocks are grouped into the same major haplotype. Within a major haplotype, contigs differing by at least one paralogy block are further separated into minor haplotypes. Haplotypes are indicated by labels on the left. The representative contig of each haplotype is positioned at the top and labelled with golden text. Contigs are arranged by starting with the representative, then taking the contig with the nearest start and end positions. All contigs are visualised with the telomeric end on the left and the centromeric end on the right.

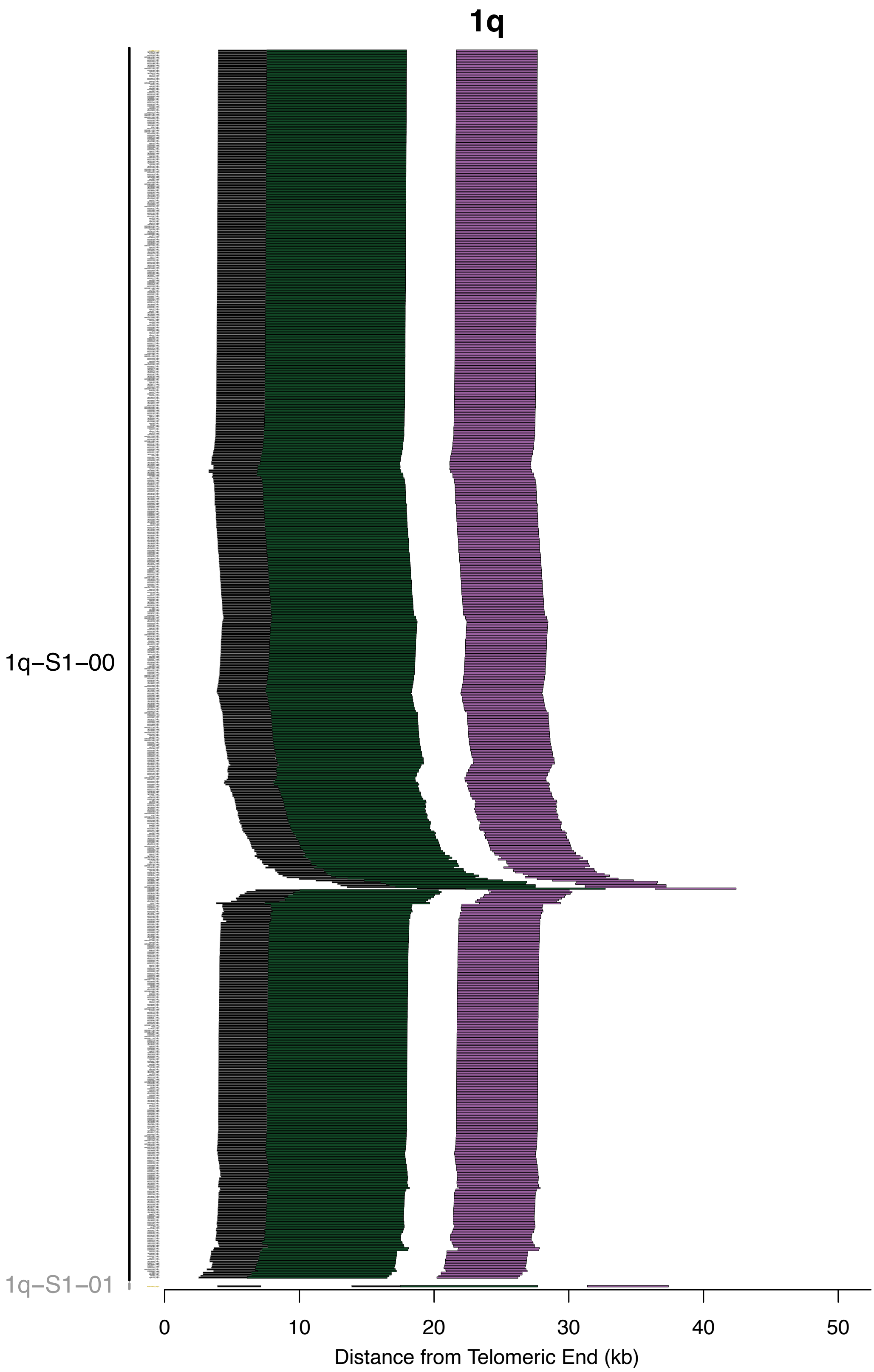


**Supplementary Fig. S10. Subtelomere contigs in 1q grouped based on paralogy block sequence.** Figure interpretation follows that of Supplementary Figure S9, but for chromosome arm 1q.

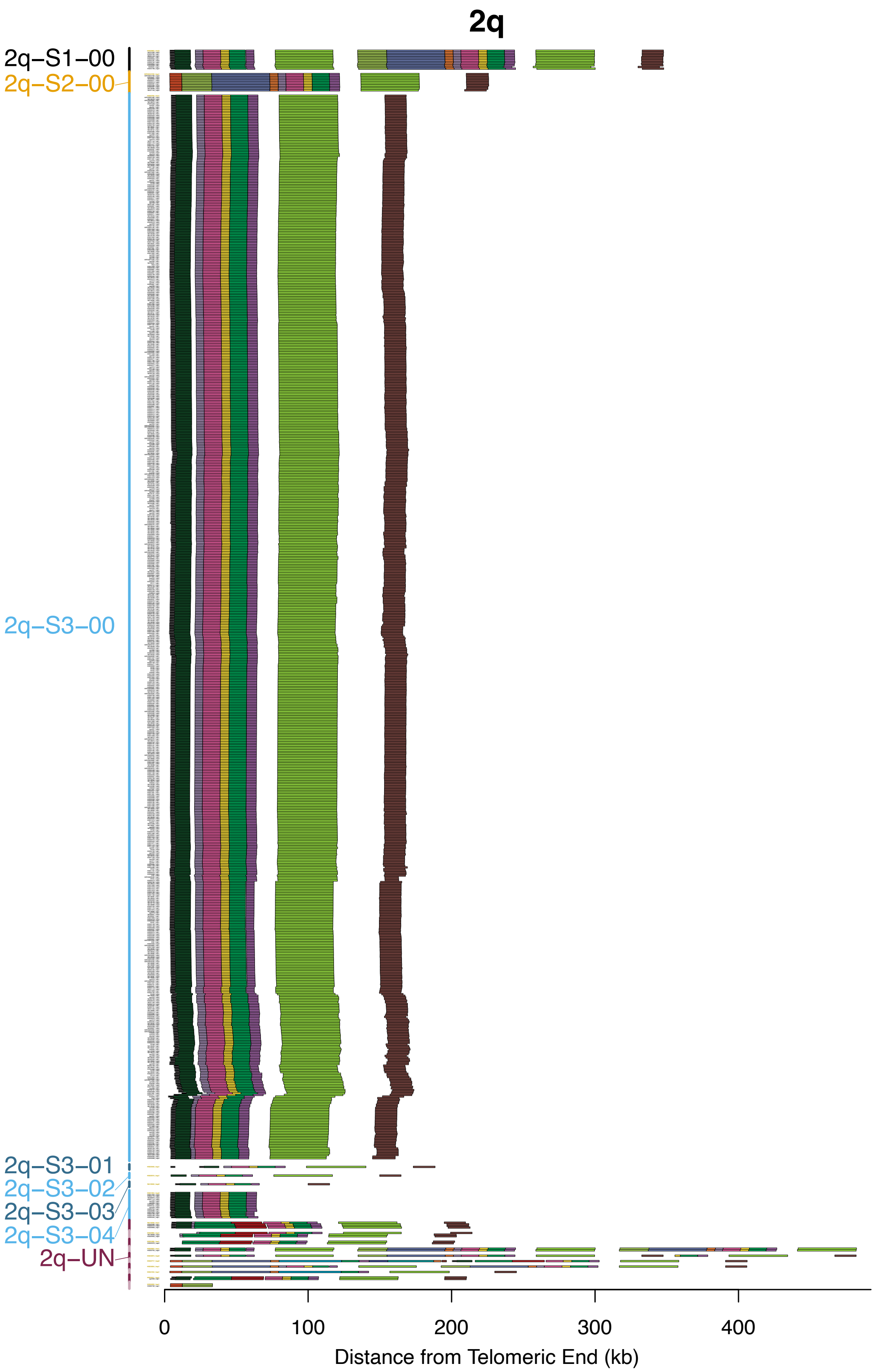


**Supplementary Fig. S11. Subtelomere contigs in 2q grouped based on paralogy block sequence.** Figure interpretation follows that of Supplementary Figure S9, but for chromosome arm 2q.

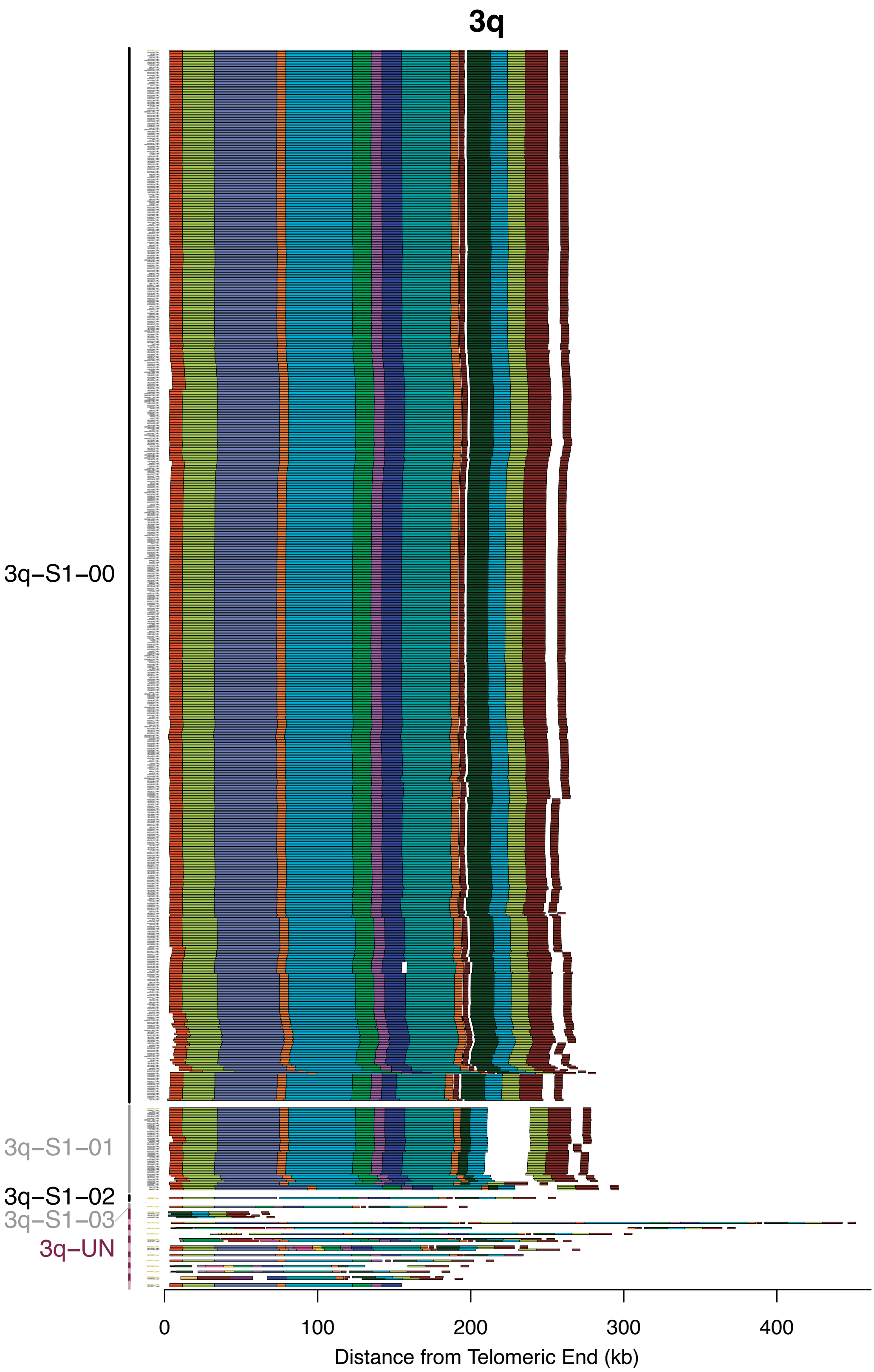


**Supplementary Fig. S12. Subtelomere contigs in 3q grouped based on paralogy block sequence.** Figure interpretation follows that of Supplementary Figure S9, but for chromosome arm 3q.

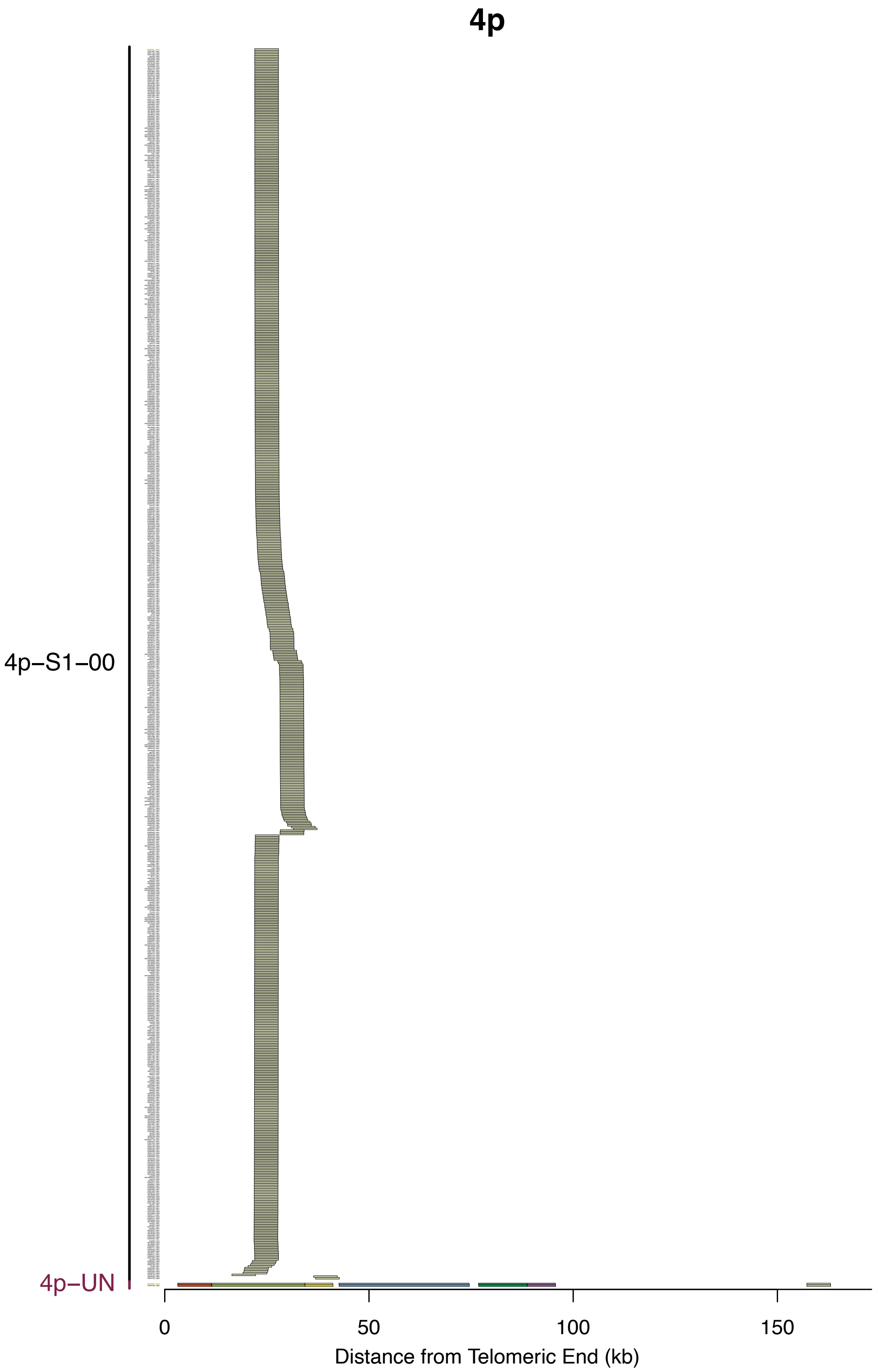


**Supplementary Fig. S13. Subtelomere contigs in 4p grouped based on paralogy block sequence.** Figure interpretation follows that of Supplementary Figure S9, but for chromosome arm 4p.

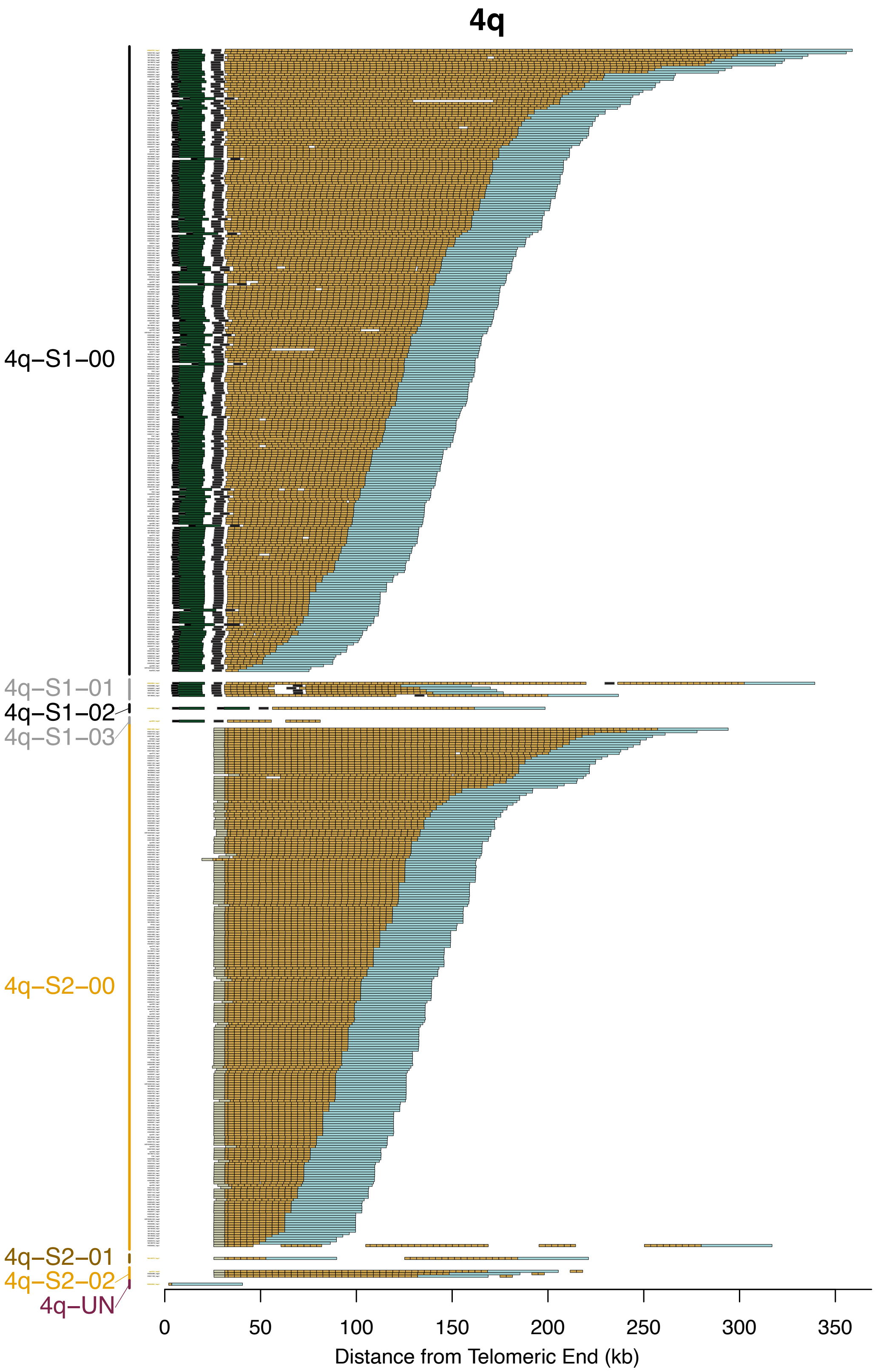


**Supplementary Fig. S14. Subtelomere contigs in 4q grouped based on paralogy block sequence.** Figure interpretation follows that of Supplementary Figure S9, but for chromosome arm 4q.

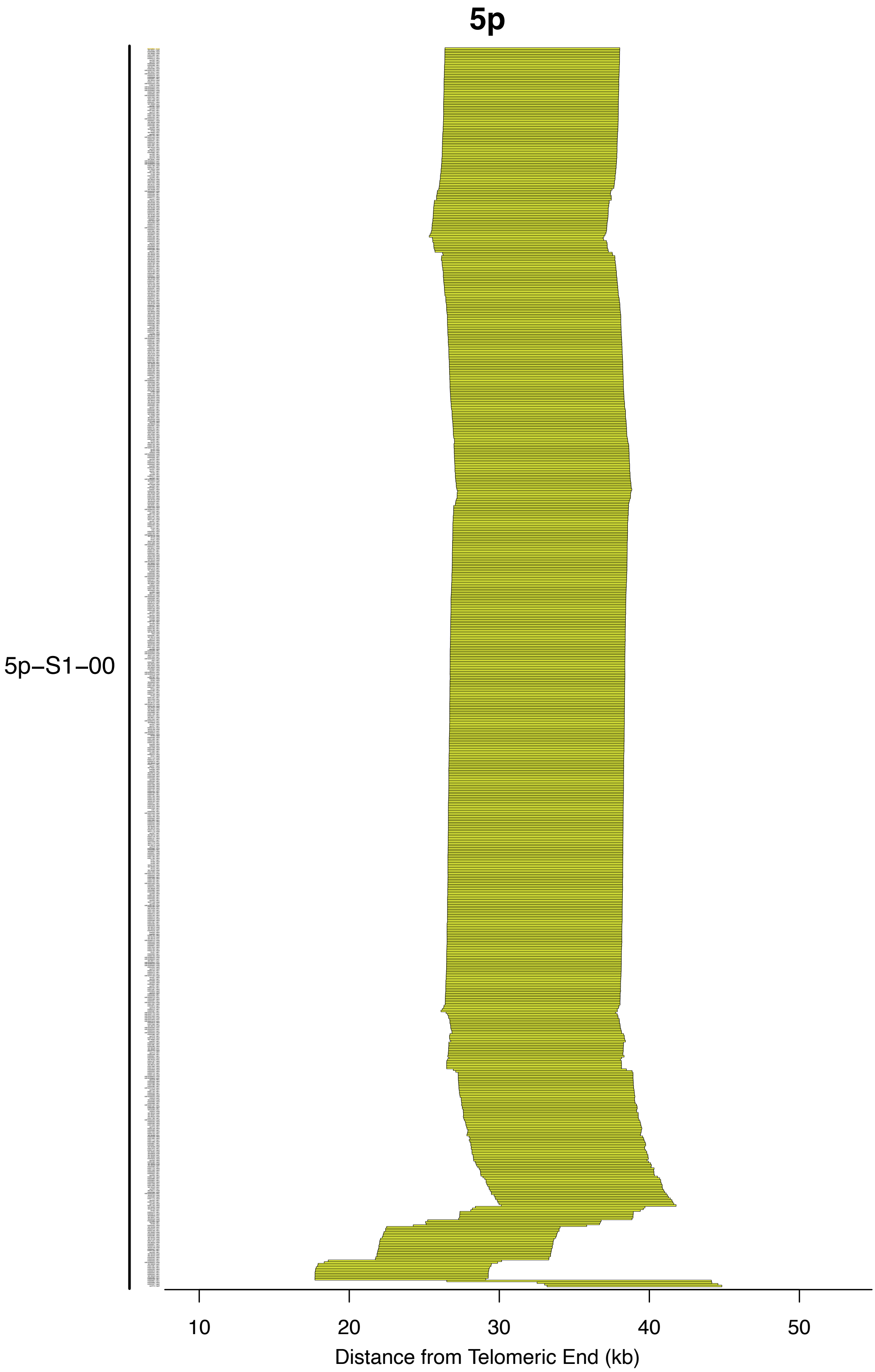


**Supplementary Fig. S15. Subtelomere contigs in 5p grouped based on paralogy block sequence.** Figure interpretation follows that of Supplementary Figure S9, but for chromosome arm 5p.

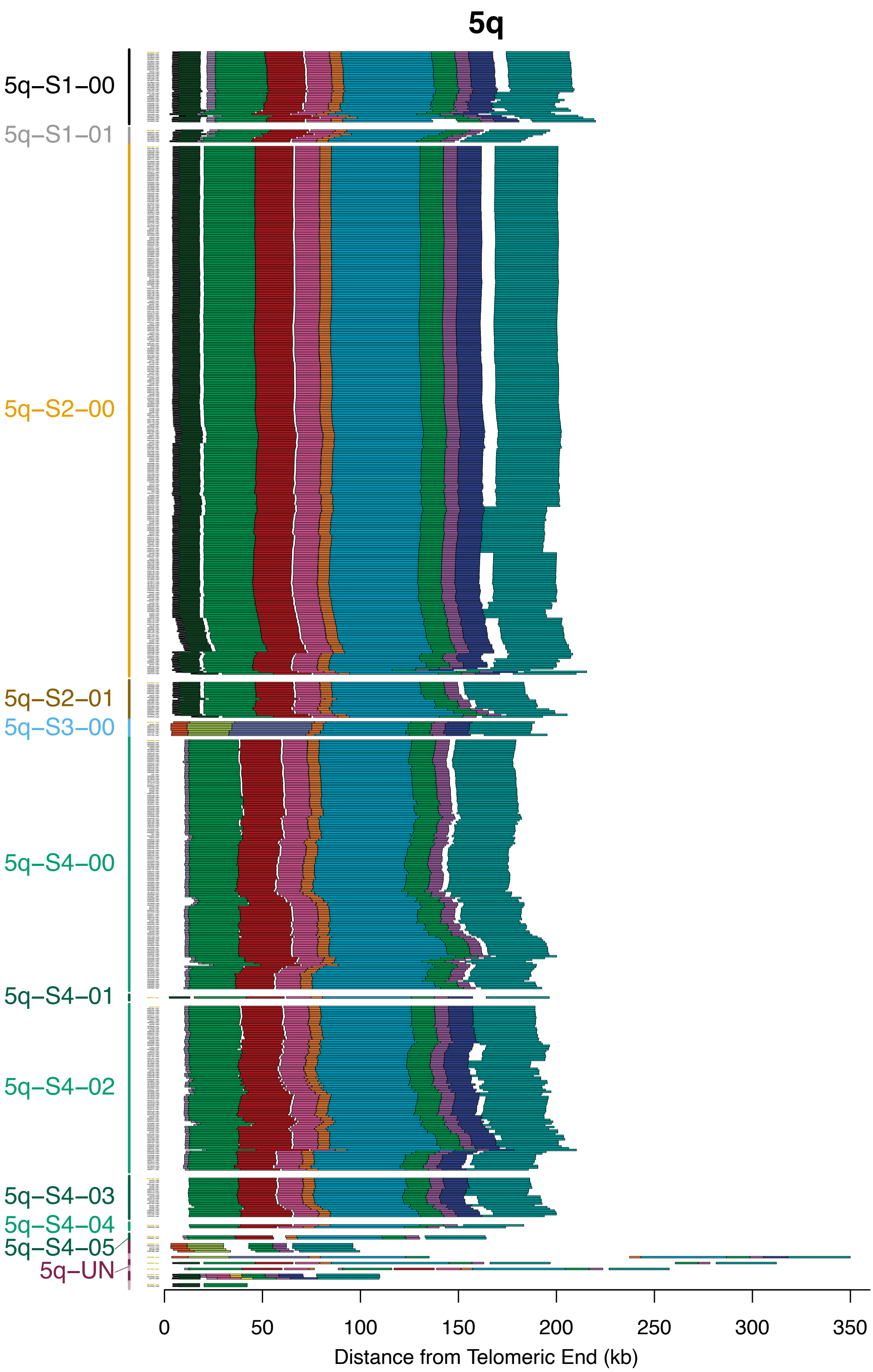


**Supplementary Fig. S16. Subtelomere contigs in 5q grouped based on paralogy block sequence.** Figure interpretation follows that of Supplementary Figure S9, but for chromosome arm 5q.

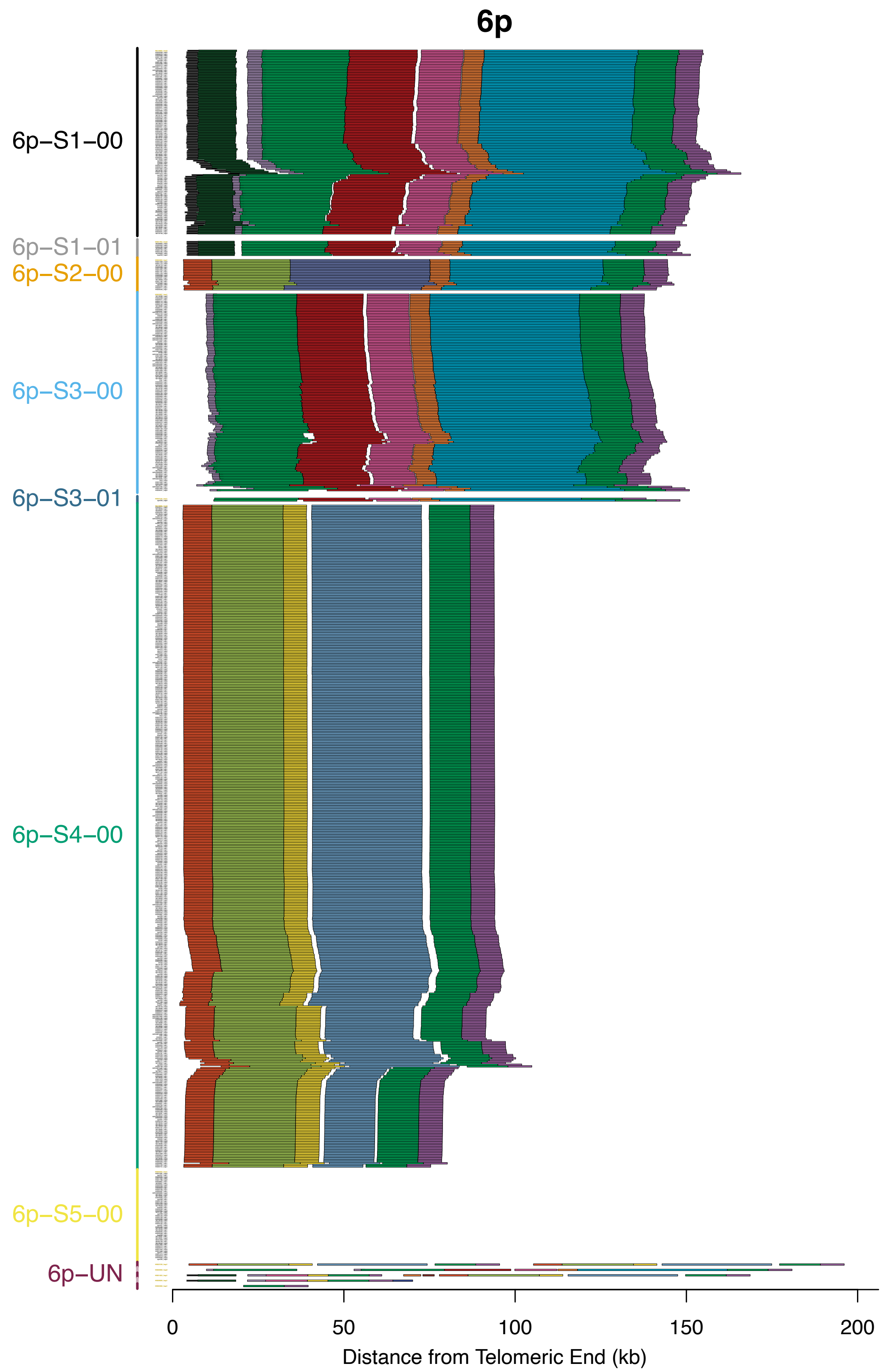


**Supplementary Fig. S17. Subtelomere contigs in 6p grouped based on paralogy block sequence.** Figure interpretation follows that of Supplementary Figure S9, but for chromosome arm 6p.

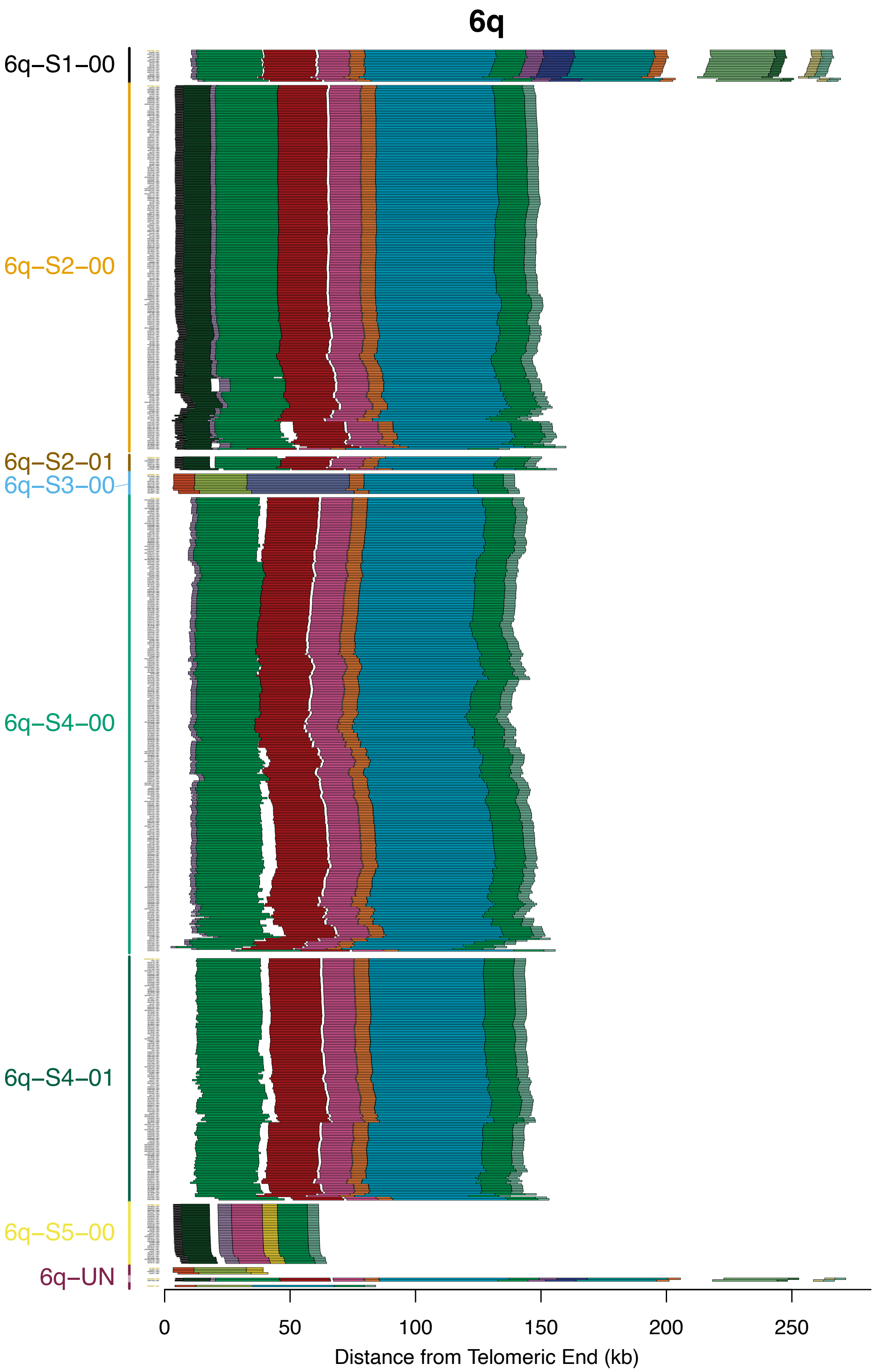


**Supplementary Fig. S18. Subtelomere contigs in 6q grouped based on paralogy block sequence.** Figure interpretation follows that of Supplementary Figure S9, but for chromosome arm 6q.

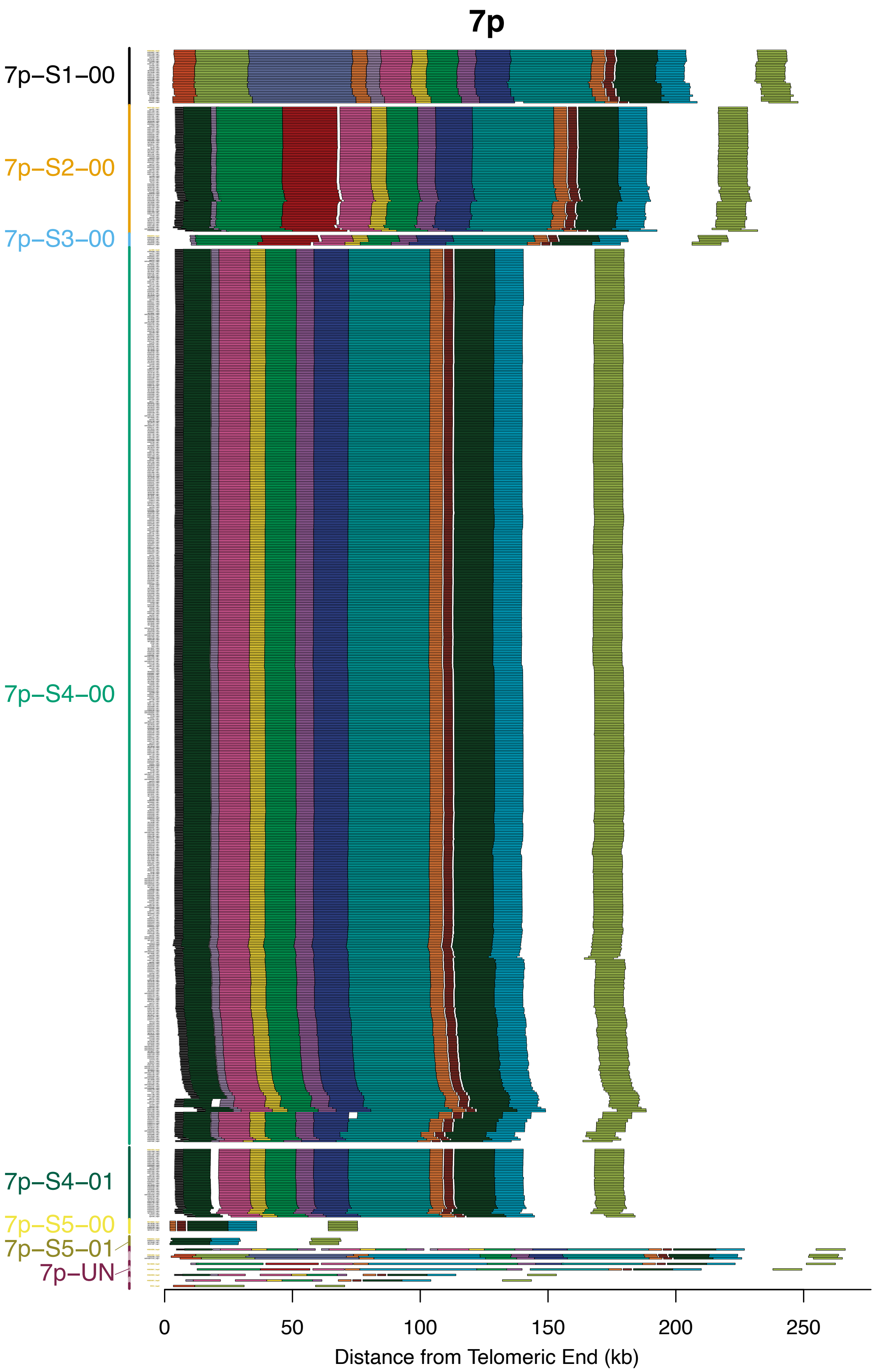


**Supplementary Fig. S19. Subtelomere contigs in 7p grouped based on paralogy block sequence.** Figure interpretation follows that of Supplementary Figure S9, but for chromosome arm 7p.

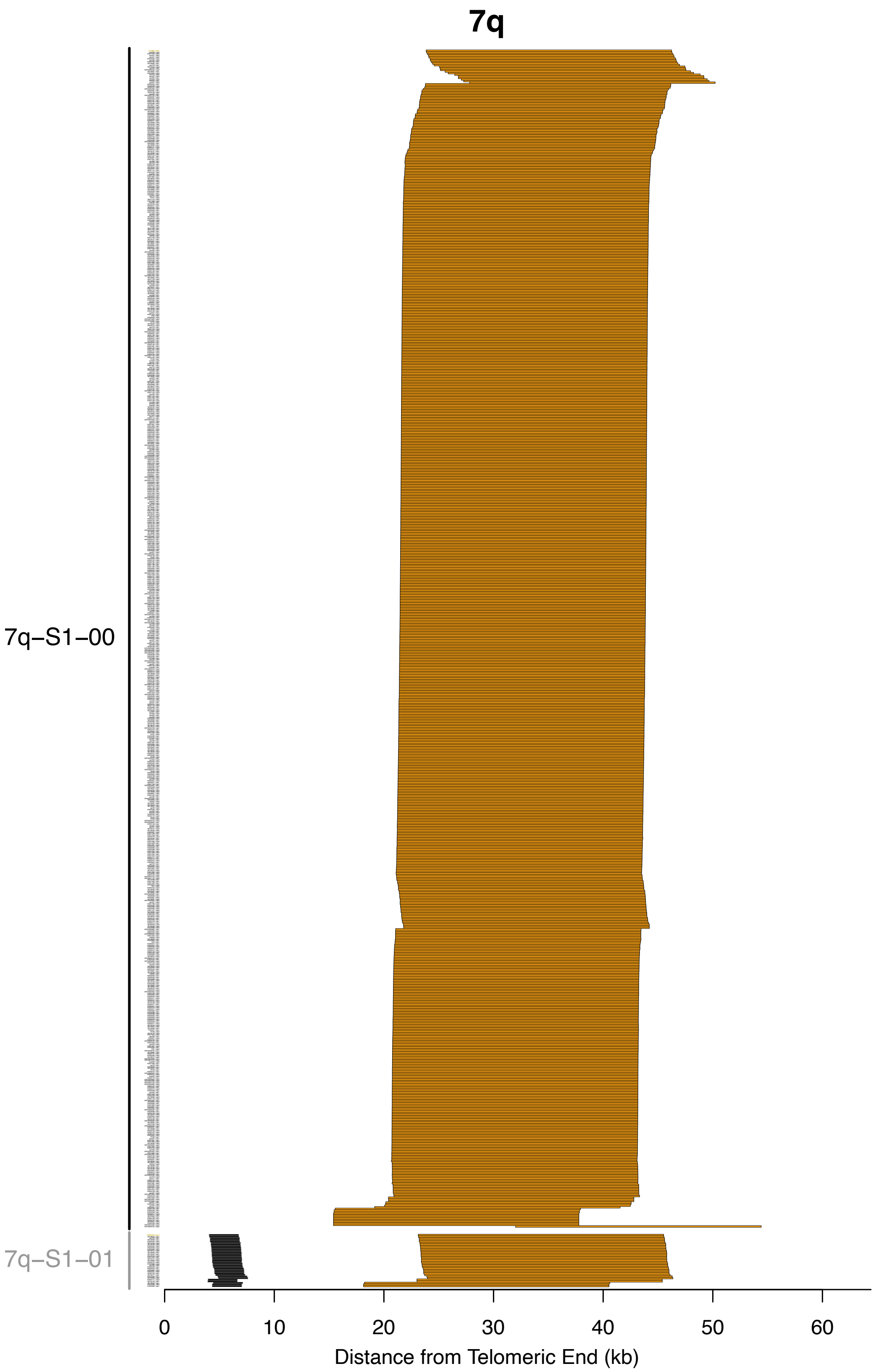


**Supplementary Fig. S20. Subtelomere contigs in 7q grouped based on paralogy block sequence.** Figure interpretation follows that of Supplementary Figure S9, but for chromosome arm 7q.

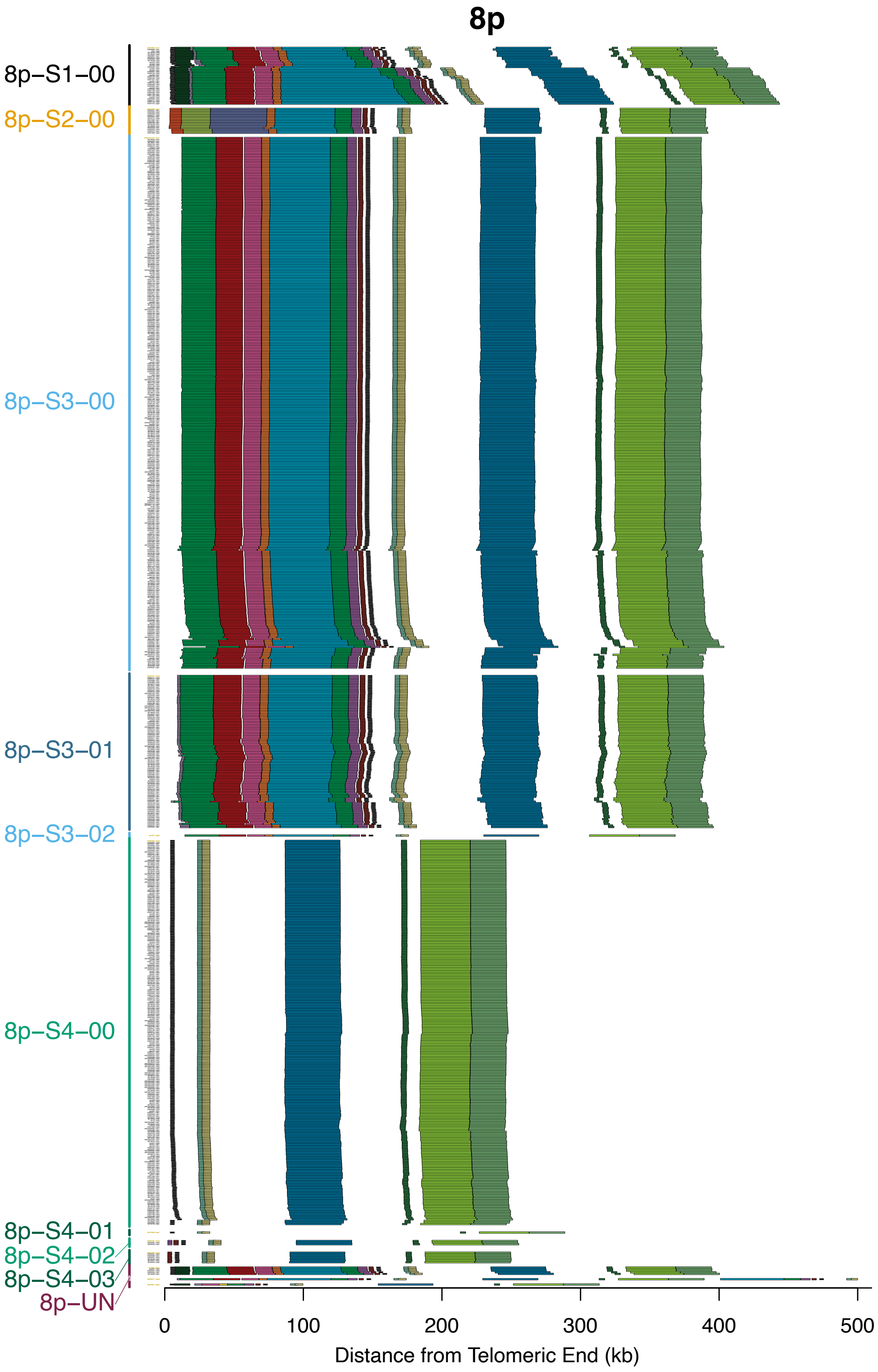


**Supplementary Fig. S21. Subtelomere contigs in 8p grouped based on paralogy block sequence.** Figure interpretation follows that of Supplementary Figure S9, but for chromosome arm 8p.

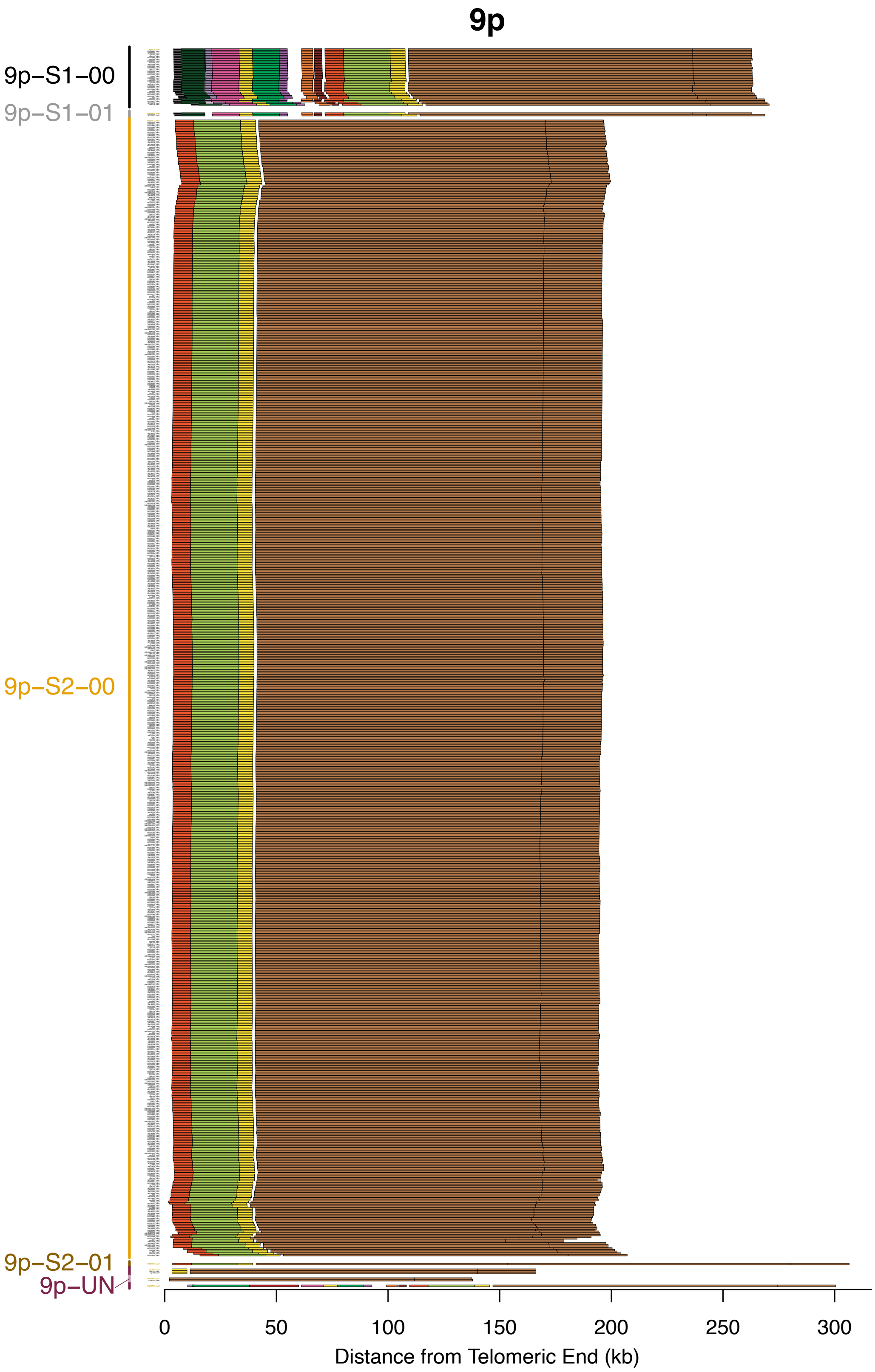


**Supplementary Fig. S22. Subtelomere contigs in 9p grouped based on paralogy block sequence.** Figure interpretation follows that of Supplementary Figure S9, but for chromosome arm 9p.

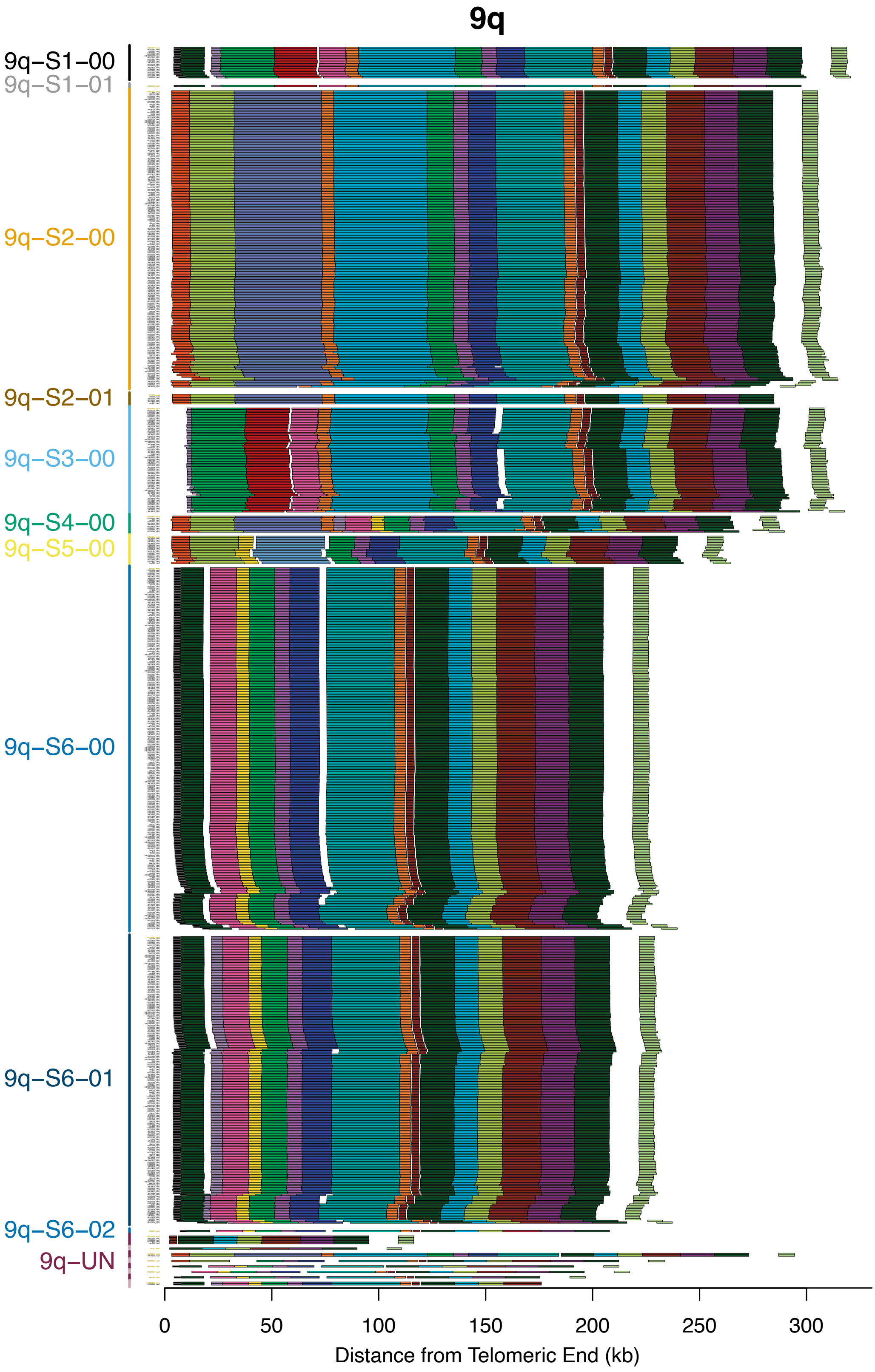


**Supplementary Fig. S23. Subtelomere contigs in 9q grouped based on paralogy block sequence.** Figure interpretation follows that of Supplementary Figure S9, but for chromosome arm 9q.

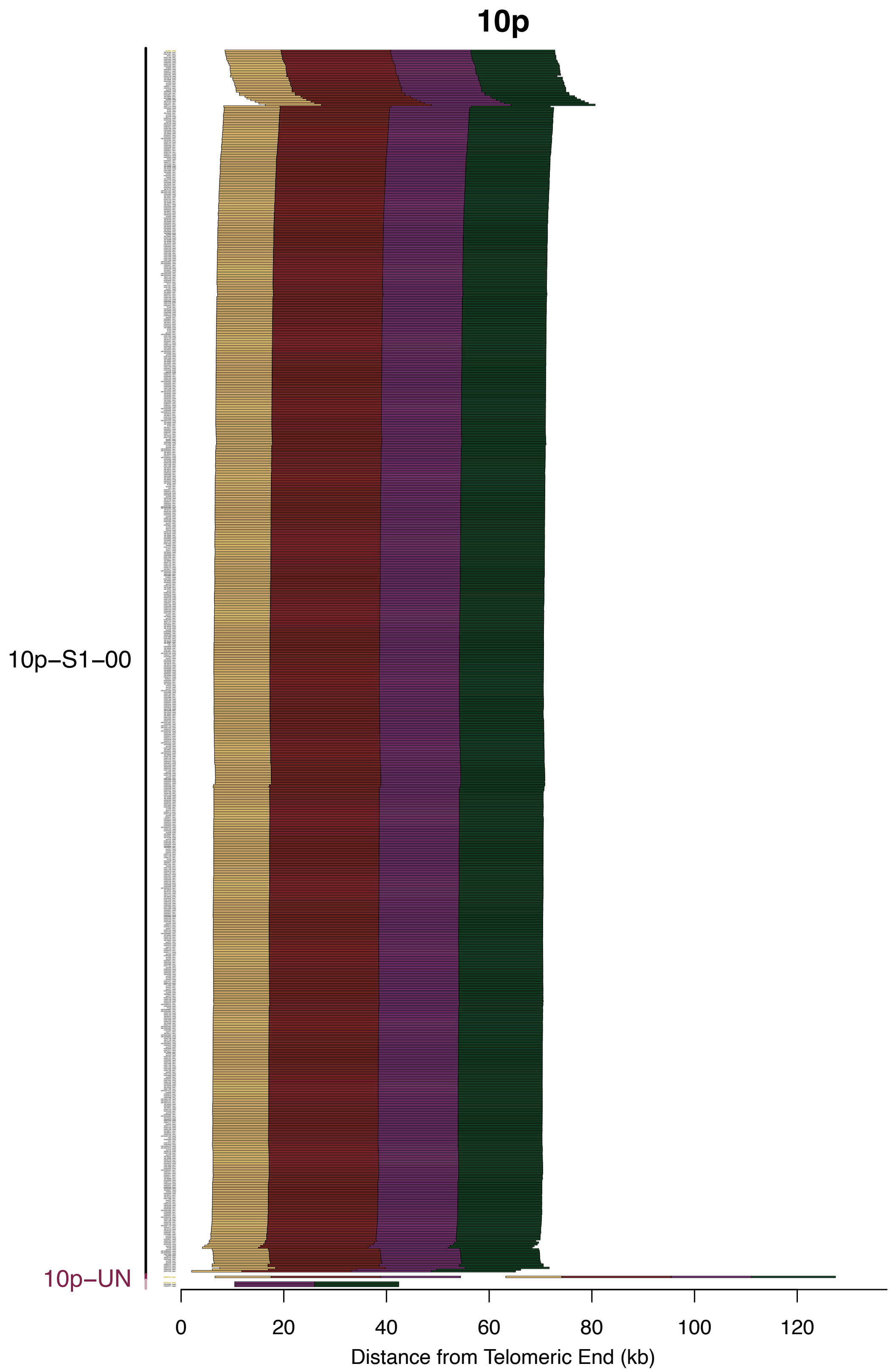


**Supplementary Fig. S24. Subtelomere contigs in 10p grouped based on paralogy block sequence.** Figure interpretation follows that of Supplementary Figure S9, but for chromosome arm 10p.

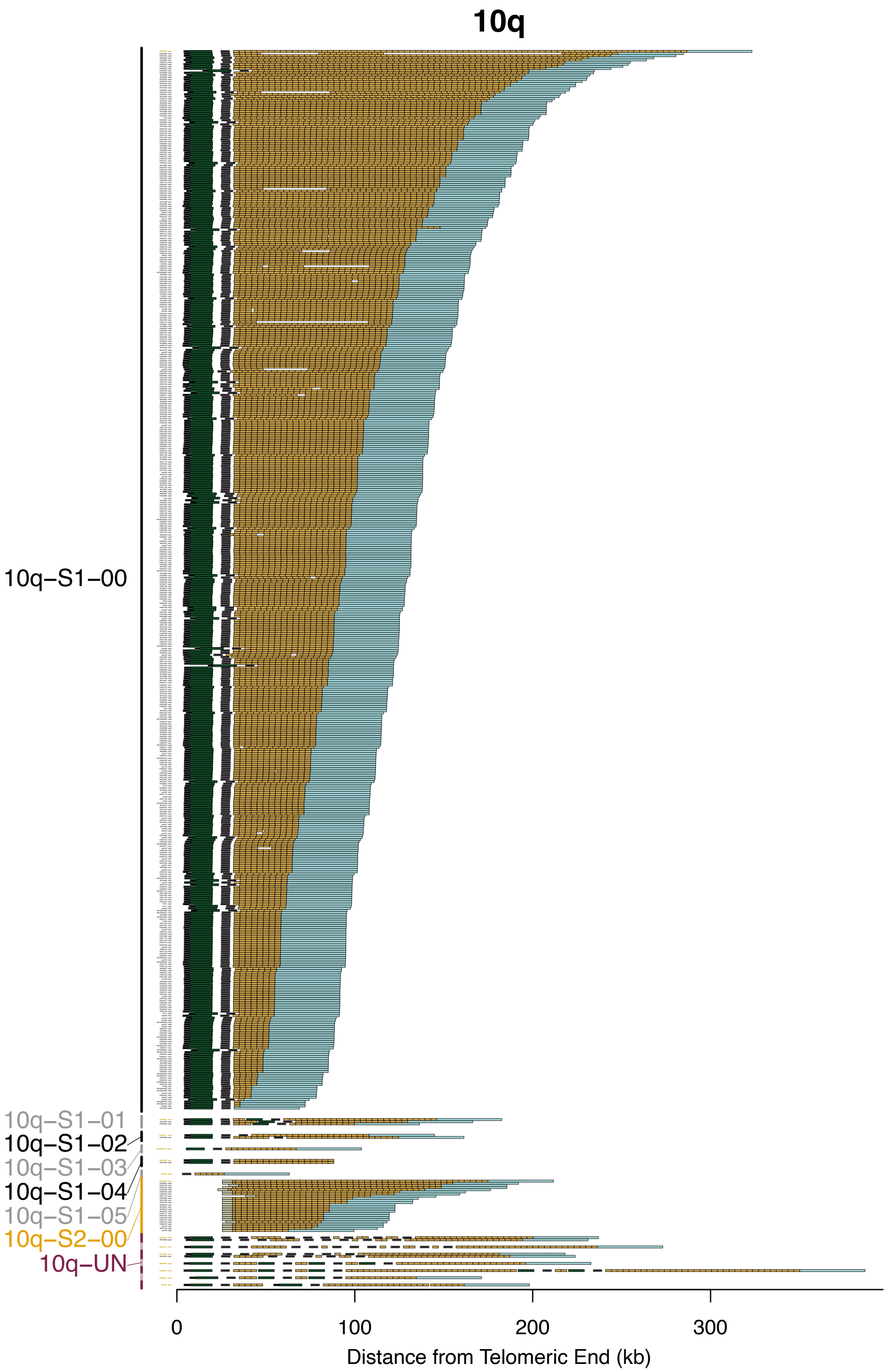


**Supplementary Fig. S25. Subtelomere contigs in 10q grouped based on paralogy block sequence.** Figure interpretation follows that of Supplementary Figure S9, but for chromosome arm 10q.

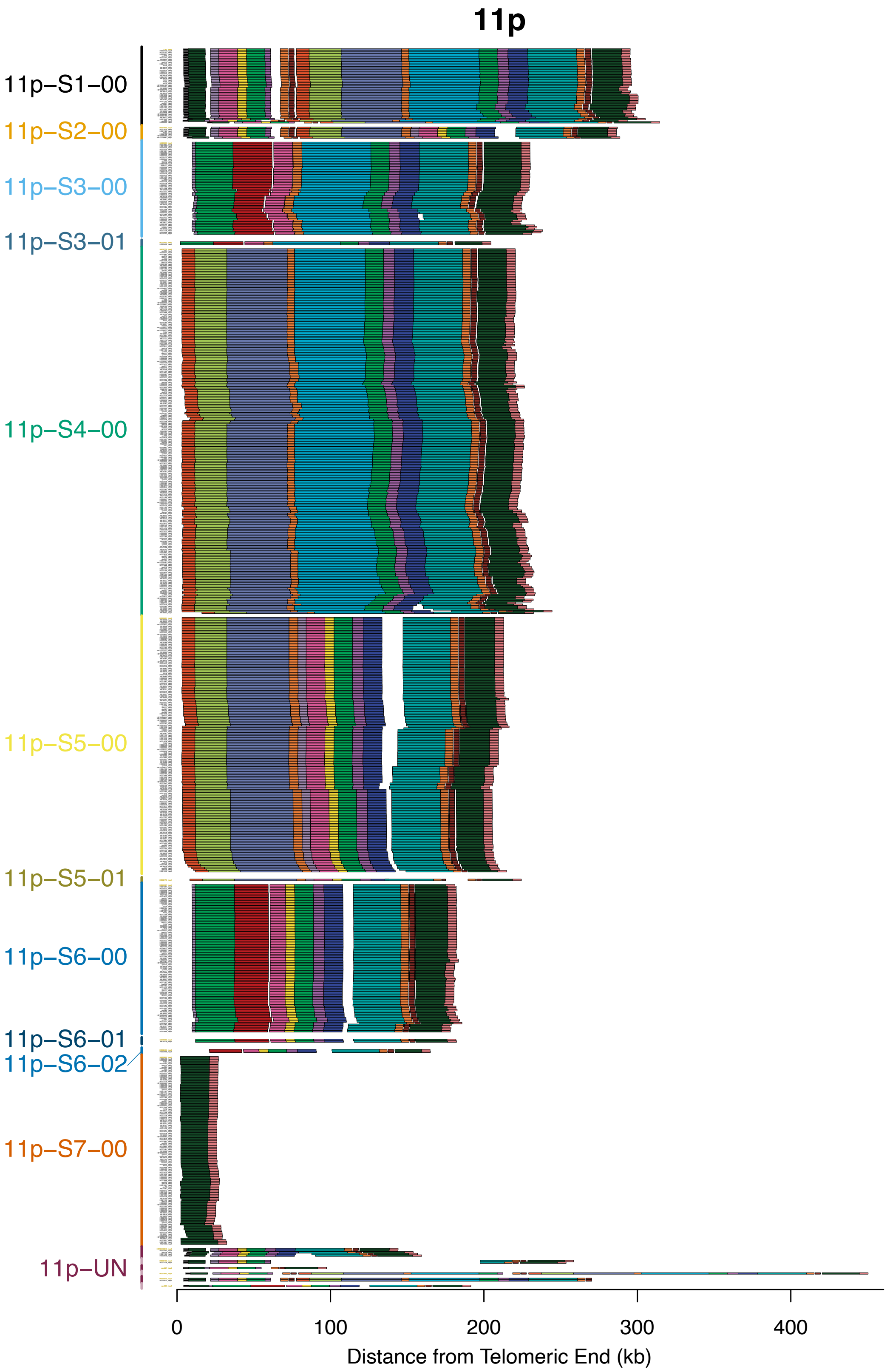


**Supplementary Fig. S26. Subtelomere contigs in 11p grouped based on paralogy block sequence.** Figure interpretation follows that of Supplementary Figure S9, but for chromosome arm 11p.

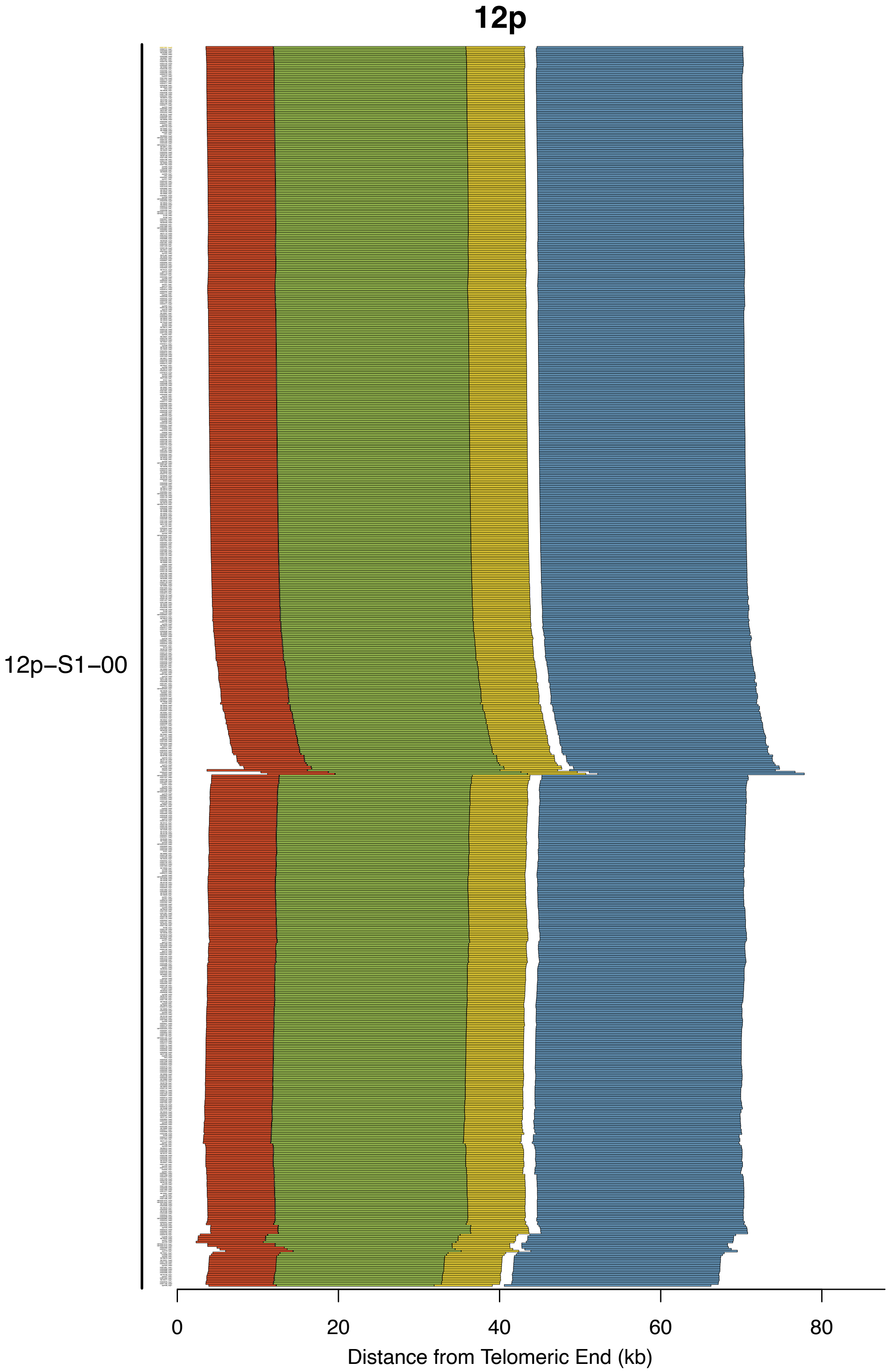


**Supplementary Fig. S27. Subtelomere contigs in 12p grouped based on paralogy block sequence.** Figure interpretation follows that of Supplementary Figure S9, but for chromosome arm 12p.

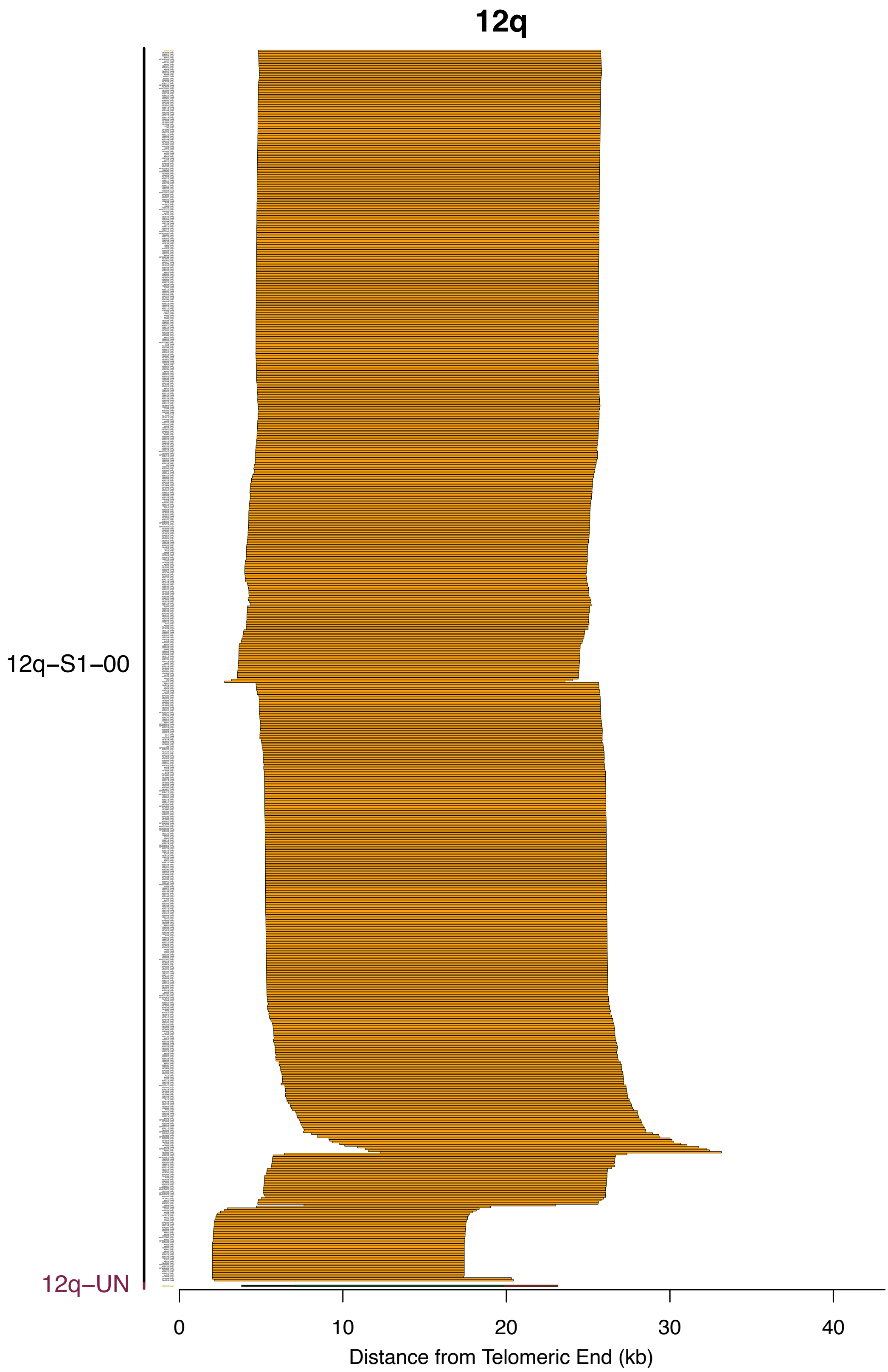


**Supplementary Fig. S28. Subtelomere contigs in 12q grouped based on paralogy block sequence.** Figure interpretation follows that of Supplementary Figure S9, but for chromosome arm 12q.

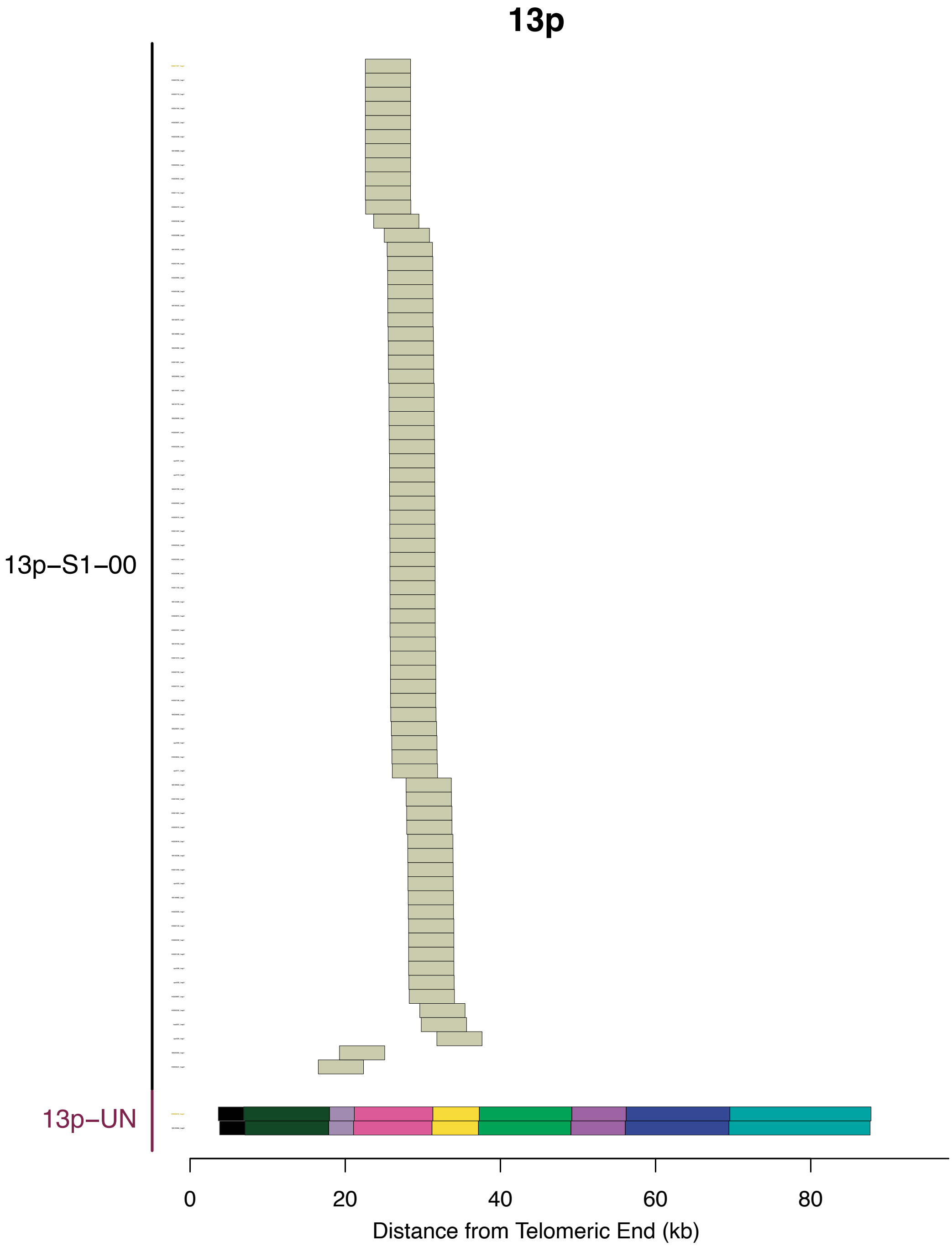


**Supplementary Fig. S29. Subtelomere contigs in 13p grouped based on paralogy block sequence.** Figure interpretation follows that of Supplementary Figure S9, but for chromosome arm 13p.

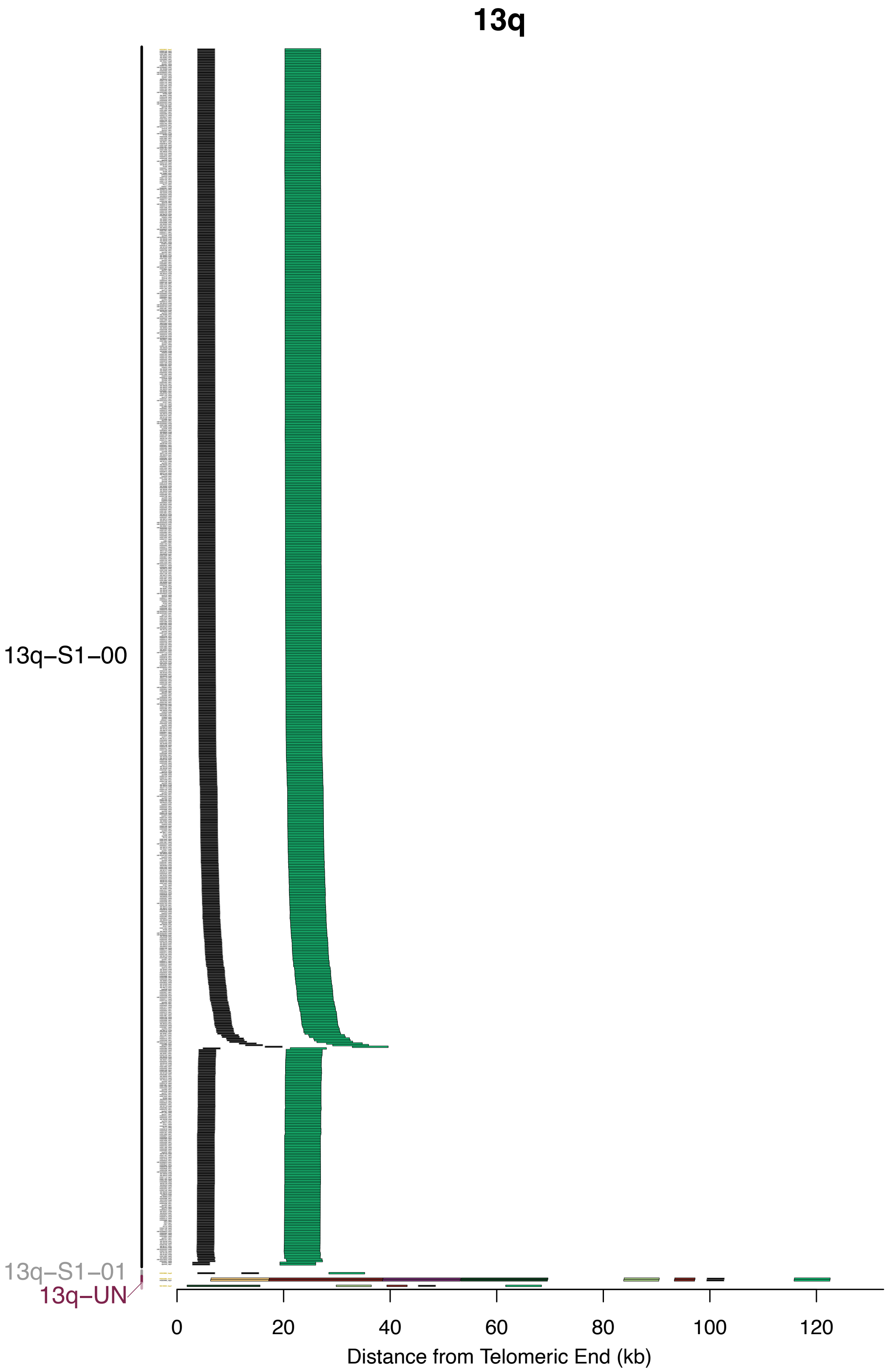


**Supplementary Fig. S30. Subtelomere contigs in 13q grouped based on paralogy block sequence.** Figure interpretation follows that of Supplementary Figure S9, but for chromosome arm 13q.

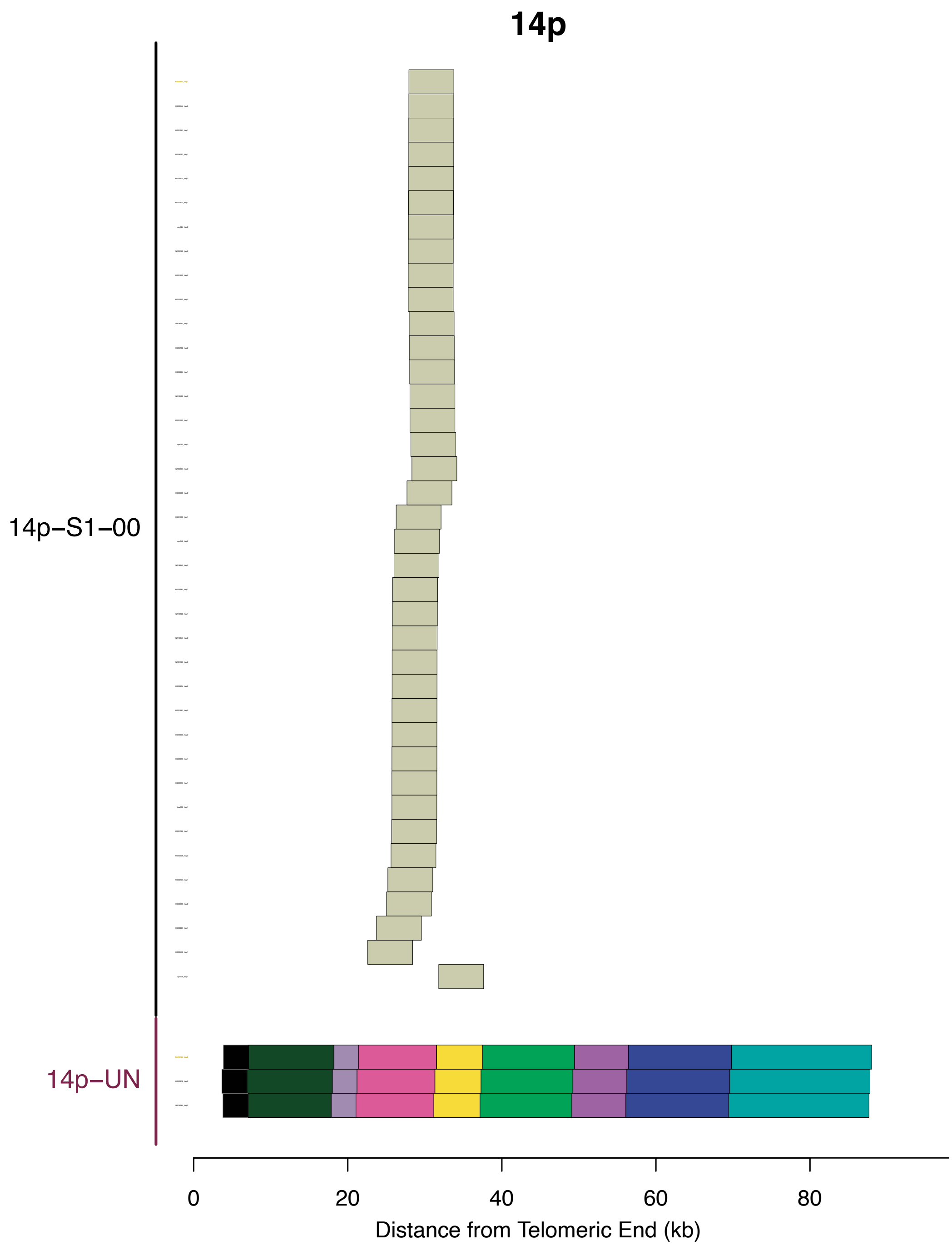


**Supplementary Fig. S31. Subtelomere contigs in 14p grouped based on paralogy block sequence.** Figure interpretation follows that of Supplementary Figure S9, but for chromosome arm 14p.

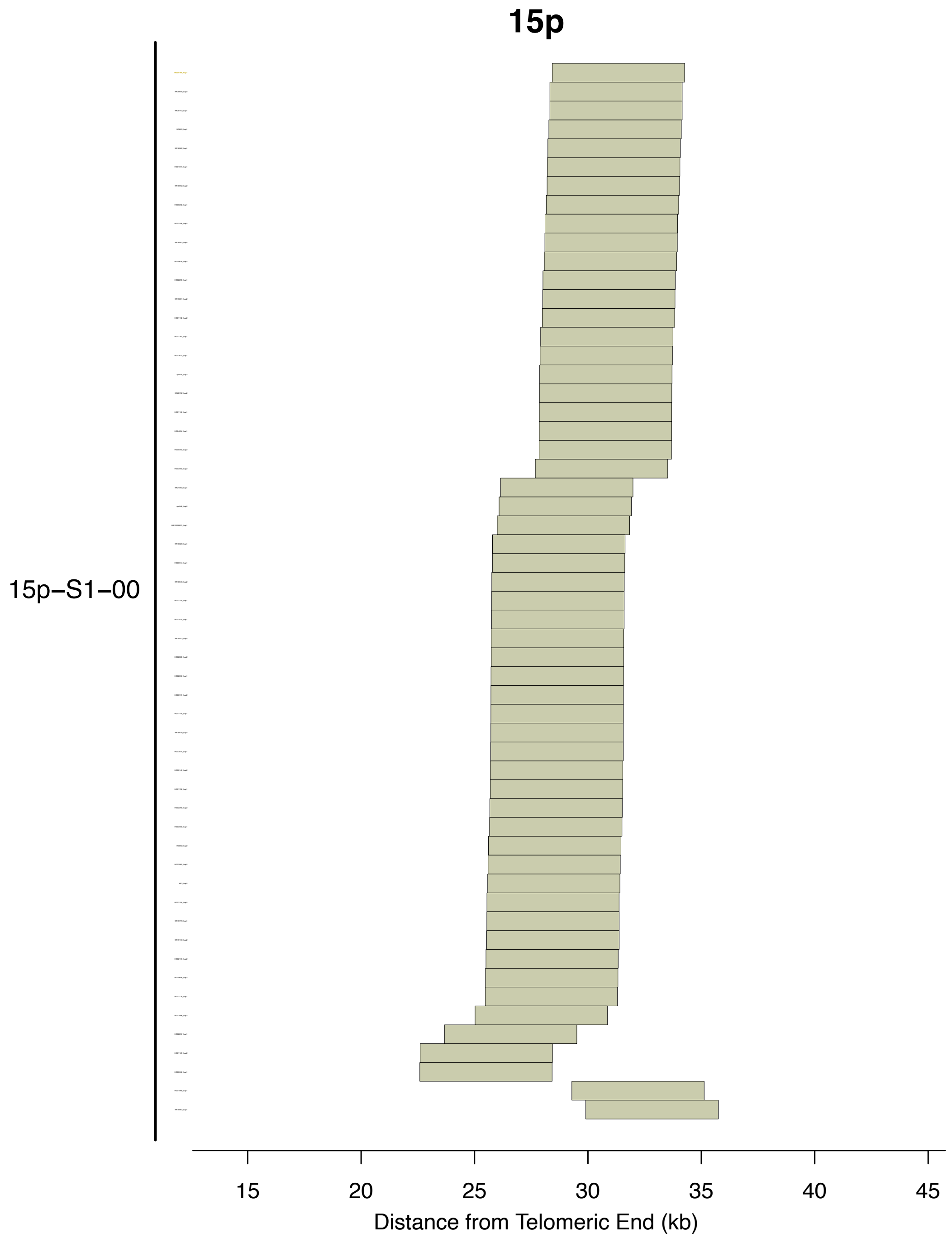


**Supplementary Fig. S32. Subtelomere contigs in 15p grouped based on paralogy block sequence.** Figure interpretation follows that of Supplementary Figure S9, but for chromosome arm 15p.

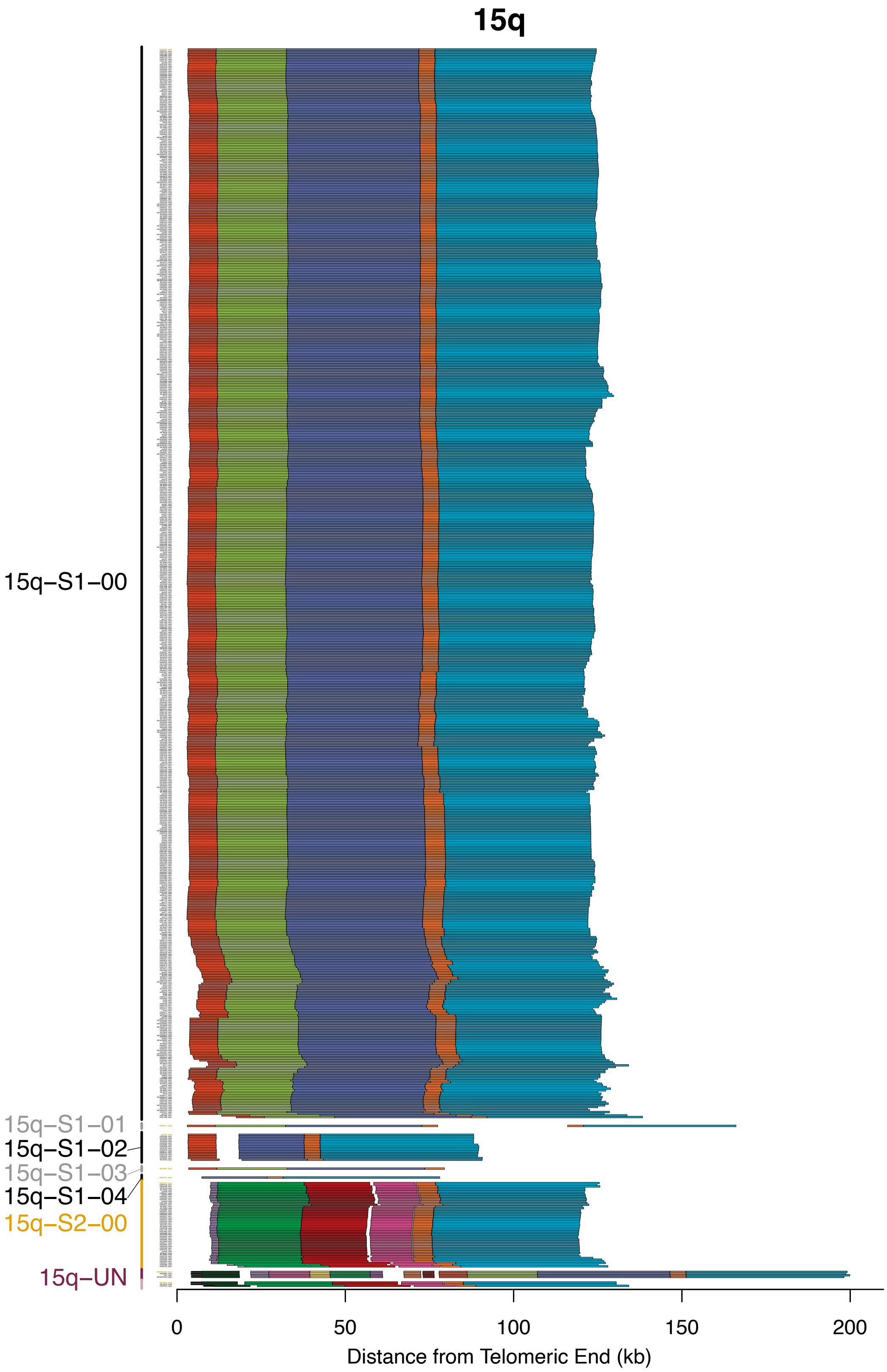


**Supplementary Fig. S33. Subtelomere contigs in 15q grouped based on paralogy block sequence.** Figure interpretation follows that of Supplementary Figure S9, but for chromosome arm 15q.

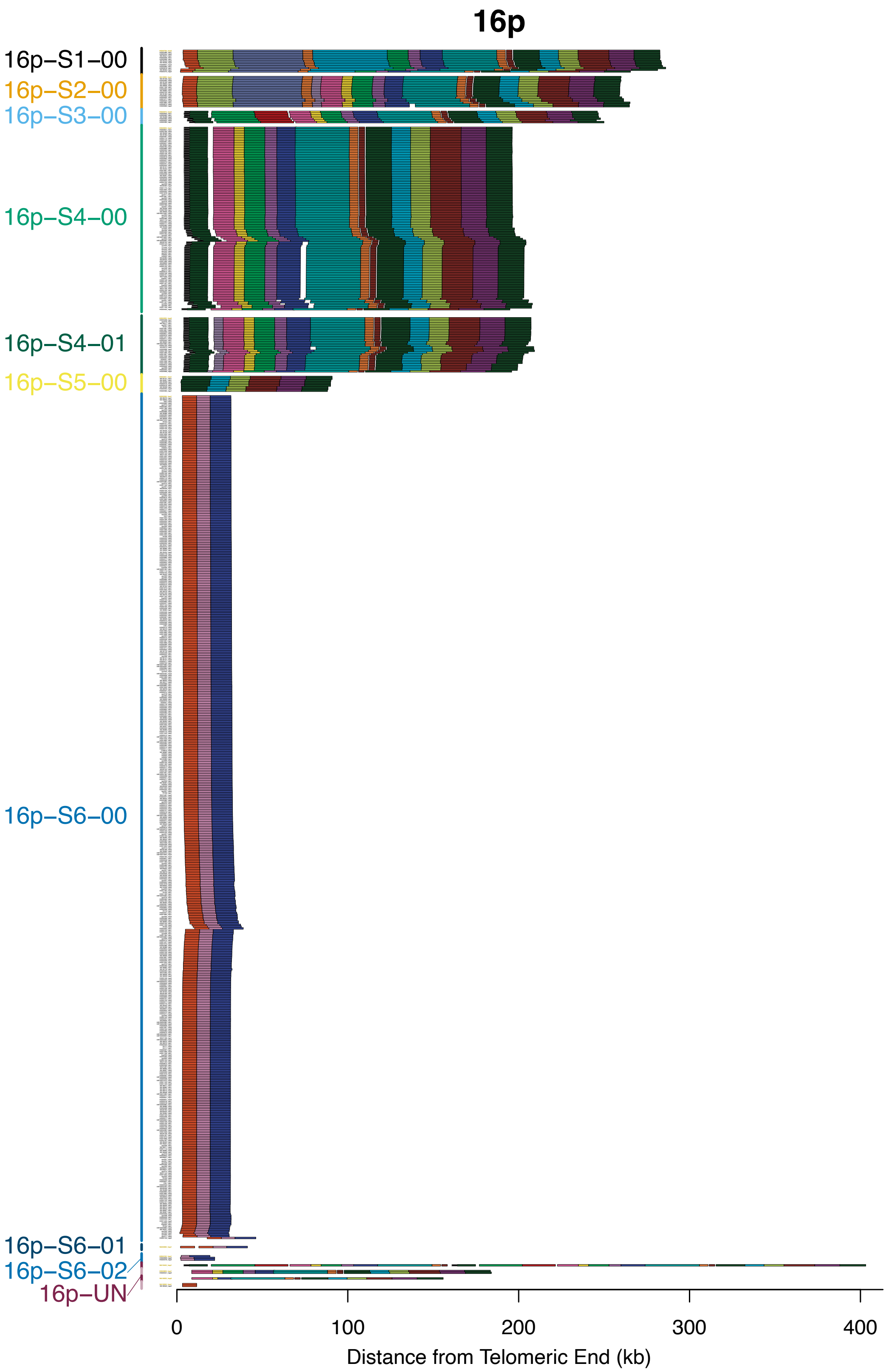


**Supplementary Fig. S34. Subtelomere contigs in 16p grouped based on paralogy block sequence.** Figure interpretation follows that of Supplementary Figure S9, but for chromosome arm 16p.

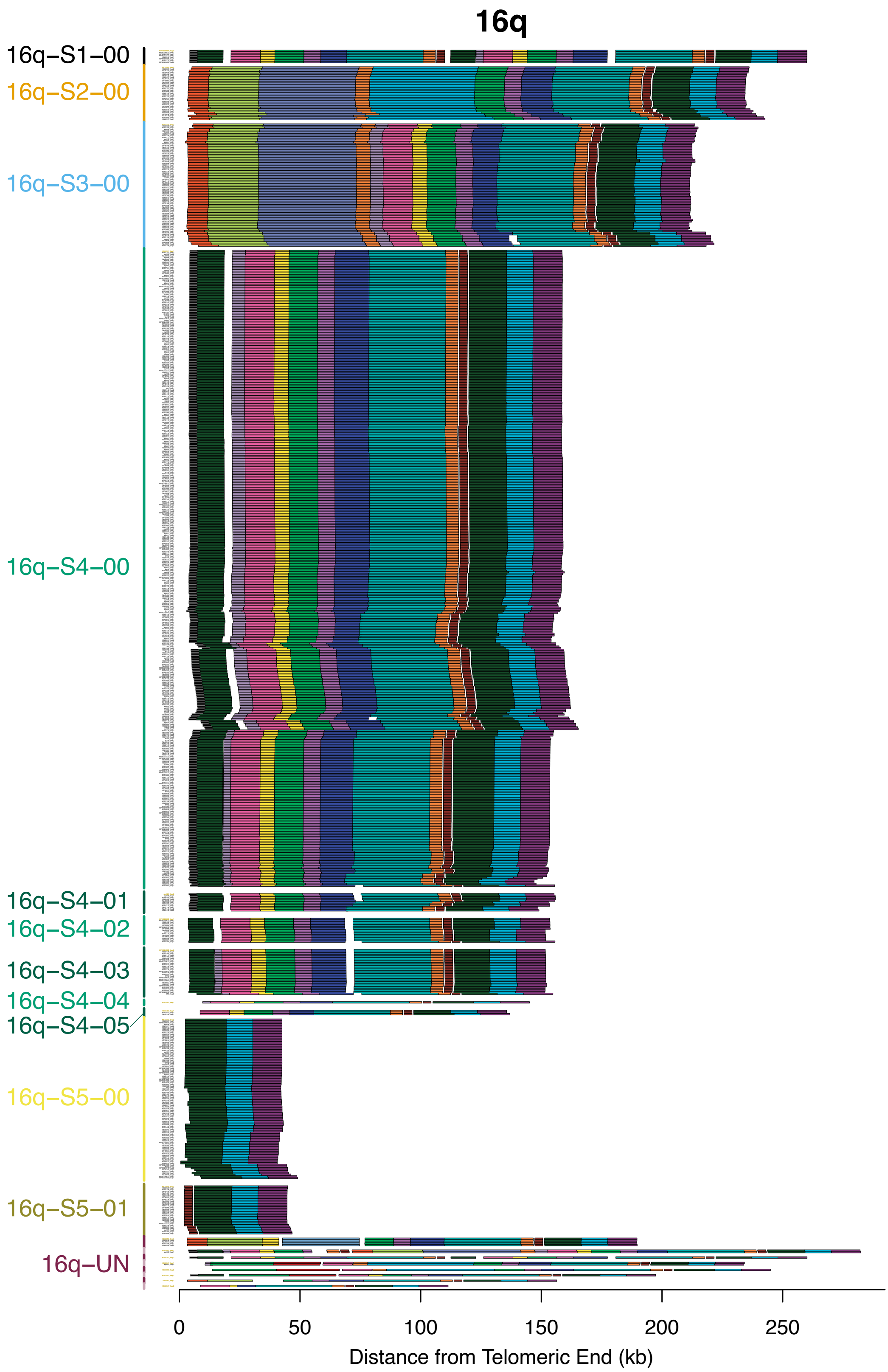


**Supplementary Fig. S35. Subtelomere contigs in 16q grouped based on paralogy block sequence.** Figure interpretation follows that of Supplementary Figure S9, but for chromosome arm 16q.

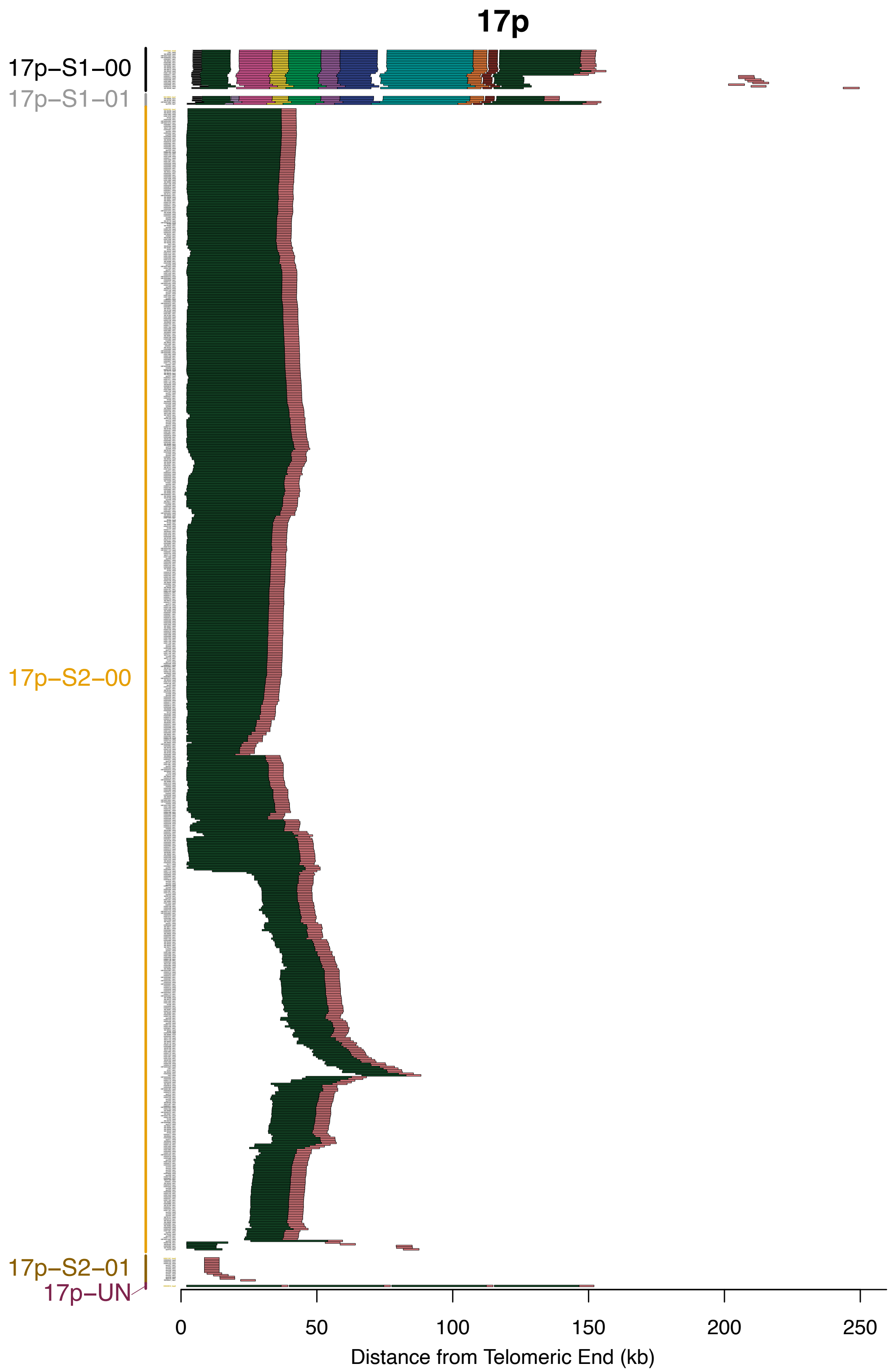


**Supplementary Fig. S36. Subtelomere contigs in 17p grouped based on paralogy block sequence.** Figure interpretation follows that of Supplementary Figure S9, but for chromosome arm 17p.

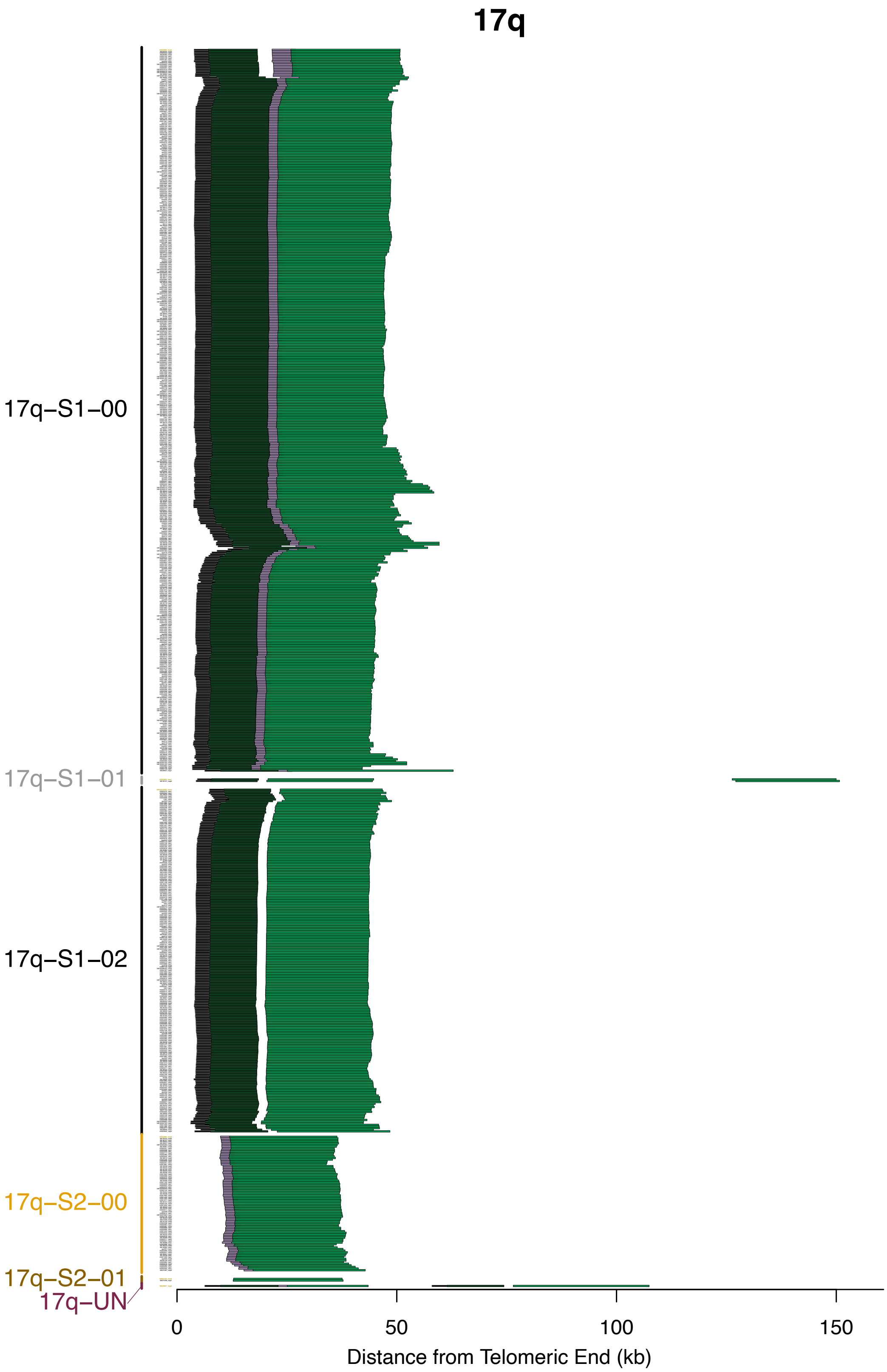


**Supplementary Fig. S37. Subtelomere contigs in 17q grouped based on paralogy block sequence.** Figure interpretation follows that of Supplementary Figure S9, but for chromosome arm 17q.

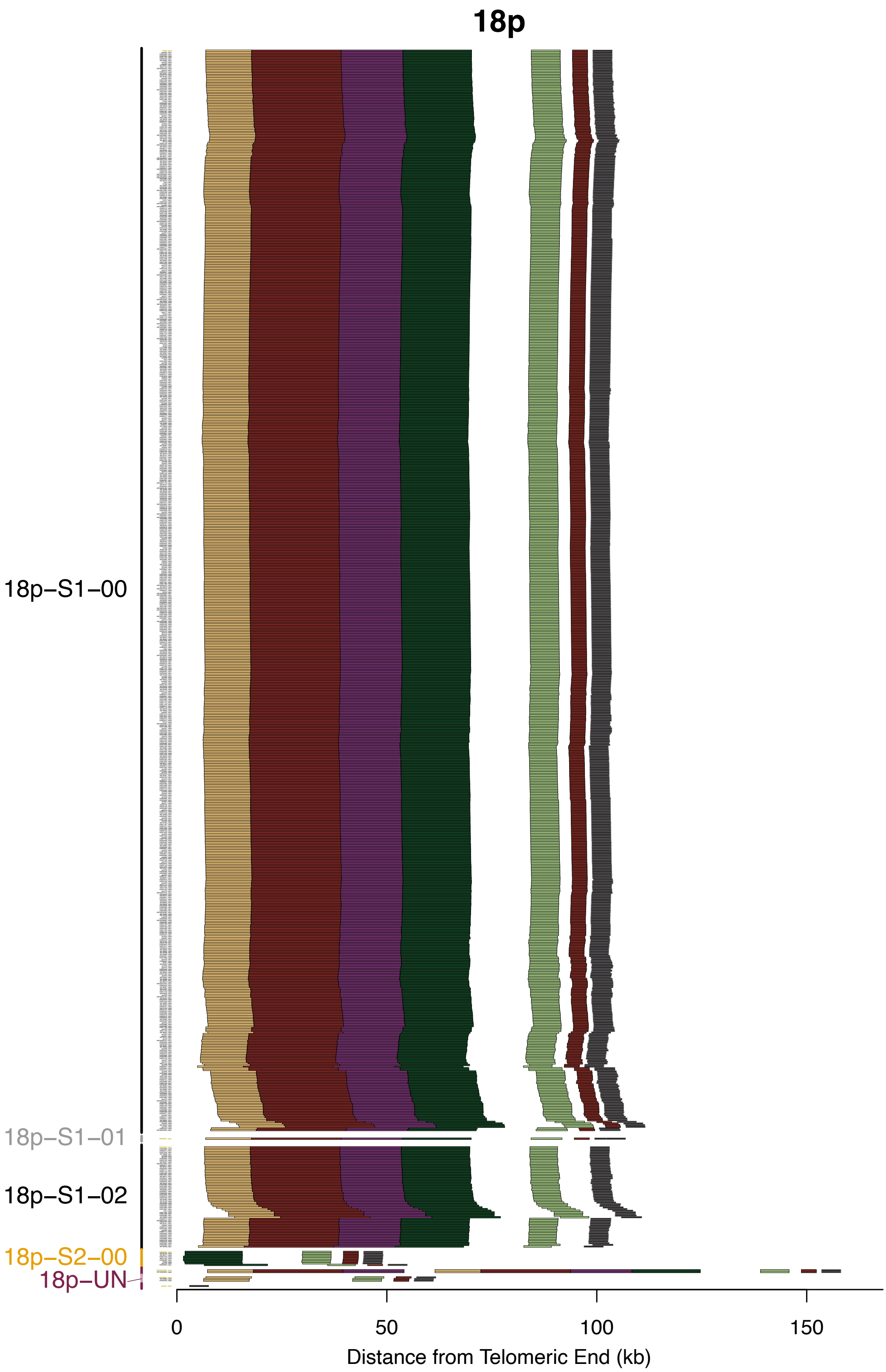


**Supplementary Fig. S38. Subtelomere contigs in 18p grouped based on paralogy block sequence.** Figure interpretation follows that of Supplementary Figure S9, but for chromosome arm 18p.

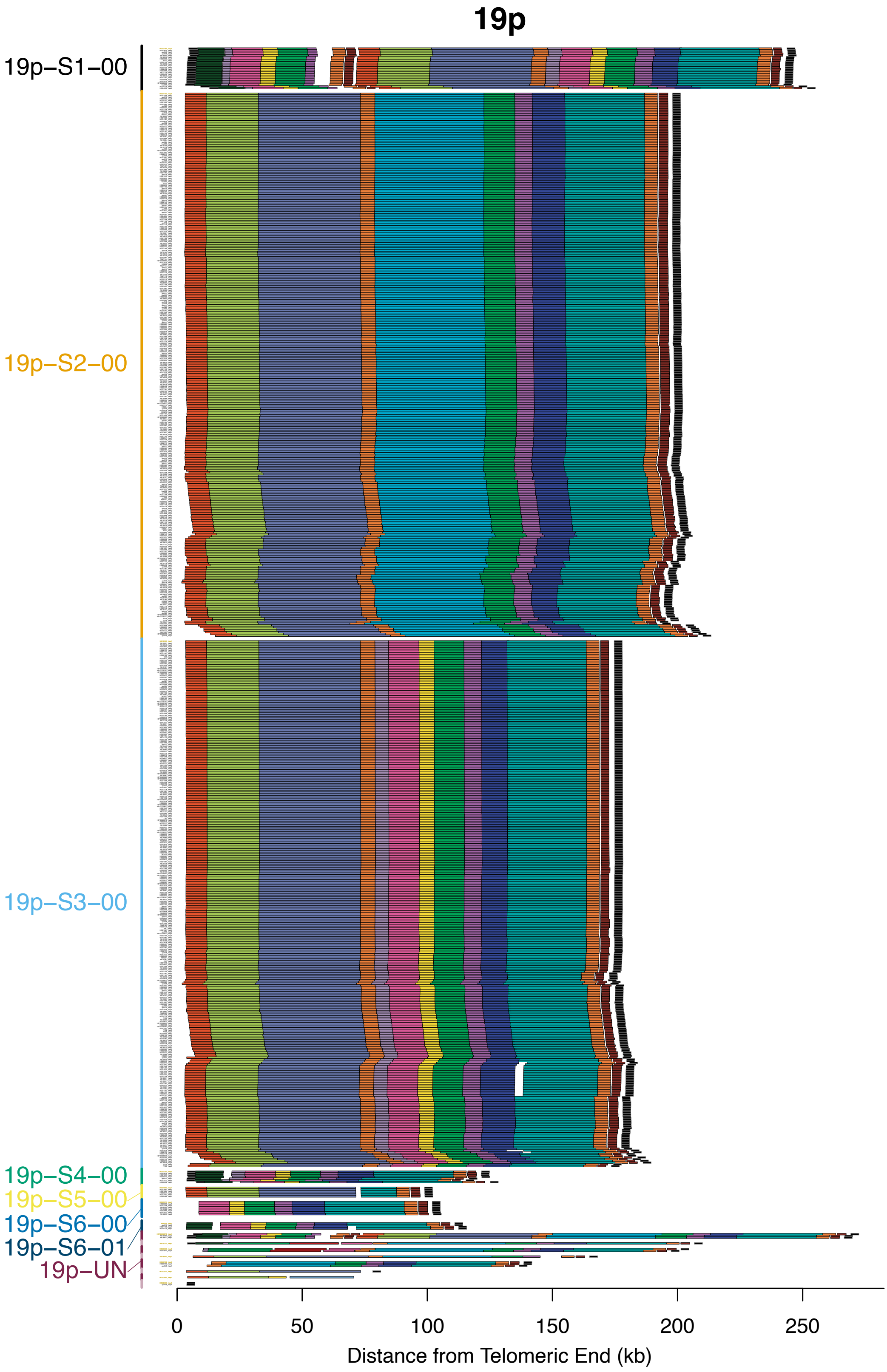


**Supplementary Fig. S39. Subtelomere contigs in 19p grouped based on paralogy block sequence.** Figure interpretation follows that of Supplementary Figure S9, but for chromosome arm 19p.

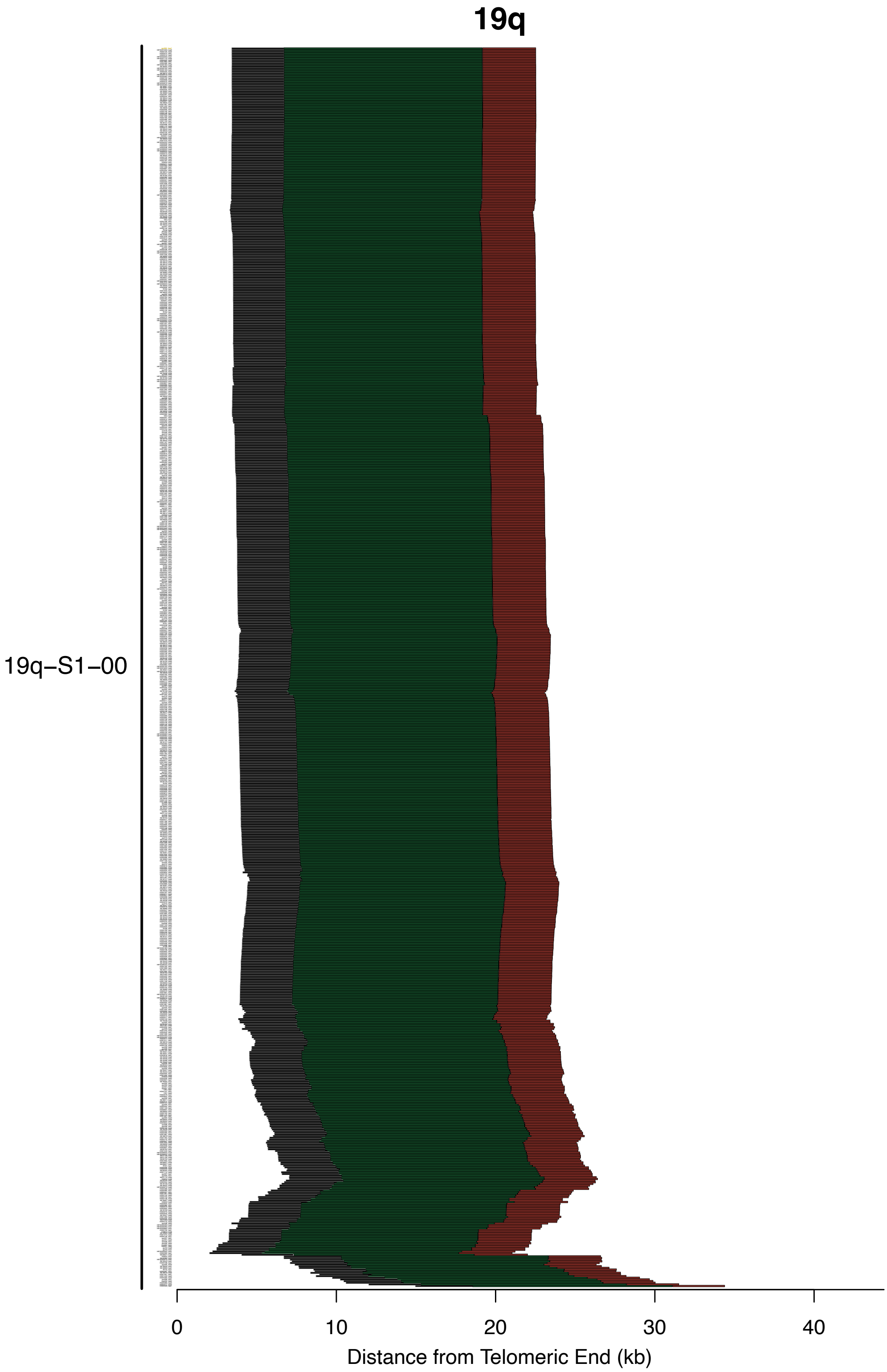


**Supplementary Fig. S40. Subtelomere contigs in 19q grouped based on paralogy block sequence.** Figure interpretation follows that of Supplementary Figure S9, but for chromosome arm 19q.

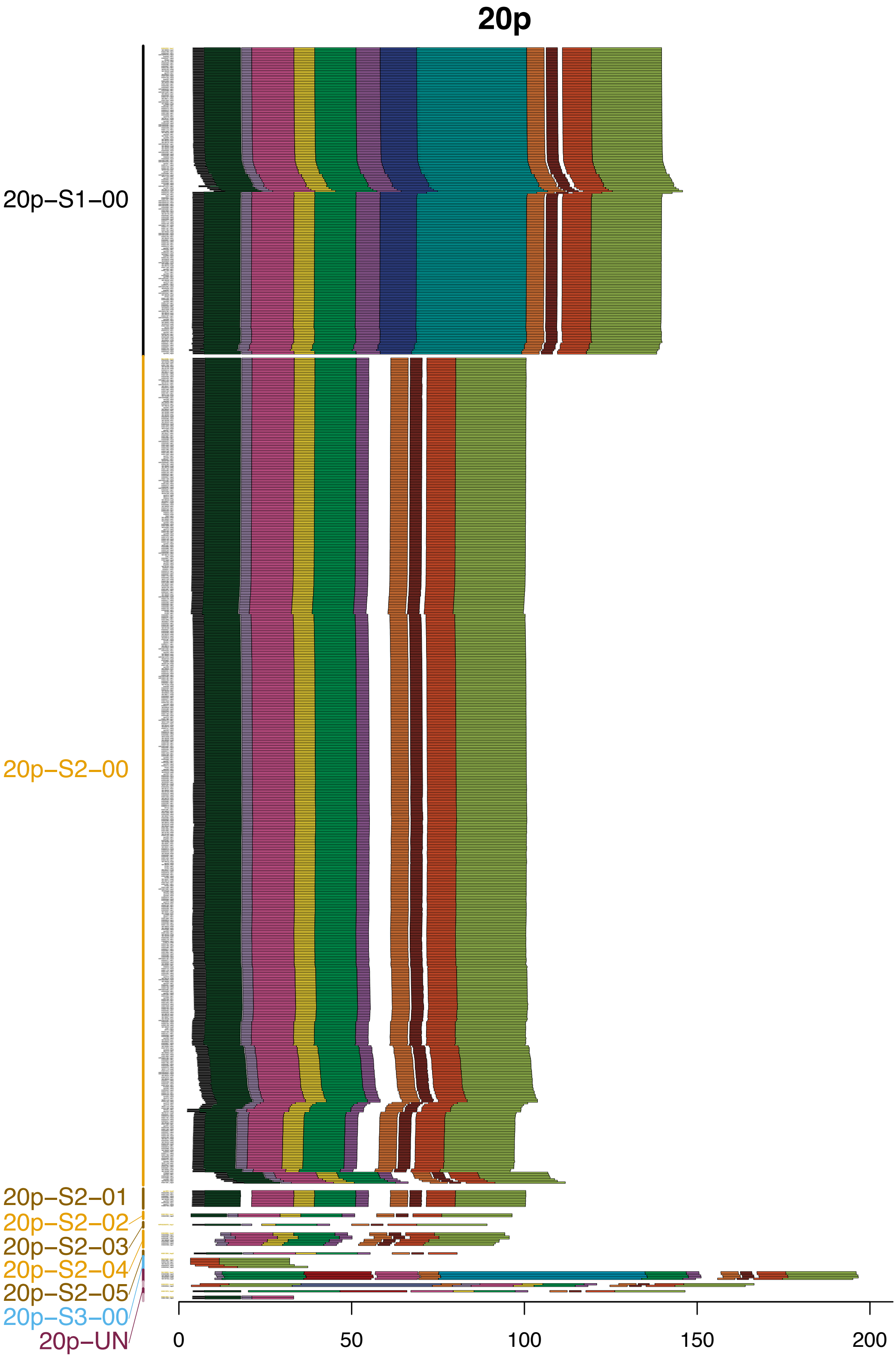


**Supplementary Fig. S41. Subtelomere contigs in 20p grouped based on paralogy block sequence.** Figure interpretation follows that of Supplementary Figure S9, but for chromosome arm 20p.

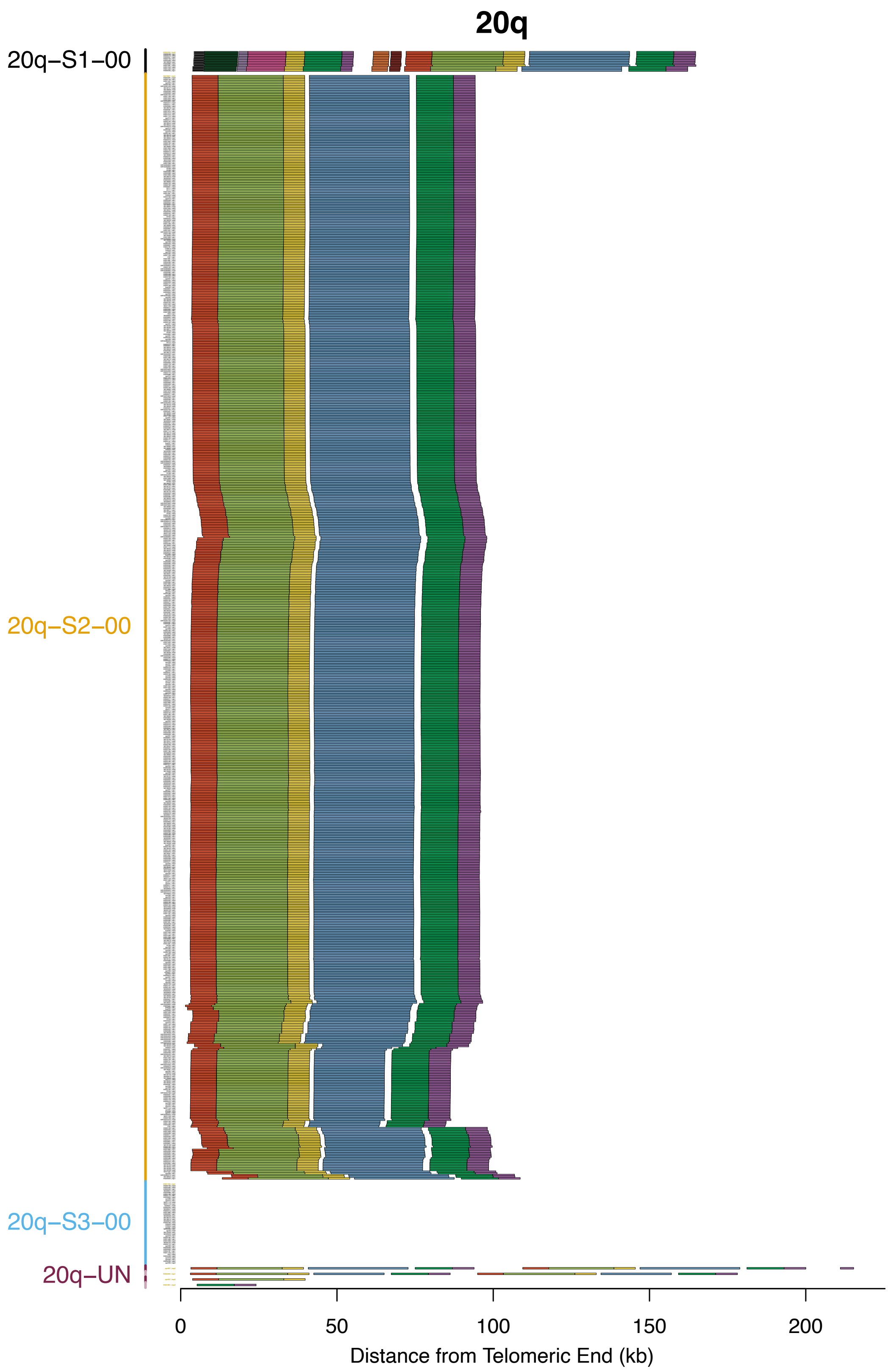


**Supplementary Fig. S42. Subtelomere contigs in 20q grouped based on paralogy block sequence.** Figure interpretation follows that of Supplementary Figure S9, but for chromosome arm 20q.

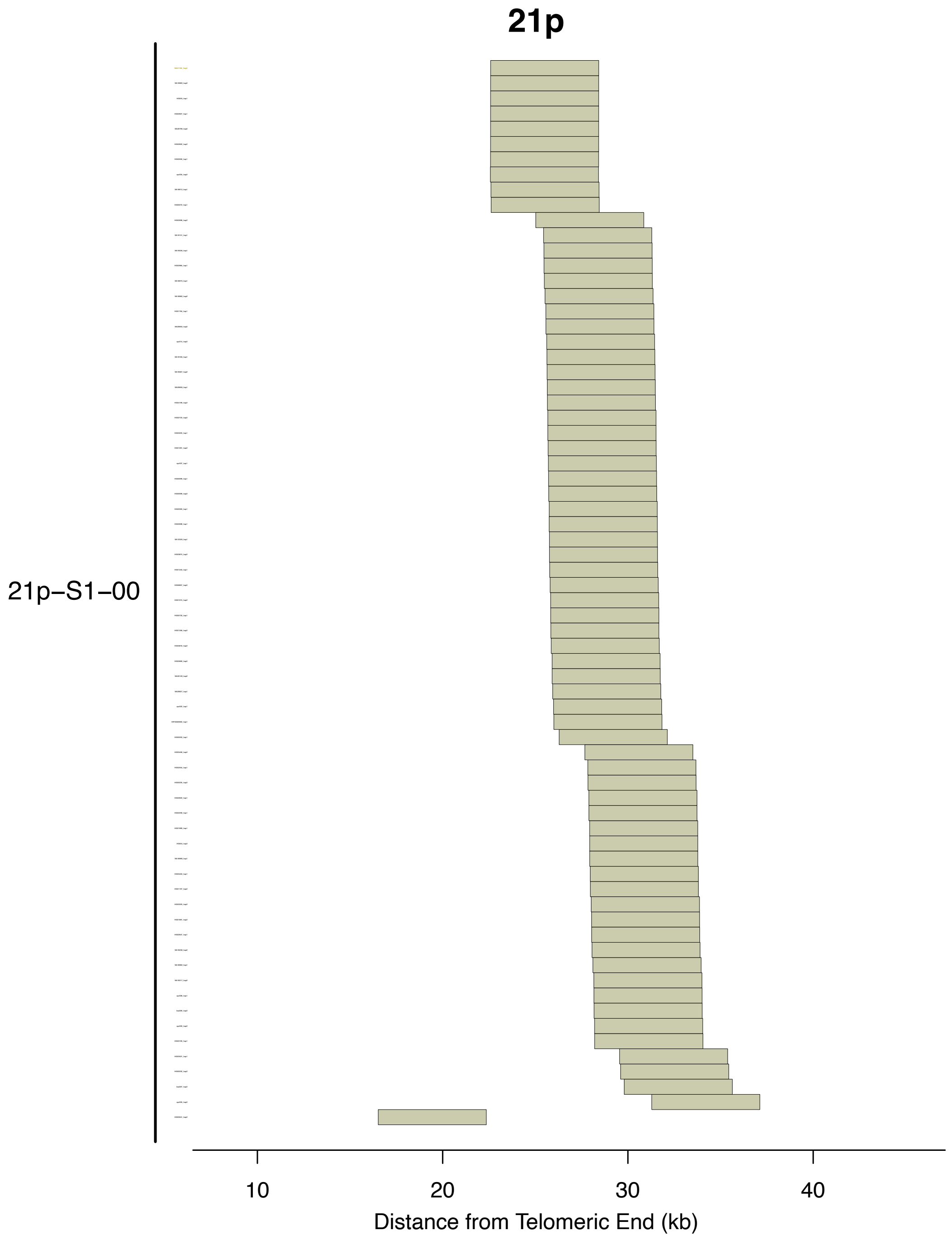


**Supplementary Fig. S43. Subtelomere contigs in 21p grouped based on paralogy block sequence.** Figure interpretation follows that of Supplementary Figure S9, but for chromosome arm 21p.

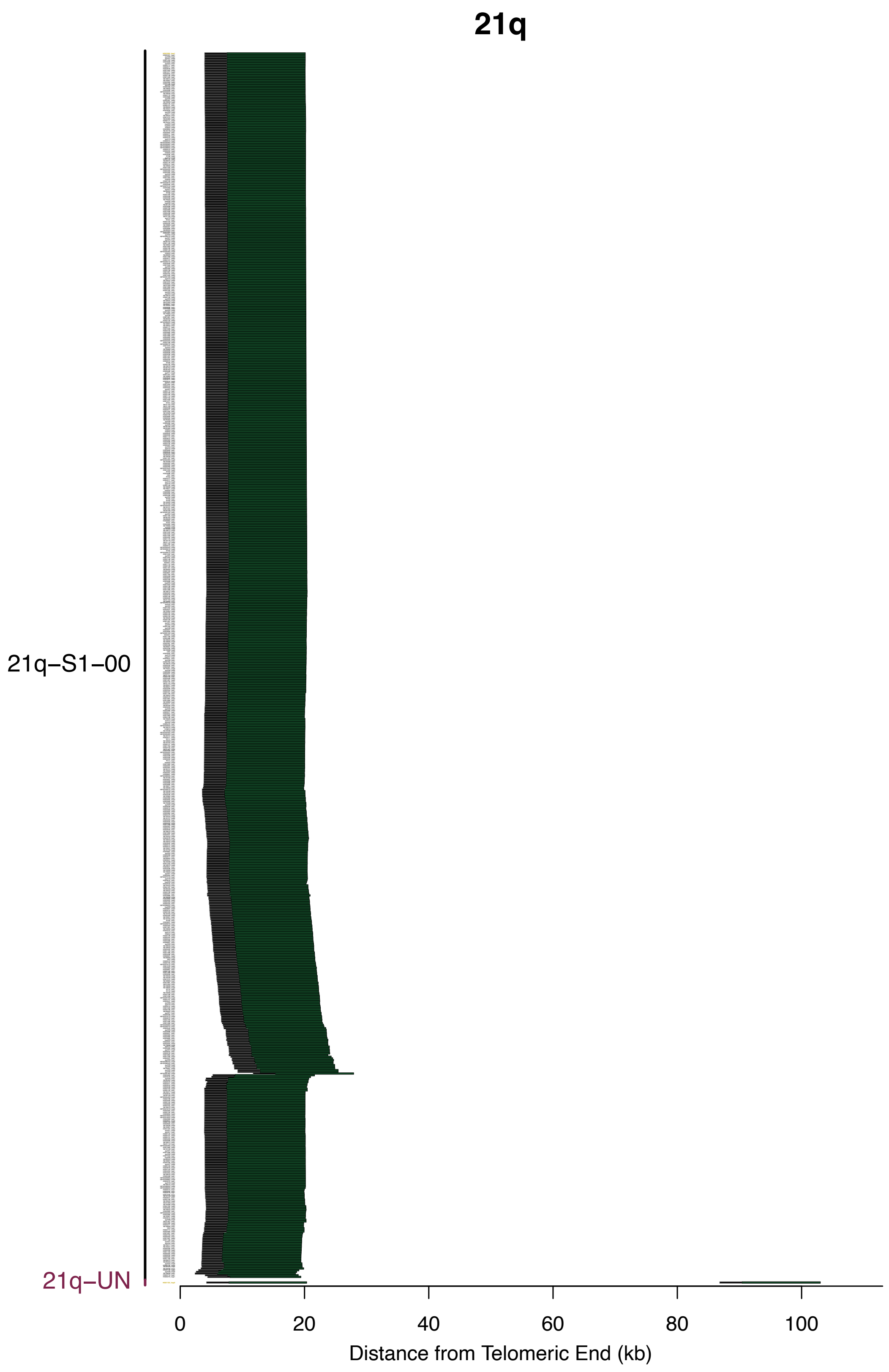


**Supplementary Fig. S44. Subtelomere contigs in 21q grouped based on paralogy block sequence.** Figure interpretation follows that of Supplementary Figure S9, but for chromosome arm 21q.

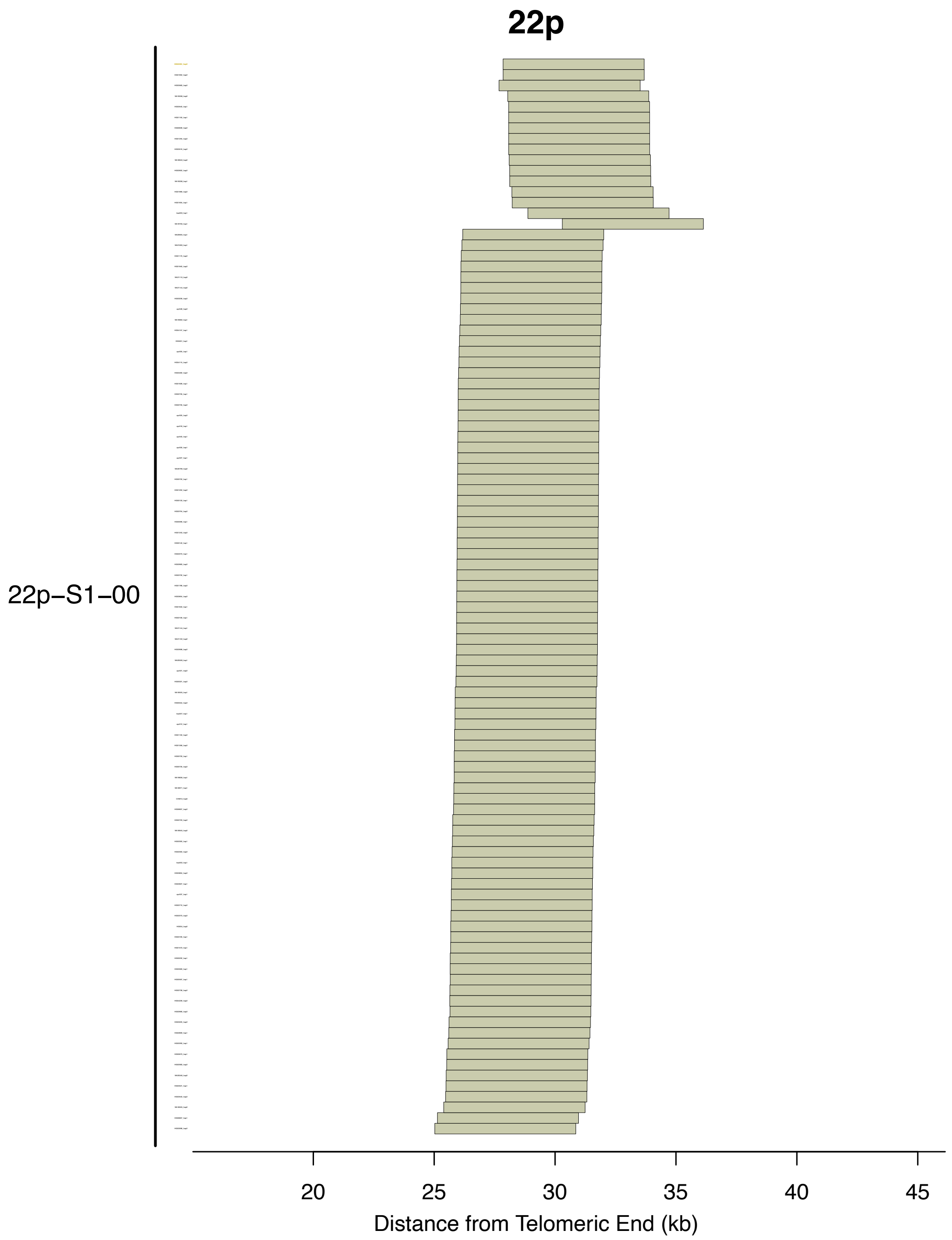


**Supplementary Fig. S45. Subtelomere contigs in 22p grouped based on paralogy block sequence.** Figure interpretation follows that of Supplementary Figure S9, but for chromosome arm 22p.

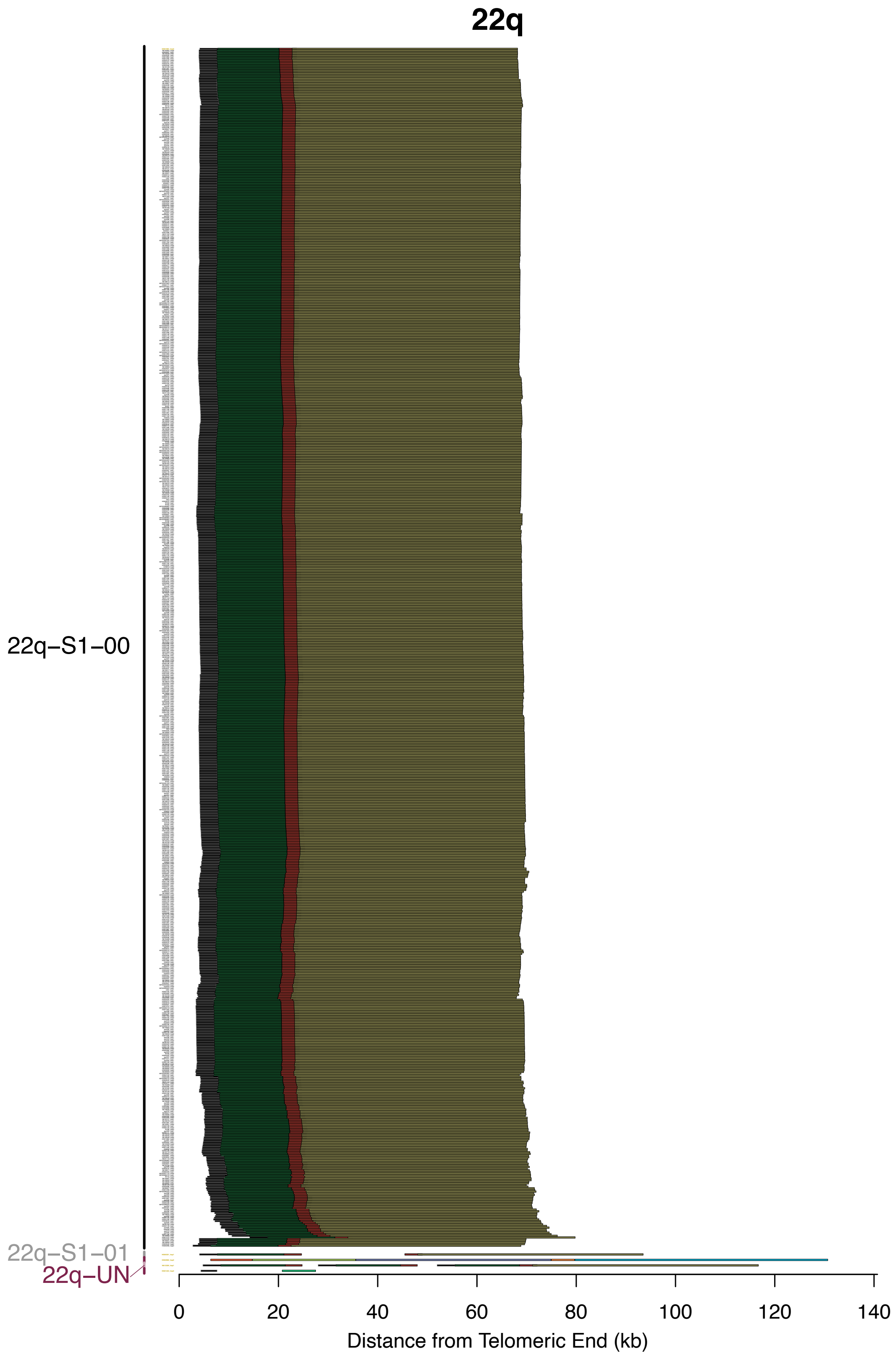


**Supplementary Fig. S46. Subtelomere contigs in 22q grouped based on paralogy block sequence.** Figure interpretation follows that of Supplementary Figure S9, but for chromosome arm 22q.

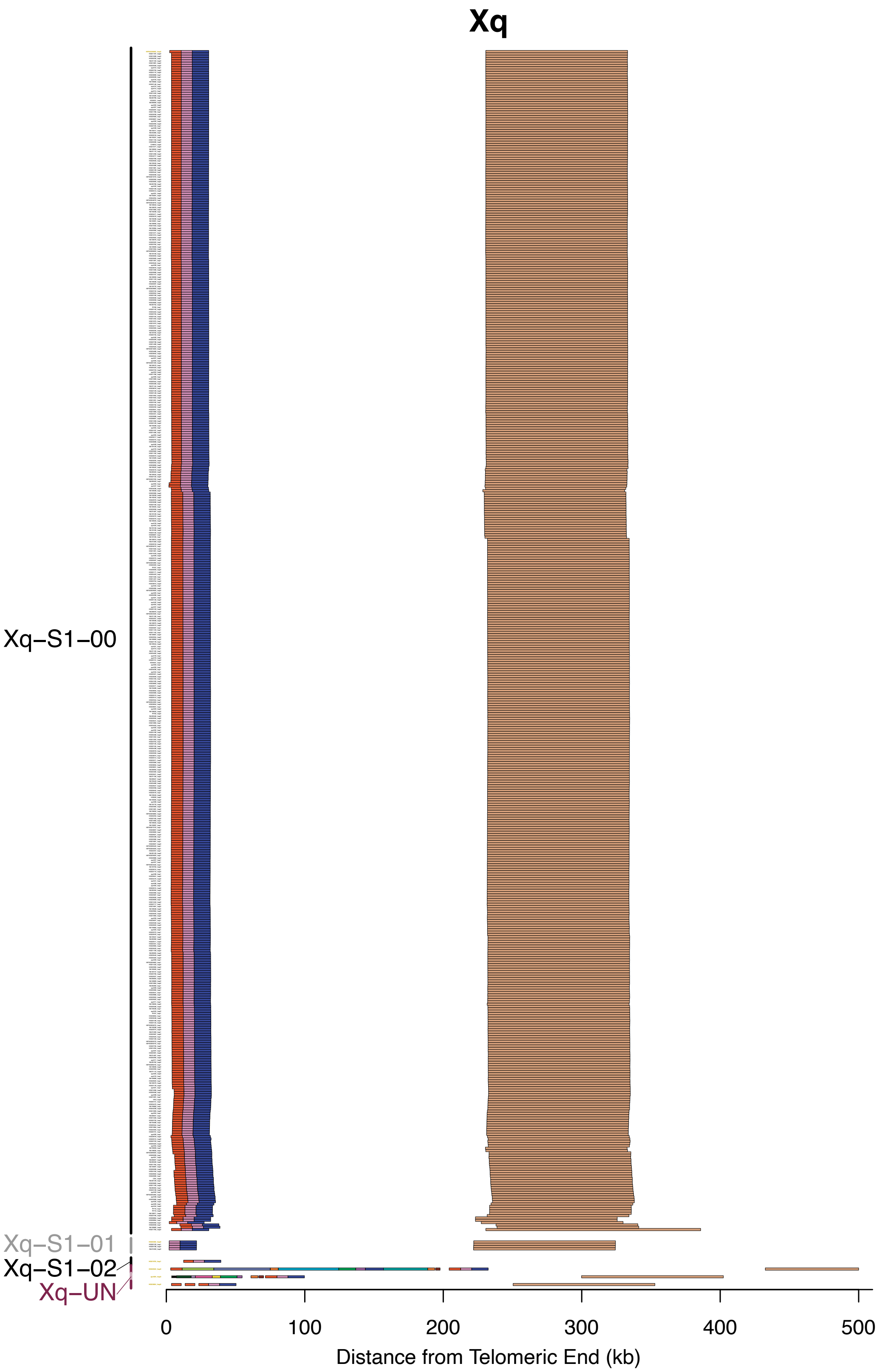


**Supplementary Fig. S47. Subtelomere contigs in Xq grouped based on paralogy block sequence.** Figure interpretation follows that of Supplementary Figure S9, but for chromosome arm Xq.

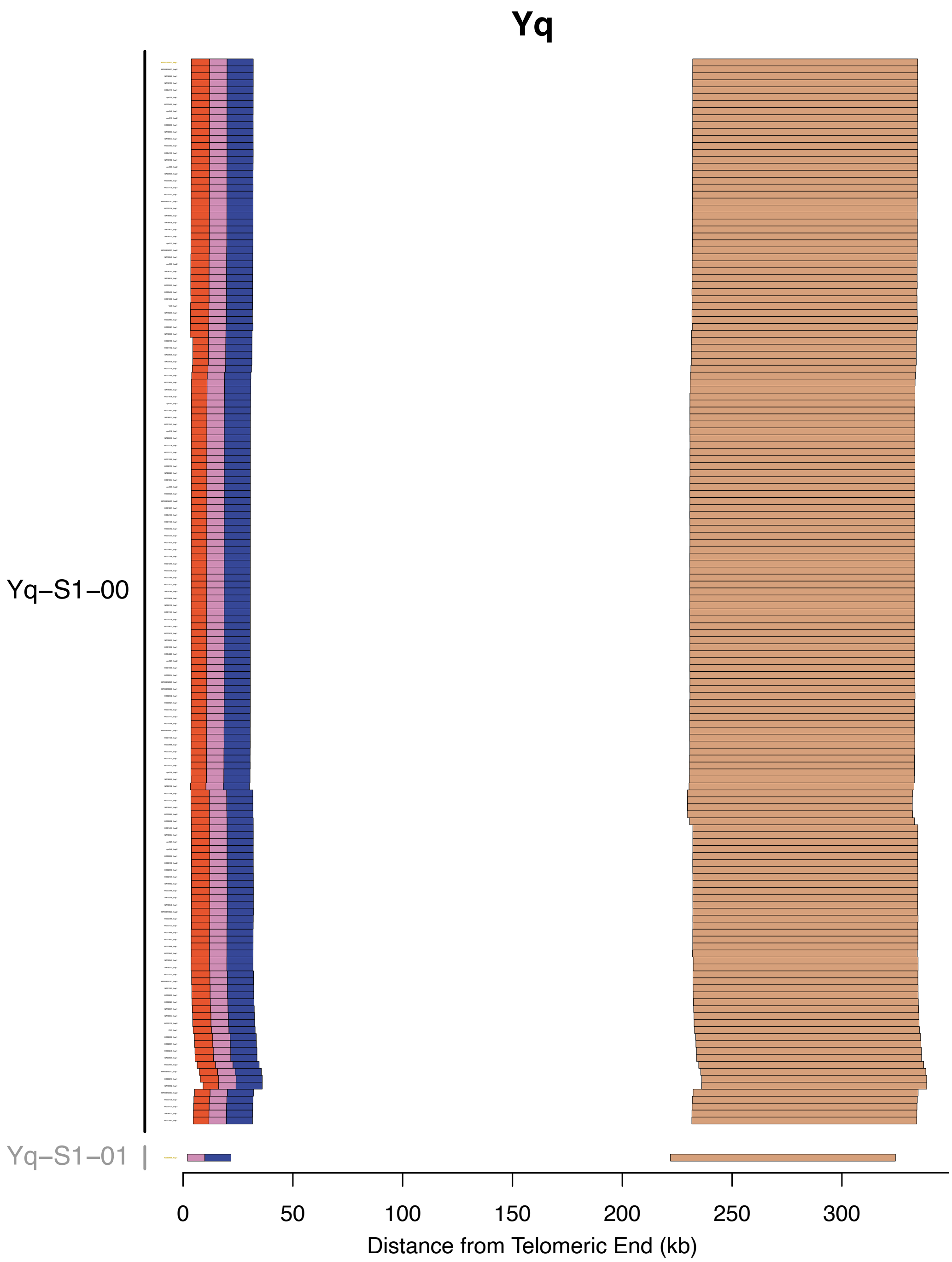


**Supplementary Fig. S48. Subtelomere contigs in Yq grouped based on paralogy block sequence.** Figure interpretation follows that of Supplementary Figure S9, but for chromosome arm Yq.

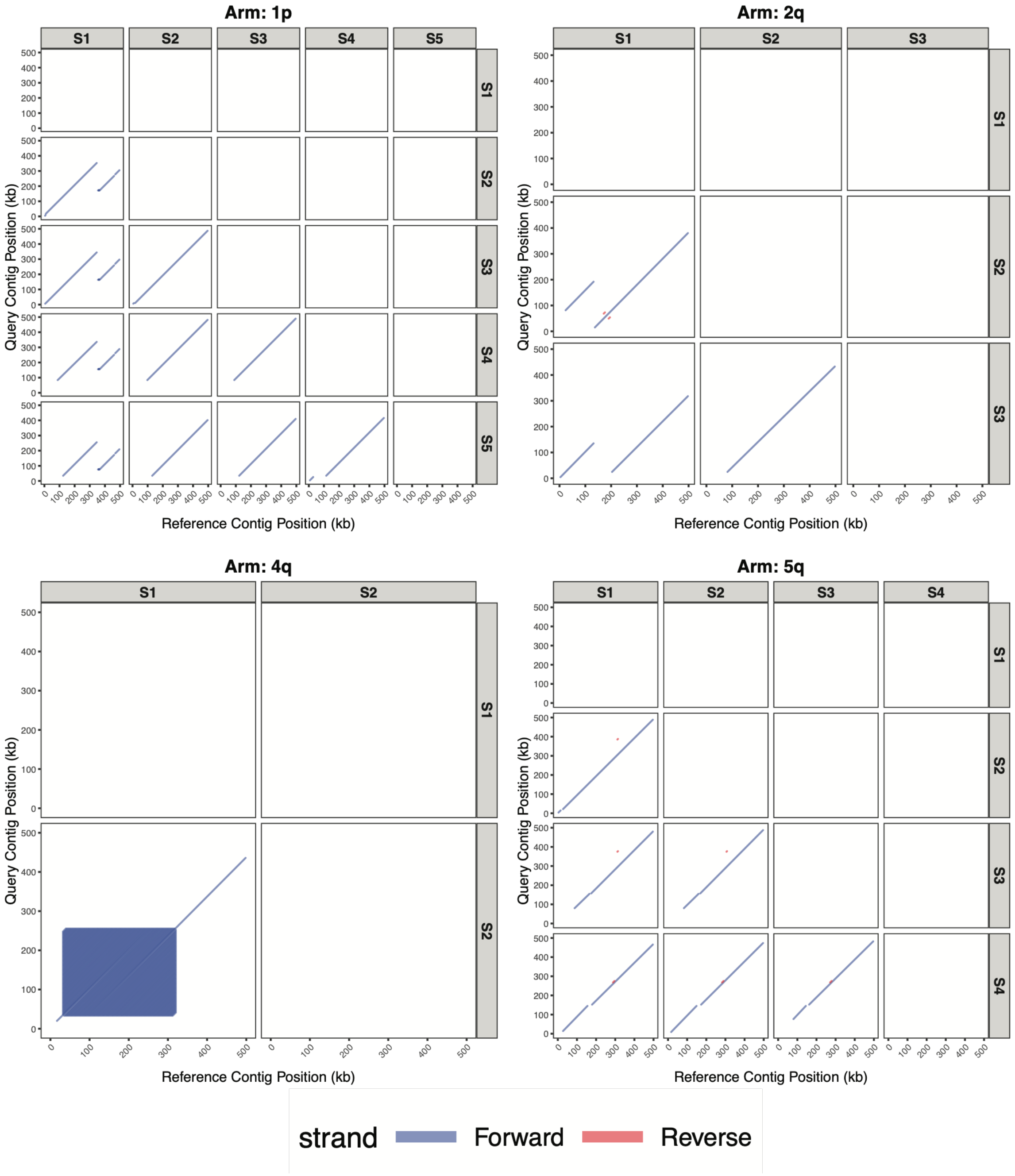


**Supplementary Fig. S49. Pairwise synteny dot-plot comparisons of representative contig sequences corresponding to each major haplotype, spanning chromosome arms 1p, 2q, 4q, and 5q.** Rows and columns denote major haplotypes S1 to S5. Each panel plots the reference contig position (kb, x-axis) against the query contig position (kb, y-axis), with forward-strand alignments shown as diagonal blue segments. Continuous diagonals indicate collinear alignment between contigs, while shorter offset segments reflect partial alignments or structural discontinuities between haplotype assemblies.

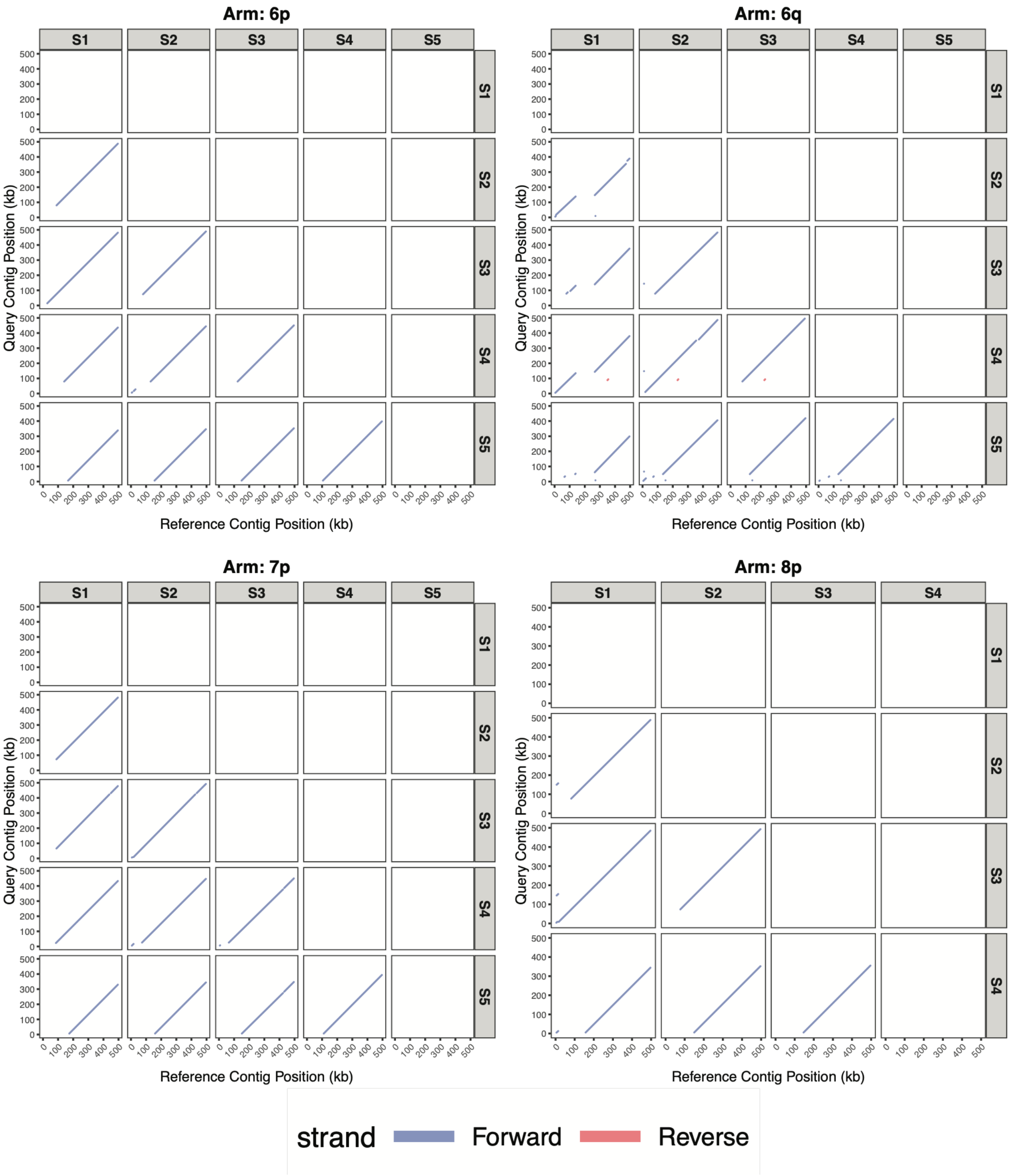


**Supplementary Fig. S50. Pairwise synteny dot-plot comparisons of representative contig sequences corresponding to each major haplotype, spanning chromosome arms 6p, 6q, 7p, and 8p.** Figure interpretation follows that of Supplementary Figure S49, but for chromosome arms 6p, 6q, 7p and 8p.

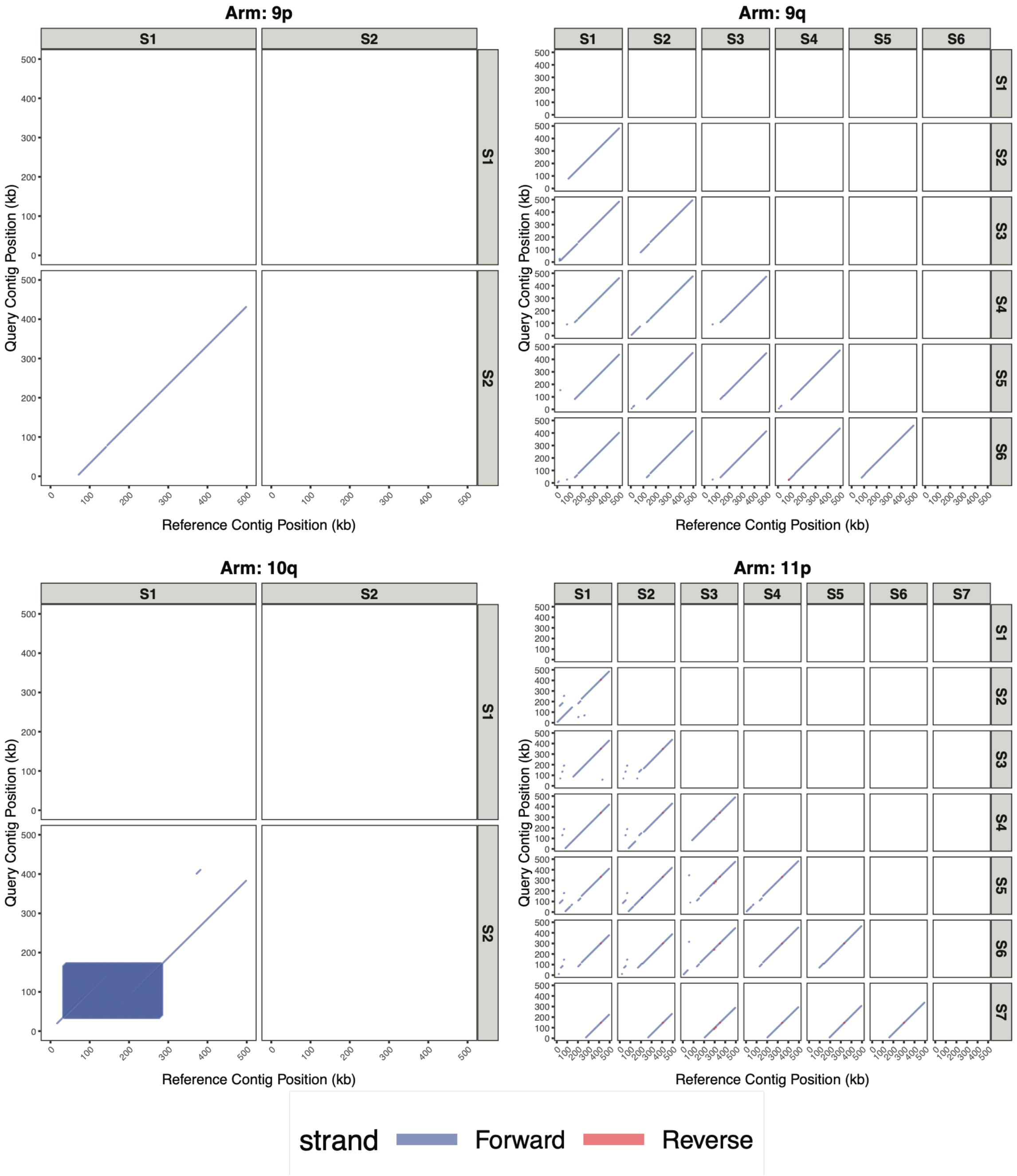


**Supplementary Fig. S51. Pairwise synteny dot-plot comparisons of representative contig sequences corresponding to each major haplotype, spanning chromosome arms 9p, 9q, 10q, and 11p.** Figure interpretation follows that of Supplementary Figure S49, but for chromosome arms 9p, 9q, 10q and 11p.

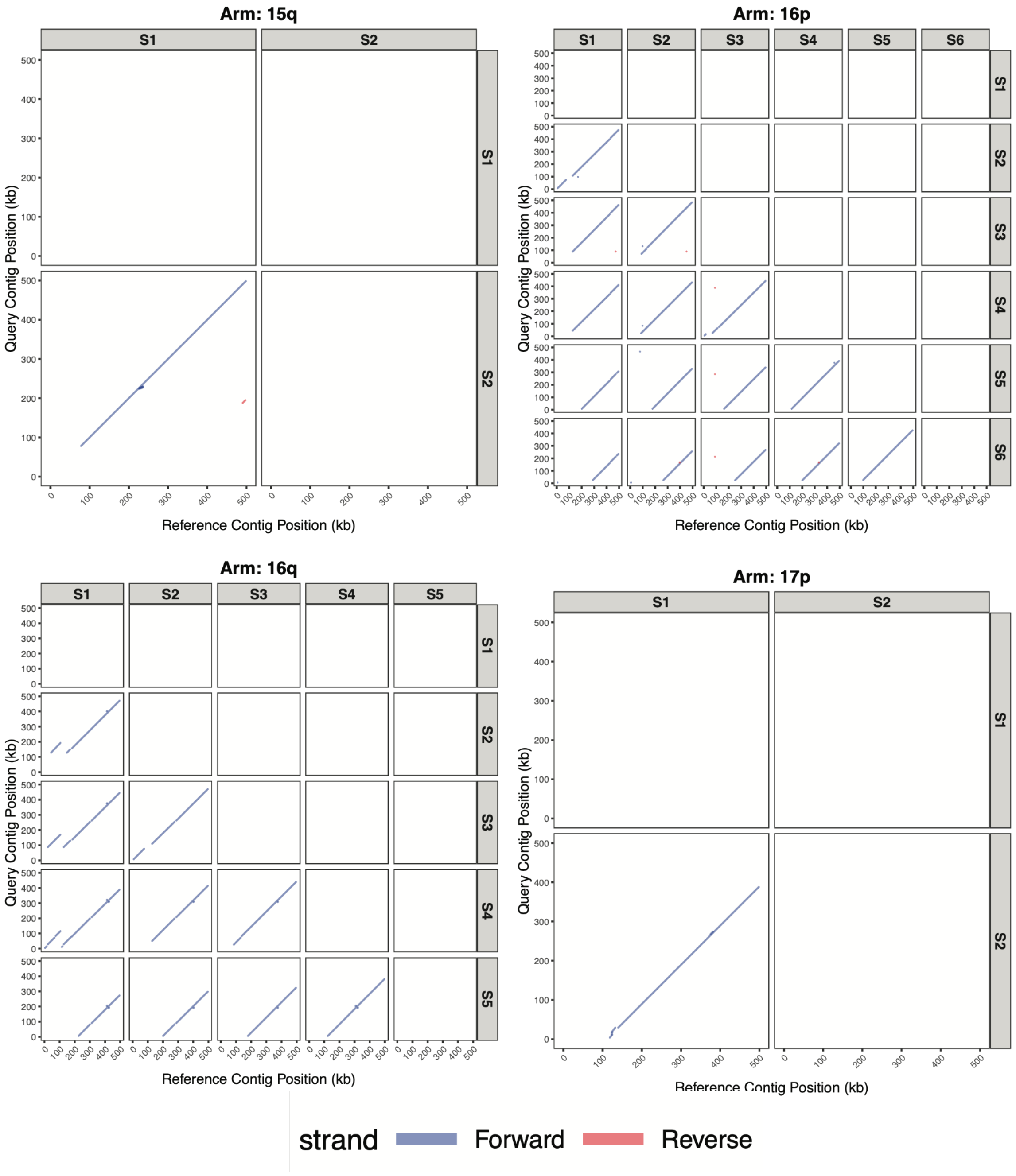


**Supplementary Fig. S52. Pairwise synteny dot-plot comparisons of representative contig sequences corresponding to each major haplotype, spanning chromosome arms 15q, 16p, 16q, and 17p.** Figure interpretation follows that of Supplementary Figure S49, but for chromosome arms 15q, 16p, 16q and 17p.

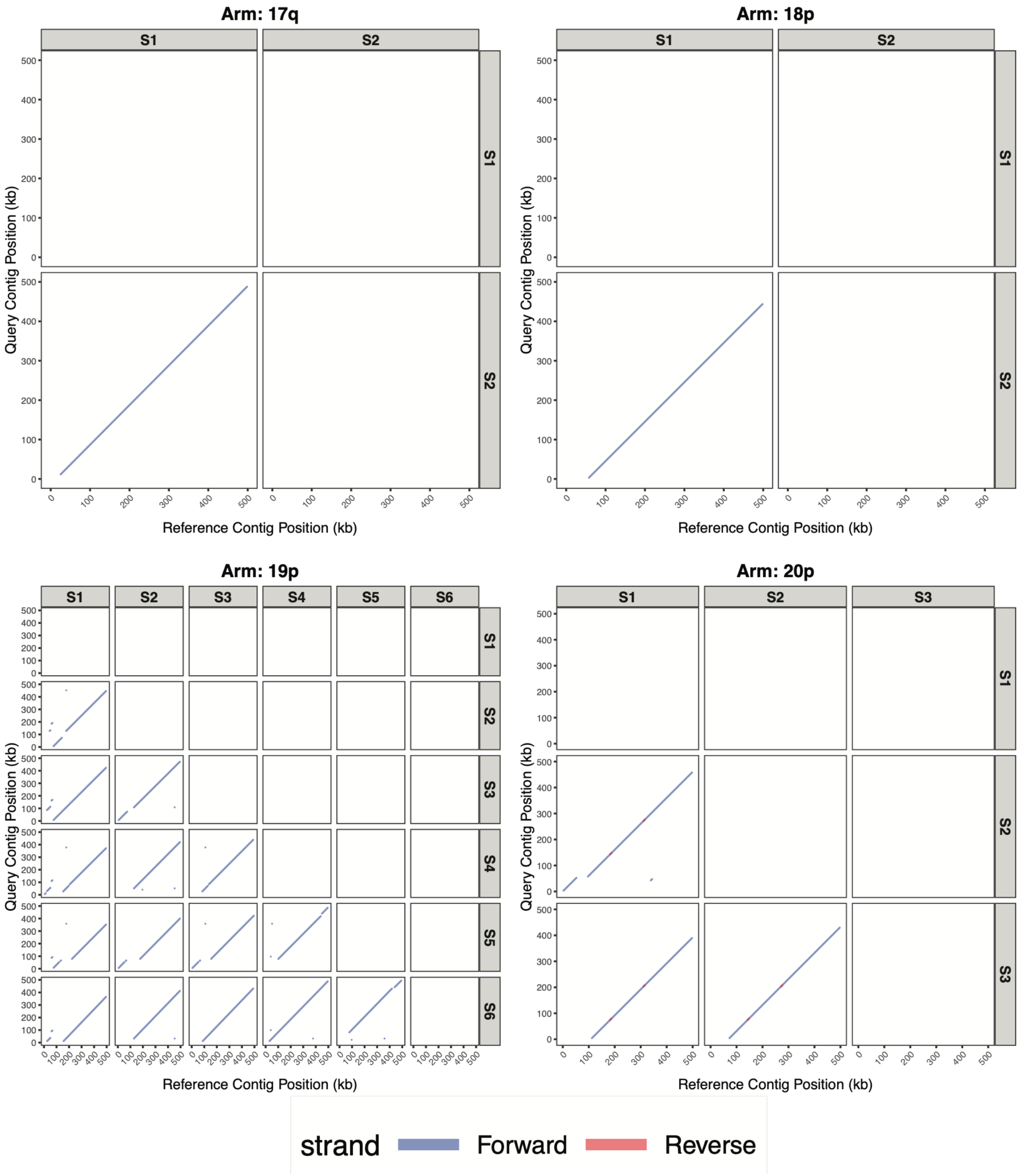


**Supplementary Fig. S53. Pairwise synteny dot-plot comparisons of representative contig sequences corresponding to each major haplotype, spanning chromosome arms 17q, 18p, 19p, and 20p.** Figure interpretation follows that of Supplementary Figure S49, but for chromosome arms 17q, 18p, 19p and 20p.

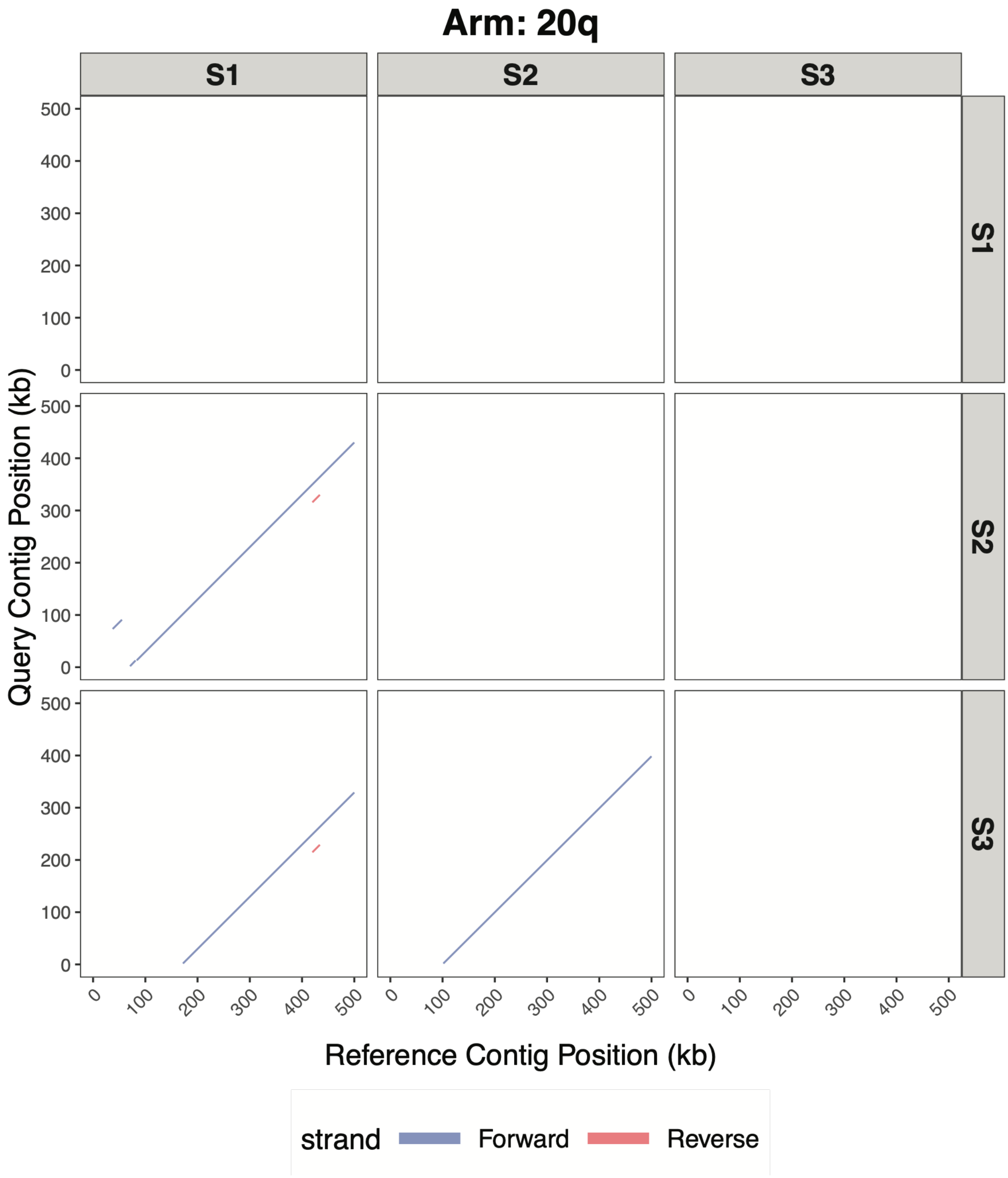


**Supplementary Fig. S54. Pairwise synteny dot-plot comparisons of representative contig sequences corresponding to each major haplotype, spanning chromosome arm 20q.** Figure interpretation follows that of Supplementary Figure S49, but for chromosome arm 20q.

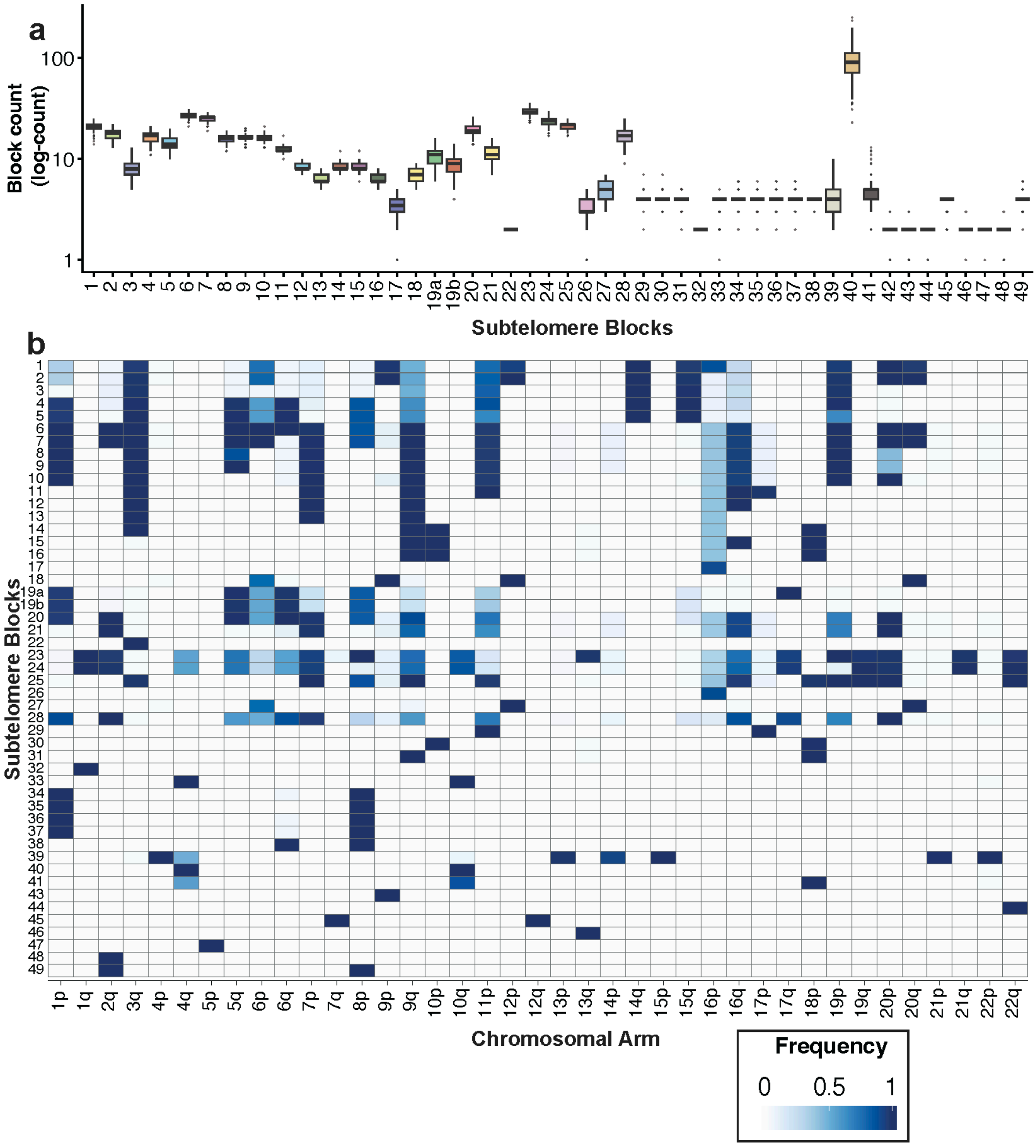


**Supplementary Fig. S55. Subtelomeric block distribution across chromosomal arms.** (a) Log-scaled boxplot of block counts per subtelomeric block derived from the HPRC cohort (n = 211) using complete contigs. Each box represents the distribution of copy number observed across haplotypes for a given subtelomeric block, with outliers shown as individual points. (a) Frequency heatmap of subtelomeric block presence across chromosomal arms in the full population cohort, where cell colour intensity reflects the proportion of haplotypes carrying each block at a given chromosomal arm.

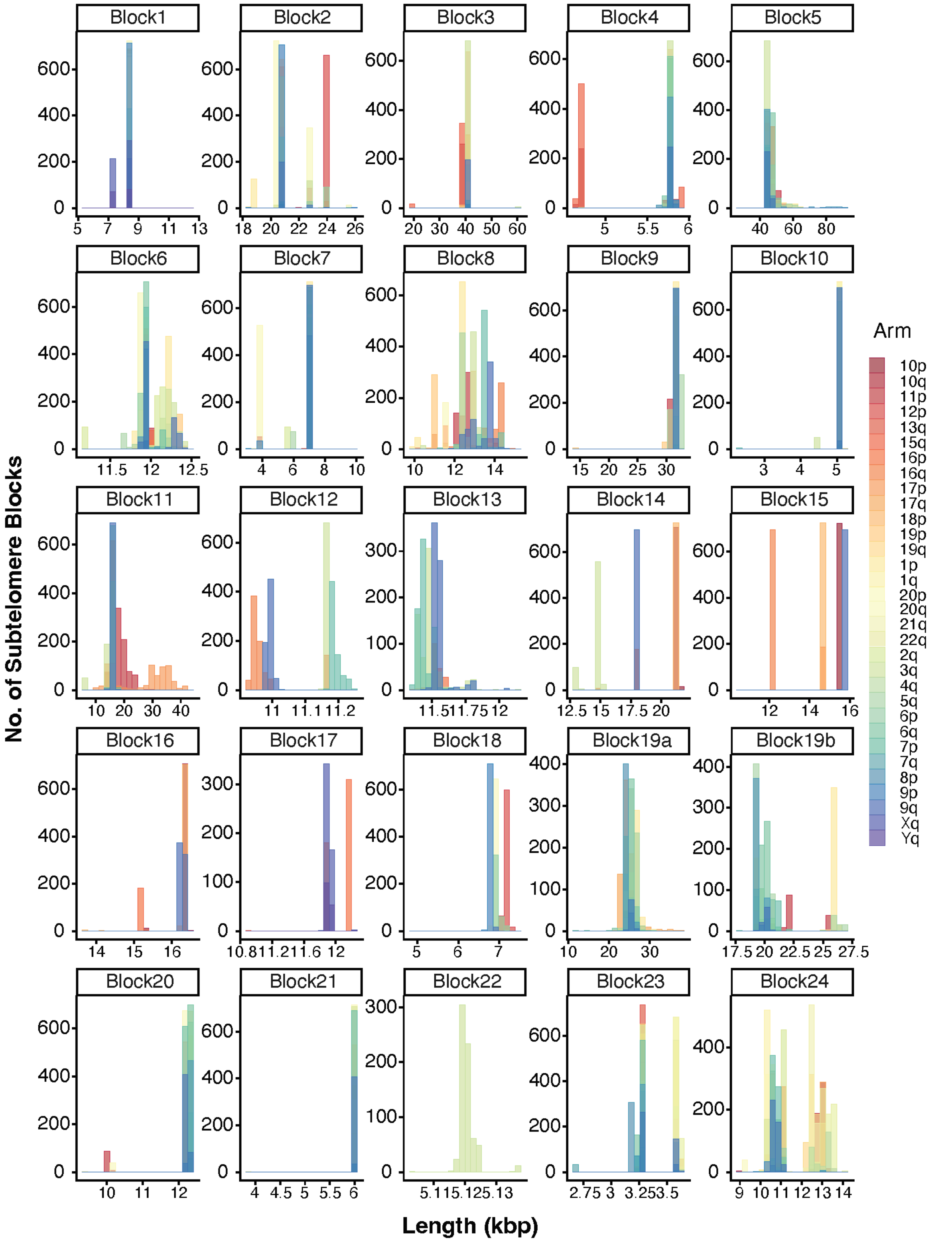


**Supplementary Fig. S56. Subtelomeric block length distributions (I)** Histograms of sequence length (kbp) for each subtelomeric block (Block 1 to Block 24) across the population, coloured by chromosomal arm of origin. Each panel represents a single block, with the x-axis showing sequence length in kilobase pairs and the y-axis showing the count of subtelomere blocks.

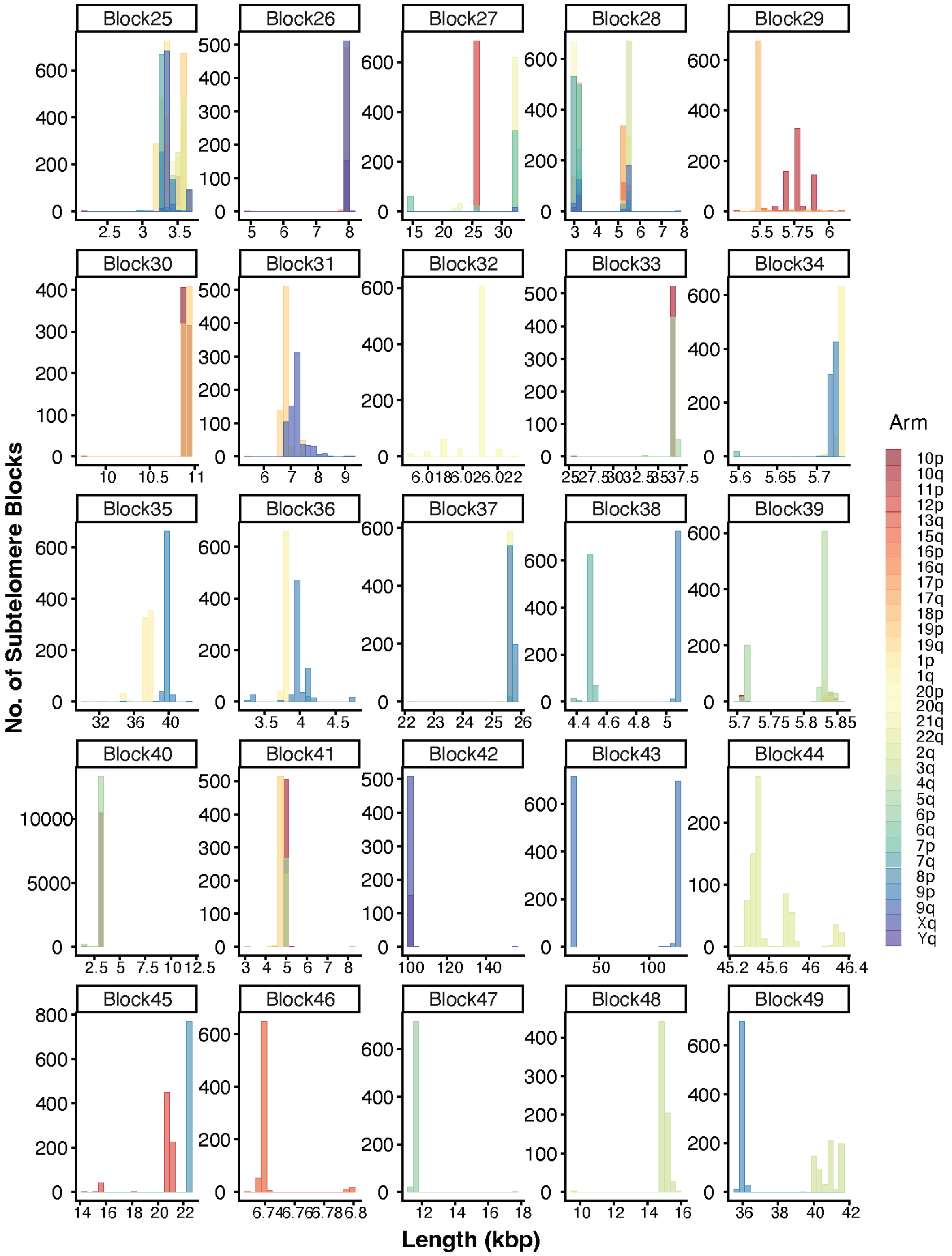


**Supplementary Fig. S57. Subtelomeric block length distribution (II)** Figure interpretation follows that of Supplementary Figure S56, but for Block 25 to Block 49.

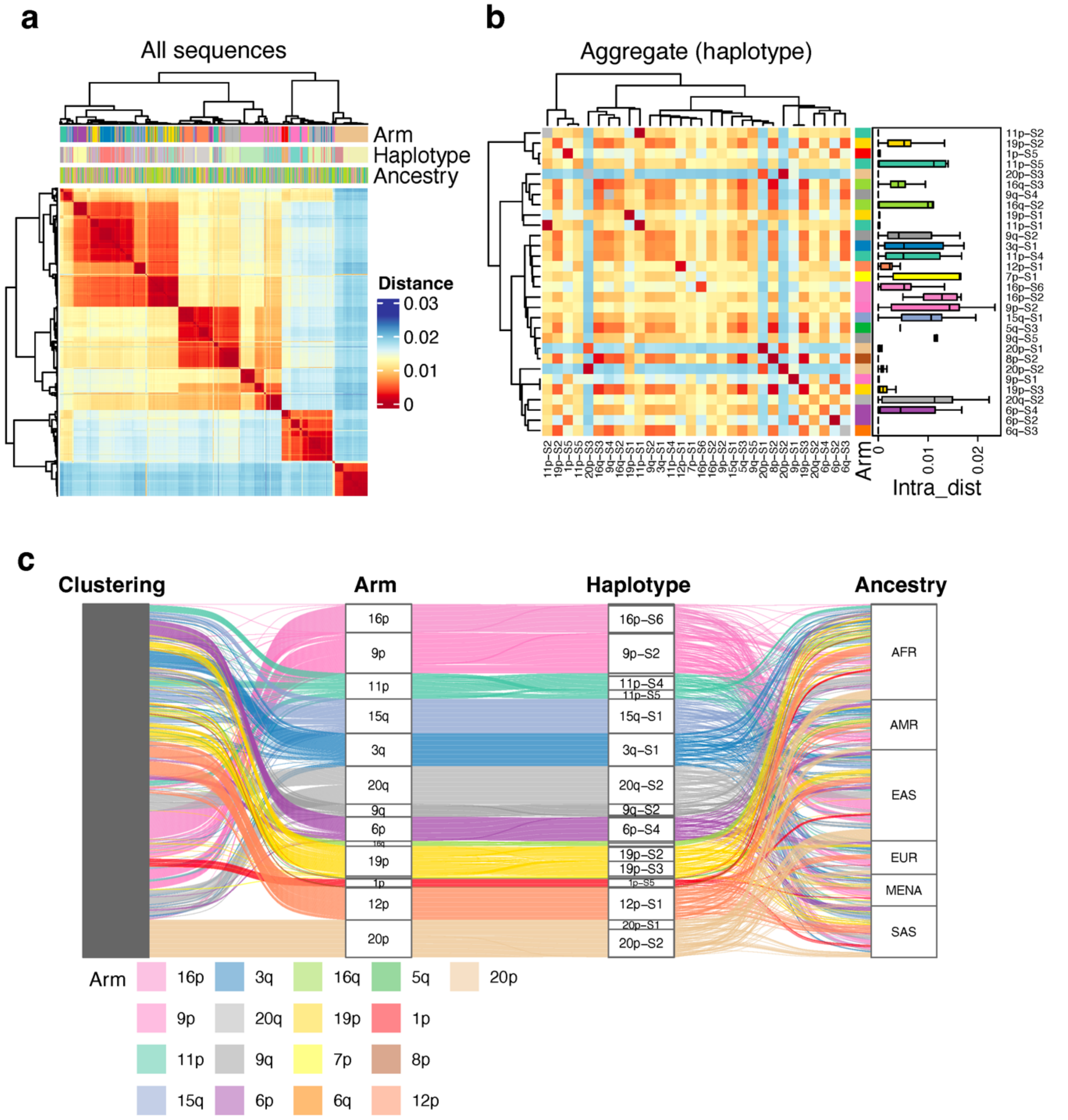


**Supplementary Fig. S58. Pairwise Mash distance analysis for 1000 randomly subsampled subtelomeric of Block 1.** (a) Pairwise Mash distance heatmap across all individual sequences, hierarchically clustered using Ward's method. Annotation bars indicate chromosomal arm of origin, haplotype, and population ancestry. (b) Aggregate heatmap of mean inter-haplotype Mash distances, collapsed to the haplotype level. The boxplot on the right shows the distribution of intra-haplotype pairwise distances per haplotype, coloured by chromosomal arm of origin. (c) Alluvial plot linking hierarchical clustering rank, chromosomal arm, haplotype identity, and population ancestry.

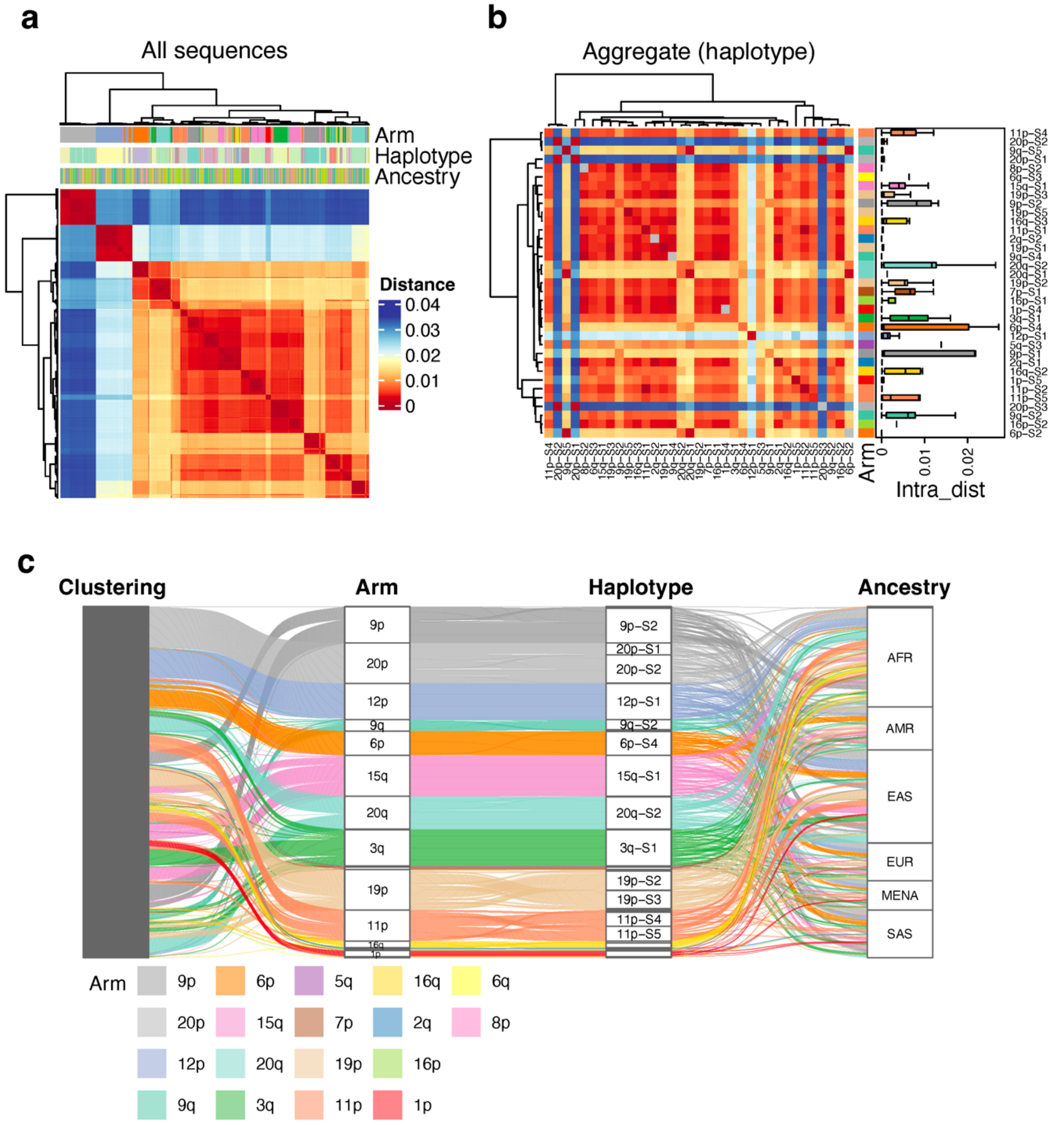


**Supplementary Fig. S59. Pairwise Mash distance analysis for 1000 randomly subsampled subtelomeric of Block 2.** Figure interpretation follows that of Supplementary Figure S58, but for Block 2.

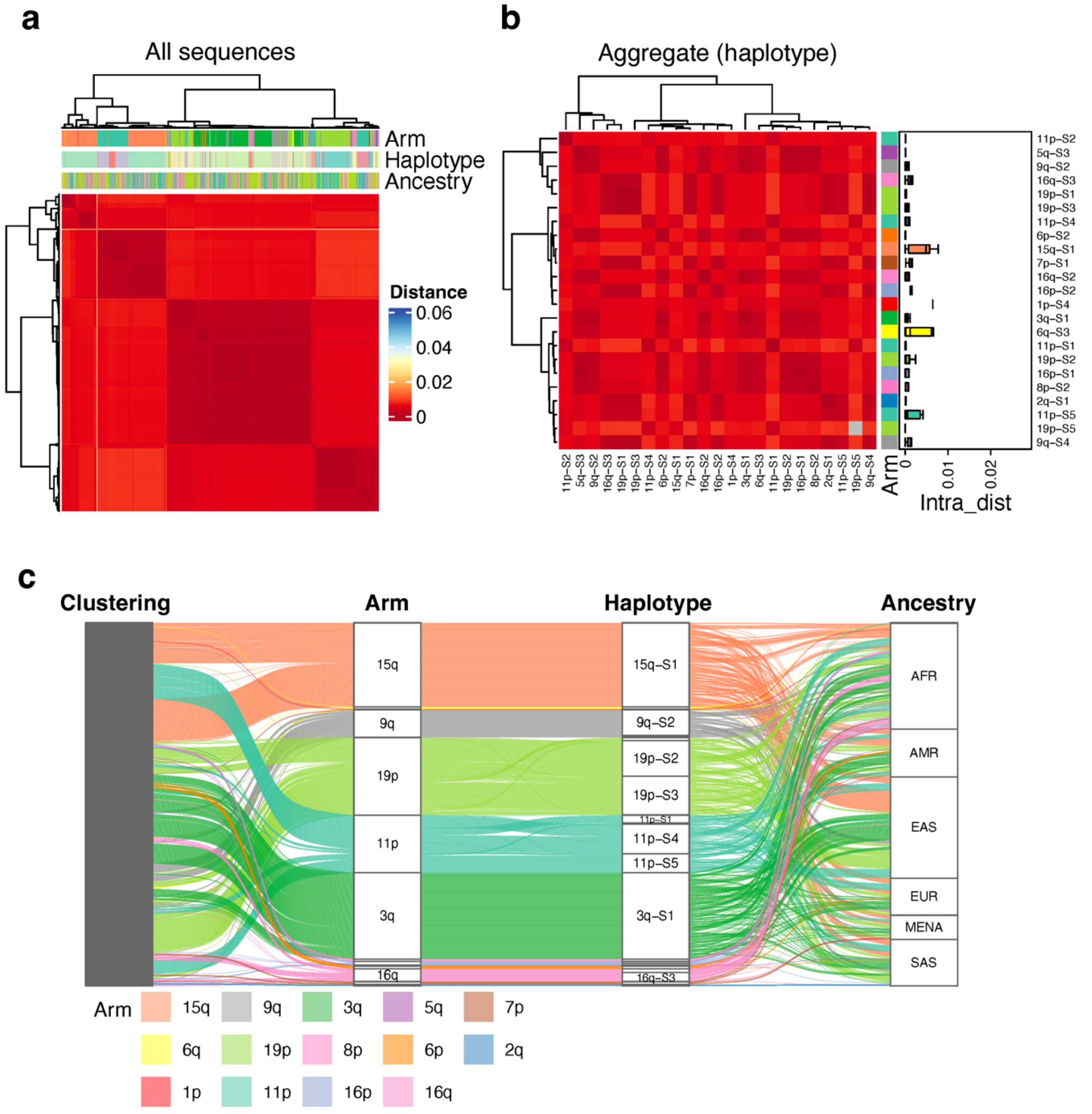


**Supplementary Fig. S60. Pairwise Mash distance analysis for 1000 randomly subsampled subtelomeric of Block 3.** Figure interpretation follows that of Supplementary Figure S58, but for Block 3.

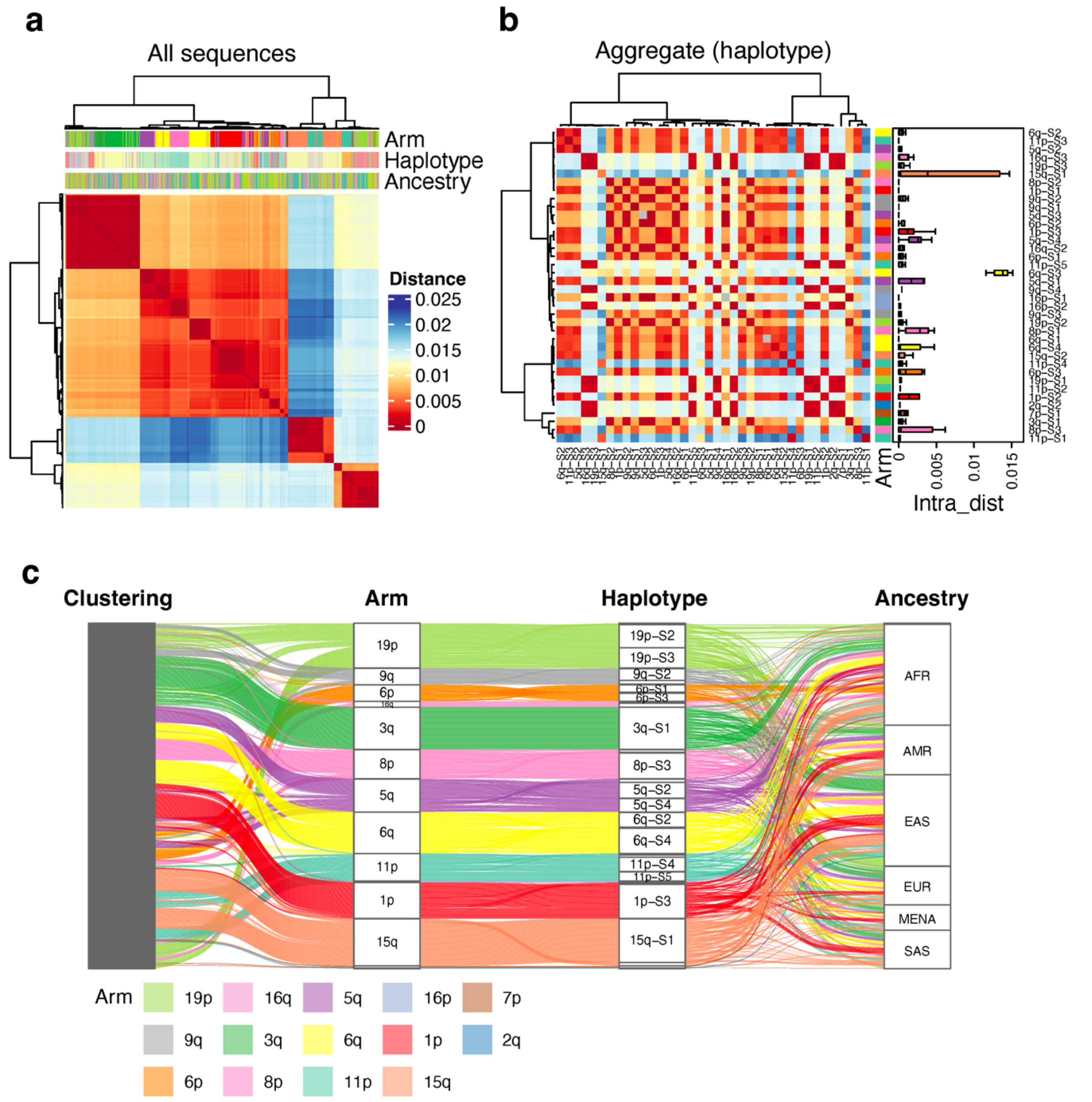


**Supplementary Fig. S61. Pairwise Mash distance analysis for 1000 randomly subsampled subtelomeric of Block 4.** Figure interpretation follows that of Supplementary Figure S58, but for Block 4.

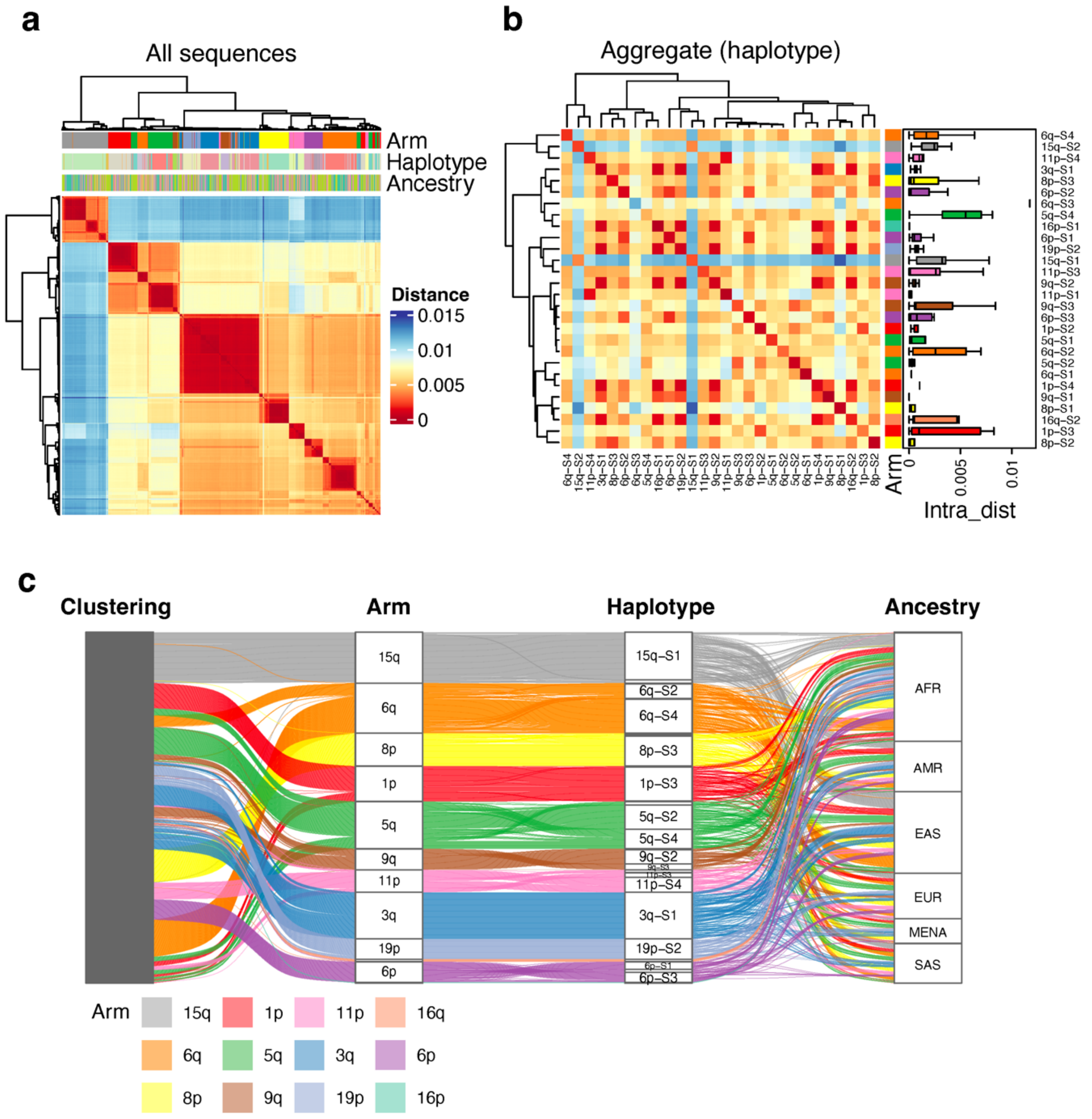


**Supplementary Fig. S62. Pairwise Mash distance analysis for 1000 randomly subsampled subtelomeric of Block 5.** Figure interpretation follows that of Supplementary Figure S58, but for Block 5.

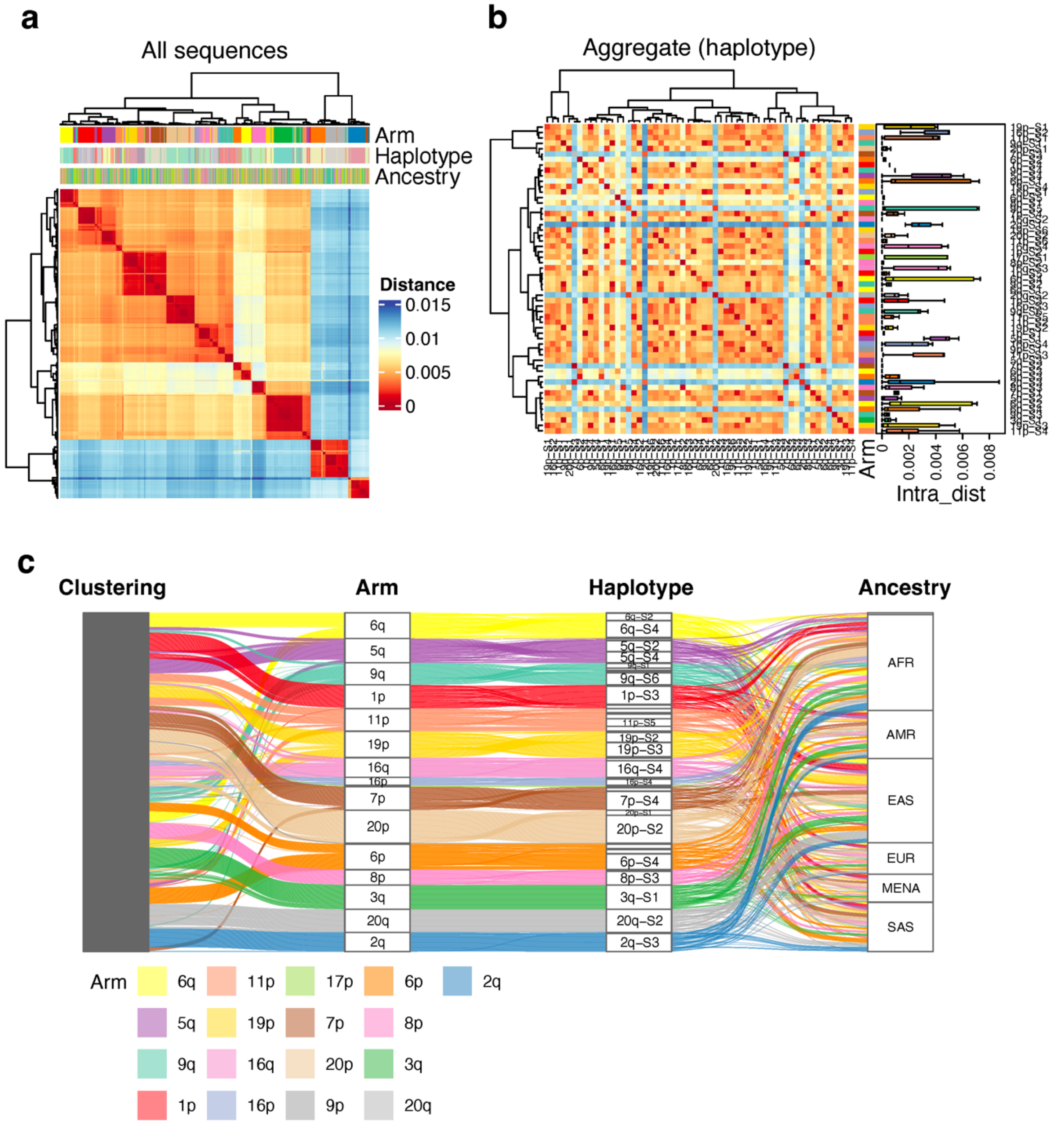


**Supplementary Fig. S63. Pairwise Mash distance analysis for 1000 randomly subsampled subtelomeric of Block 6.** Figure interpretation follows that of Supplementary Figure S58, but for Block 6.

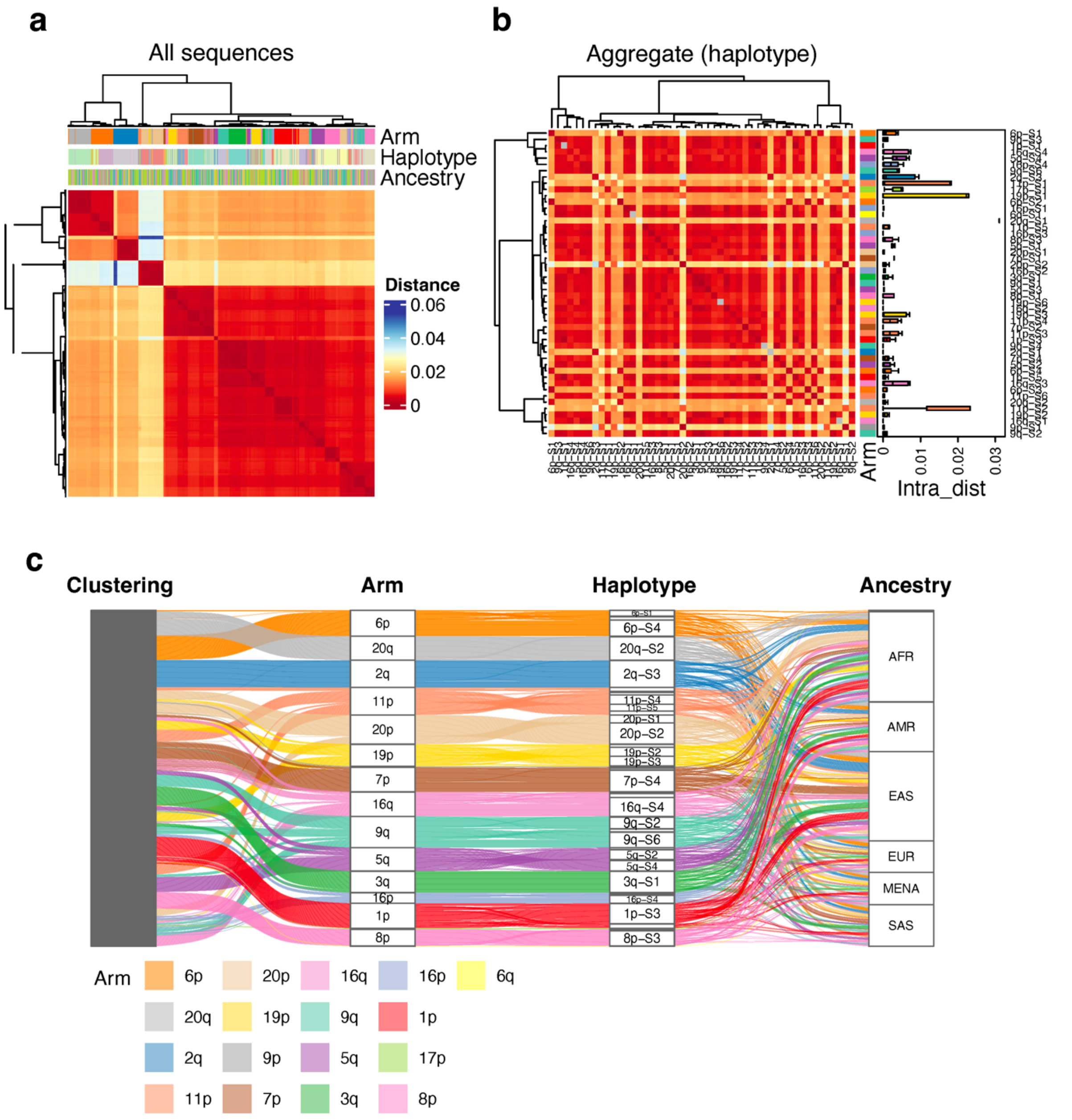


**Supplementary Fig. S64. Pairwise Mash distance analysis for 1000 randomly subsampled subtelomeric of Block 7.** Figure interpretation follows that of Supplementary Figure S58, but for Block 7.

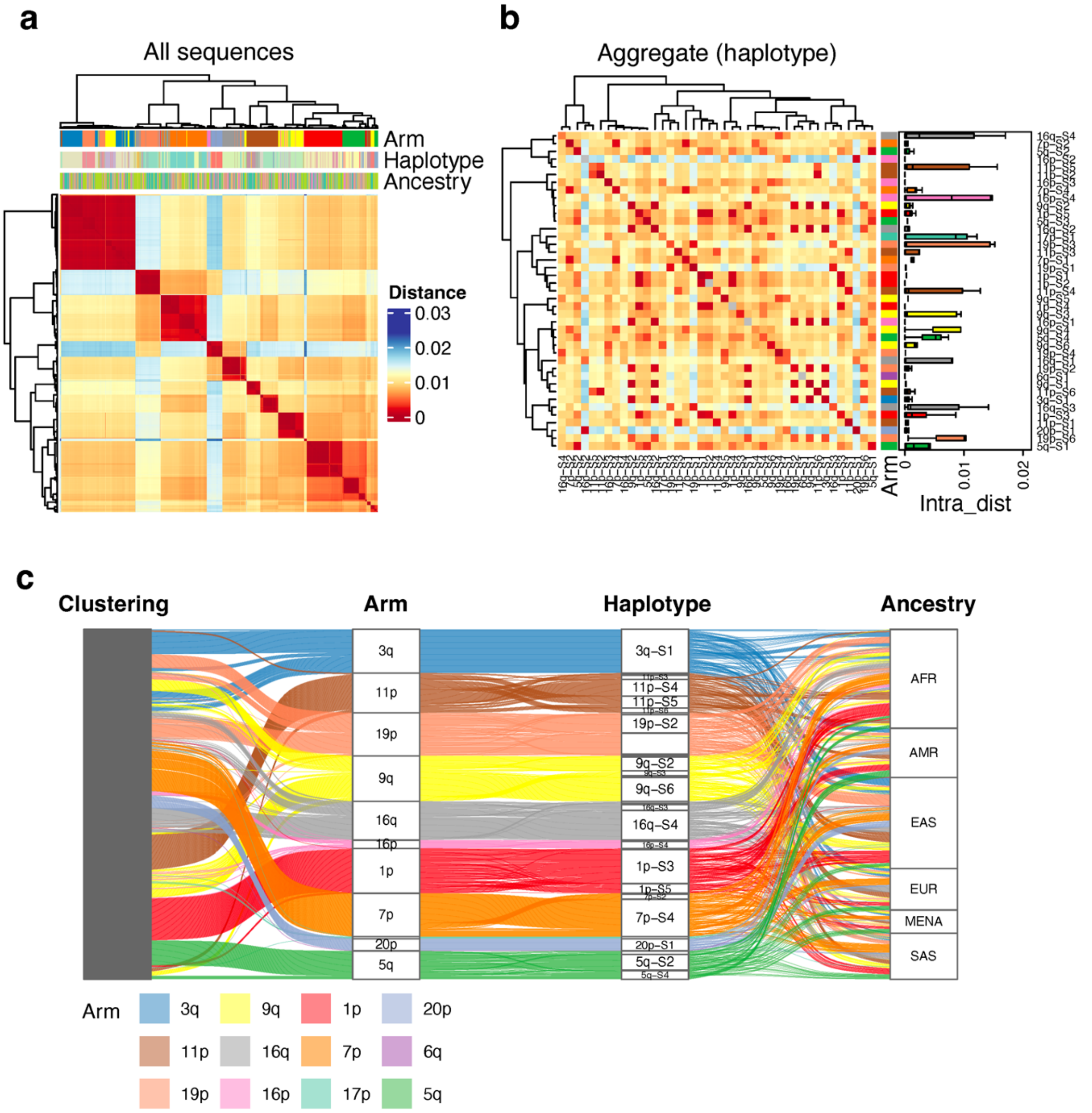


**Supplementary Fig. S65. Pairwise Mash distance analysis for 1000 randomly subsampled subtelomeric of Block 8.** Figure interpretation follows that of Supplementary Figure S58, but for Block 8.

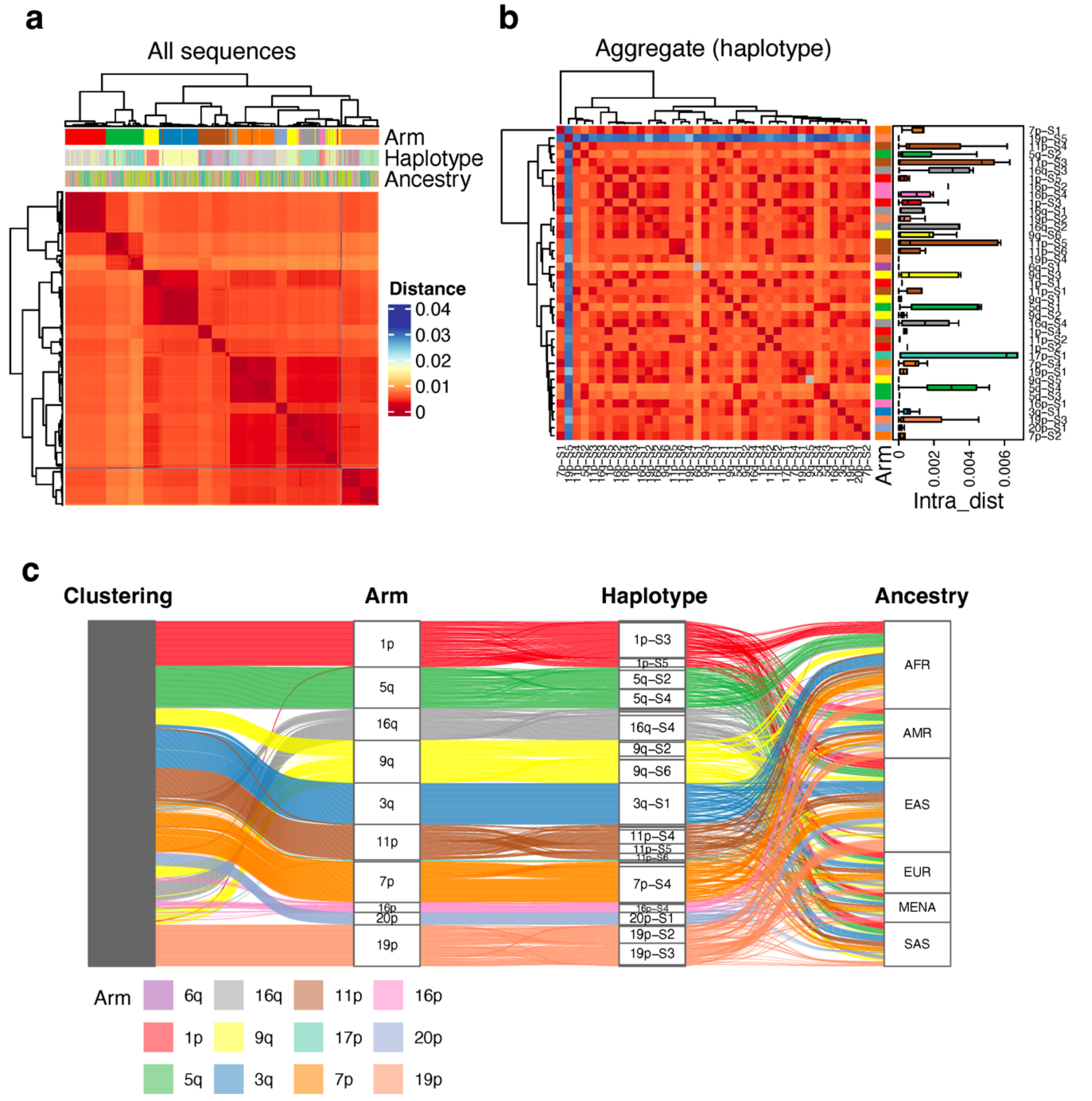


**Supplementary Fig. S66. Pairwise Mash distance analysis for 1000 randomly subsampled subtelomeric of Block 9.** Figure interpretation follows that of Supplementary Figure S58, but for Block 9.

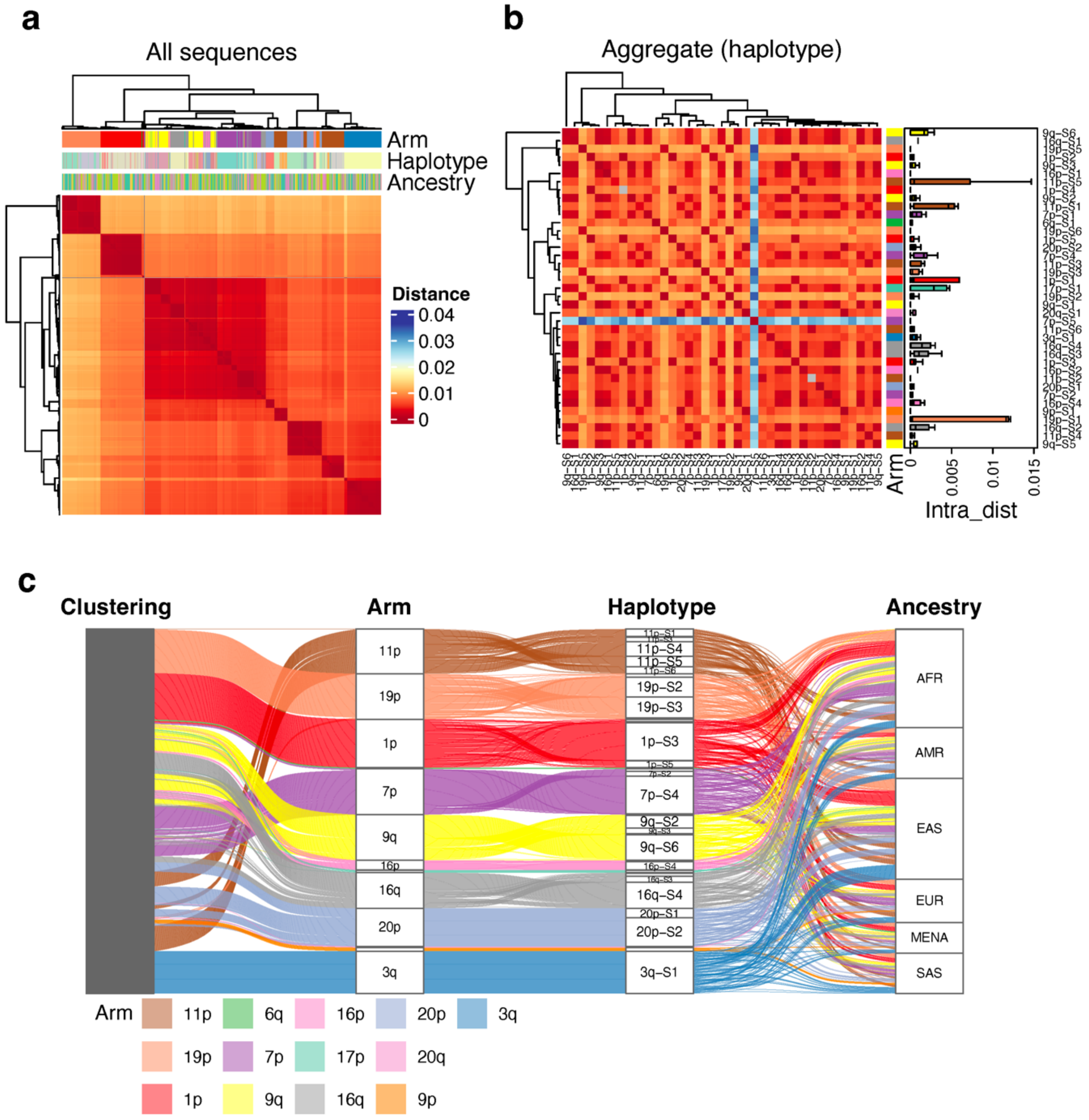


**Supplementary Fig. S67. Pairwise Mash distance analysis for 1000 randomly subsampled subtelomeric of Block 10.** Figure interpretation follows that of Supplementary Figure S58, but for Block 10.

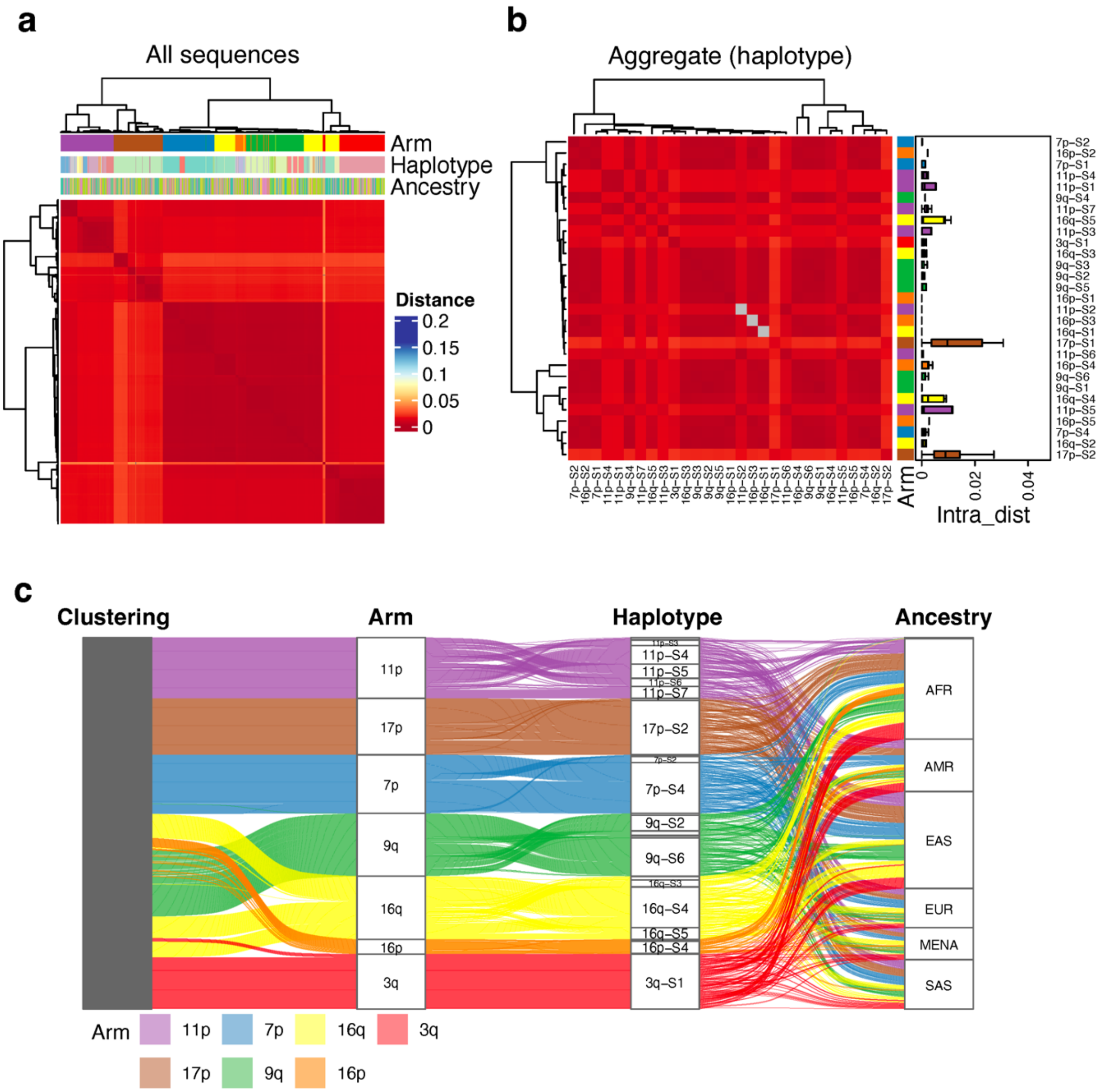


**Supplementary Fig. S68. Pairwise Mash distance analysis for 1000 randomly subsampled subtelomeric of Block 11.** Figure interpretation follows that of Supplementary Figure S58, but for Block 11.

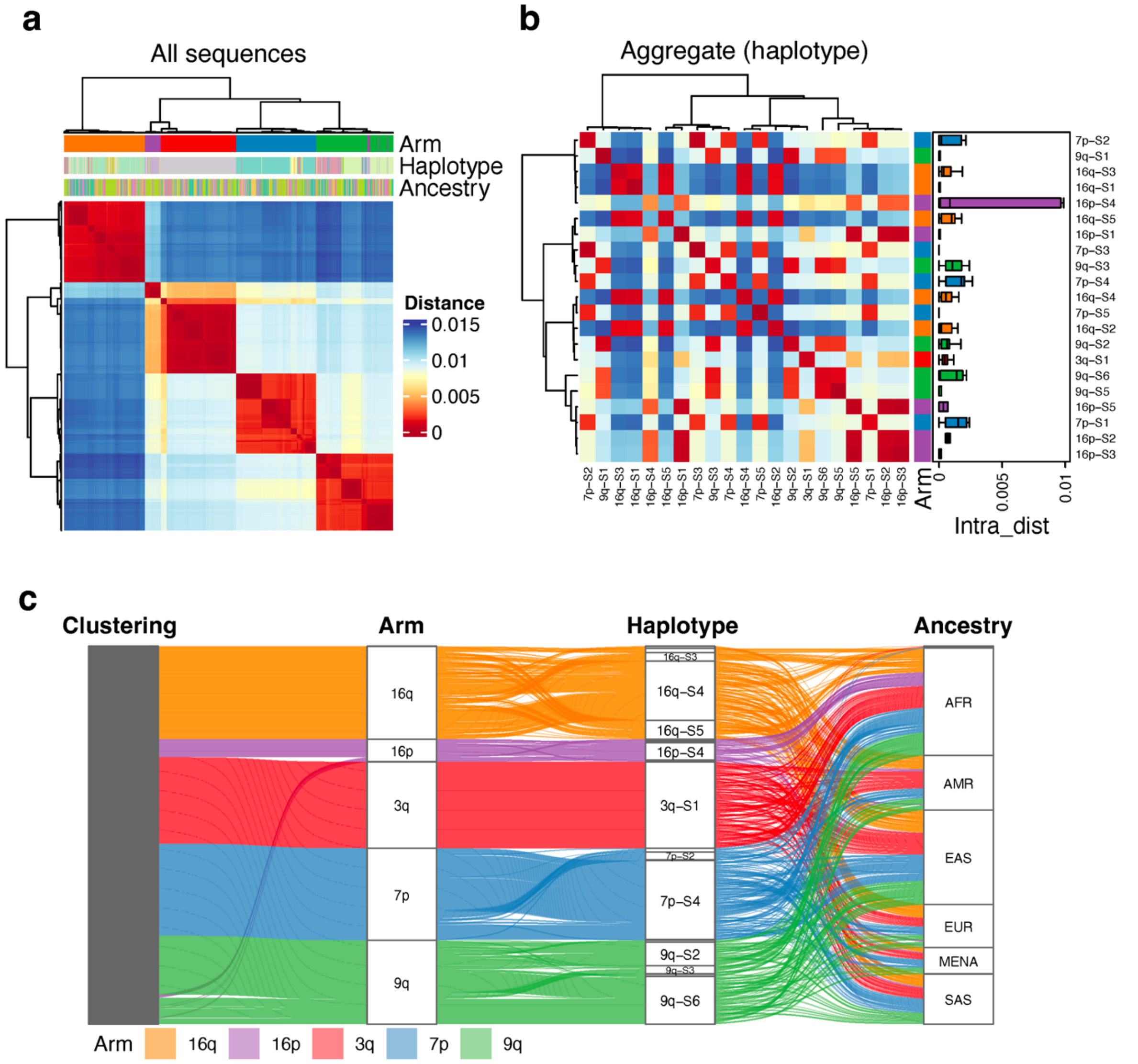


**Supplementary Fig. S69. Pairwise Mash distance analysis for 1000 randomly subsampled subtelomeric of Block 12.** Figure interpretation follows that of Supplementary Figure S58, but for Block 12.

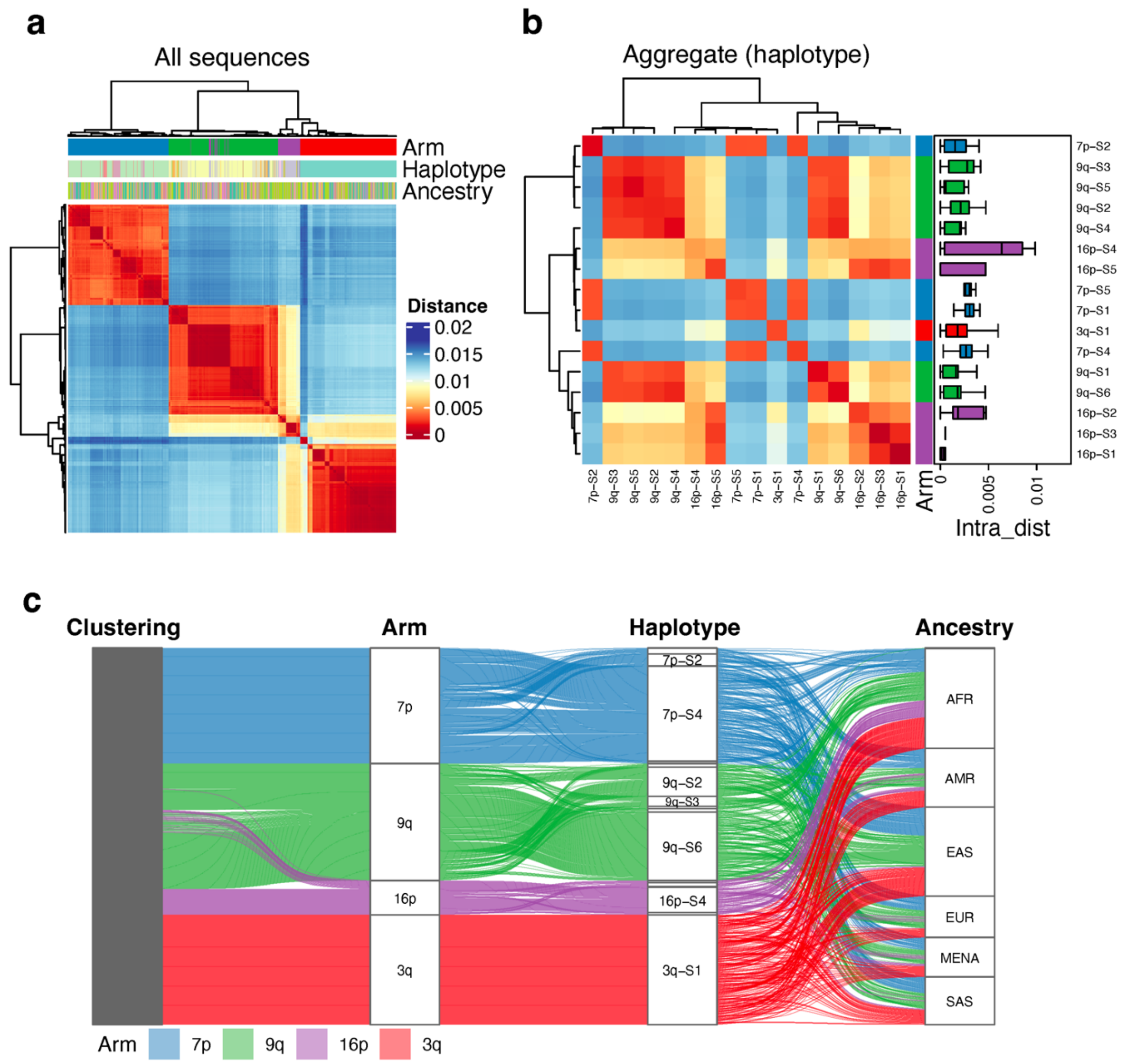


**Supplementary Fig. S70. Pairwise Mash distance analysis for 1000 randomly subsampled subtelomeric of Block 13.** Figure interpretation follows that of Supplementary Figure S58, but for Block 13.

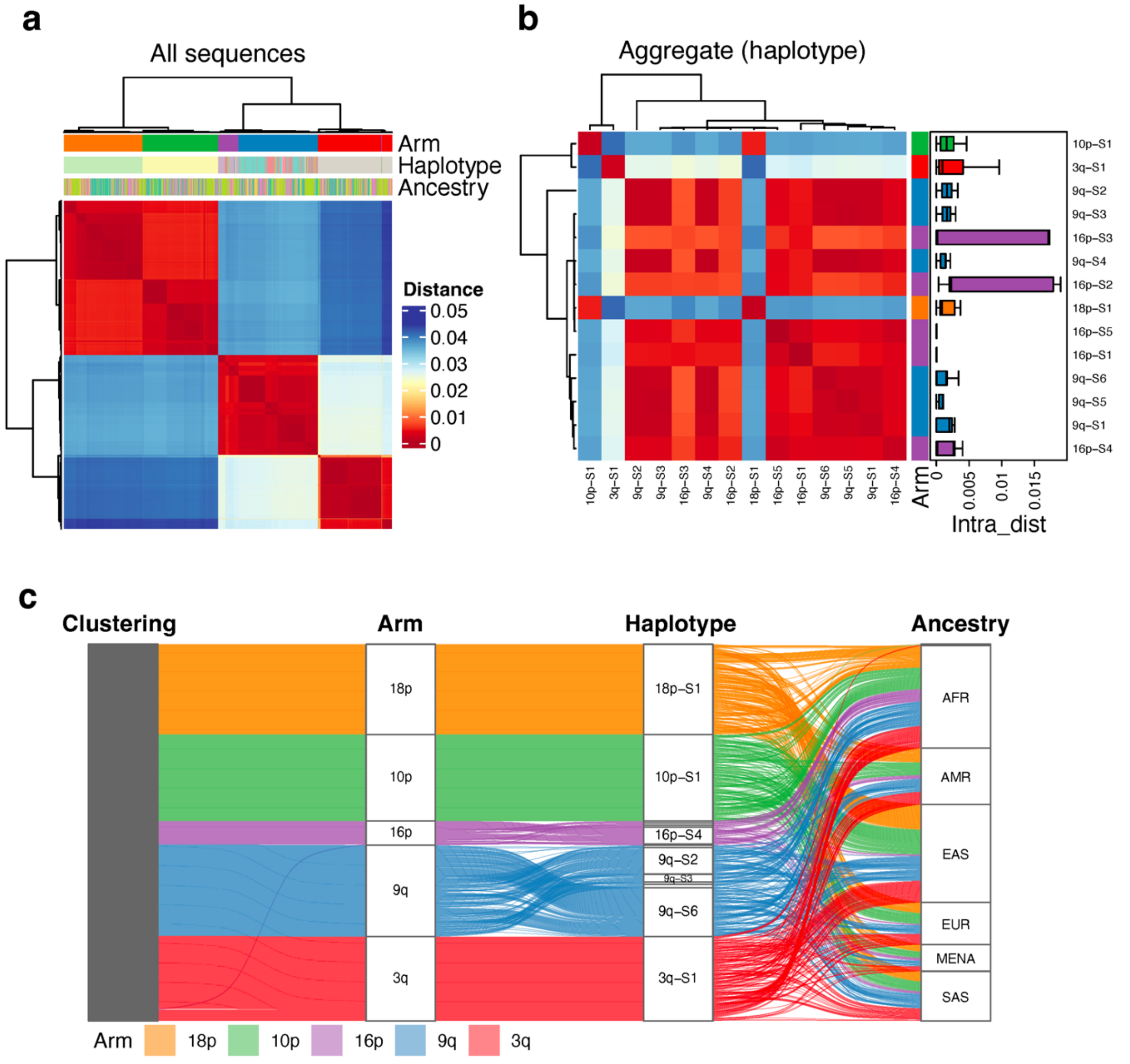


**Supplementary Fig. S71. Pairwise Mash distance analysis for 1000 randomly subsampled subtelomeric of Block 14.** Figure interpretation follows that of Supplementary Figure S58, but for Block 14.

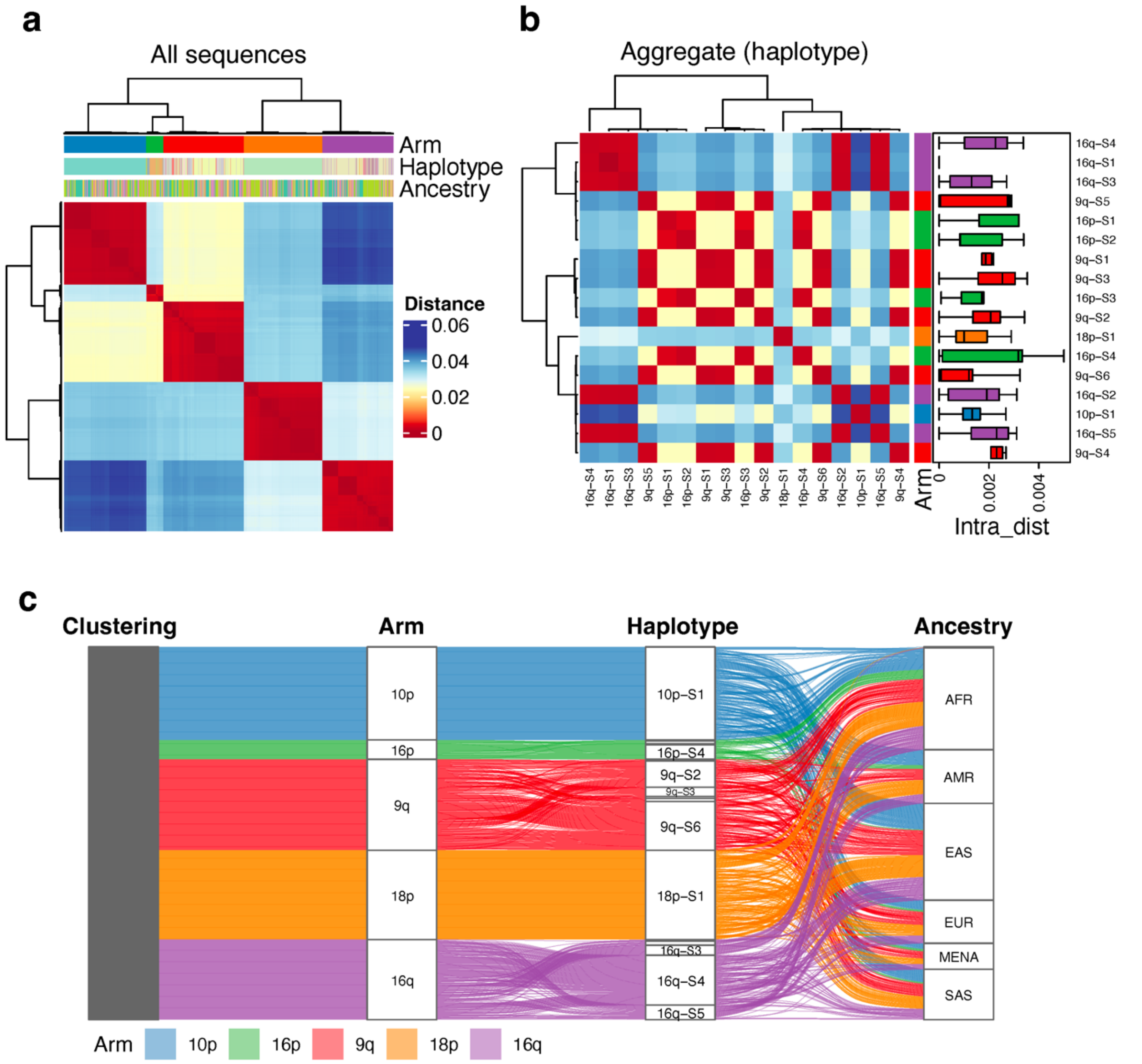


**Supplementary Fig. S72. Pairwise Mash distance analysis for 1000 randomly subsampled subtelomeric of Block 15.** Figure interpretation follows that of Supplementary Figure S58, but for Block 15.

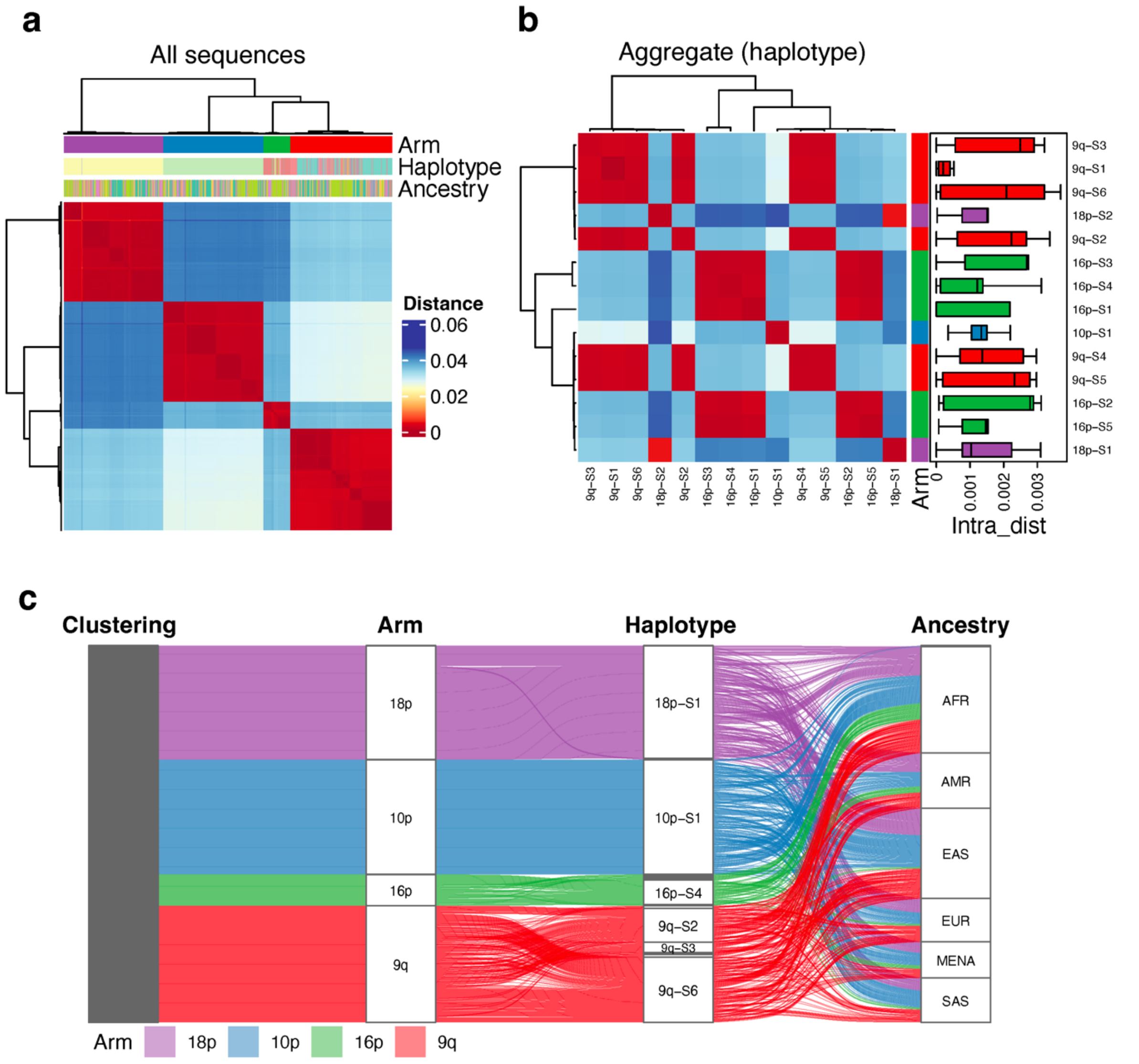


**Supplementary Fig. S73. Pairwise Mash distance analysis for 1000 randomly subsampled subtelomeric of Block 16.** Figure interpretation follows that of Supplementary Figure S58, but for Block 16.

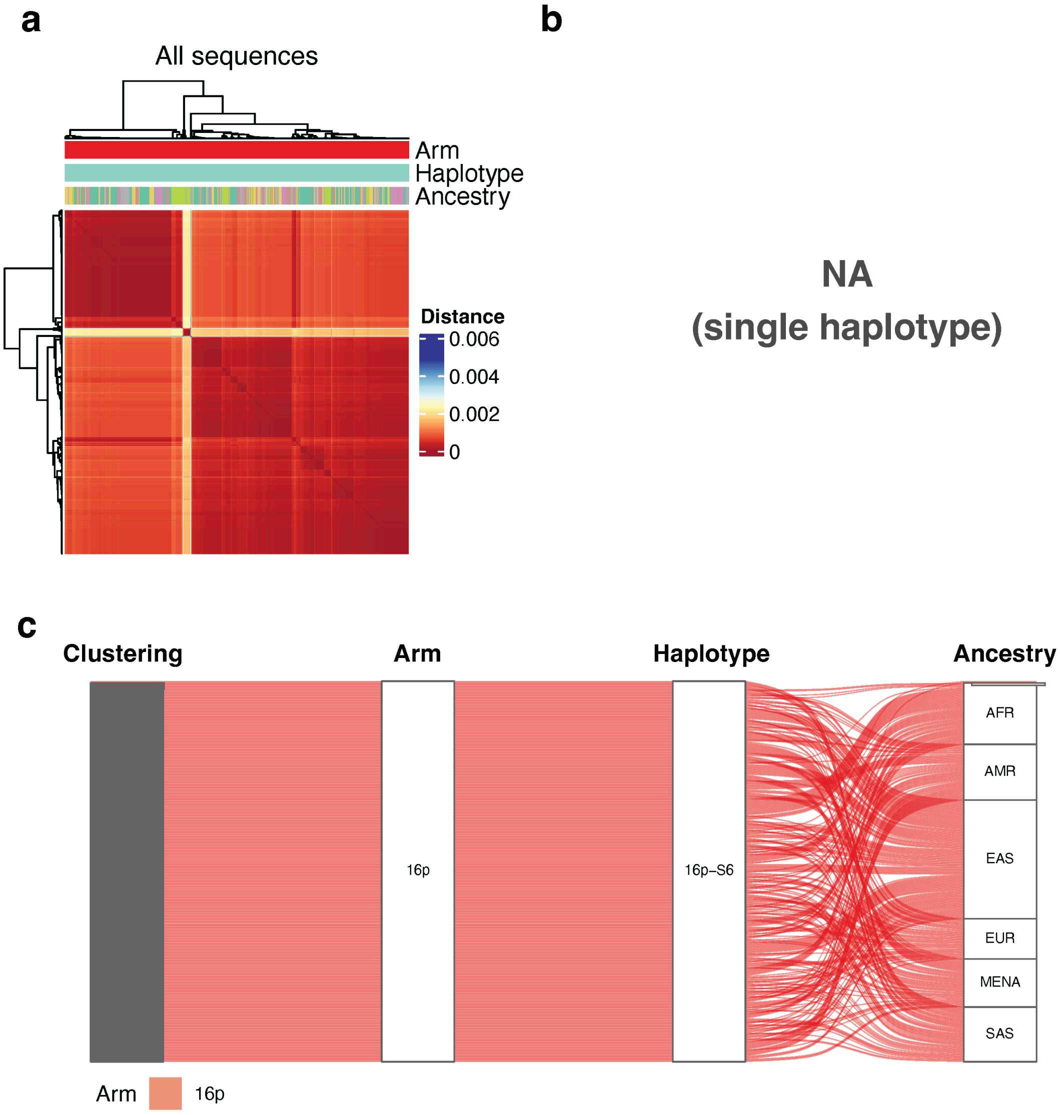


**Supplementary Fig. S74. Pairwise Mash distance analysis for 1000 randomly subsampled subtelomeric of Block 17.** Figure interpretation follows that of Supplementary Figure S58, but for Block 17.

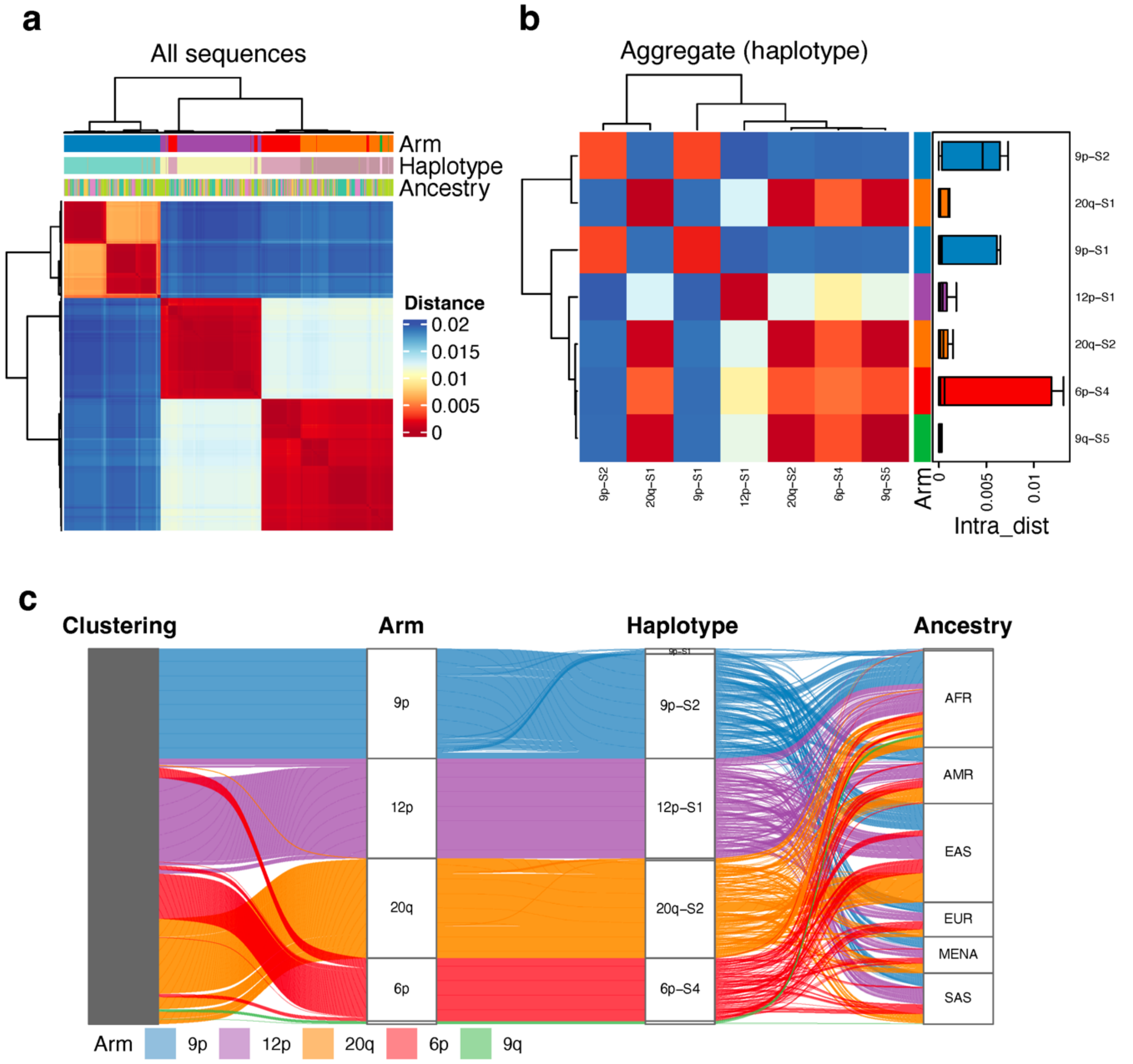


**Supplementary Fig. S75. Pairwise Mash distance analysis for 1000 randomly subsampled subtelomeric of Block 18.** Figure interpretation follows that of Supplementary Figure S58, but for Block 18.

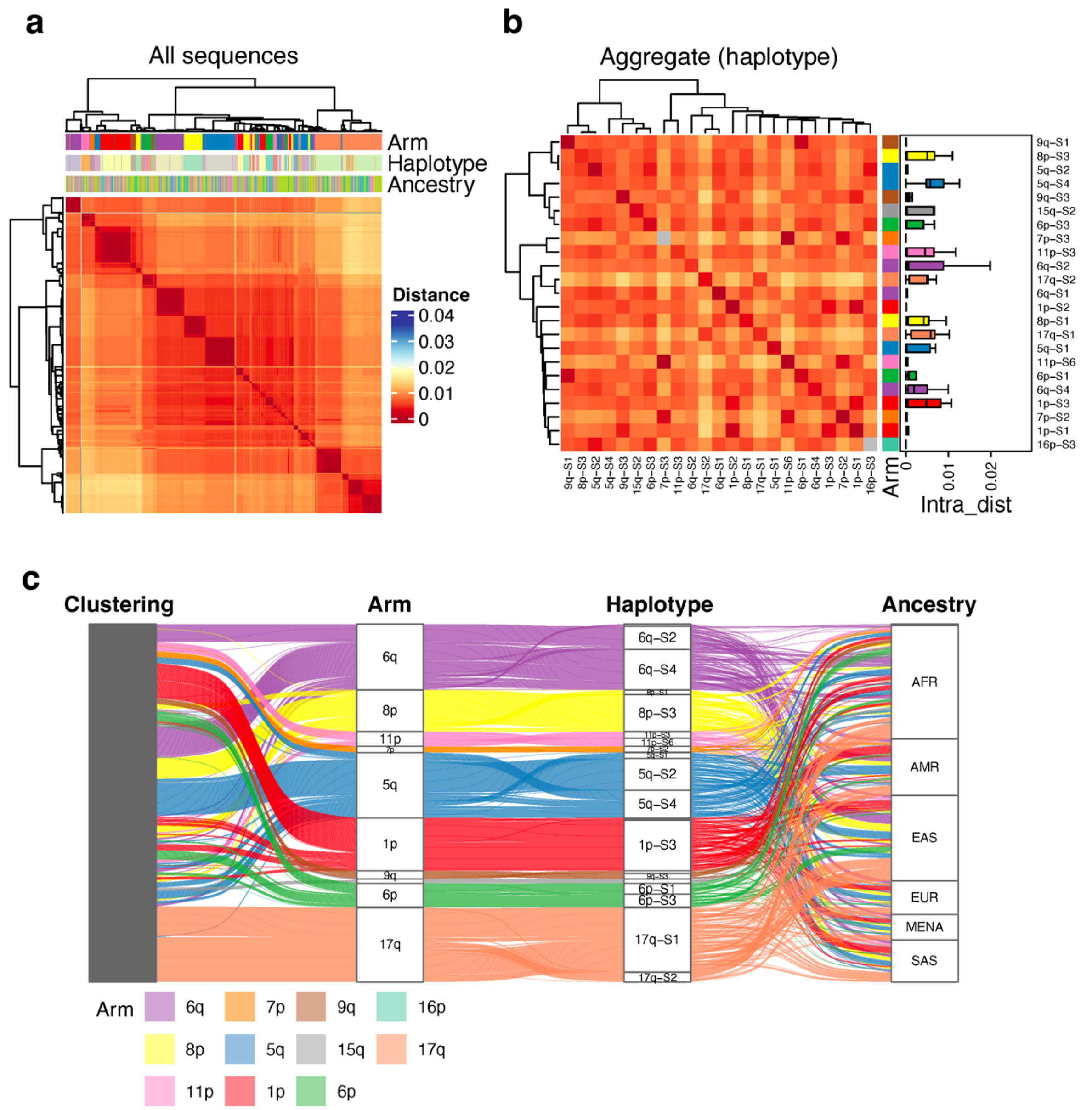


**Supplementary Fig. S76. Pairwise Mash distance analysis for 1000 randomly subsampled subtelomeric of Block 19a.** Figure interpretation follows that of Supplementary Figure S58, but for Block 19a.

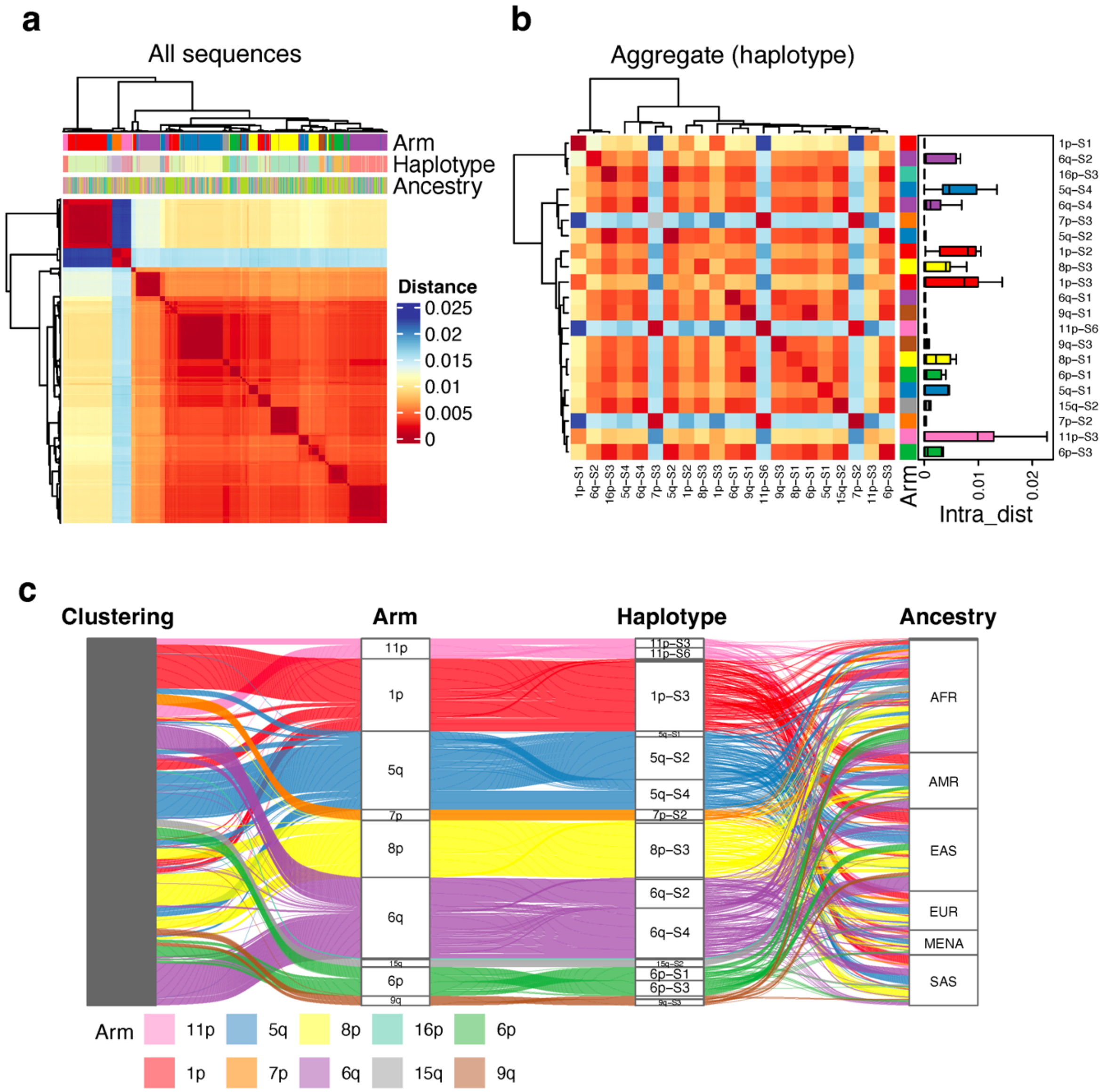


**Supplementary Fig. S77. Pairwise Mash distance analysis for 1000 randomly subsampled subtelomeric of Block 19b.** Figure interpretation follows that of Supplementary Figure S58, but for Block 19b.

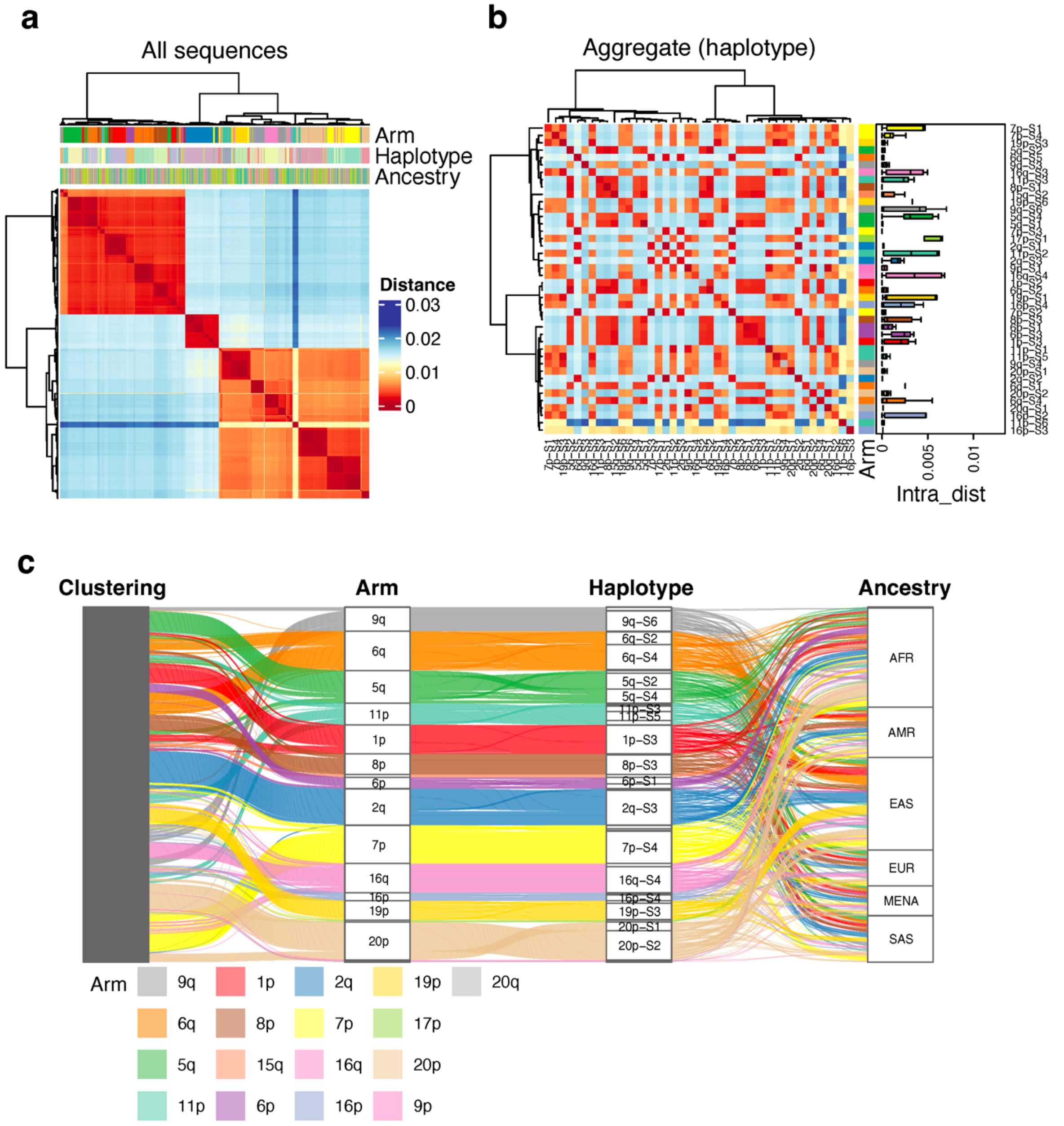


**Supplementary Fig. S78. Pairwise Mash distance analysis for 1000 randomly subsampled subtelomeric of Block 20.** Figure interpretation follows that of Supplementary Figure S58, but for Block 20.

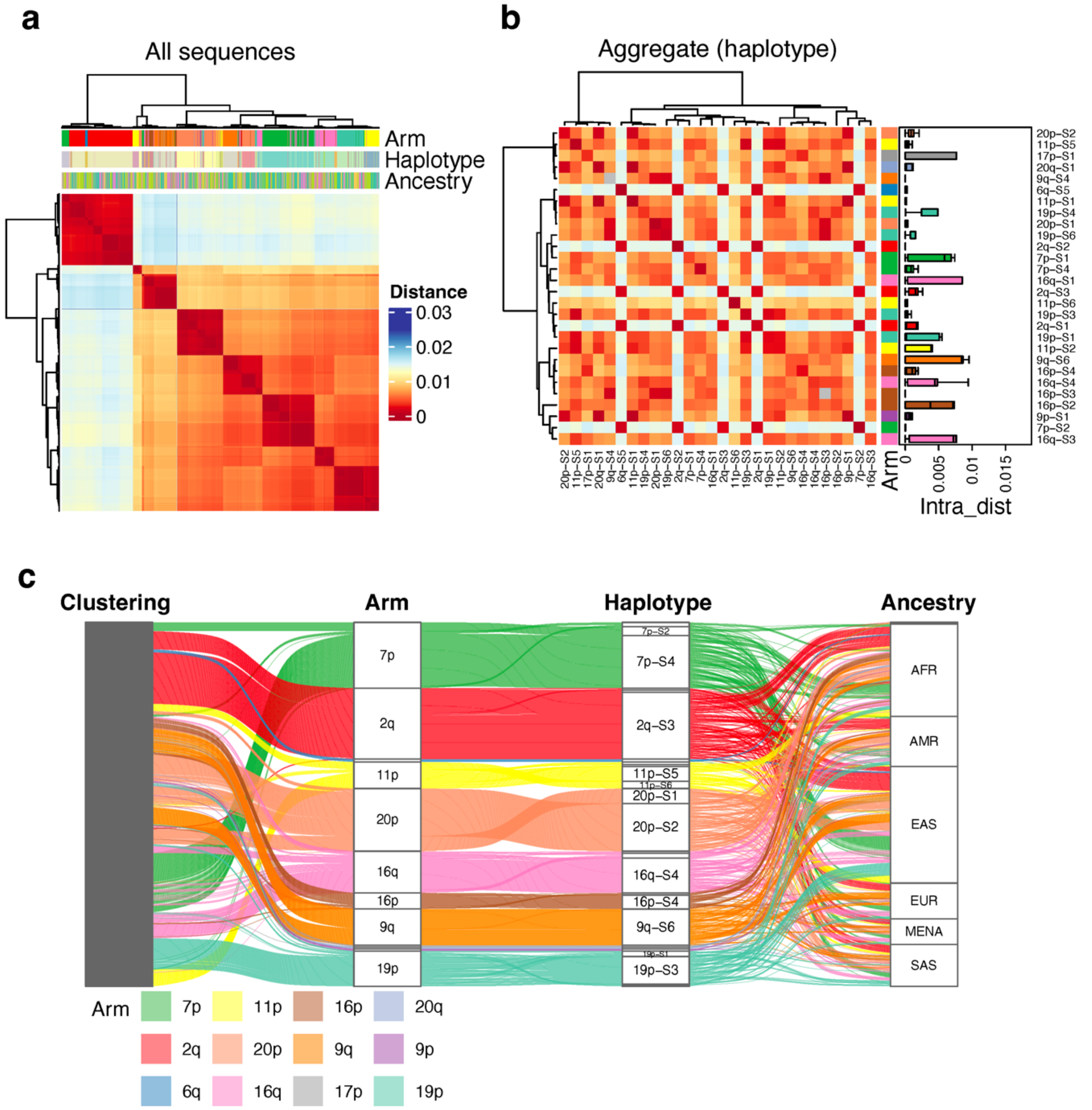


**Supplementary Fig. S79. Pairwise Mash distance analysis for 1000 randomly subsampled subtelomeric of Block 21.** Figure interpretation follows that of Supplementary Figure S58, but for Block 21.

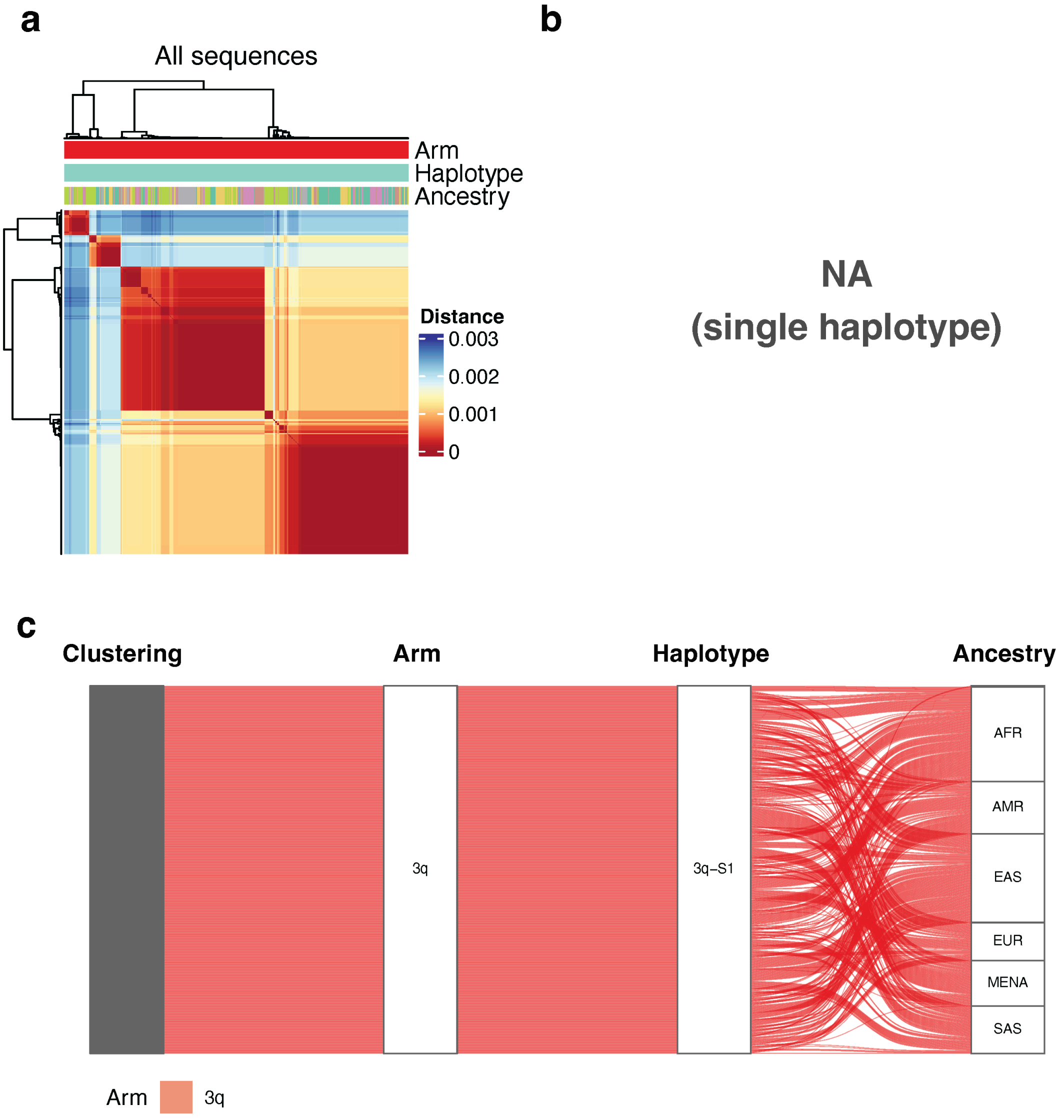


**Supplementary Fig. S80. Pairwise Mash distance analysis for 1000 randomly subsampled subtelomeric of Block 22.** Figure interpretation follows that of Supplementary Figure S58, but for Block 22.

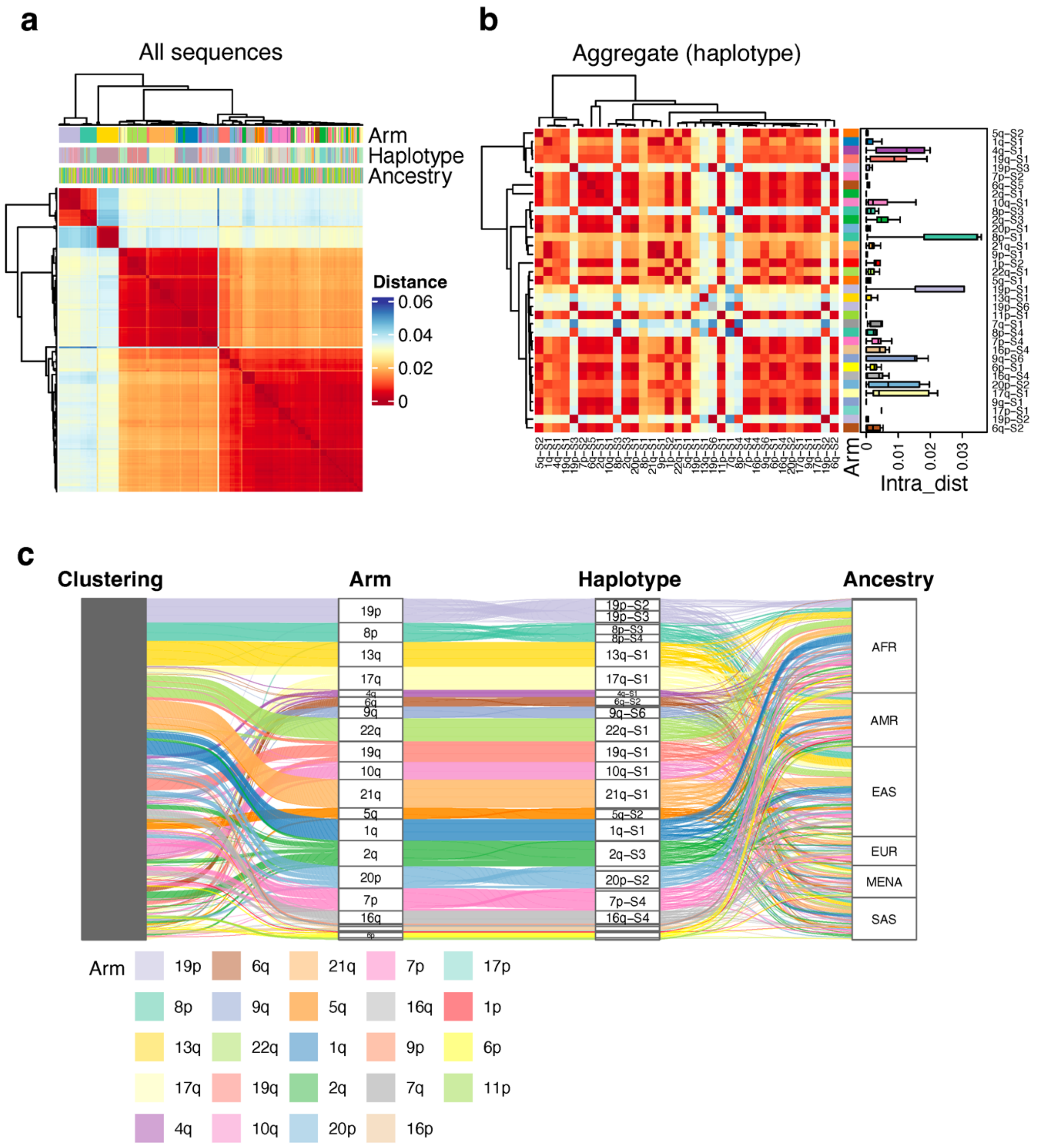


**Supplementary Fig. S81. Pairwise Mash distance analysis for 1000 randomly subsampled subtelomeric of Block 23.** Figure interpretation follows that of Supplementary Figure S58, but for Block 23.

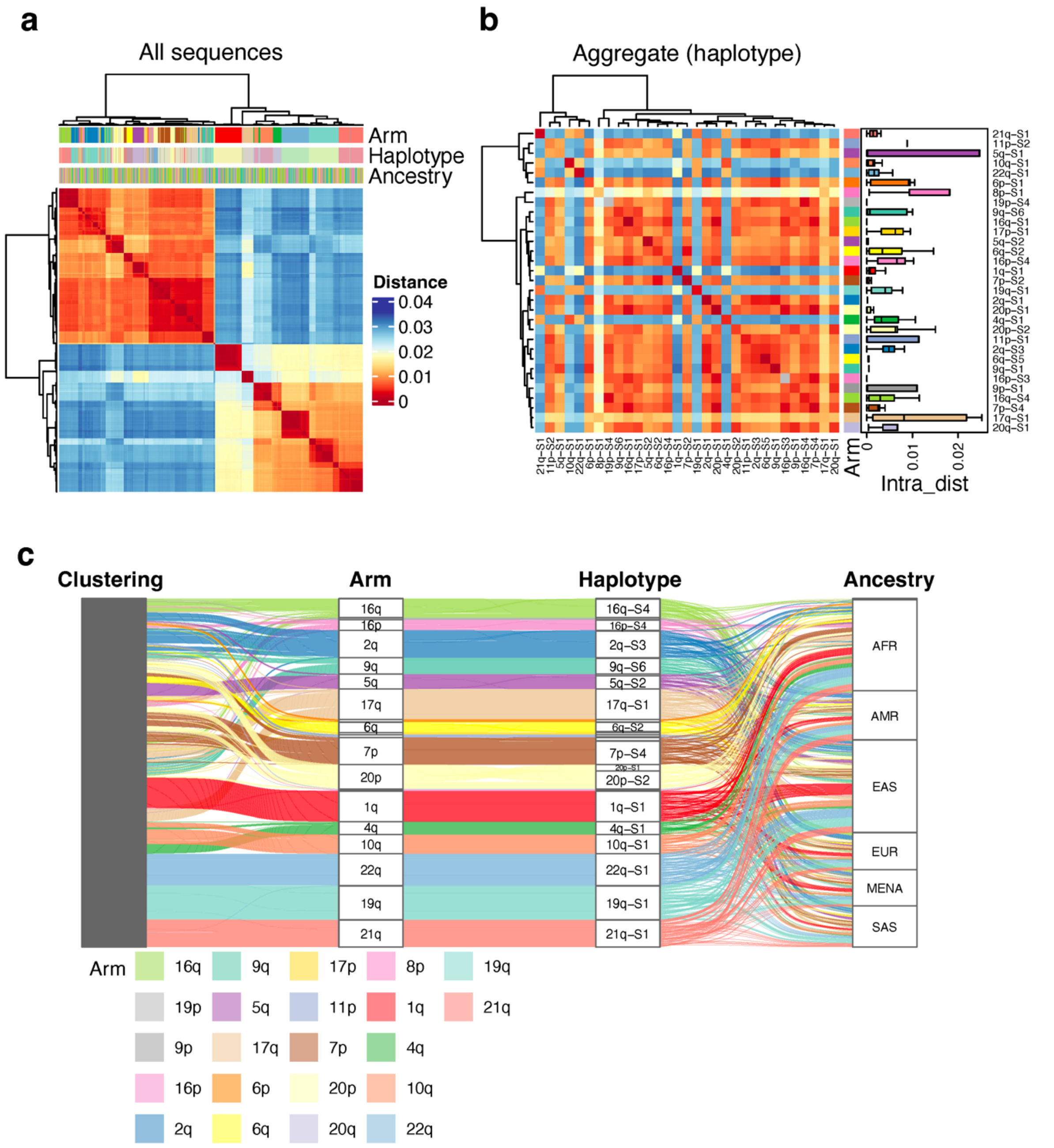


**Supplementary Fig. S82. Pairwise Mash distance analysis for 1000 randomly subsampled subtelomeric of Block 24.** Figure interpretation follows that of Supplementary Figure S58, but for Block 24.

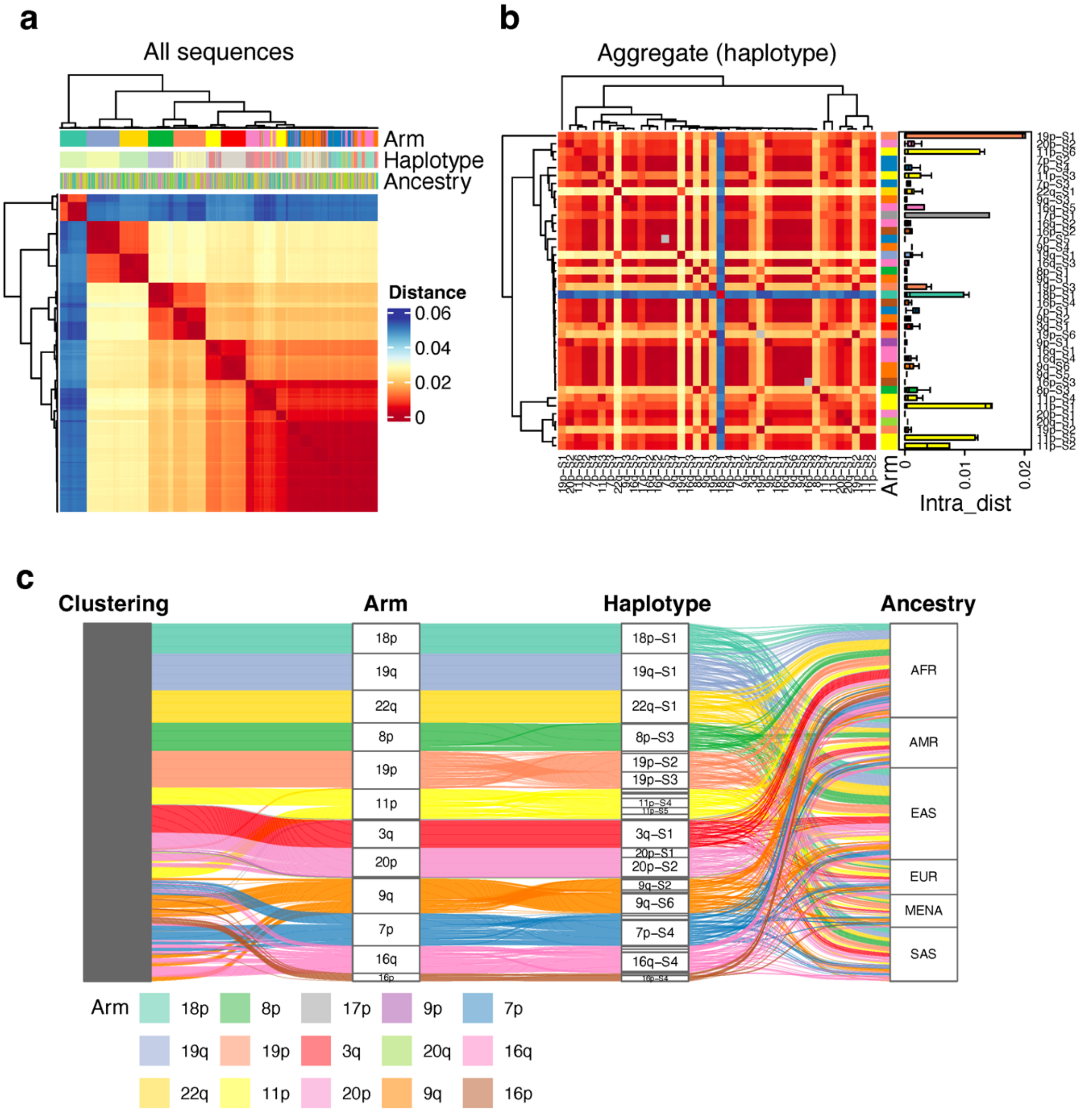


**Supplementary Fig. S83. Pairwise Mash distance analysis for 1000 randomly subsampled subtelomeric of Block 25.** Figure interpretation follows that of Supplementary Figure S58, but for Block 25.

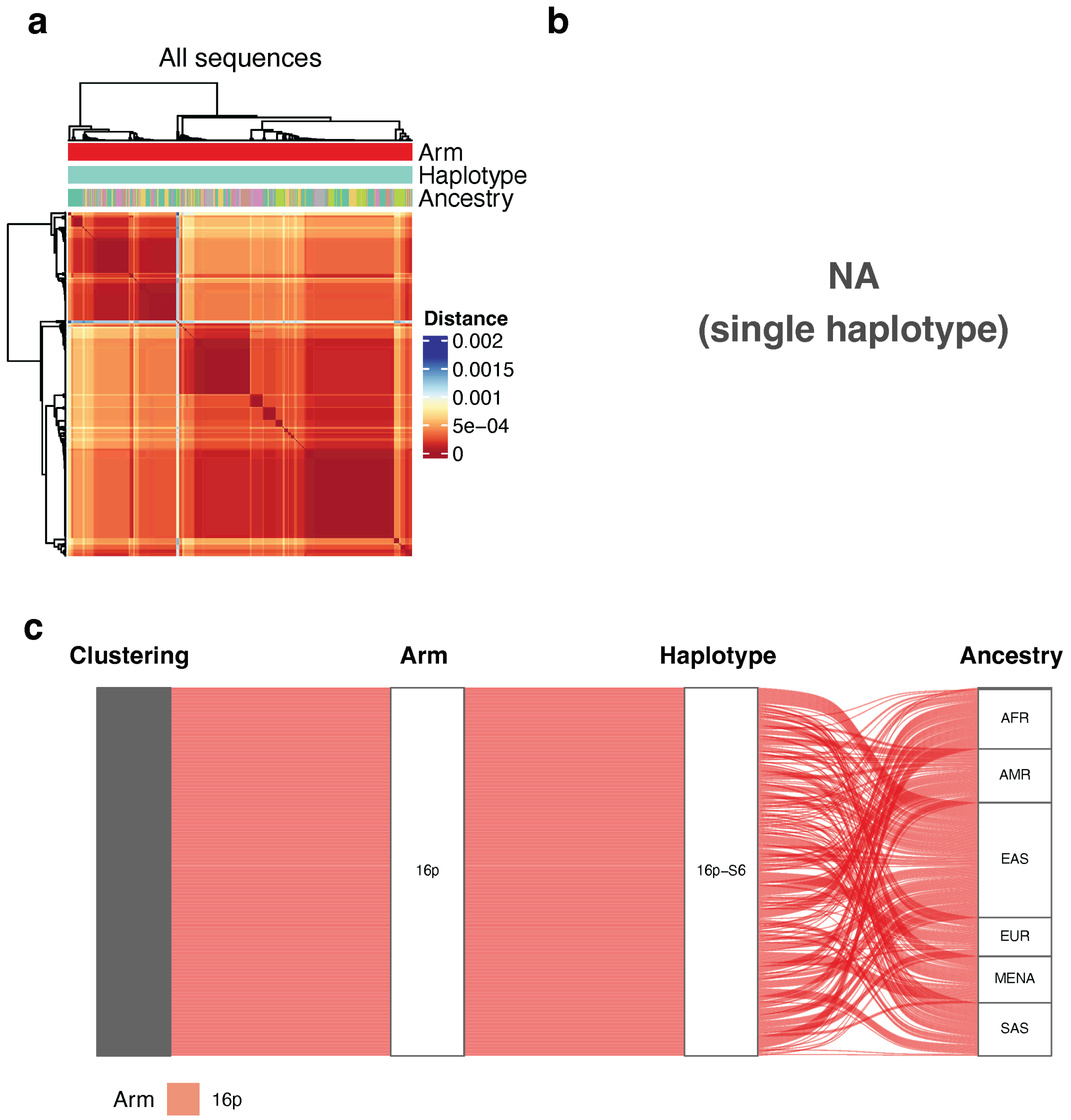


**Supplementary Fig. S84. Pairwise Mash distance analysis for 1000 randomly subsampled subtelomeric of Block 26.** Figure interpretation follows that of Supplementary Figure S58, but for Block 26.

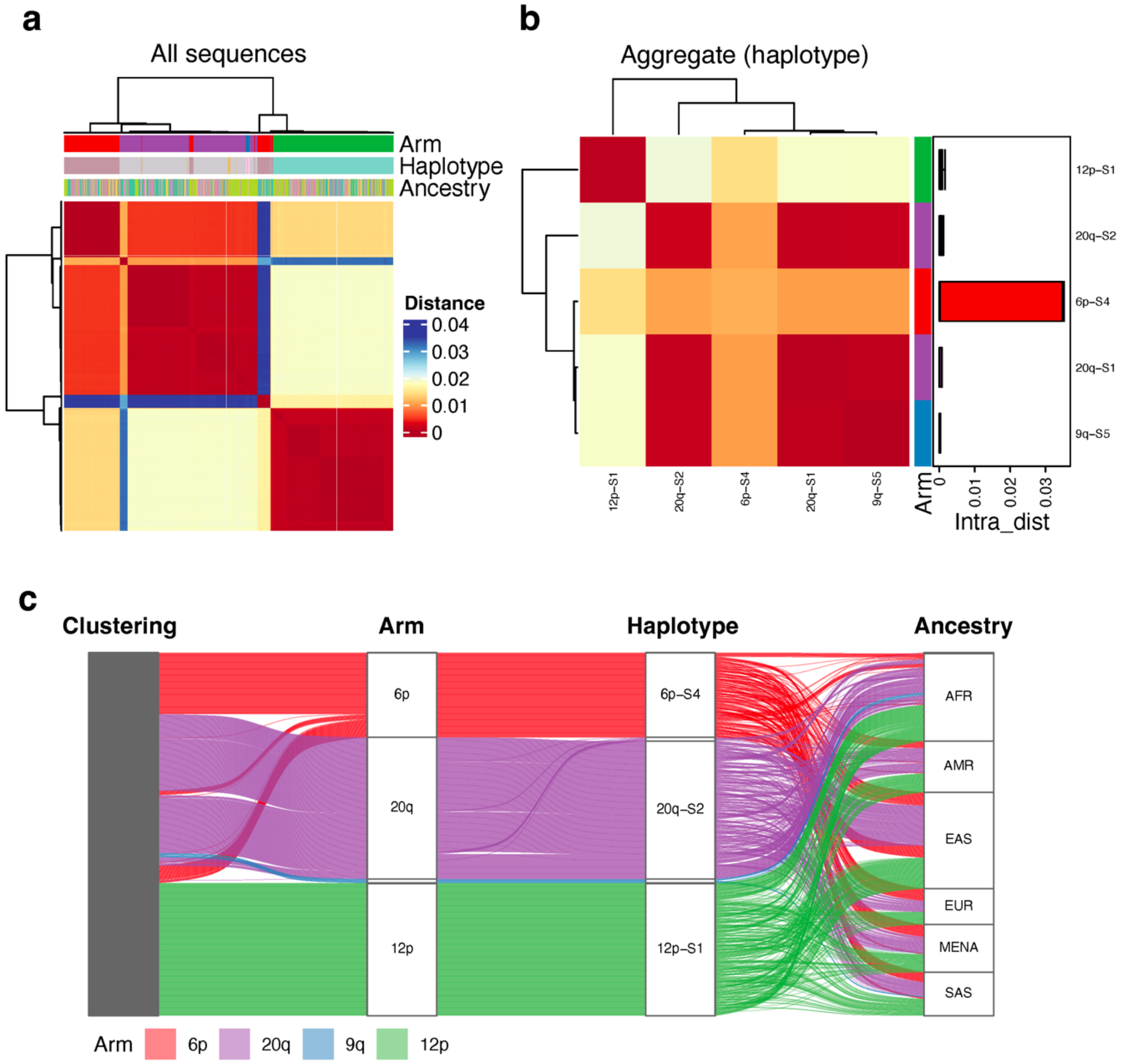


**Supplementary Fig. S85. Pairwise Mash distance analysis for 1000 randomly subsampled subtelomeric of Block 27.** Figure interpretation follows that of Supplementary Figure S58, but for Block 27.

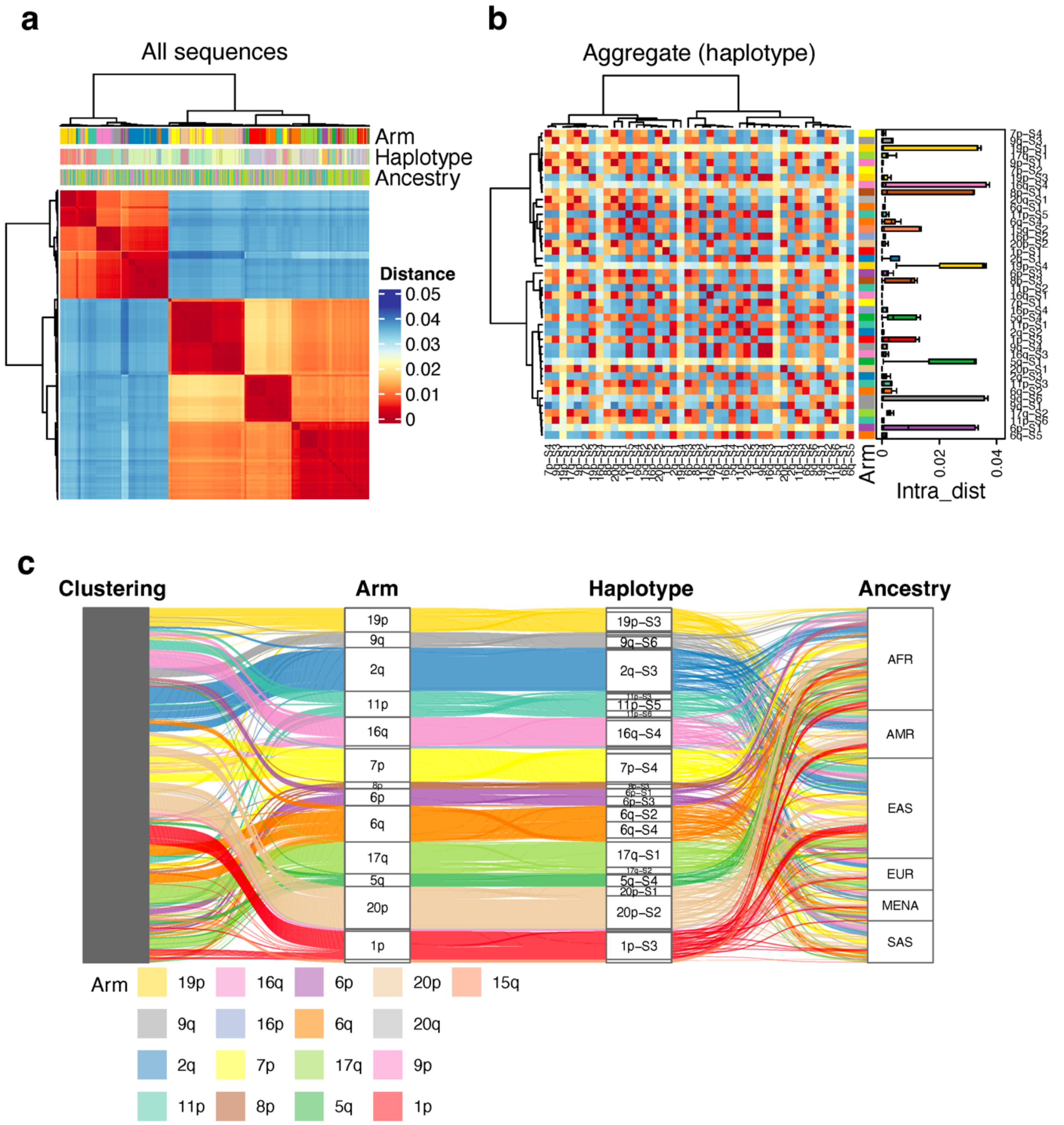


**Supplementary Fig. S86. Pairwise Mash distance analysis for 1000 randomly subsampled subtelomeric of Block 28.** Figure interpretation follows that of Supplementary Figure S58, but for Block 28.

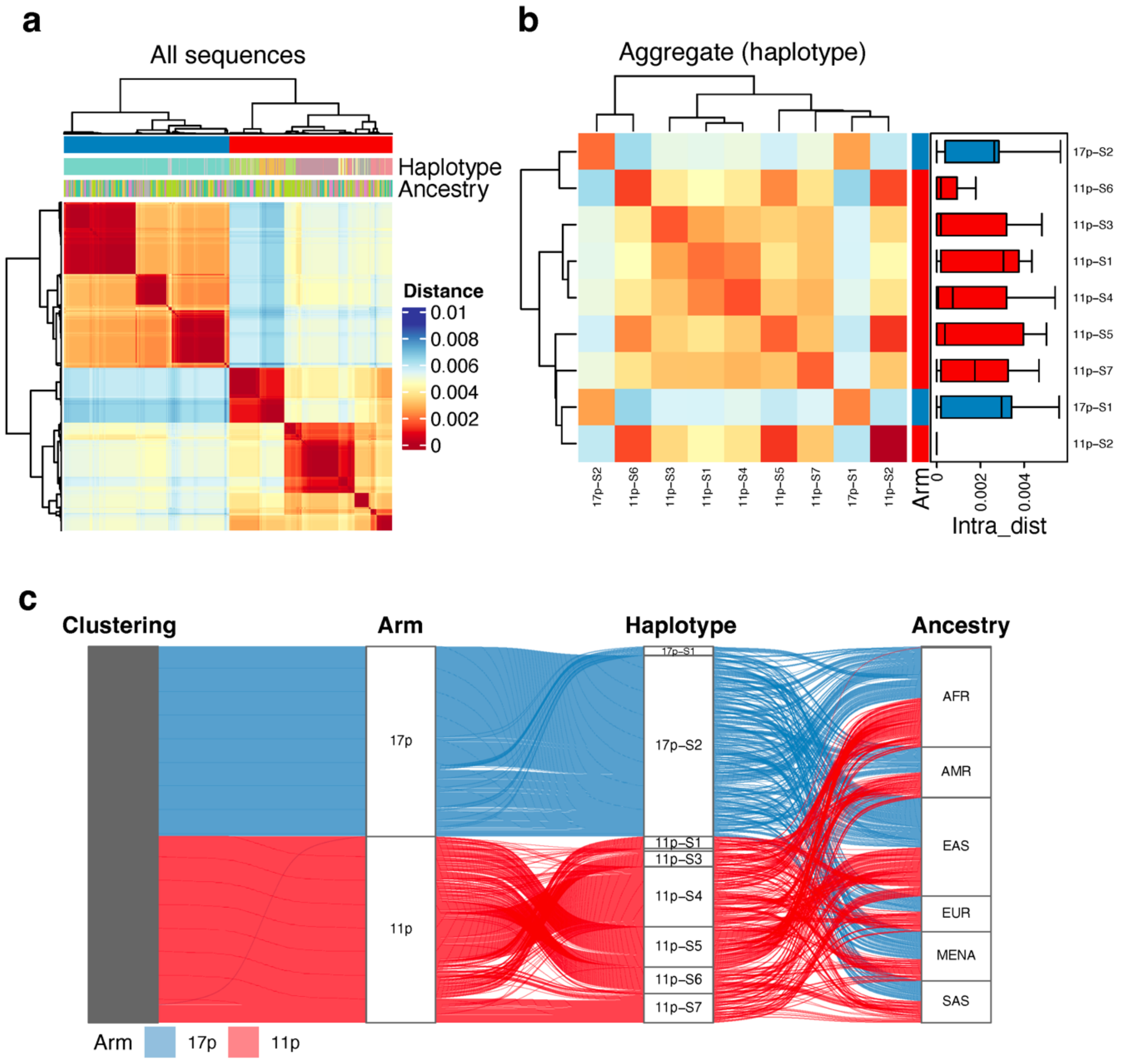


**Supplementary Fig. S87. Pairwise Mash distance analysis for 1000 randomly subsampled subtelomeric of Block 29.** Figure interpretation follows that of Supplementary Figure S58, but for Block 29.

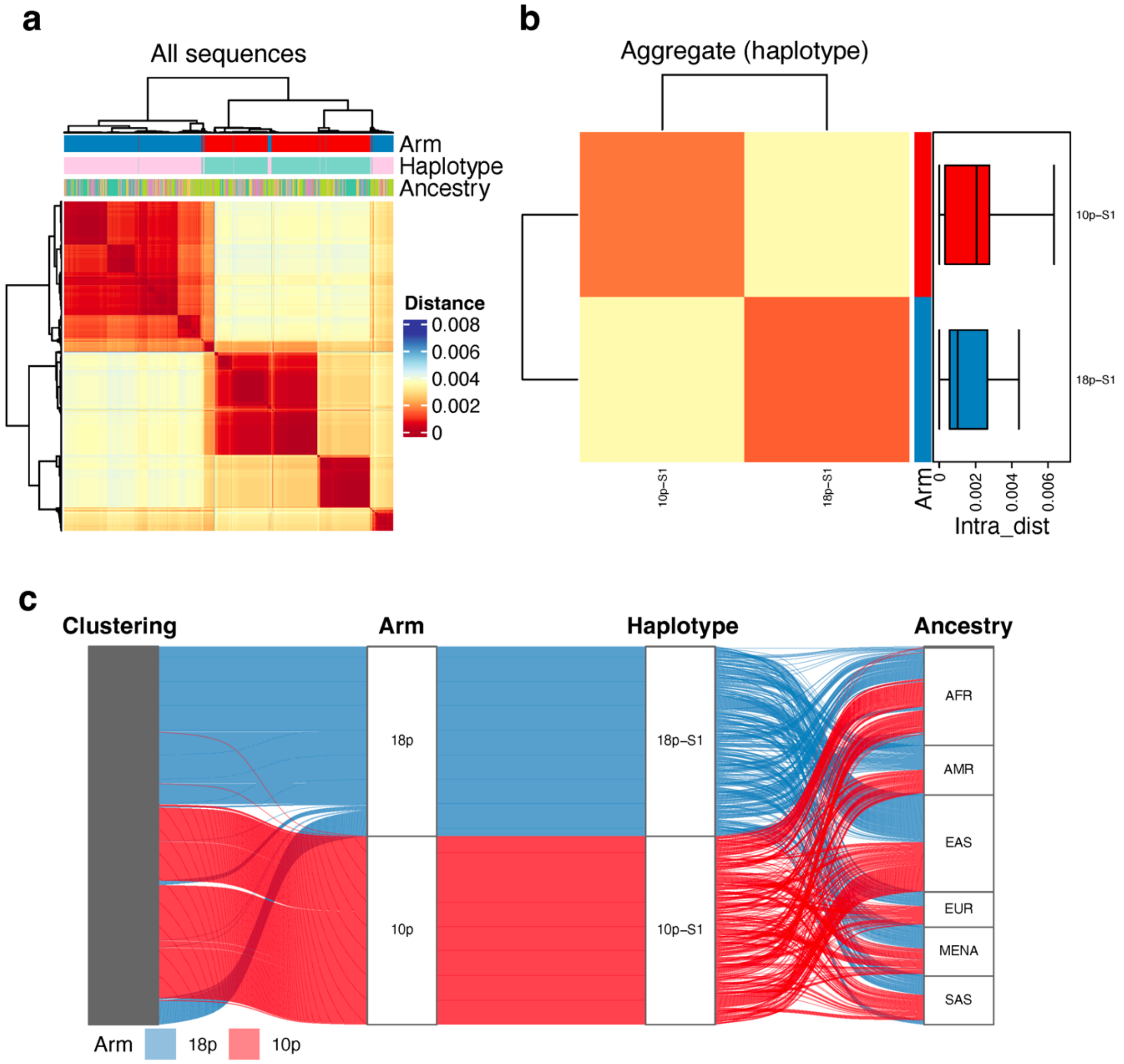


**Supplementary Fig. S88. Pairwise Mash distance analysis for 1000 randomly subsampled subtelomeric of Block 30.** Figure interpretation follows that of Supplementary Figure S58, but for Block 30.

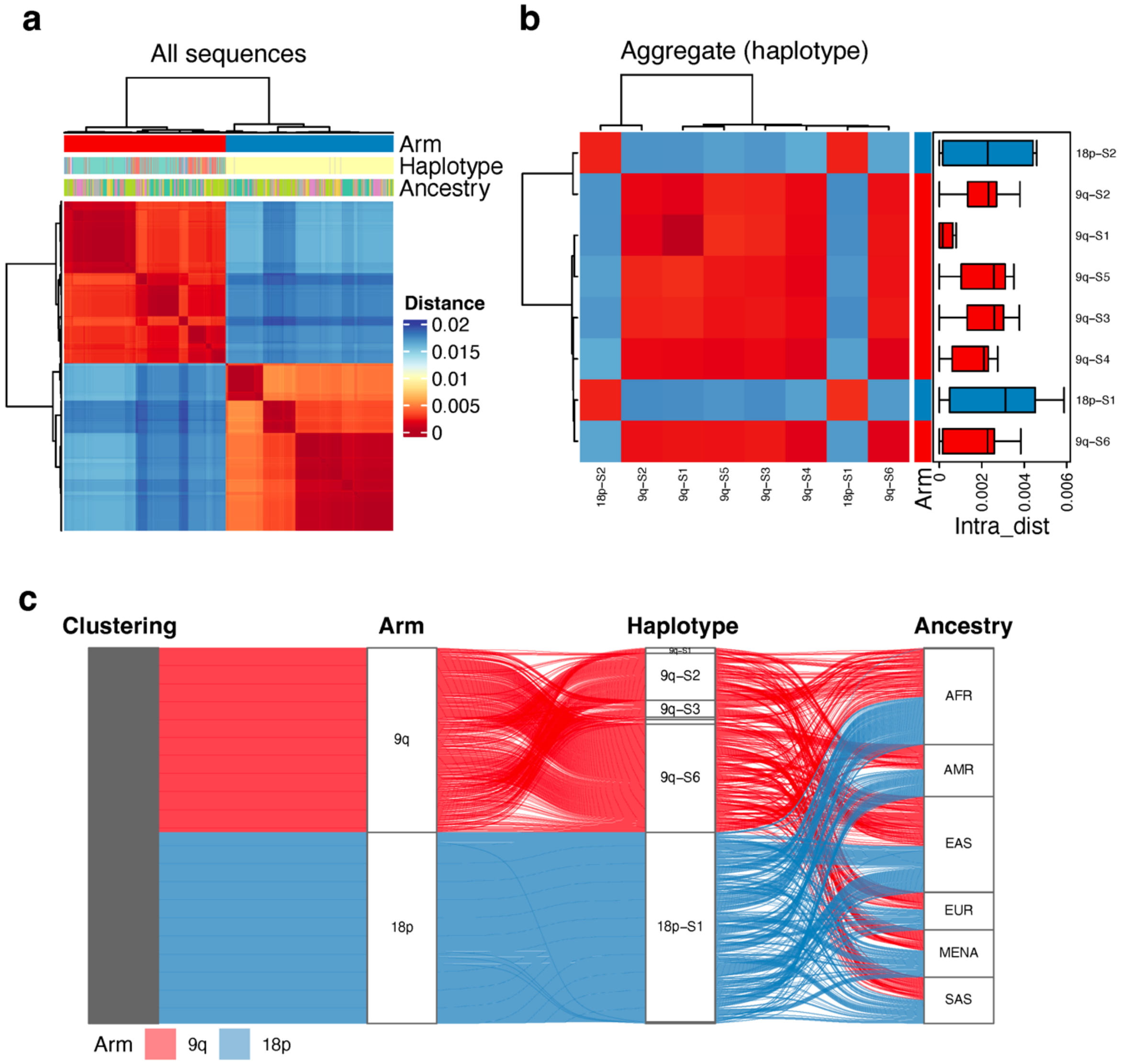


**Supplementary Fig. S89. Pairwise Mash distance analysis for 1000 randomly subsampled subtelomeric of Block 31.** Figure interpretation follows that of Supplementary Figure S58, but for Block 31.

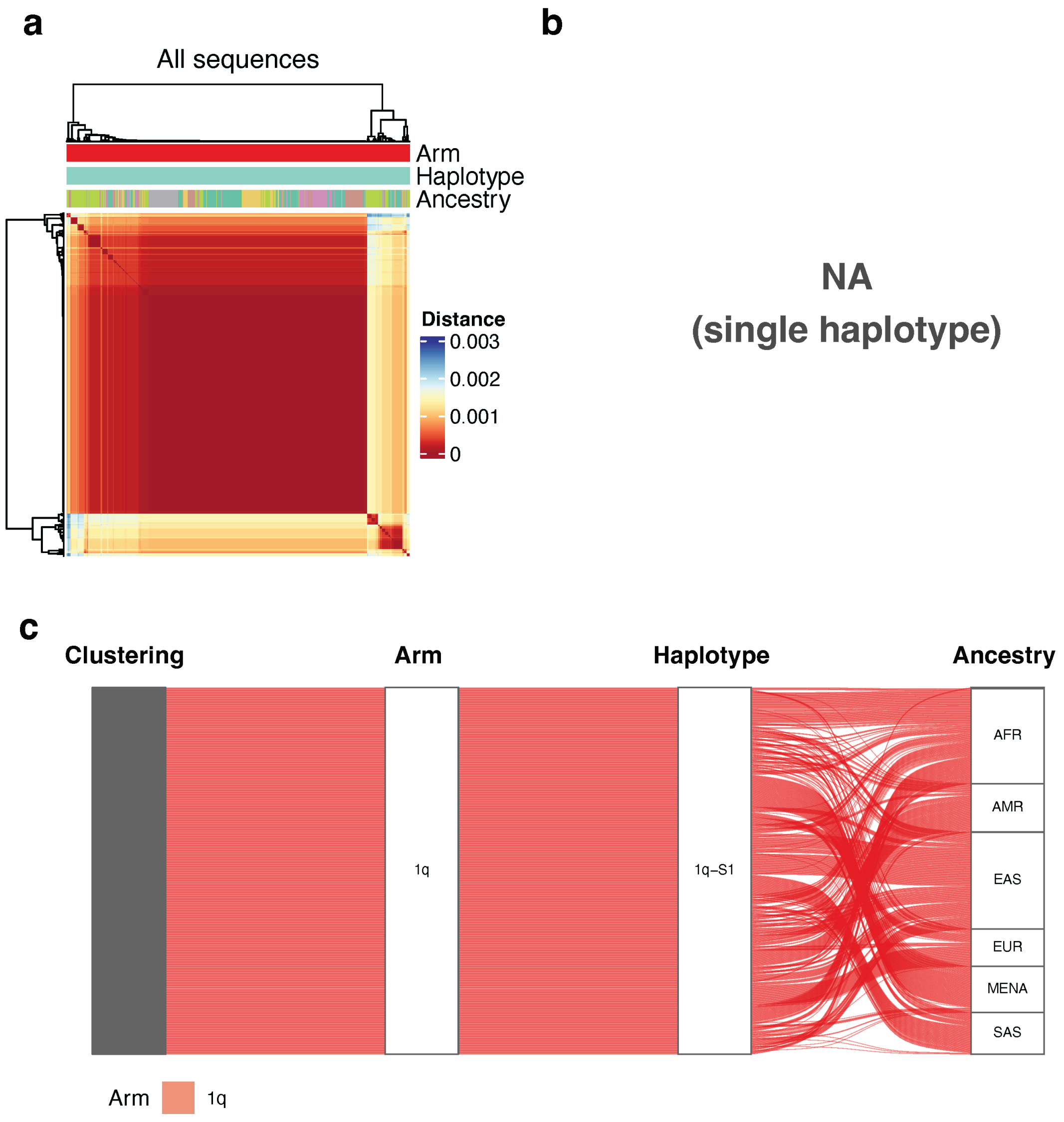


**Supplementary Fig. S90. Pairwise Mash distance analysis for 1000 randomly subsampled subtelomeric of Block 32.** Figure interpretation follows that of Supplementary Figure S58, but for Block 32.

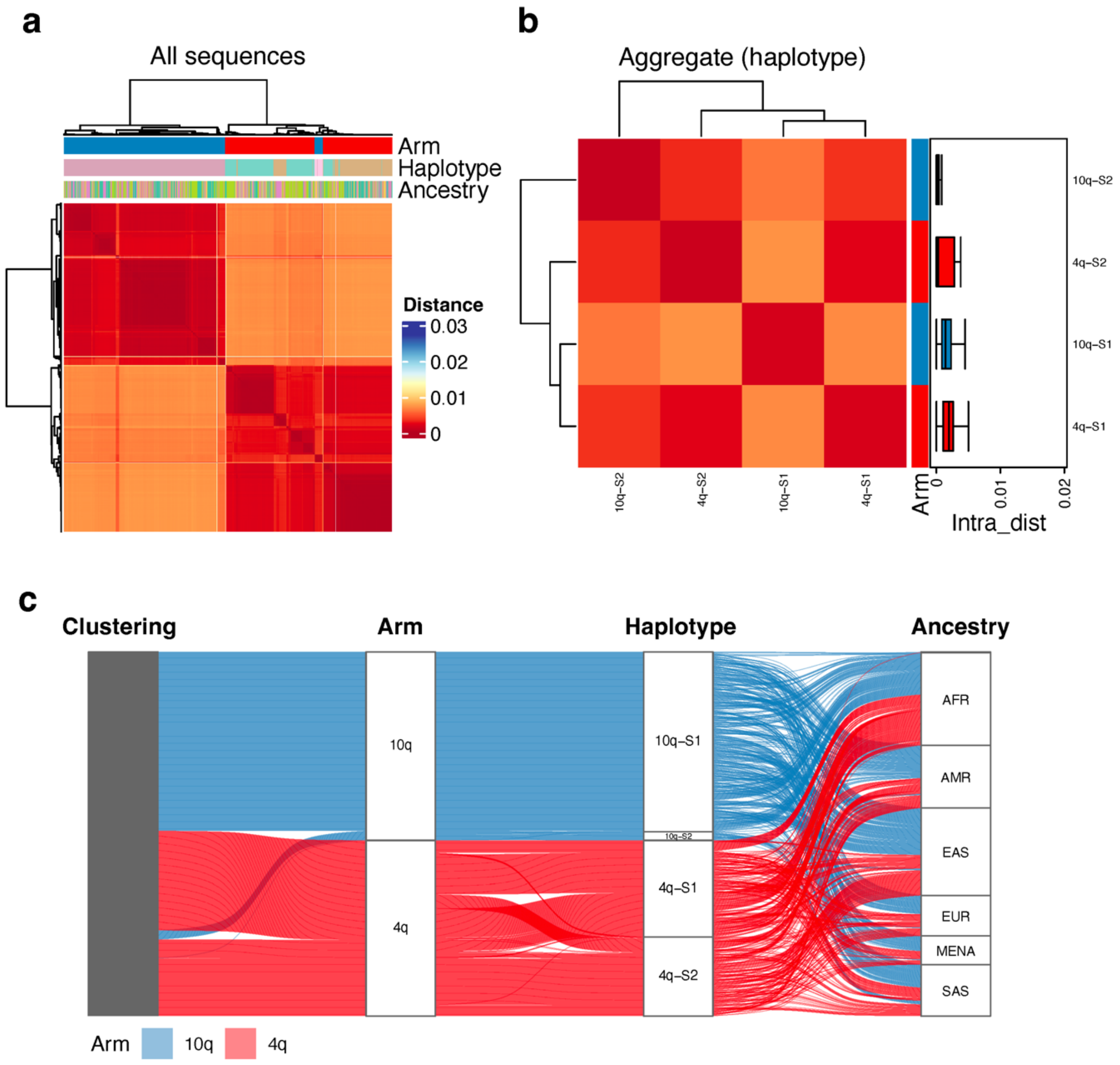


**Supplementary Fig. S91. Pairwise Mash distance analysis for 1000 randomly subsampled subtelomeric of Block 33.** Figure interpretation follows that of Supplementary Figure S58, but for Block 33.

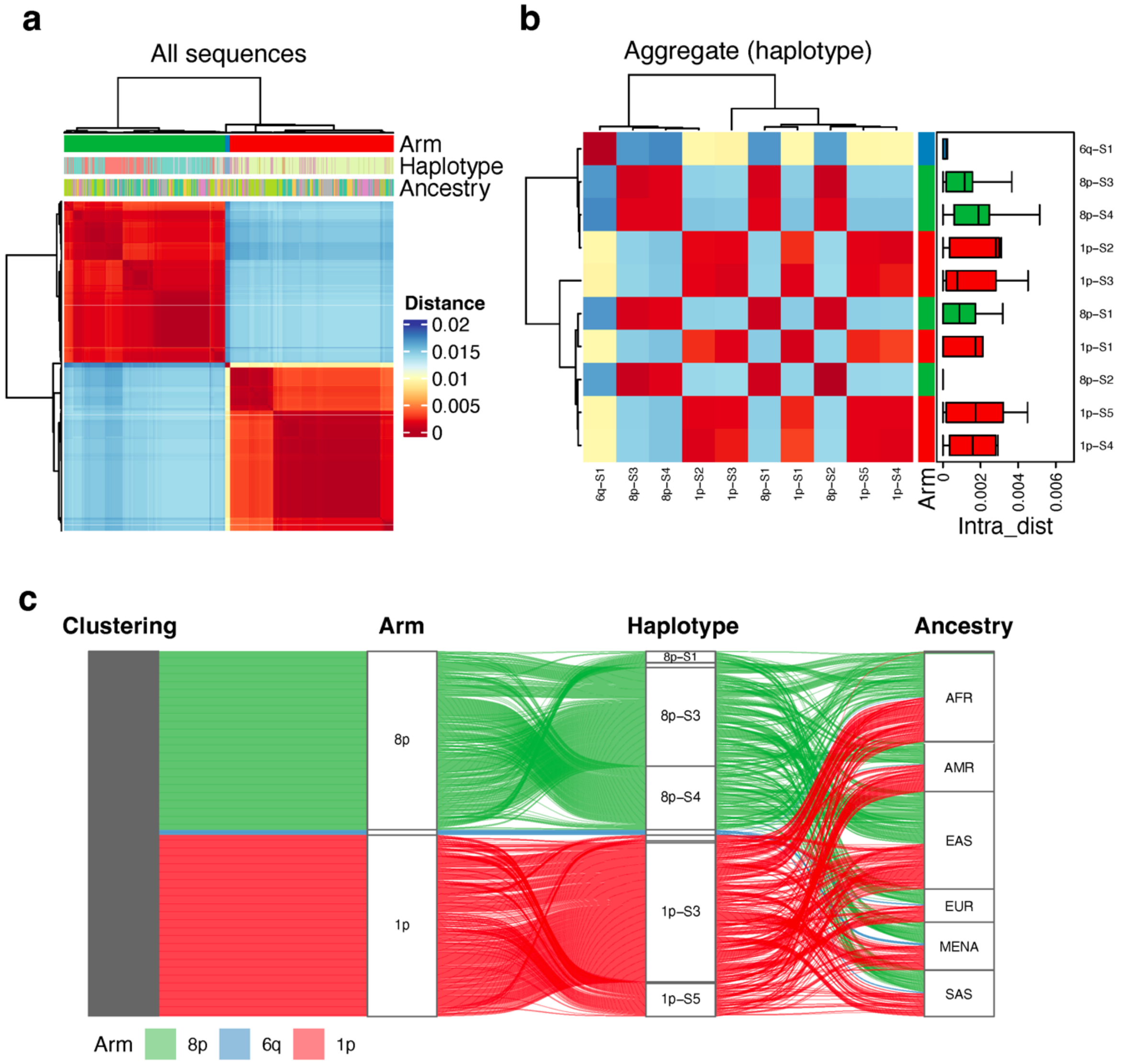


**Supplementary Fig. S92. Pairwise Mash distance analysis for 1000 randomly subsampled subtelomeric of Block 34.** Figure interpretation follows that of Supplementary Figure S58, but for Block 34.

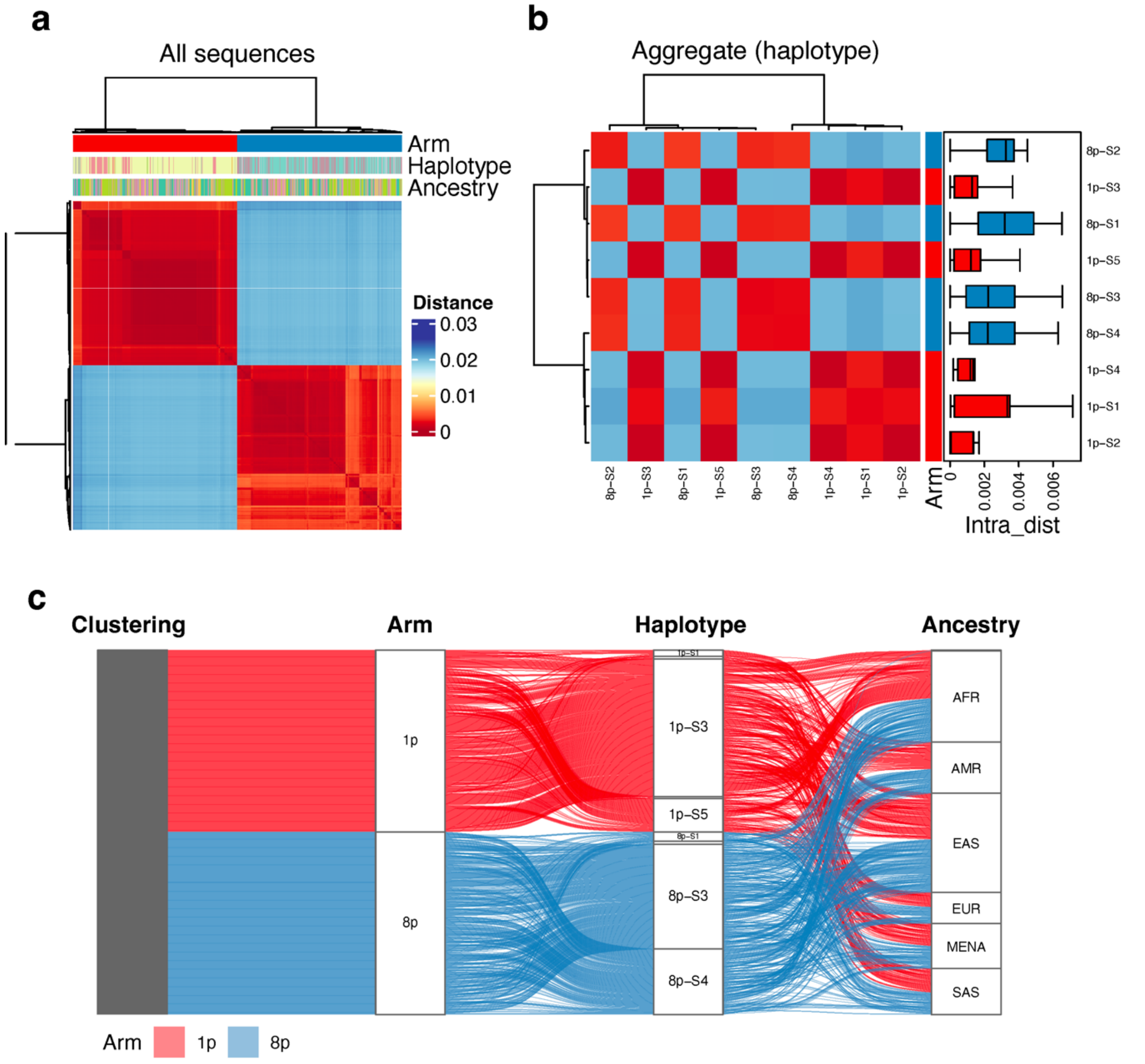


**Supplementary Fig. S93. Pairwise Mash distance analysis for 1000 randomly subsampled subtelomeric of Block 35.** Figure interpretation follows that of Supplementary Figure S58, but for Block 35.

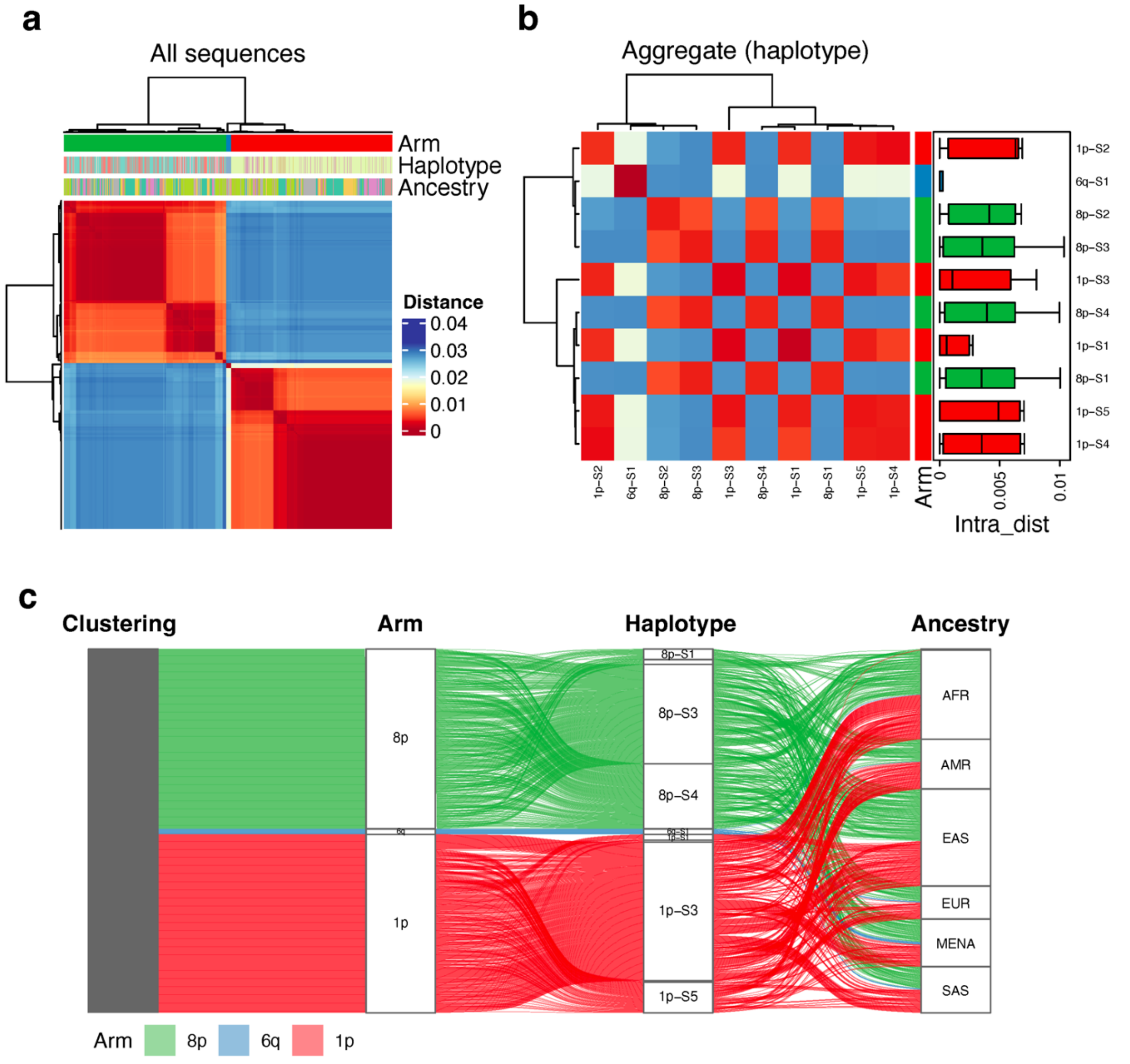


**Supplementary Fig. S94. Pairwise Mash distance analysis for 1000 randomly subsampled subtelomeric of Block 36.** Figure interpretation follows that of Supplementary Figure S58, but for Block 36.

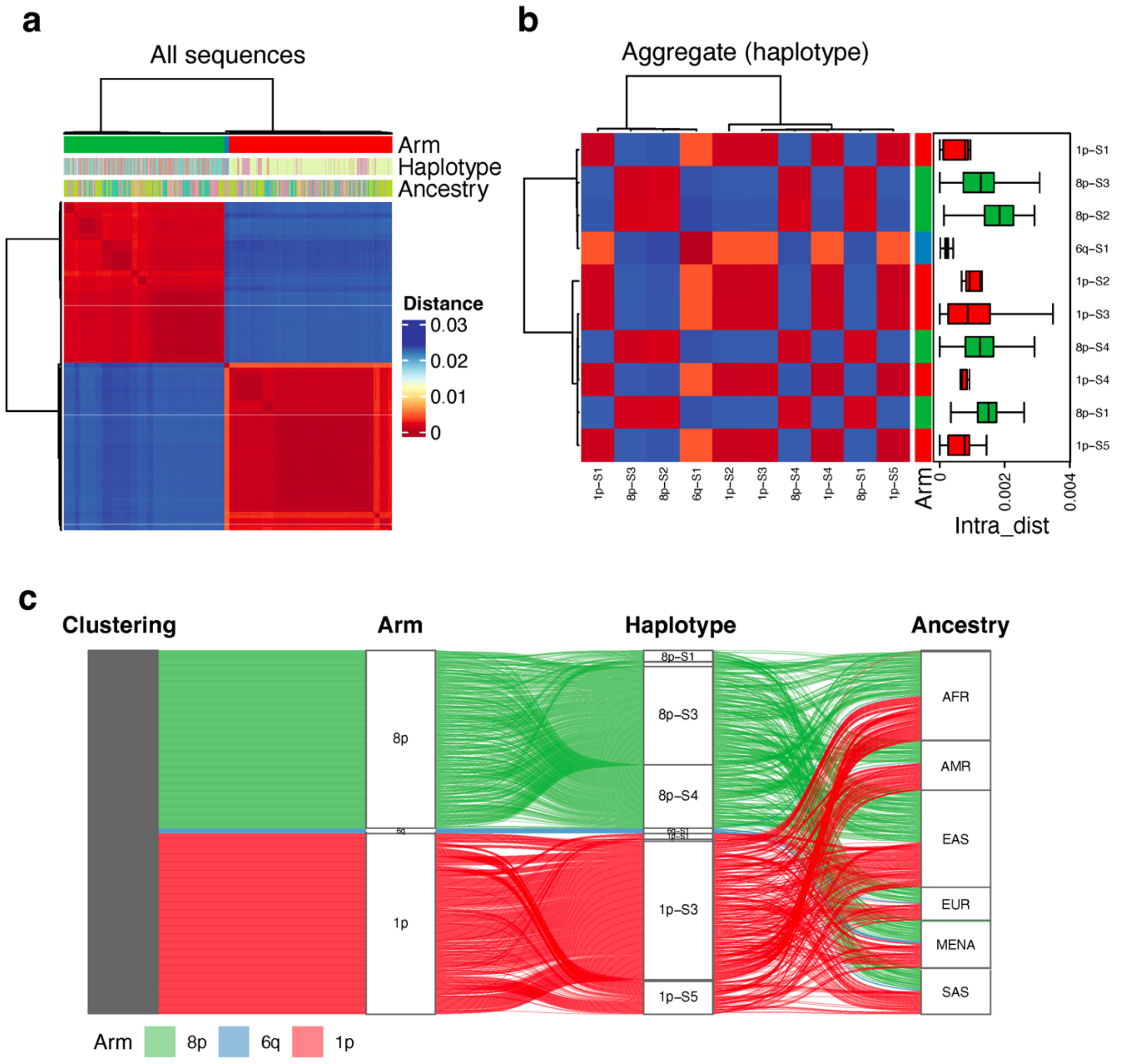


**Supplementary Fig. S95. Pairwise Mash distance analysis for 1000 randomly subsampled subtelomeric of Block 37.** Figure interpretation follows that of Supplementary Figure S58, but for Block 37.

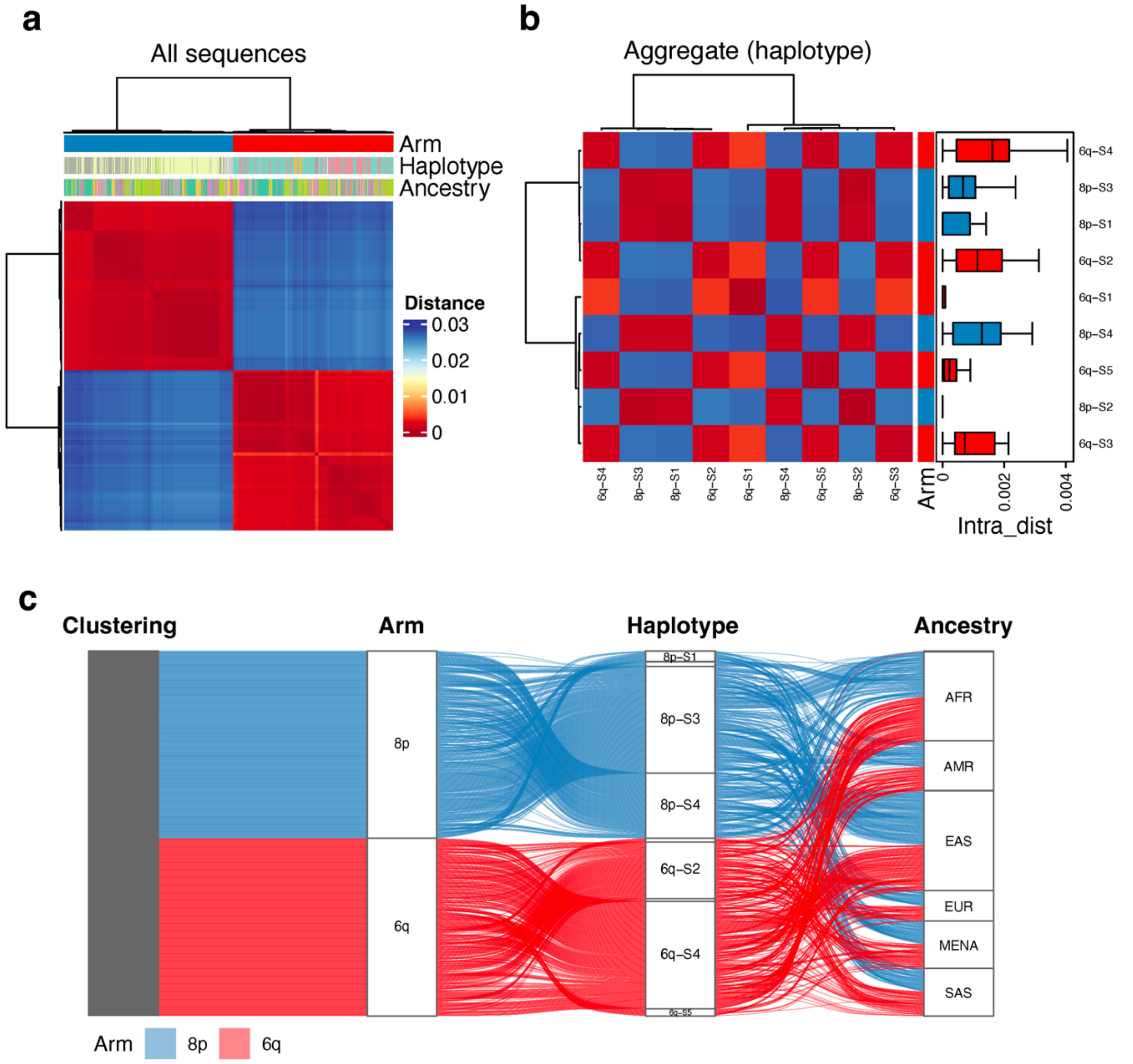


**Supplementary Fig. S96. Pairwise Mash distance analysis for 1000 randomly subsampled subtelomeric of Block 38.** Figure interpretation follows that of Supplementary Figure S58, but for Block 38.

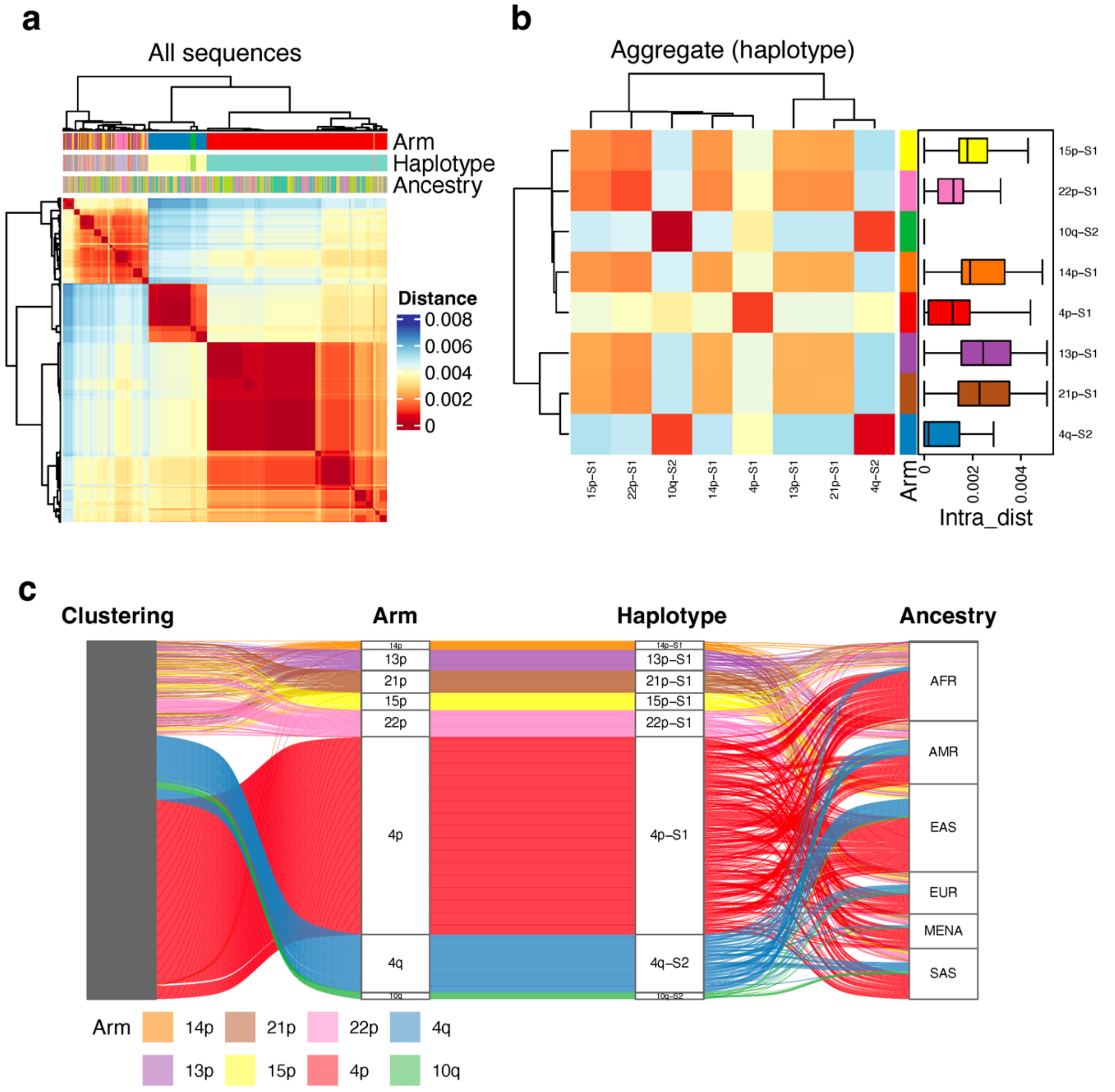


**Supplementary Fig. S97. Pairwise Mash distance analysis for 1000 randomly subsampled subtelomeric of Block 39.** Figure interpretation follows that of Supplementary Figure S58, but for Block 39.

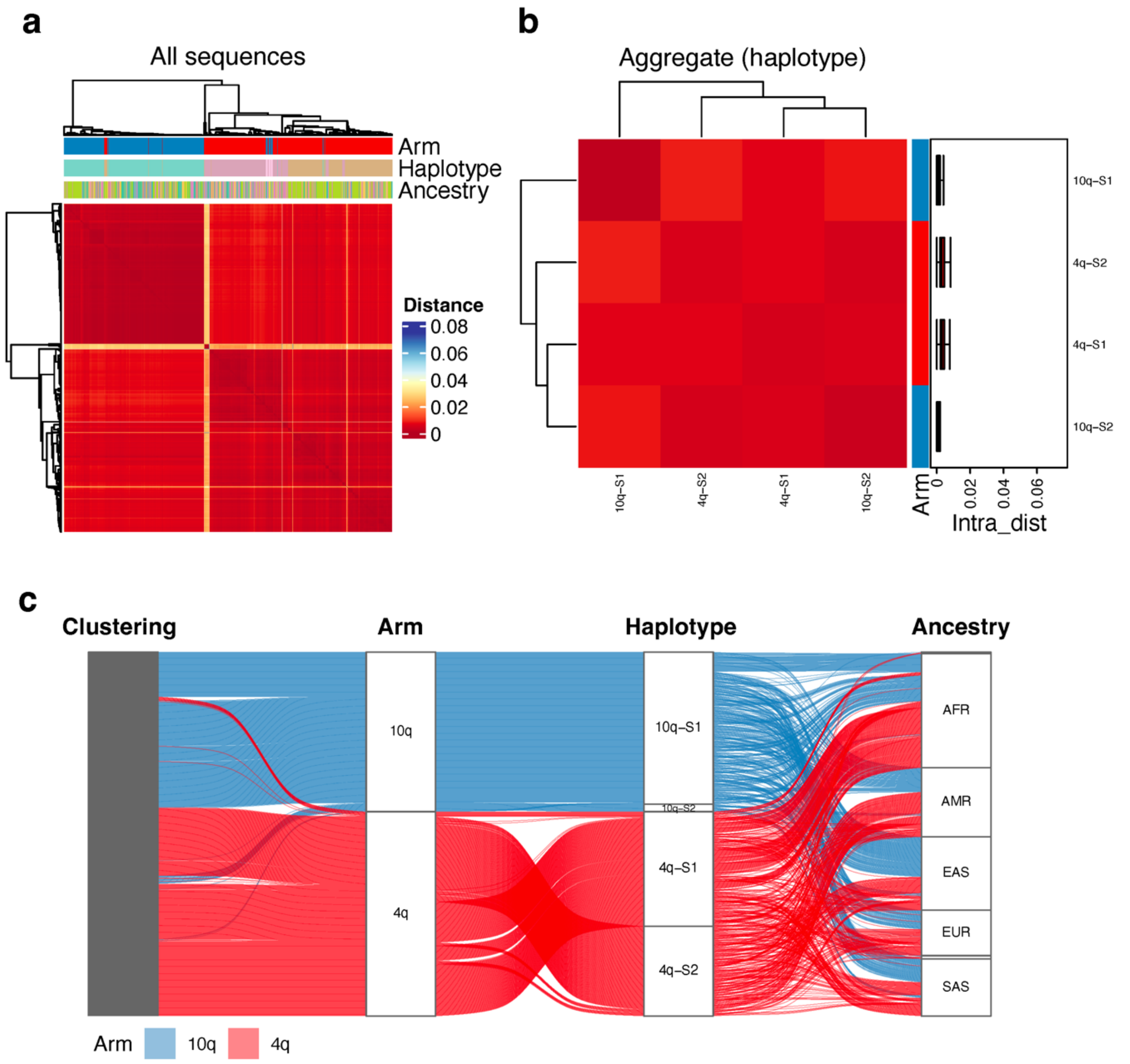


**Supplementary Fig. S98. Pairwise Mash distance analysis for 1000 randomly subsampled subtelomeric of Block 40.** Figure interpretation follows that of Supplementary Figure S58, but for Block 40.

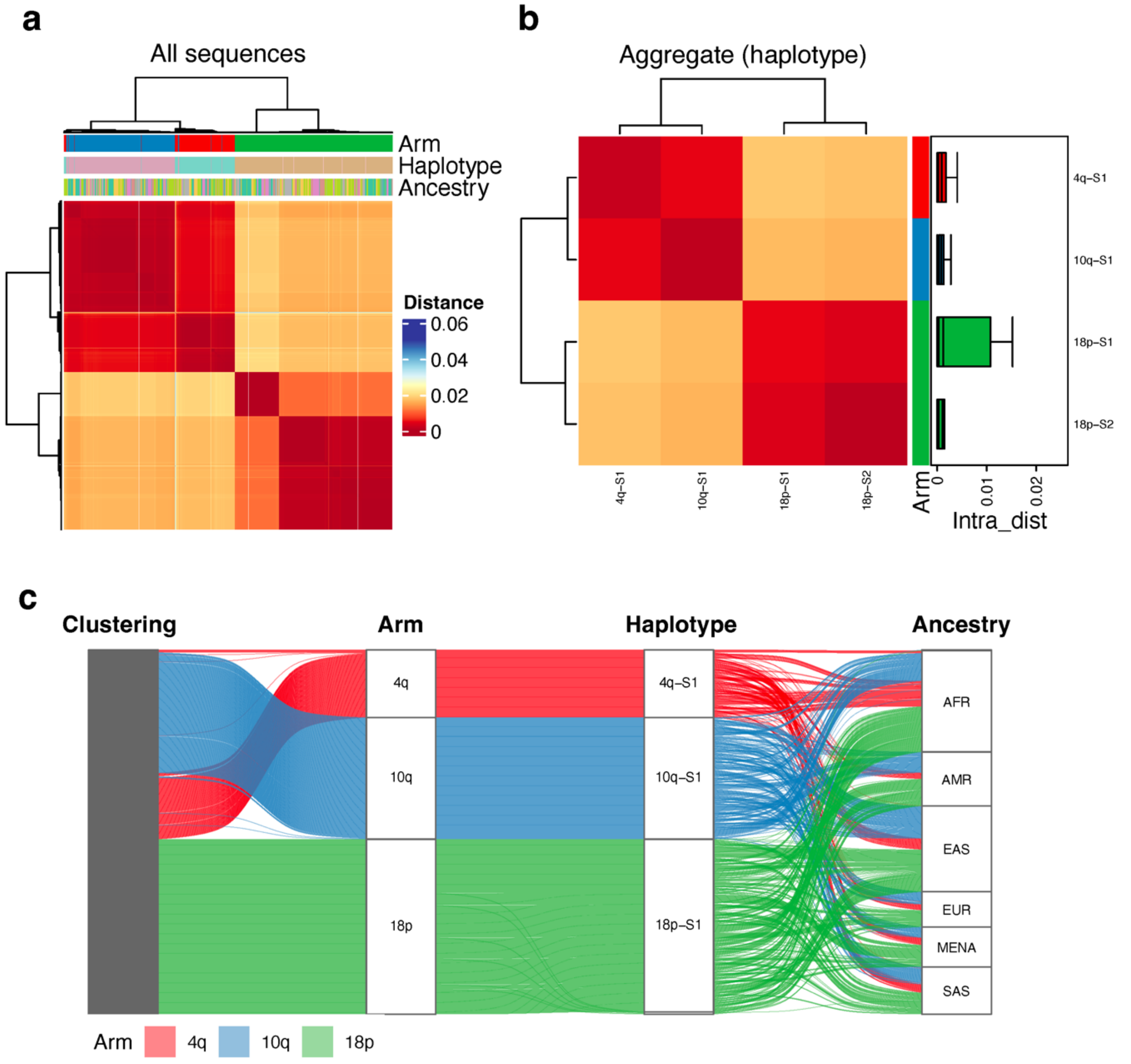


**Supplementary Fig. S99. Pairwise Mash distance analysis for 1000 randomly subsampled subtelomeric of Block 41.** Figure interpretation follows that of Supplementary Figure S58, but for Block 41.

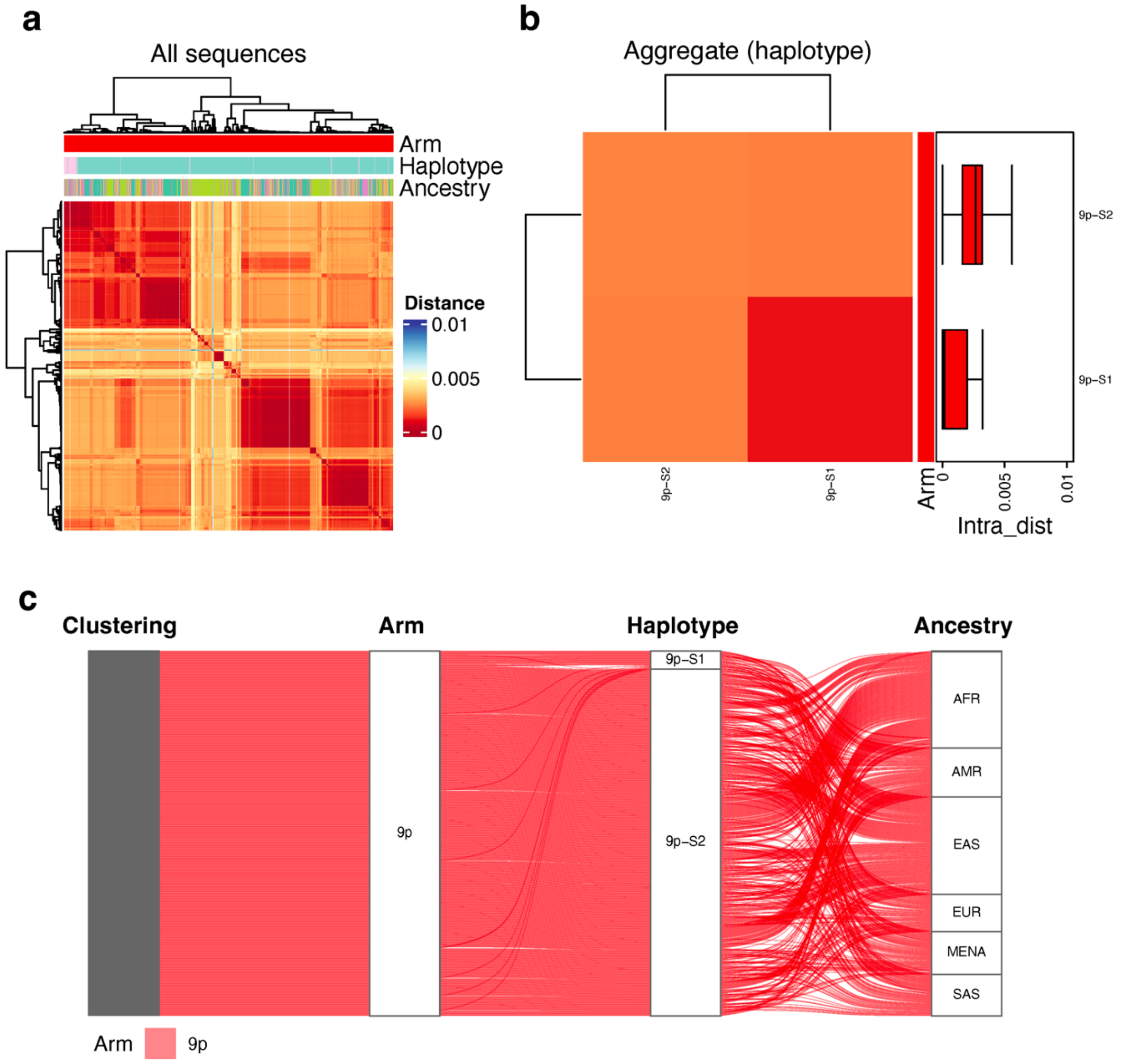


**Supplementary Fig. S100. Pairwise Mash distance analysis for 1000 randomly subsampled subtelomeric of Block 43.** Figure interpretation follows that of Supplementary Figure S58, but for Block 43.

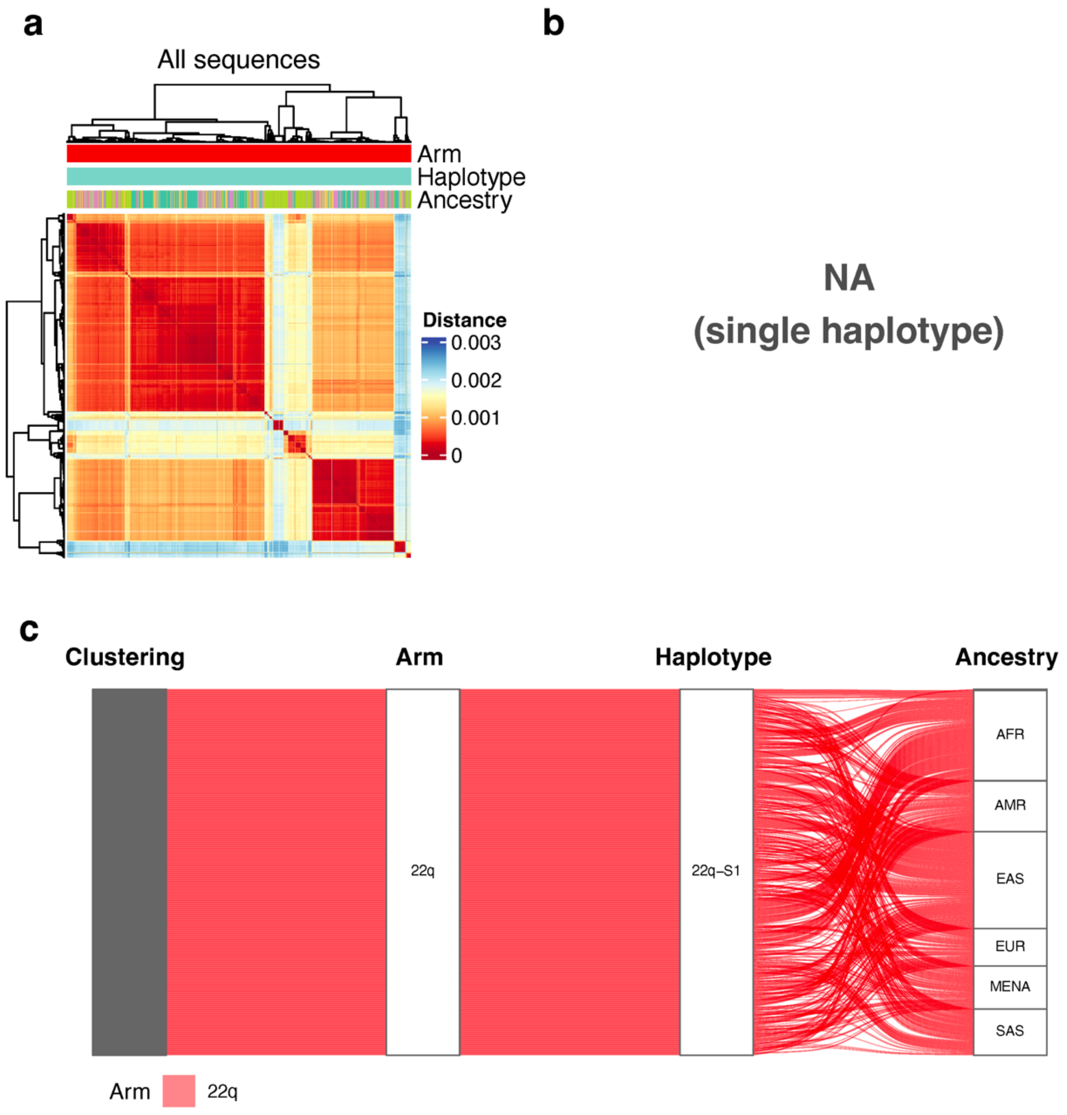


**Supplementary Fig. S101. Pairwise Mash distance analysis for 1000 randomly subsampled subtelomeric of Block 44.** Figure interpretation follows that of Supplementary Figure S58, but for Block 44.

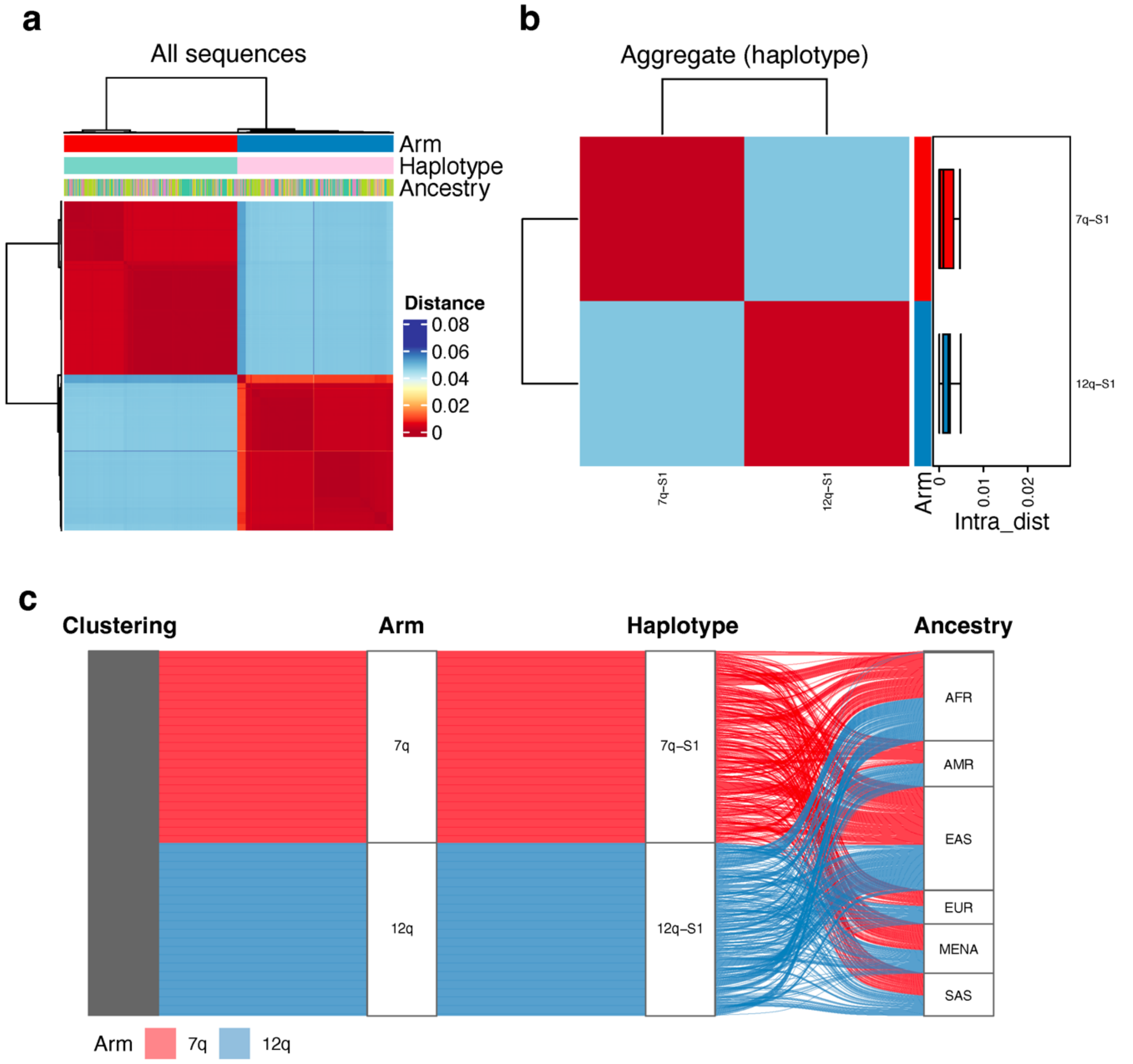


**Supplementary Fig. S102. Pairwise Mash distance analysis for 1000 randomly subsampled subtelomeric of Block 45.** Figure interpretation follows that of Supplementary Figure S58, but for Block 45.

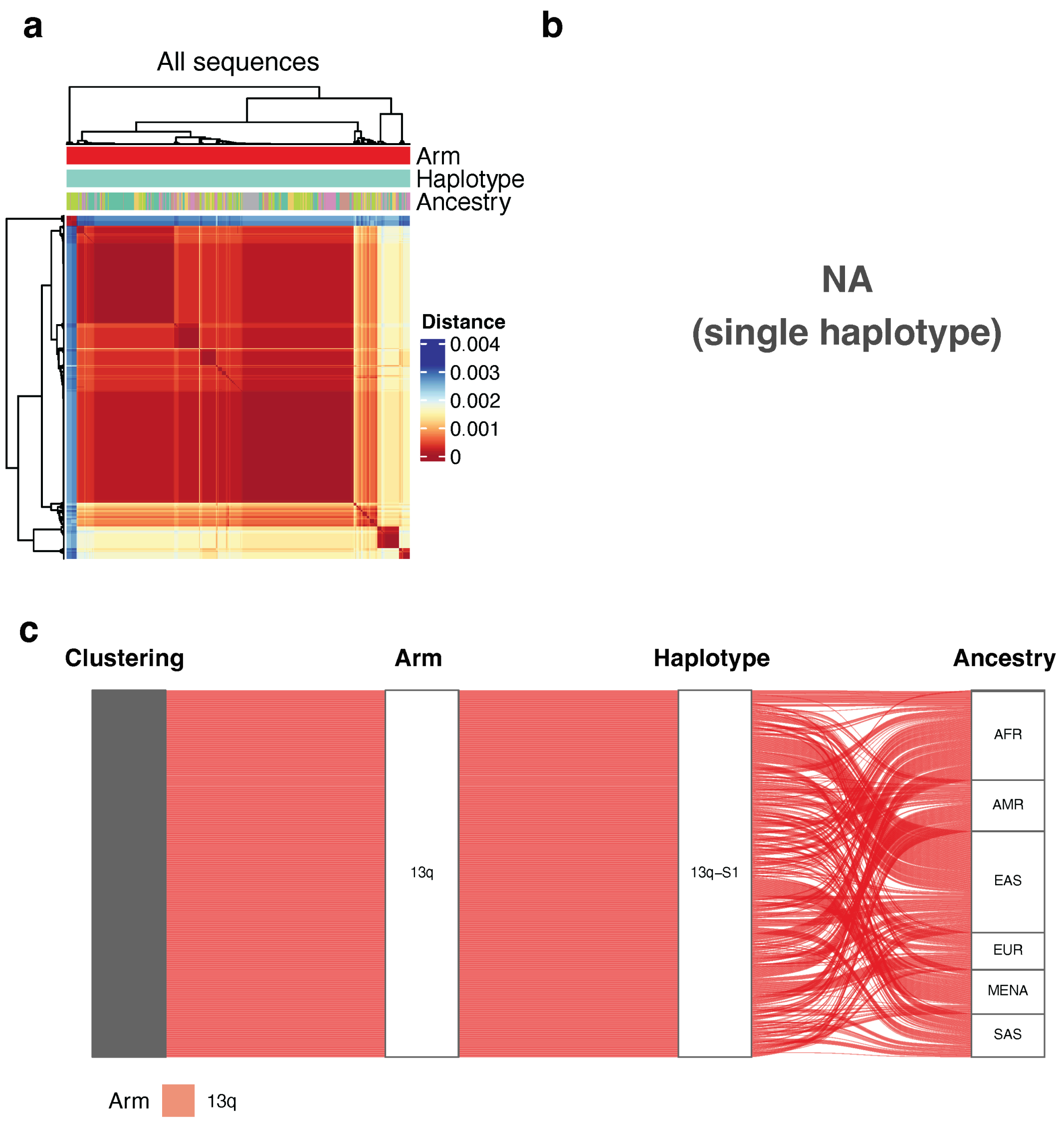


**Supplementary Fig. S103. Pairwise Mash distance analysis for 1000 randomly subsampled subtelomeric of Block 46.** Figure interpretation follows that of Supplementary Figure S58, but for Block 46.

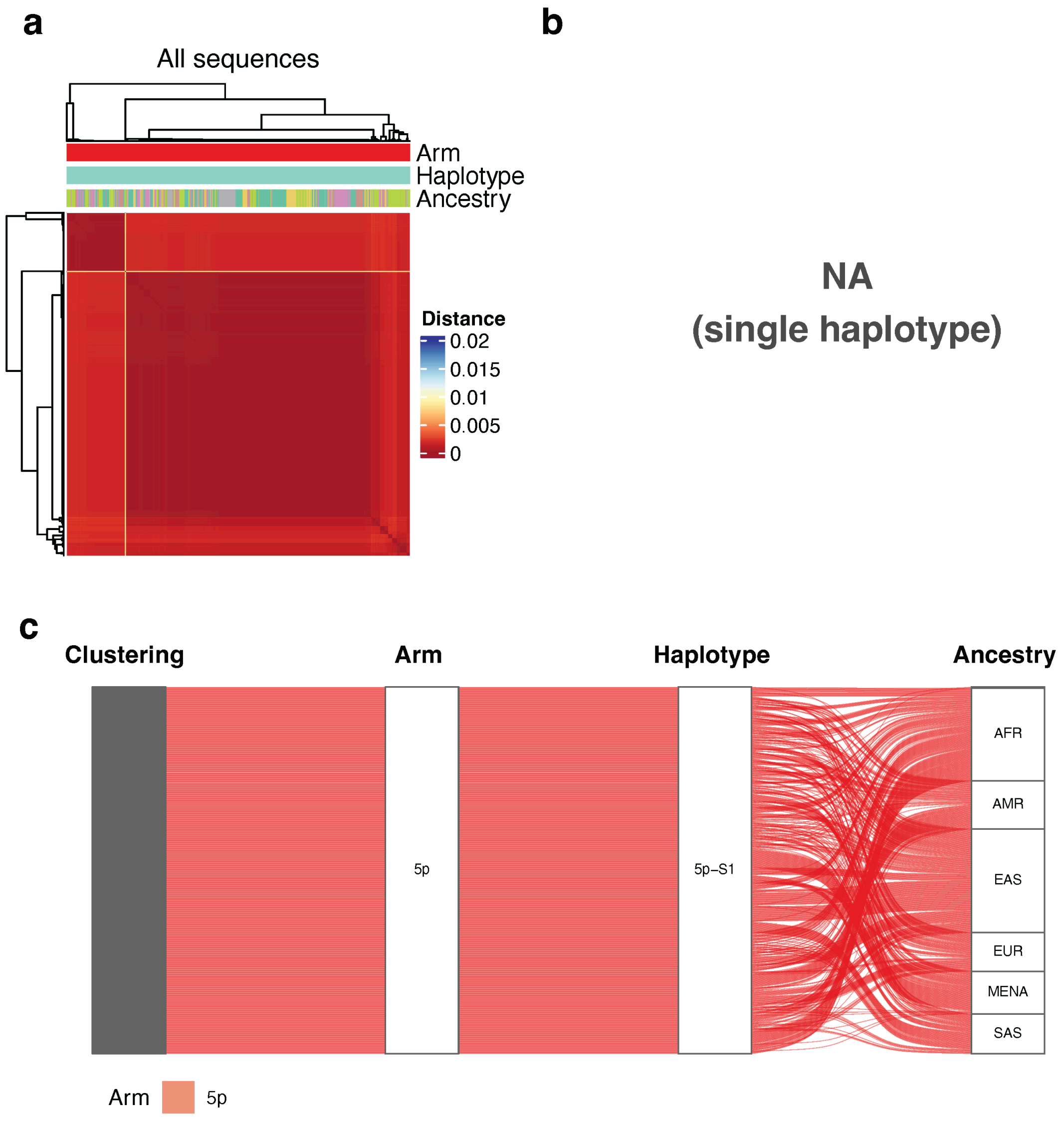


**Supplementary Fig. S104. Pairwise Mash distance analysis for 1000 randomly subsampled subtelomeric of Block 47.** Figure interpretation follows that of Supplementary Figure S58, but for Block 47.

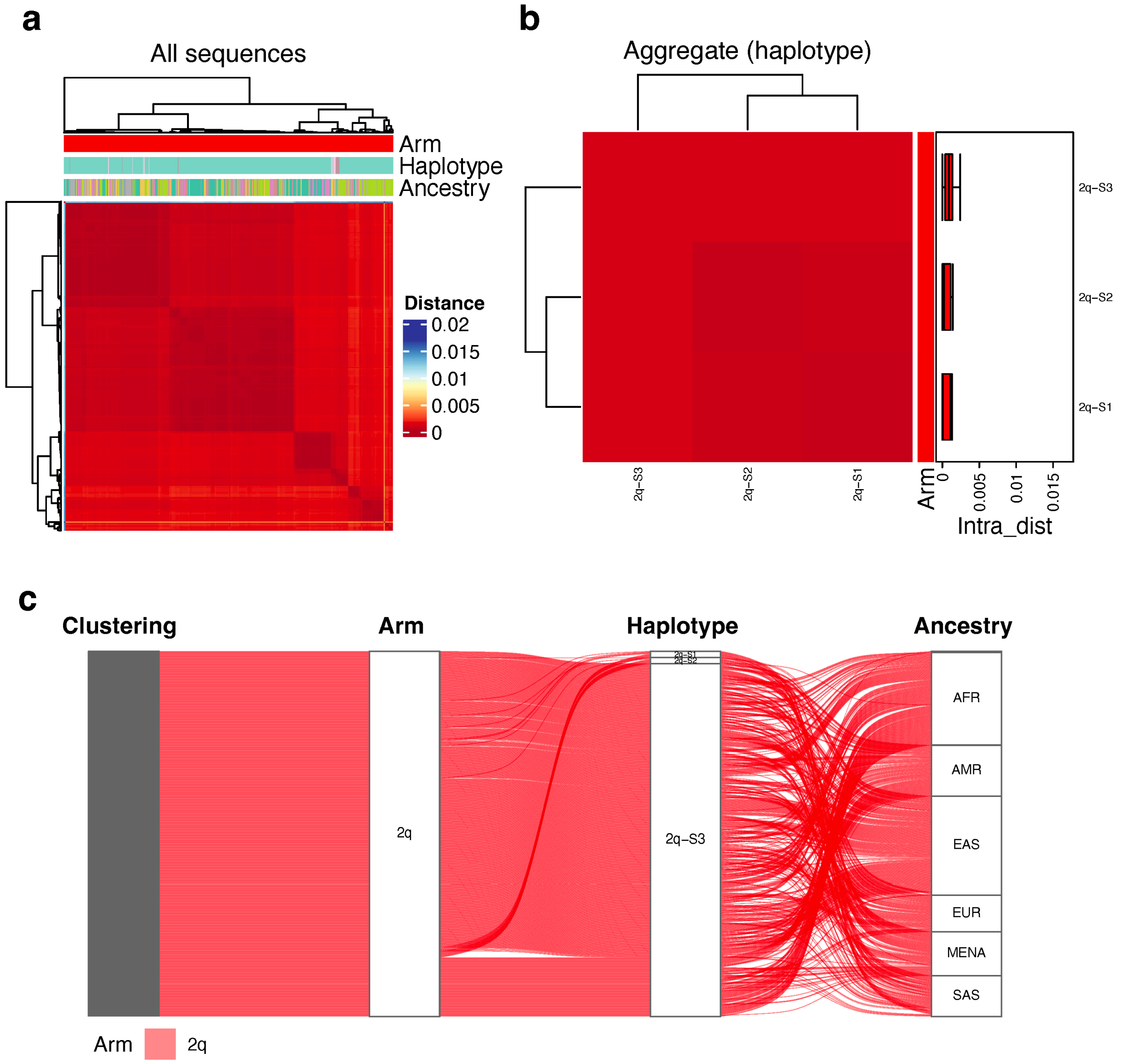


**Supplementary Fig. S105. Pairwise Mash distance analysis for 1000 randomly subsampled subtelomeric of Block 48.** Figure interpretation follows that of Supplementary Figure S58, but for Block 48.

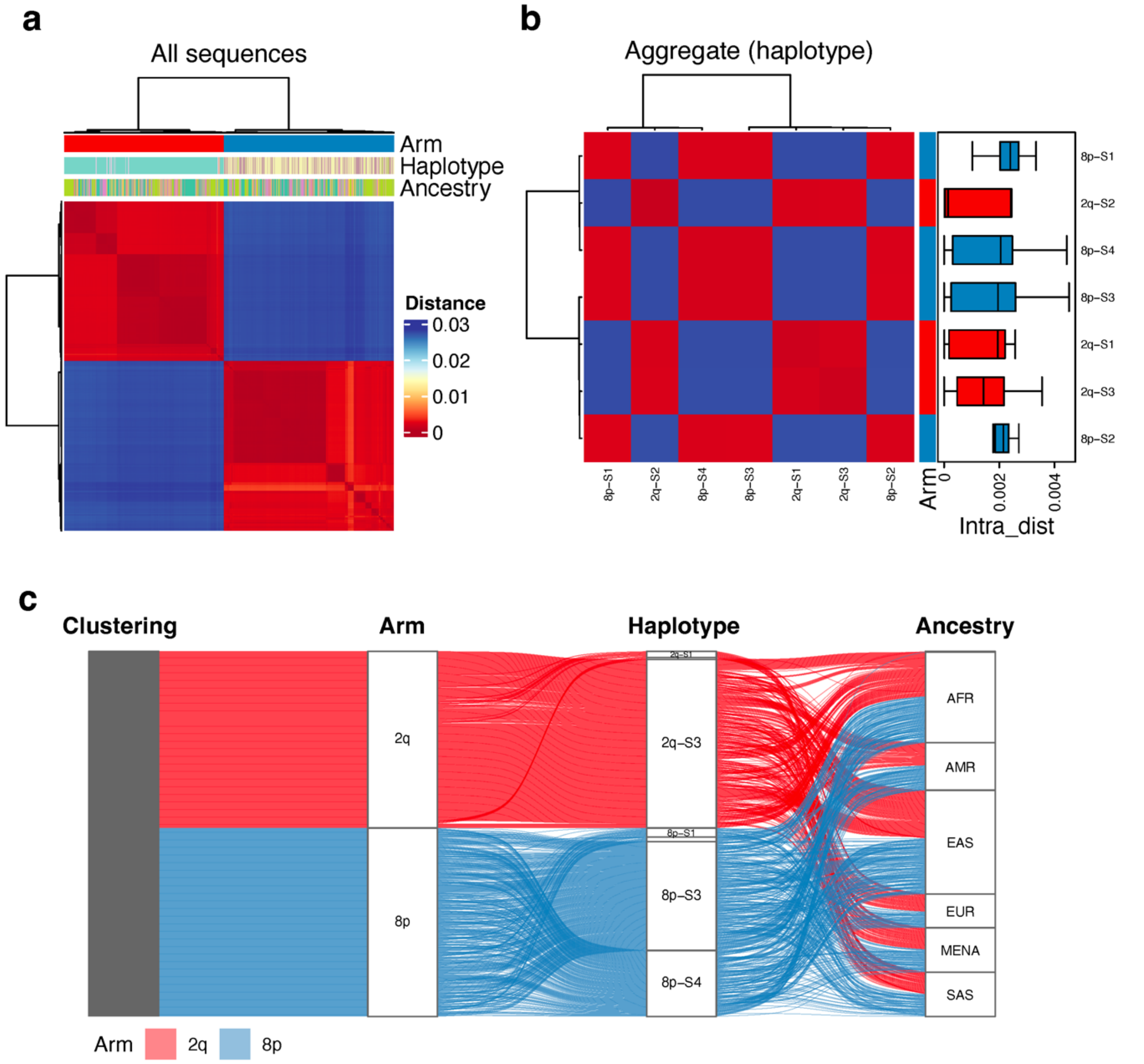


**Supplementary Fig. S106. Pairwise Mash distance analysis for 1000 randomly subsampled subtelomeric of Block 49.** Figure interpretation follows that of Supplementary Figure S58, but for Block 49.

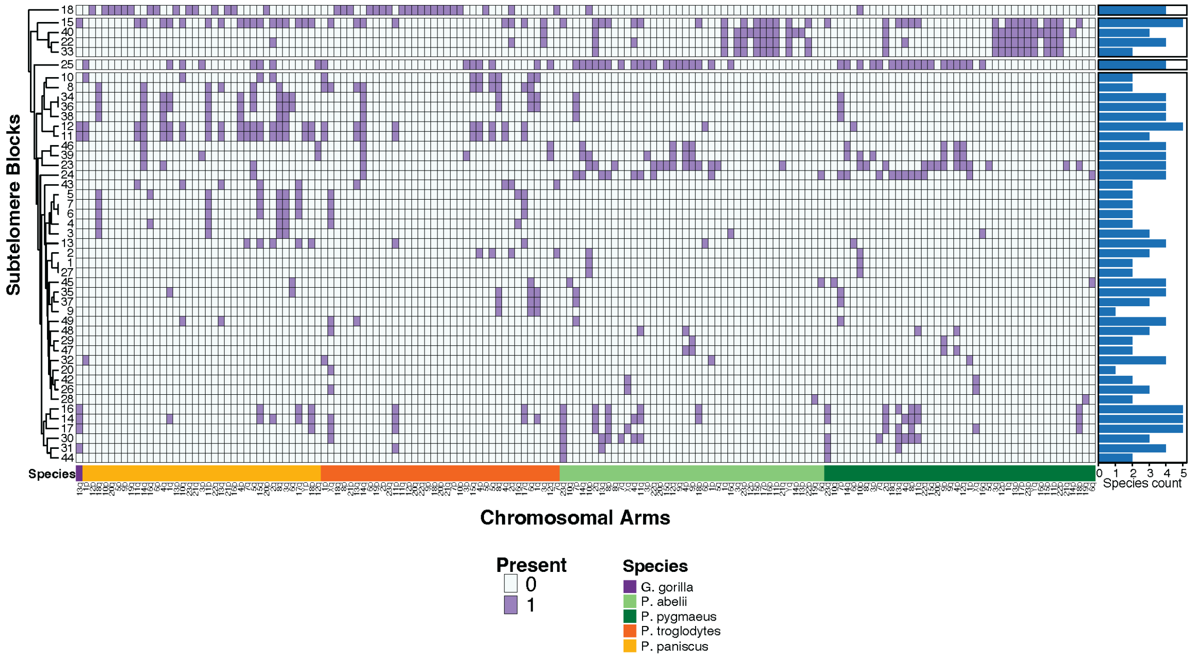


**Supplementary Fig. S107. Subtelomere block analysis in apes.** Heatmap of human subtelomeric block presence (Stong et al. 2014) across five ape species: western gorilla (Gorilla gorilla; purple), bonobo (Pan paniscus; yellow), chimpanzee (Pan troglodytes; Orange), Sumatran orangutan (Pongo abelii; light green), and Bornean orangutan (Pongo pygmaeus; dark green). Rows were clustered into four groups. The bar chart on the right indicates the number of ape species (out of five) in which each subtelomeric block was detected.

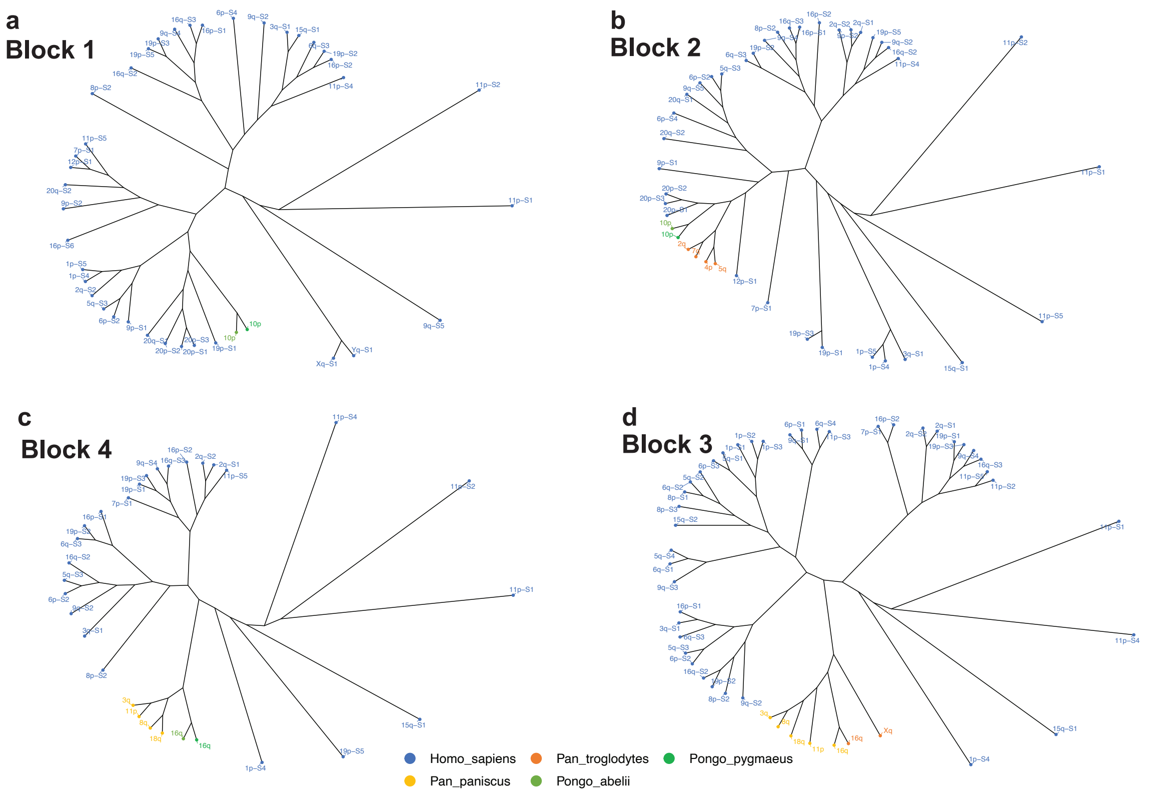


**Supplementary Fig. S108. Maximum likelihood phylogenetic trees for subtelomeric blocks in human and great apes for Blocks 1-4.** (a) Block 1, (b) Block 2, (c) Block 3 and (d) Block 4. Tip labels indicate chromosomal arm of origin, coloured by species. Human sequences represent representative sequences from each haplotype.

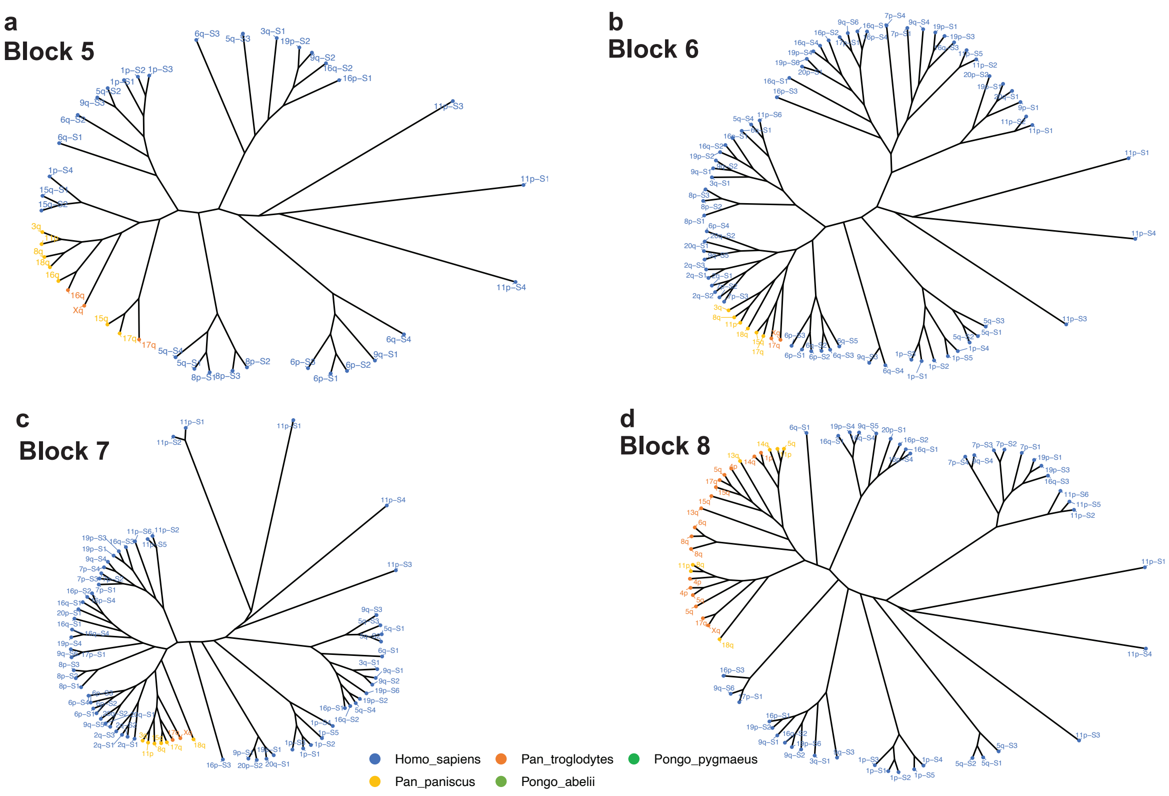


**Supplementary Fig. S109. Maximum likelihood phylogenetic trees for subtelomeric blocks in human and great apes for Block 5-8.** (a) Block 5, (b) Block 6, (c) Block 7 and (d) Block 8. Tip labels indicate chromosomal arm of origin, coloured by species. Human sequences represent representative sequences from each haplotype.

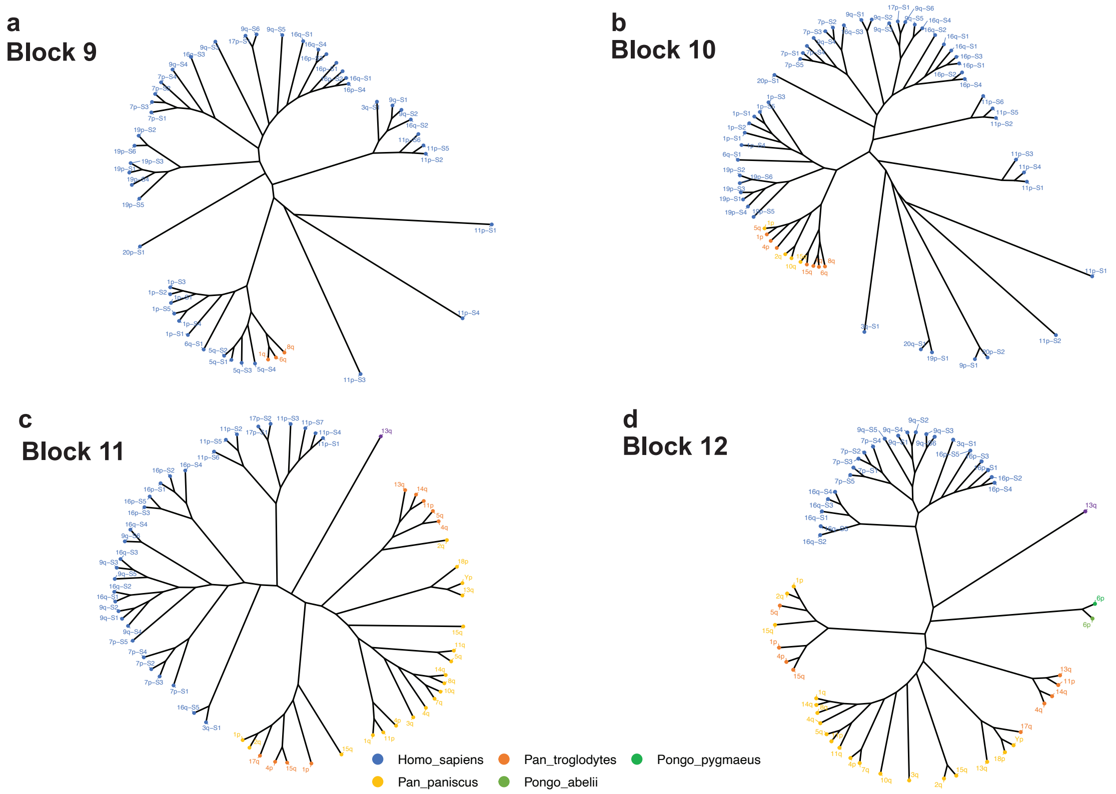


**Supplementary Fig. S110. Maximum likelihood phylogenetic trees for subtelomeric blocks in human and great apes for Block 9-12.** (a) Block 9, (b) Block 10, (c) Block 11 and (d) Block 12. Tip labels indicate chromosomal arm of origin, coloured by species. Human sequences represent representative sequences from each haplotype.

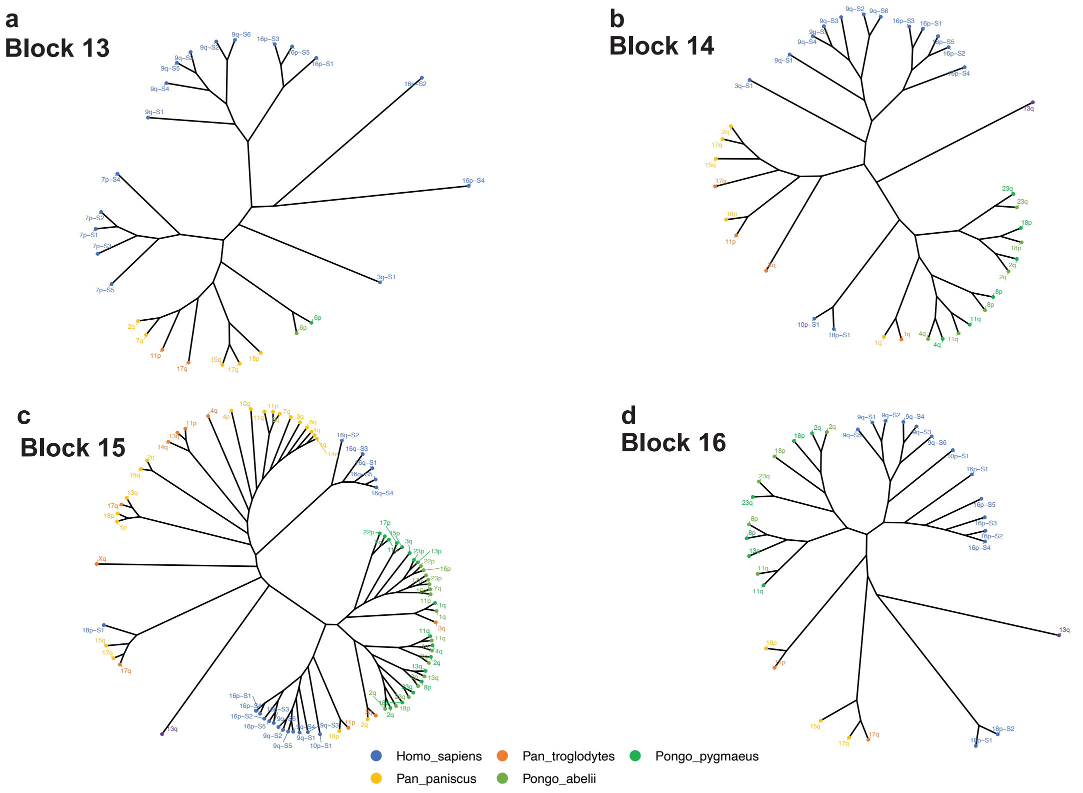


**Supplementary Fig. S111. Maximum likelihood phylogenetic trees for subtelomeric blocks in human and great apes for Block 13-16.** (a) Block 13, (b) Block 14, (c) Block 15 and (d) Block 16. Tip labels indicate chromosomal arm of origin, coloured by species. Human sequences represent representative sequences from each haplotype.

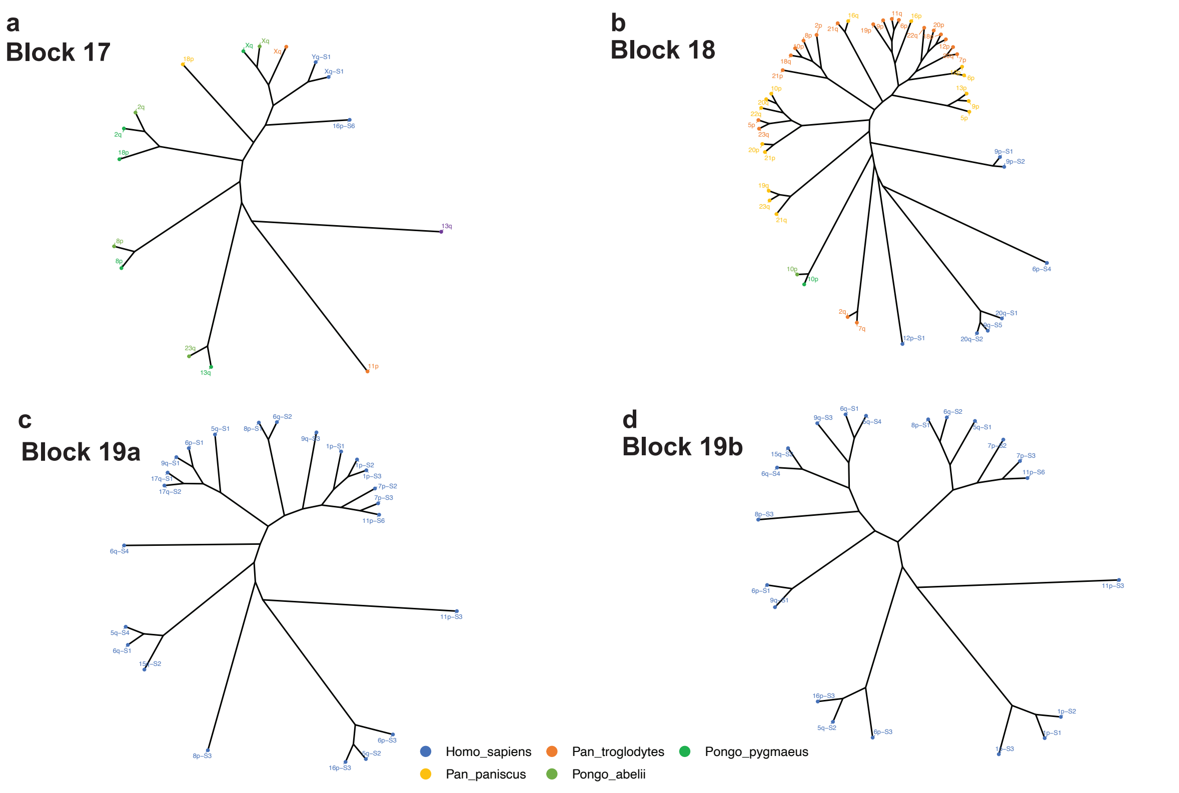


**Supplementary Fig. S112. Maximum likelihood phylogenetic trees for subtelomeric blocks in human and great apes for Block 17-19b.** (a) Block 17, (b) Block 18, (c) Block 19a and (d) Block 19b. Tip labels indicate chromosomal arm of origin, coloured by species. Human sequences represent representative sequences from each haplotype.

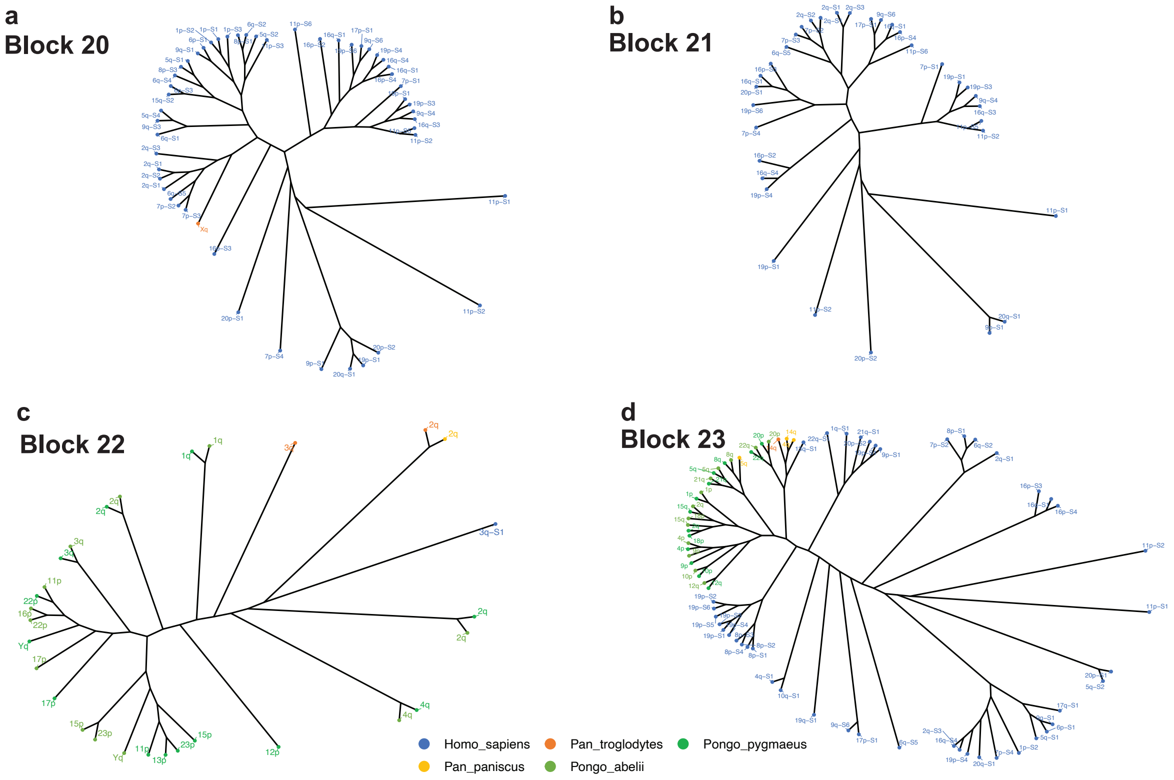


**Supplementary Fig. S113. Maximum likelihood phylogenetic trees for subtelomeric blocks in human and great apes for Block 20-23.** (a) Block 20, (b) Block 21, (c) Block 22 and (d) Block 23. Tip labels indicate chromosomal arm of origin, coloured by species. Human sequences represent representative sequences from each haplotype.

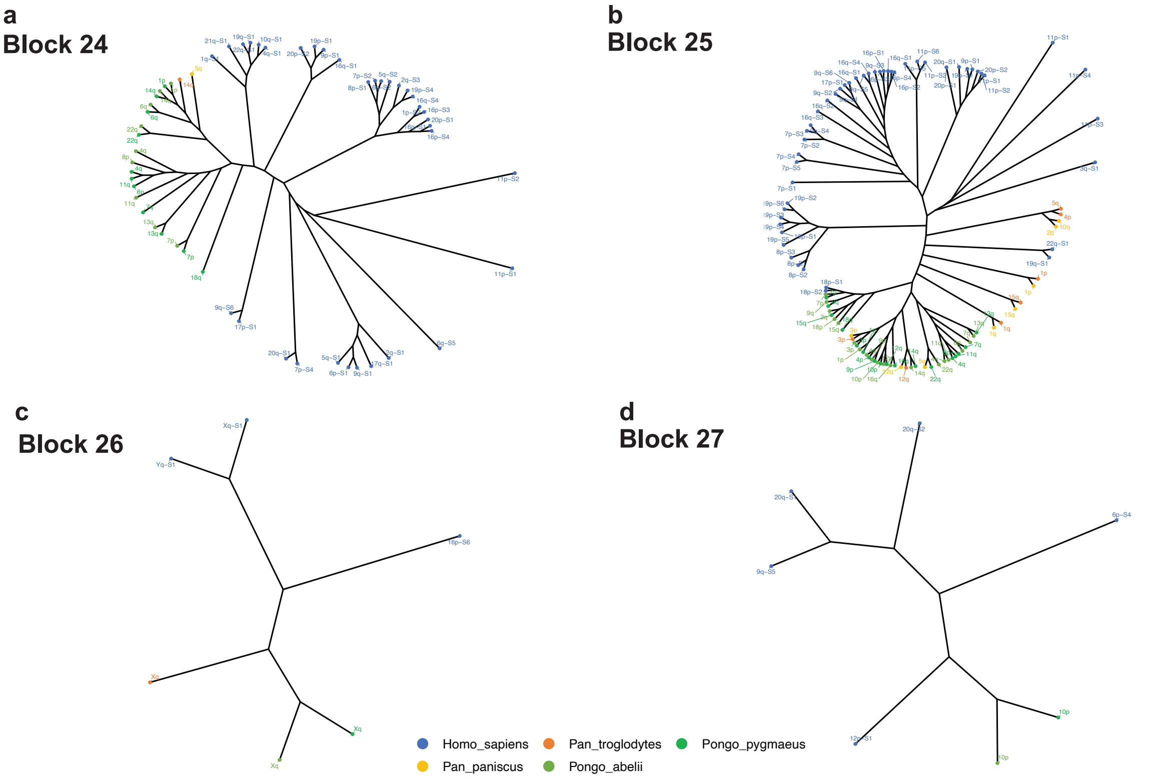


**Supplementary Fig. S114. Maximum likelihood phylogenetic trees for subtelomeric blocks in human and great apes for Block 24-27.** (a) Block 24, (b) Block 25, (c) Block 26 and (d) Block 27. Tip labels indicate chromosomal arm of origin, coloured by species. Human sequences represent representative sequences from each haplotype.

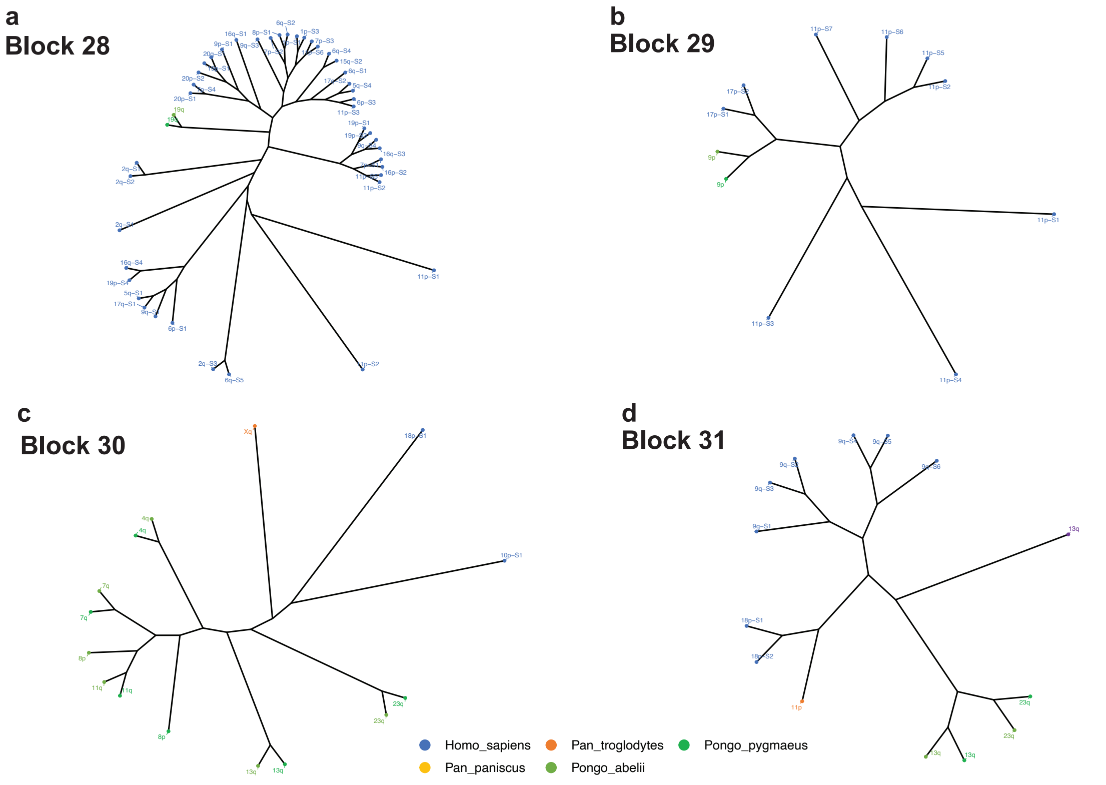


**Supplementary Fig. S115. Maximum likelihood phylogenetic trees for subtelomeric blocks in human and great apes for Block 28-31.** (a) Block 28, (b) Block 29, (c) Block 30 and (d) Block 31. Tip labels indicate chromosomal arm of origin, coloured by species. Human sequences represent representative sequences from each haplotype.

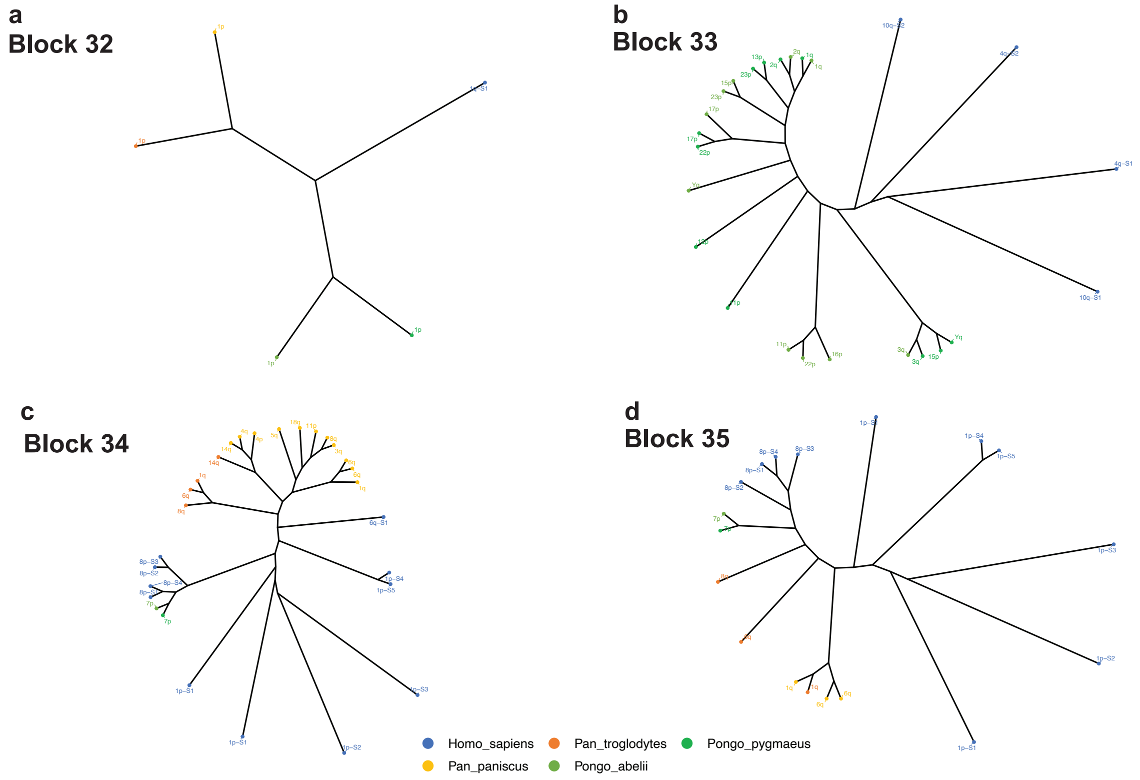


**Supplementary Fig. S116. Maximum likelihood phylogenetic trees for subtelomeric blocks in human and great apes for Block 32-35.** (a) Block 32, (b) Block 33, (c) Block 34 and (d) Block 35. Tip labels indicate chromosomal arm of origin, coloured by species. Human sequences represent representative sequences from each haplotype.

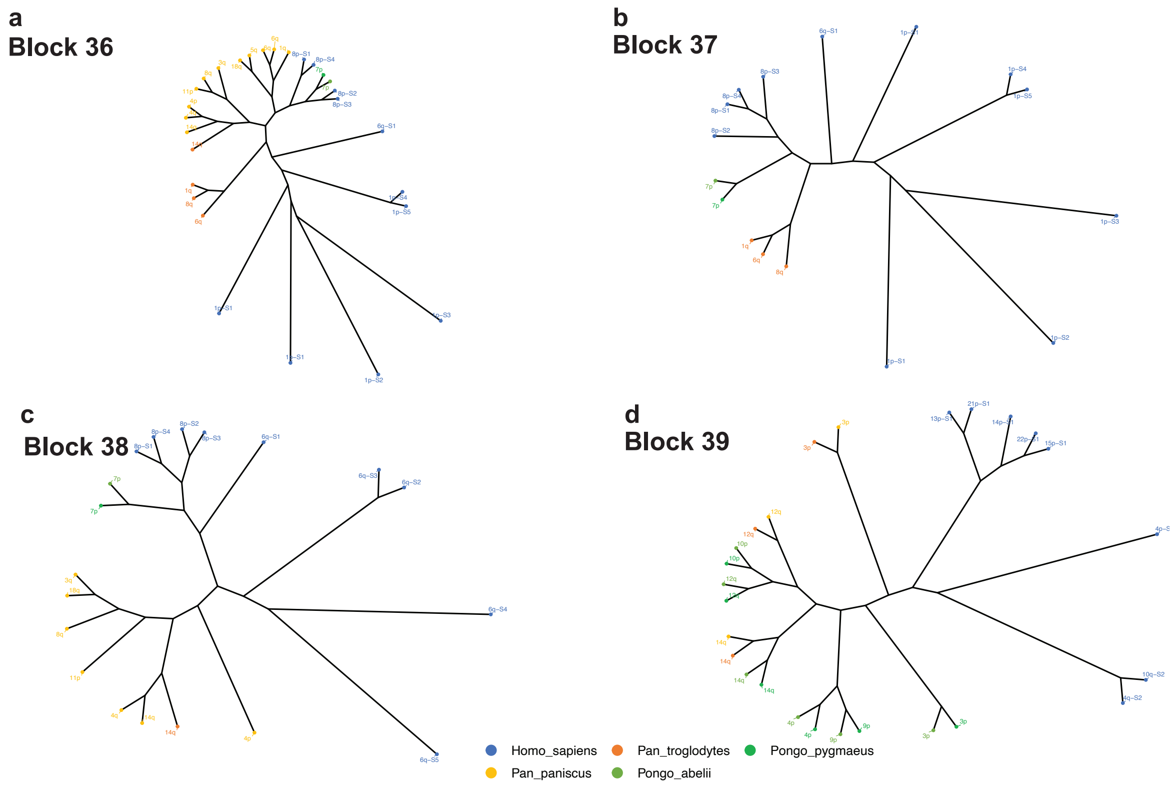


**Supplementary Fig. S117. Maximum likelihood phylogenetic trees for subtelomeric blocks in human and great apes for Block 36-39.** (a) Block 36, (b) Block 37, (c) Block 38 and (d) Block 39. Tip labels indicate chromosomal arm of origin, coloured by species. Human sequences represent representative sequences from each haplotype.

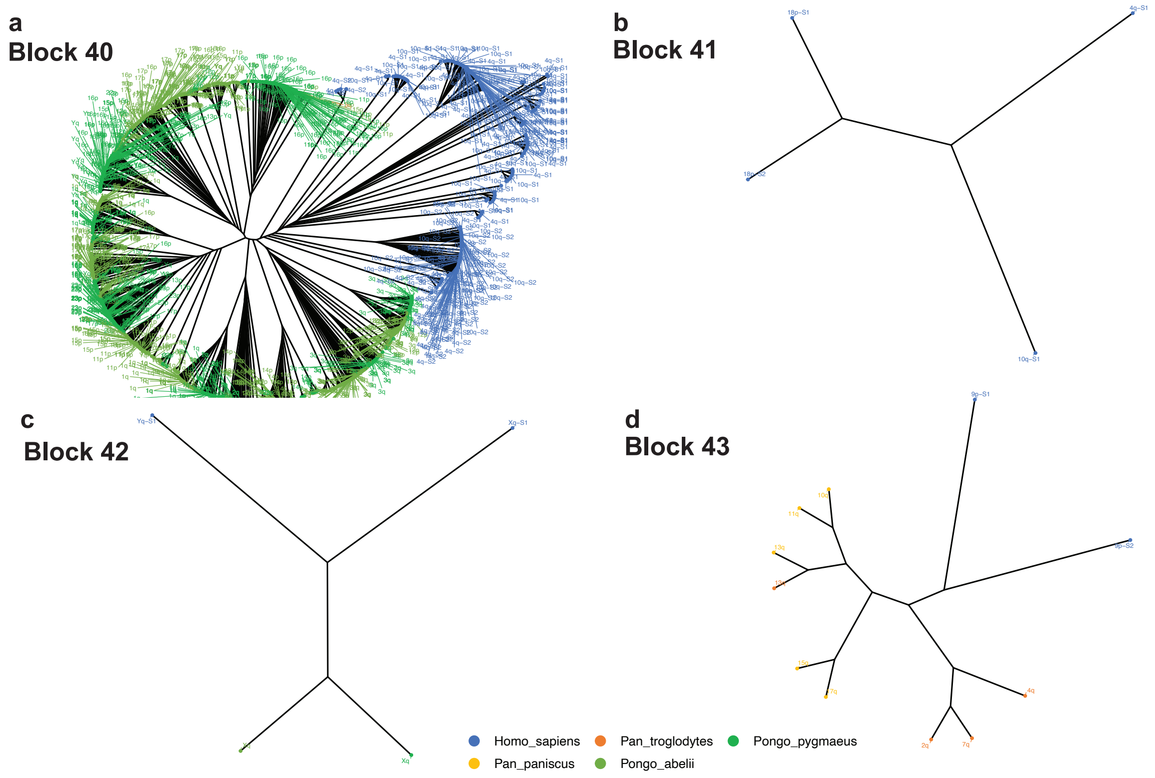


**Supplementary Fig. S118. Maximum likelihood phylogenetic trees for subtelomeric blocks in human and great apes for Block 40-43.** (a) Block 40, (b) Block 41, (c) Block 42 and (d) Block 43. Tip labels indicate chromosomal arm of origin, coloured by species. Human sequences represent representative sequences from each haplotype.

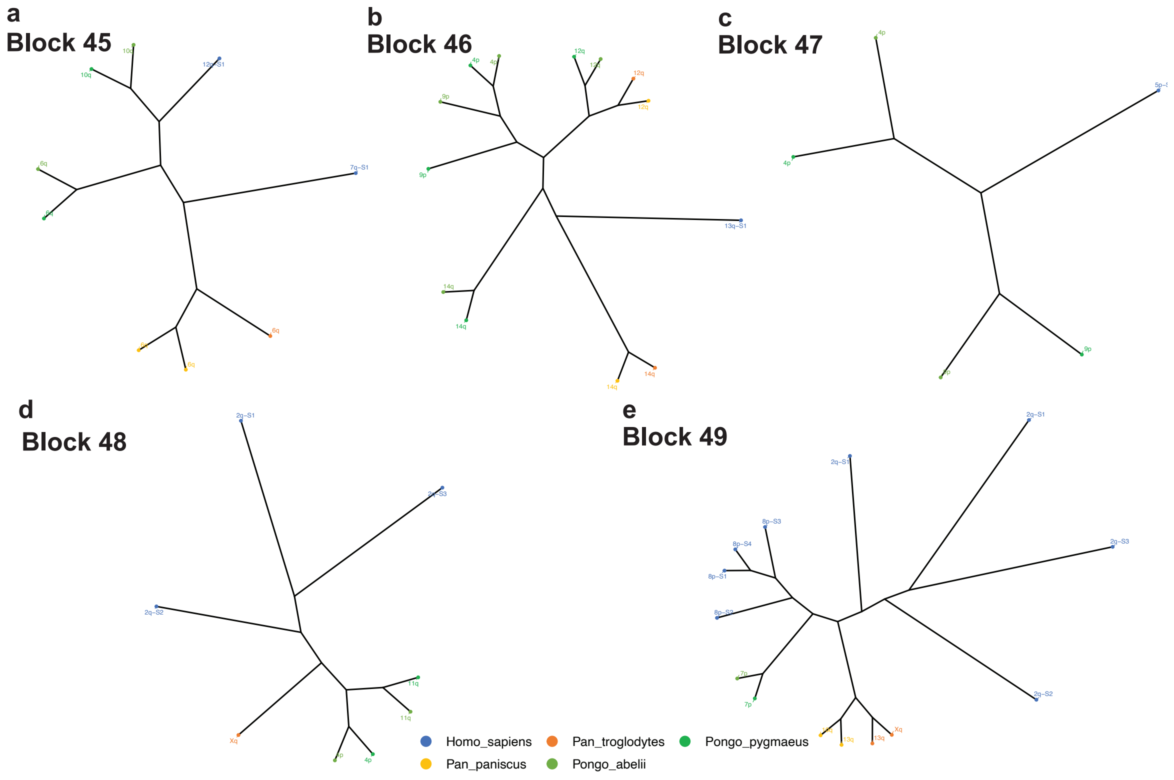


**Supplementary Fig. S119. Maximum likelihood phylogenetic trees for subtelomeric blocks in human and great apes for Block 45-49.** (a) Block 45, (b) Block 46, (c) Block 47, (d) Block 48 and (e) Block 49. Tip labels indicate chromosomal arm of origin, coloured by species. Human sequences represent representative sequences from each haplotype.

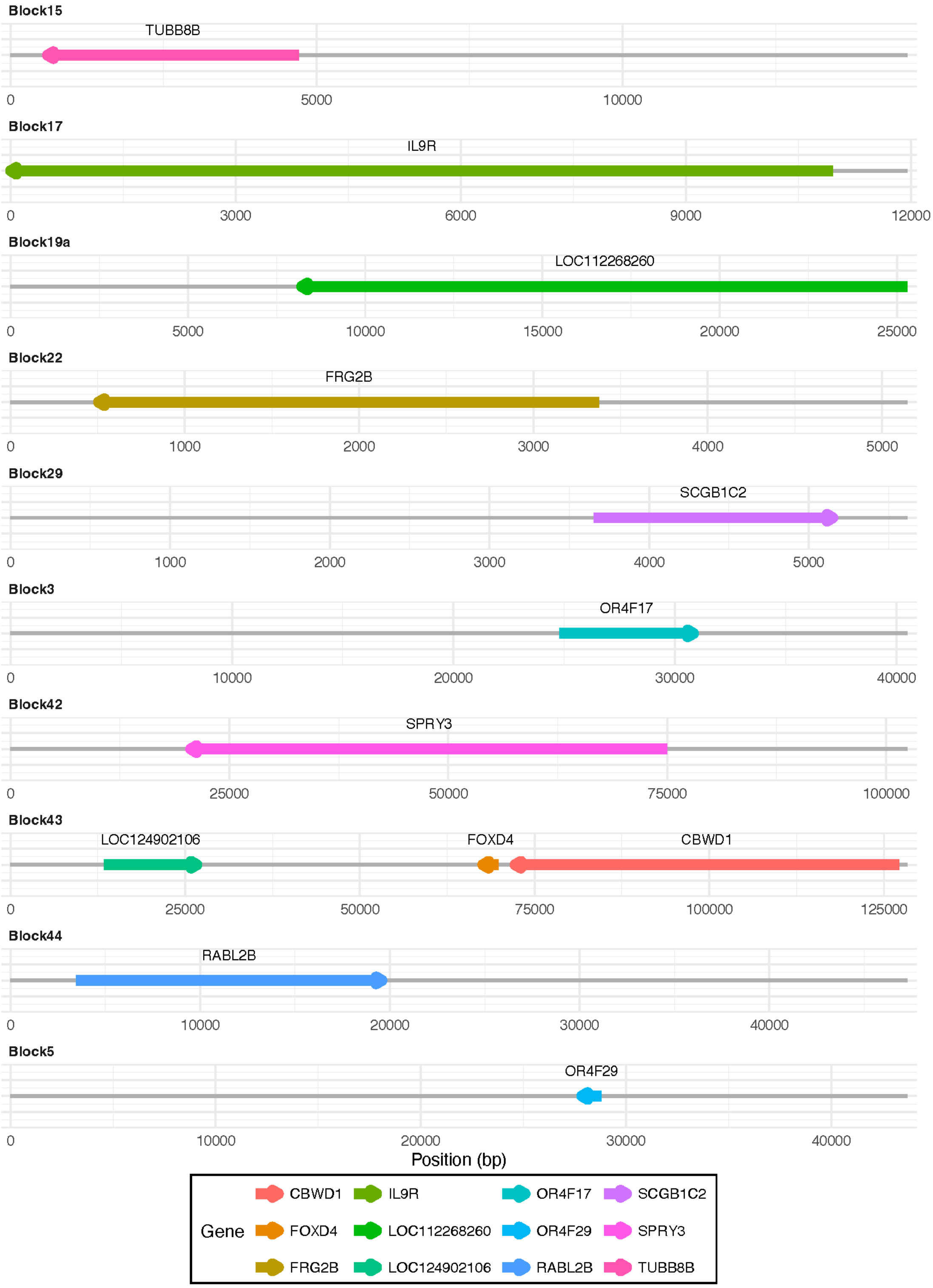


**Supplementary Fig. S120: Protein-coding genes identified within Stong et al. (2014) subtelomeric reference blocks.** Genomic feature plot of protein-coding genes detected in each subtelomeric block, identified using Liftoff with sequence identity and coverage thresholds of 0.8 against the HPRC annotation. Each panel represents a single block, with the grey horizontal bar indicating the full block span and coloured arrows indicating gene position and transcriptional orientation. Gene identity is indicated by colour as shown in the legend.

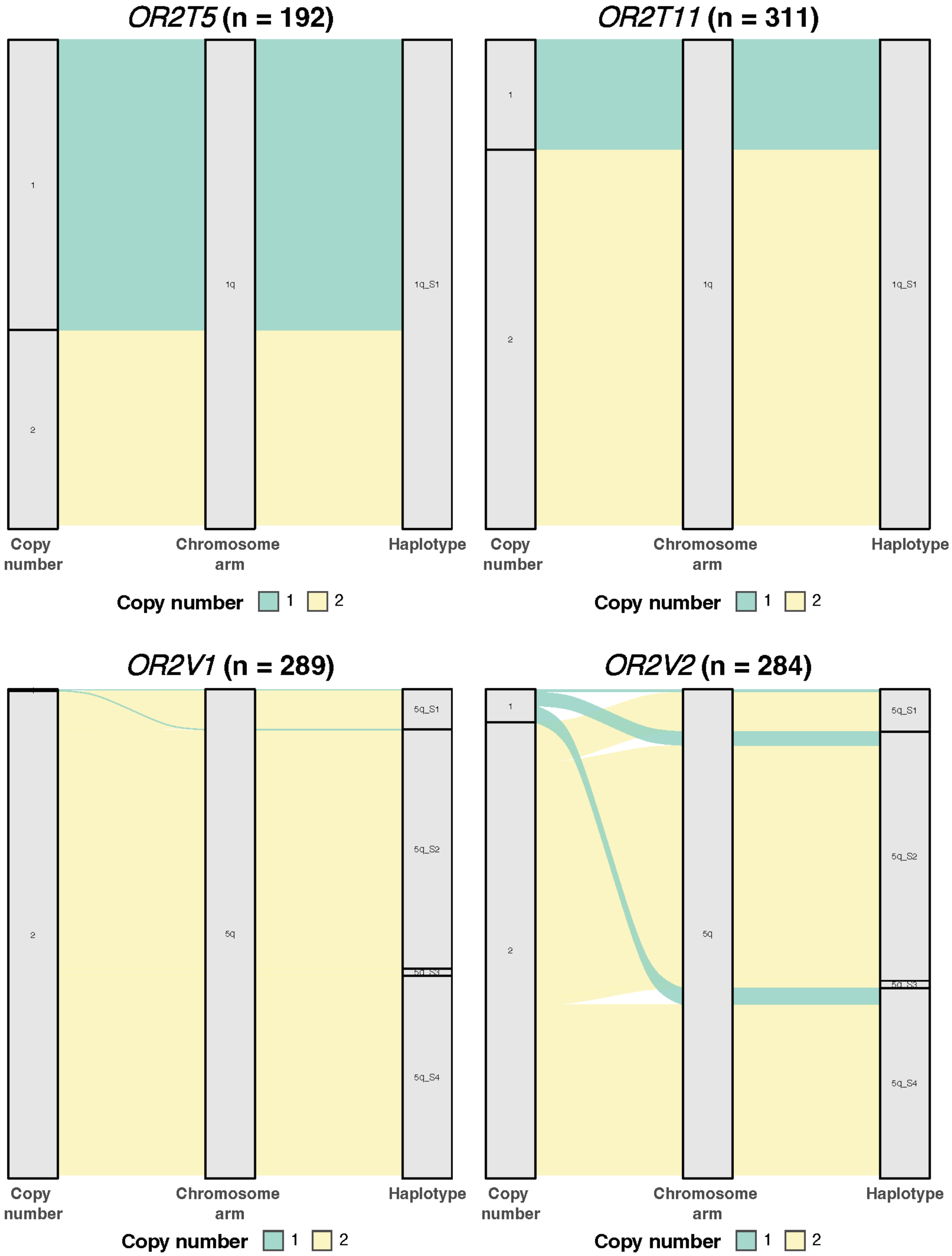


**Supplemenntary Fig. S121: Copy number, chromosomal arm and haplotype distribution of olfactory genes (I).** Alluvial plot linking, from left to right, the copy number observed per individual, the chromosomal arm carrying those copies, and the specific haplotype cluster within that arm for four olfactory genes, OR2T5, OR2T11, OR2V1 and OR2V2.

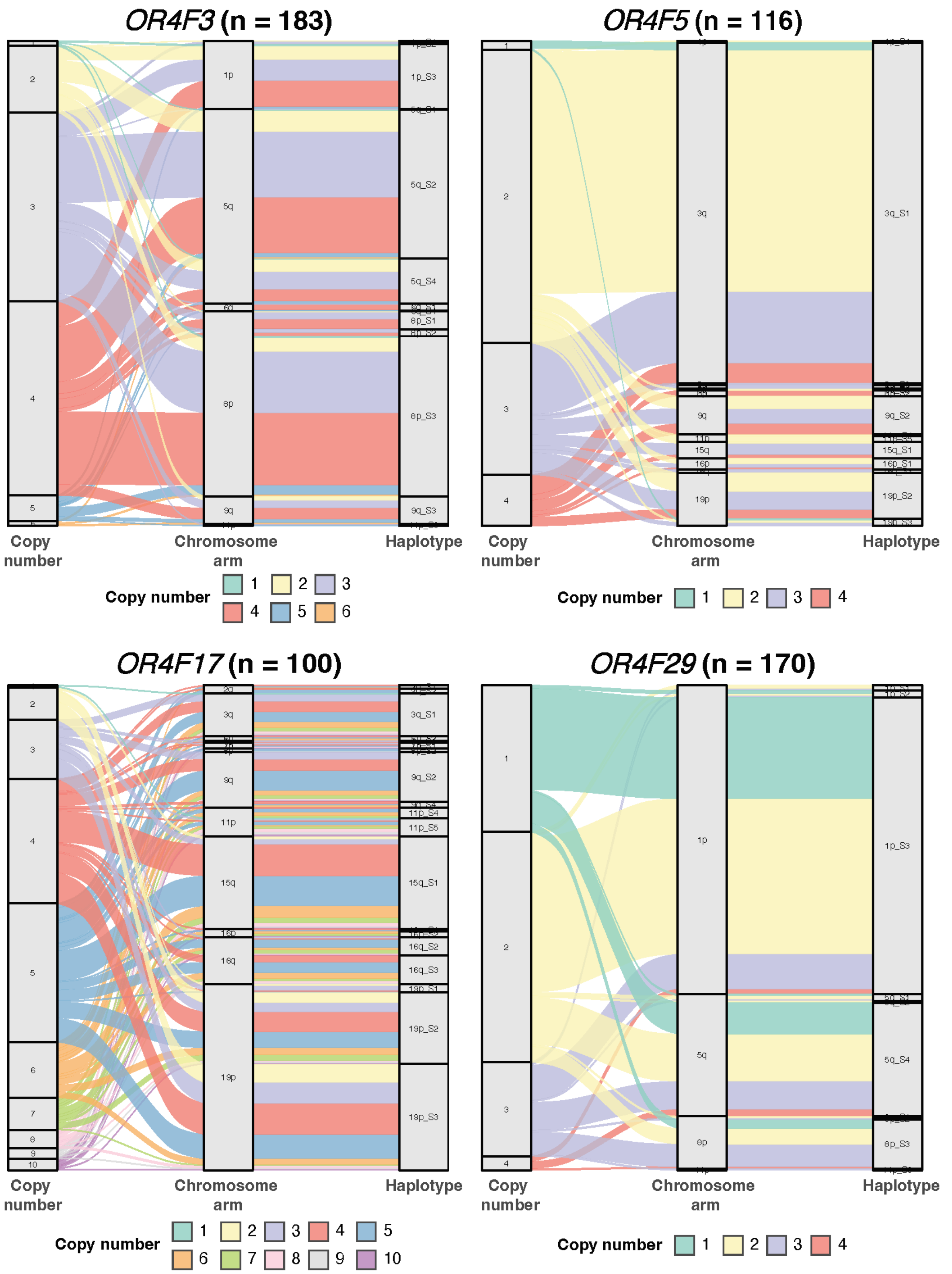


**Supplemenntary Fig. S122: Copy number, chromosomal arm and haplotype distribution of olfactory genes (II).** Figure interpretation follows that of Supplementary Figure S122, but for genes OR4F3, OR4F5, OR4F17 and OR4F29.

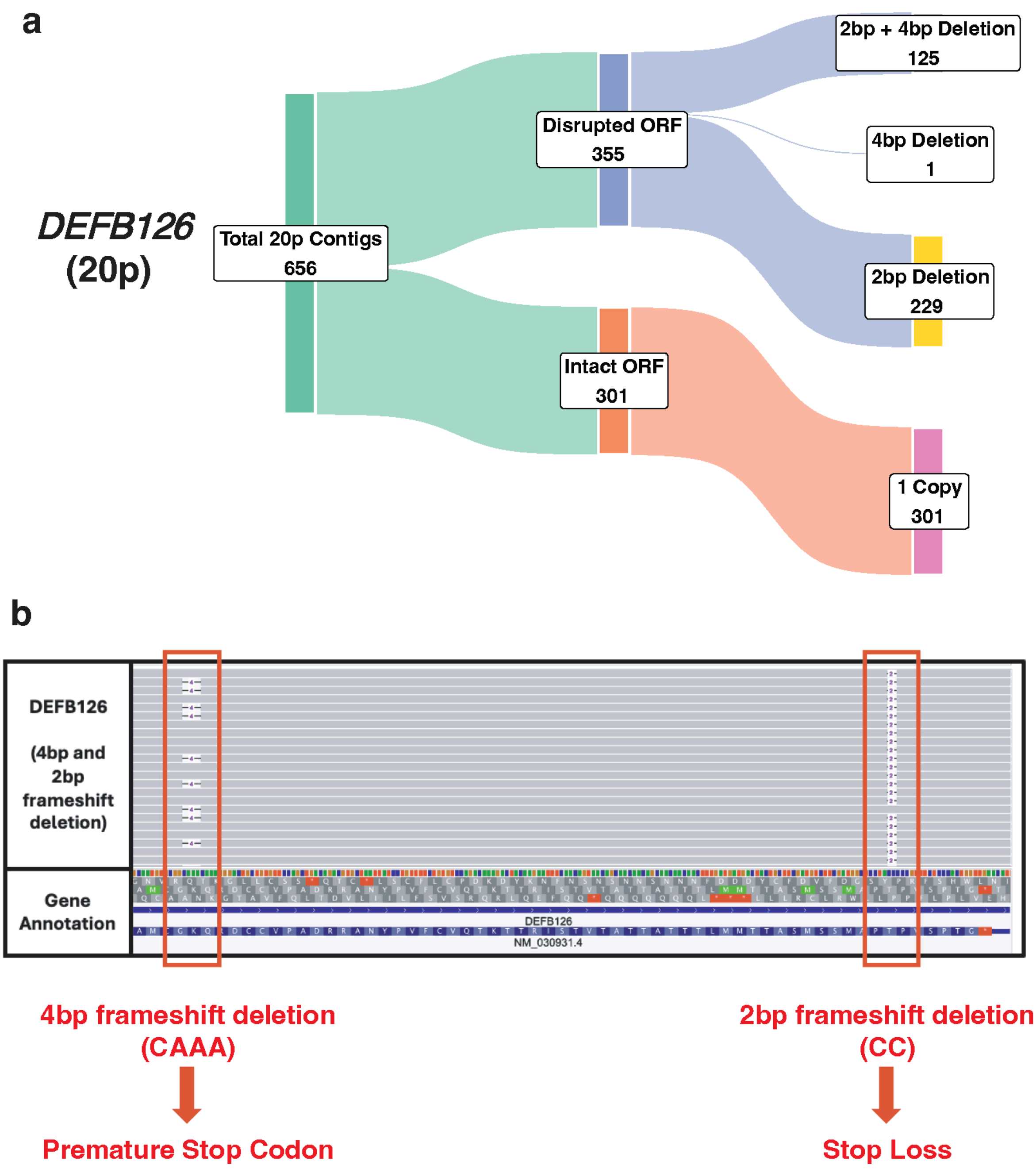


**Supplementary Fig. S123: Classification of DEFB126 contigs on chromosome arm 20p. (a)** Sankey diagram showing the classification of 20p contigs by DEFB126 open reading frame (ORF) status. Contigs with a disrupted ORF are further classified by variant type, comprising deletion status. Contigs with an intact ORF are classified by copy number. **(b)** IGV screenshot of invalid DEFB126 contigs harboring both a 4bp deletion (CAAA) resulting in a premature stop codon, and a 2bp frameshift deletion (CC) that results in stop loss.

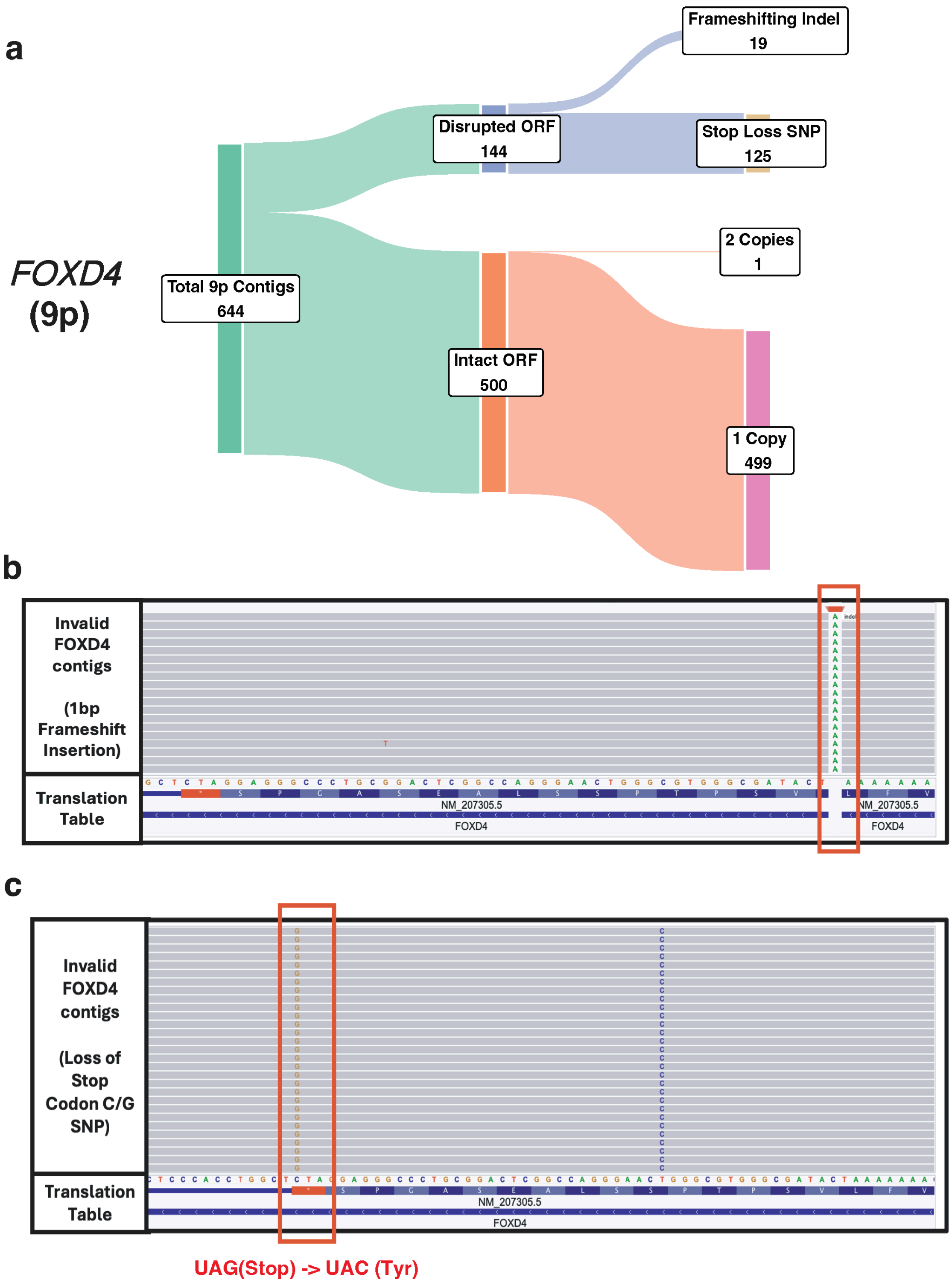


**Supplementary Fig. S124: Classification of FOXD4 contigs on chromosome arm 9p. (a)** Sankey diagram showing the classification of 9p contigs by FOXD4 open reading frame (ORF) status. Contigs with a disrupted ORF are further classified by variant type, comprising frameshifting indels and stop-loss SNPs. Contigs with an intact ORF are classified by copy number, single copy and two copies. **(b)** IGV screenshot of contigs with a disrupted ORF due to a 1bp frameshift insertion. **(c)** IGV screenshot of contigs with a C/G stop-loss SNP at the FOXD4 stop codon.

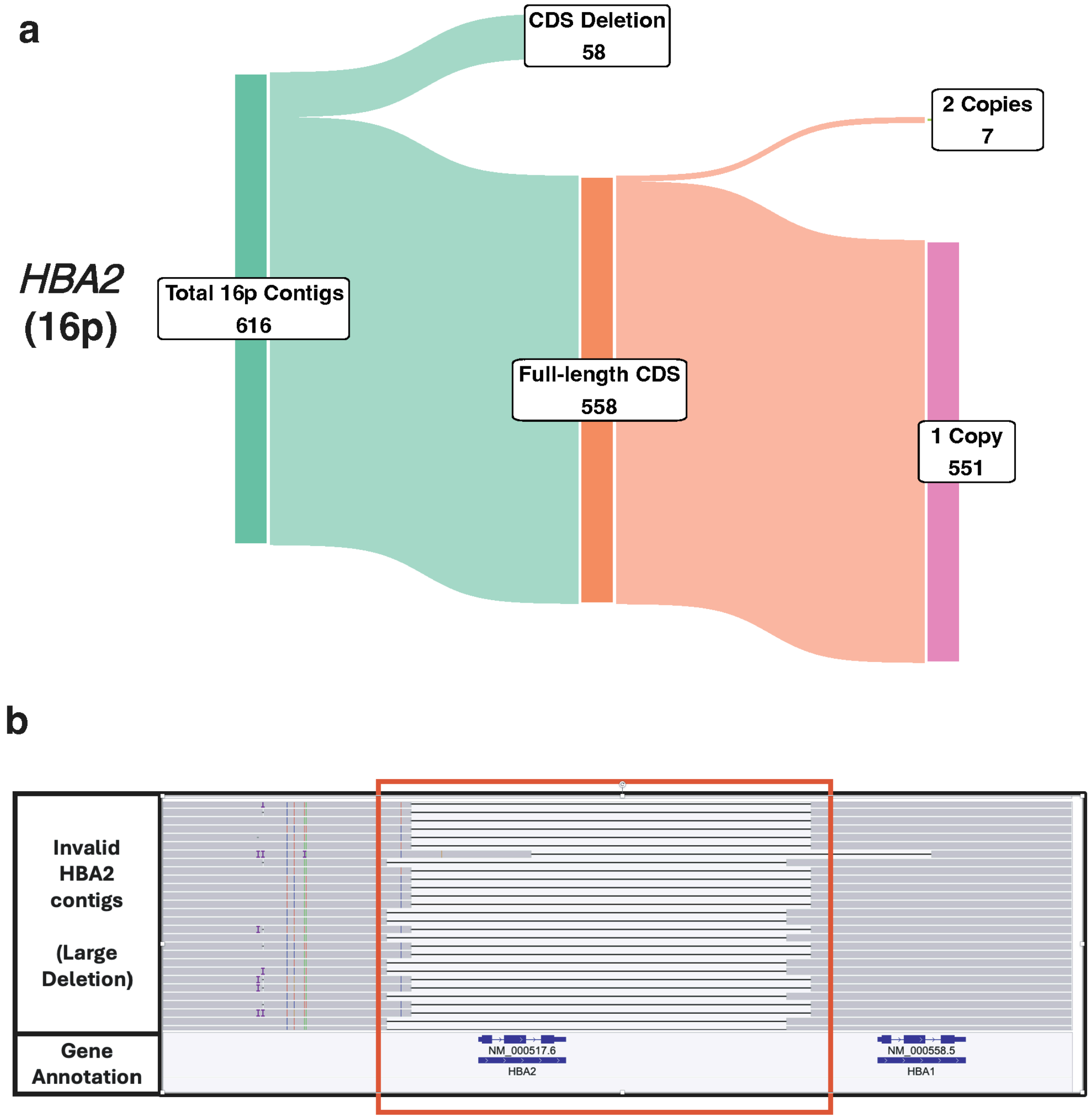


**Supplementary Fig. S125: Classification of HBA2 contigs on chromosome arm 16p. (a)** Sankey diagram showing the classification of 16p contigs by HBA2 coding sequence (CDS) status. Contigs with a CDS deletion represent invalid HBA2 assemblies, while contigs with valid an intact CDS are further classified by copy number. **(b)** IGV screenshot of invalid HBA2 contigs harboring a large deletion spanning the HBA2 locus.

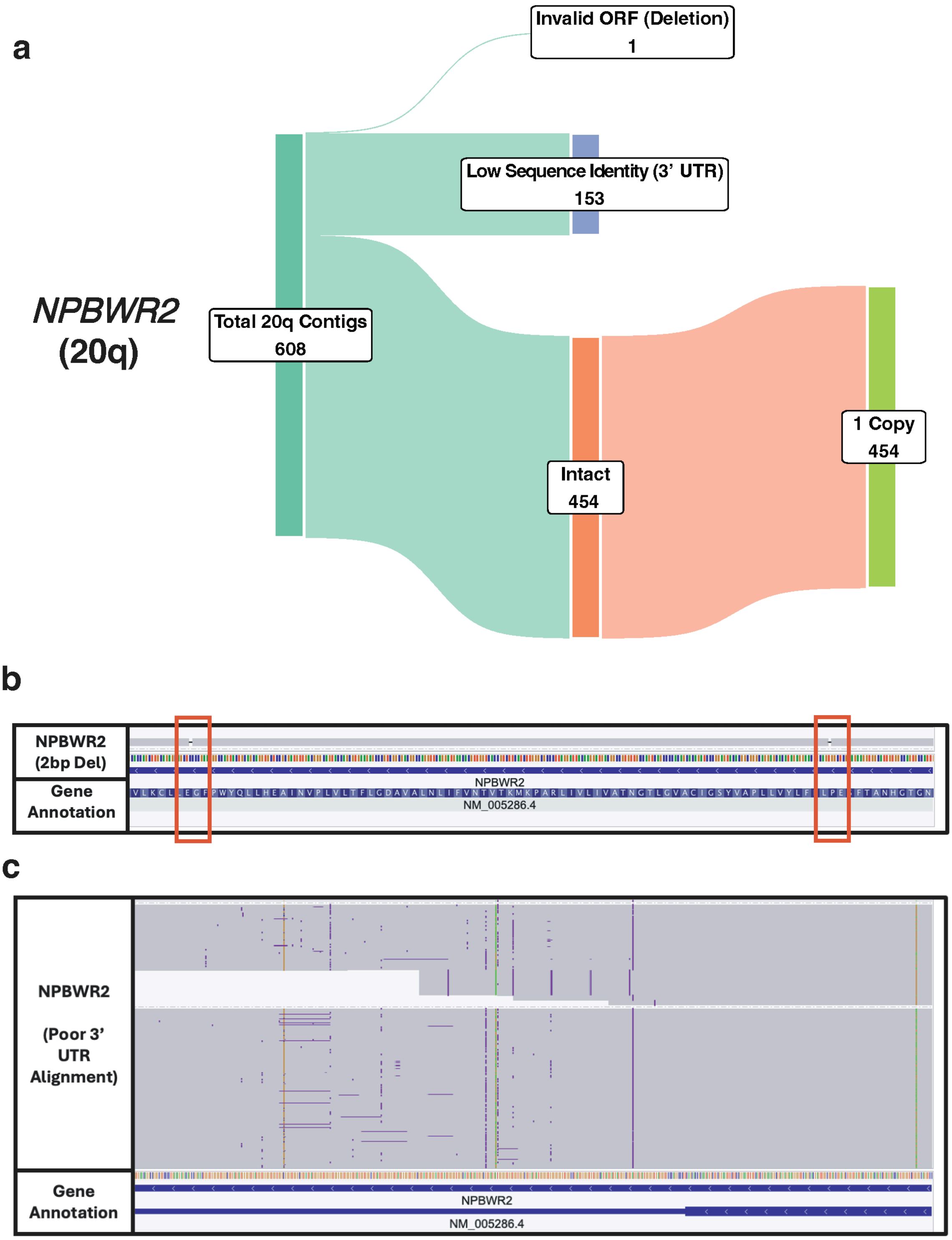


**Supplementary Fig. S126: Classification of NPBWR2 contigs on chromosome arm 20q. (a)** Sankey diagram showing the classification of 20q contigs by NPBWR2 according to Liftoff predictions. Contigs were classified into 3 categories: invalid ORF due to deletion, low sequence identity at the 3' UTR, and intact CDS. **(b)** IGV screenshot of an NPBWR2 contig harbouring a 2 bp deletion. **(c)** IGV screenshot of NPBWR2 contigs showing poor alignment identity at the 3' UTR region.

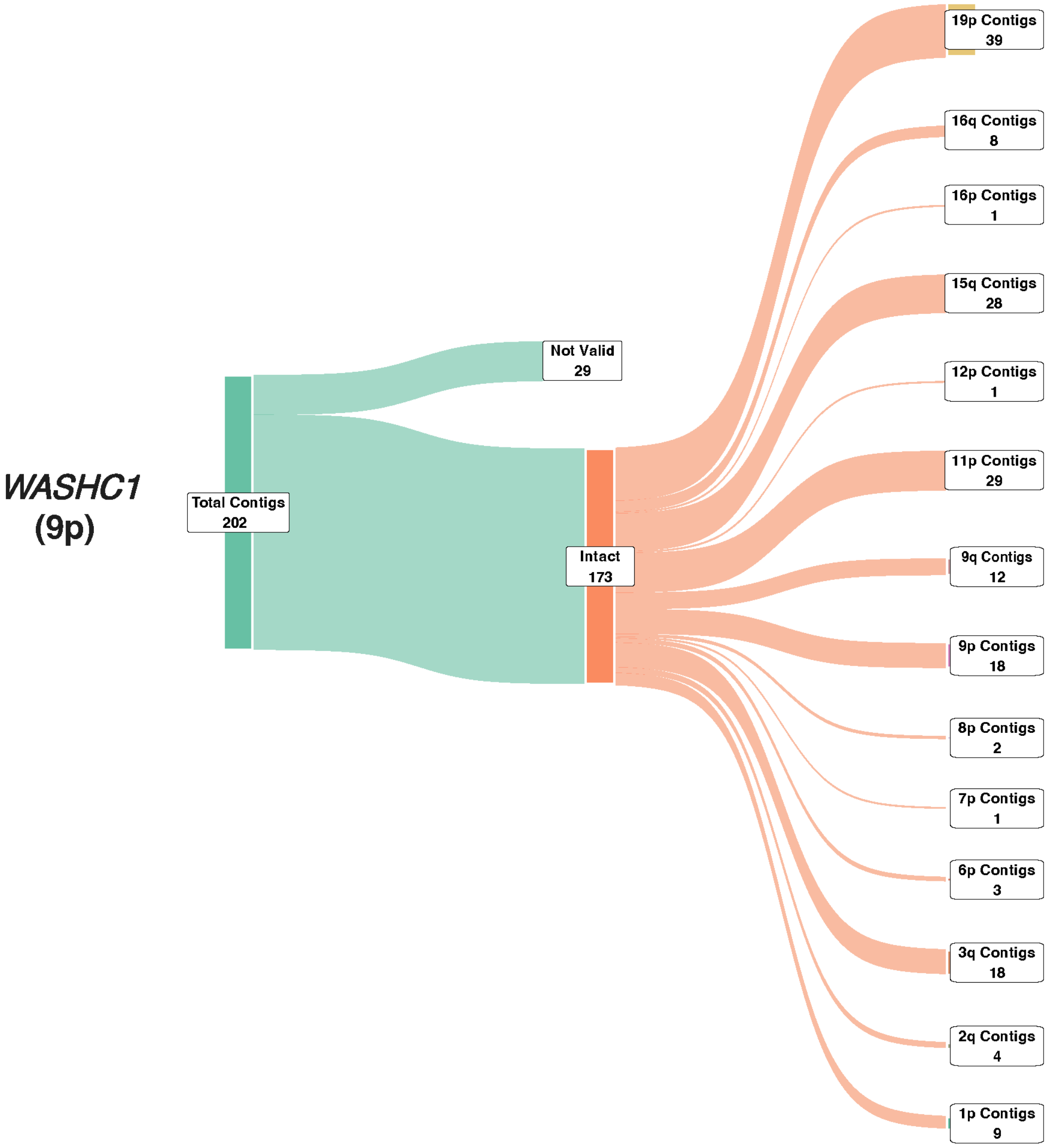


**Supplementary Fig. S127: Classification of WASHC1 contigs on multiple chromosomal arms.** Sankey diagram showing the distribution of chromosomal arms detected with WASHC1.

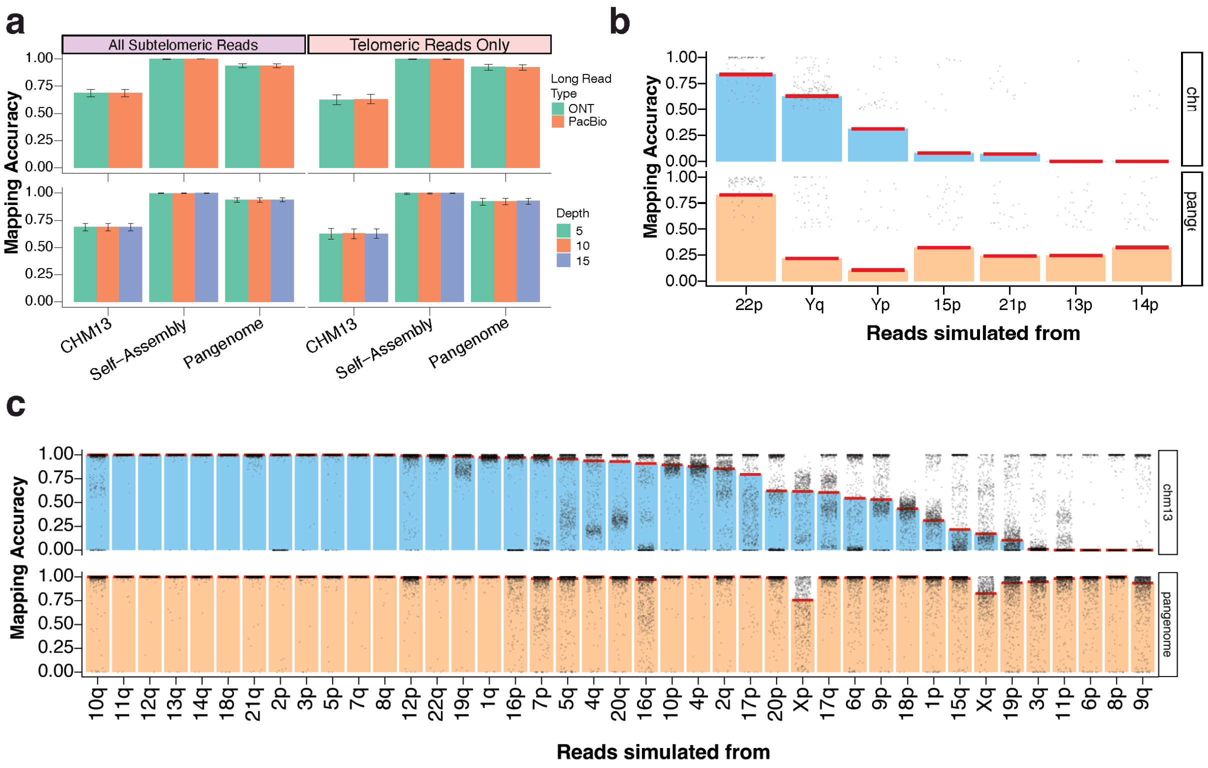


**Supplementary Fig. S128: Subtelomere Simulation with additional parameters. (a)** Barplot of mapping accuracy for all subtelomeric and telomeric-only reads across references, stratified by simulated sequencing technology (PacBio HiFi, ONT; top) and average depth (5x, 10x, 15x; bottom). **(b)** Barplot of mapping accuracy of simulated telomeric reads (15kb, 95% accuracy) to CHM13 (top, blue) and the pangenome reference (bottom, orange), for acrocentric arms and ChrY. Each point represents one sample; red line indicates the median. **(c)** Barplot of mapping accuracy of all subtelomeric reads (10kb, 95% accuracy) to CHM13 (top, blue) and the pangenome reference (bottom, orange), shown per chromosomal arm. Each point represents one sample; red line indicates the median.

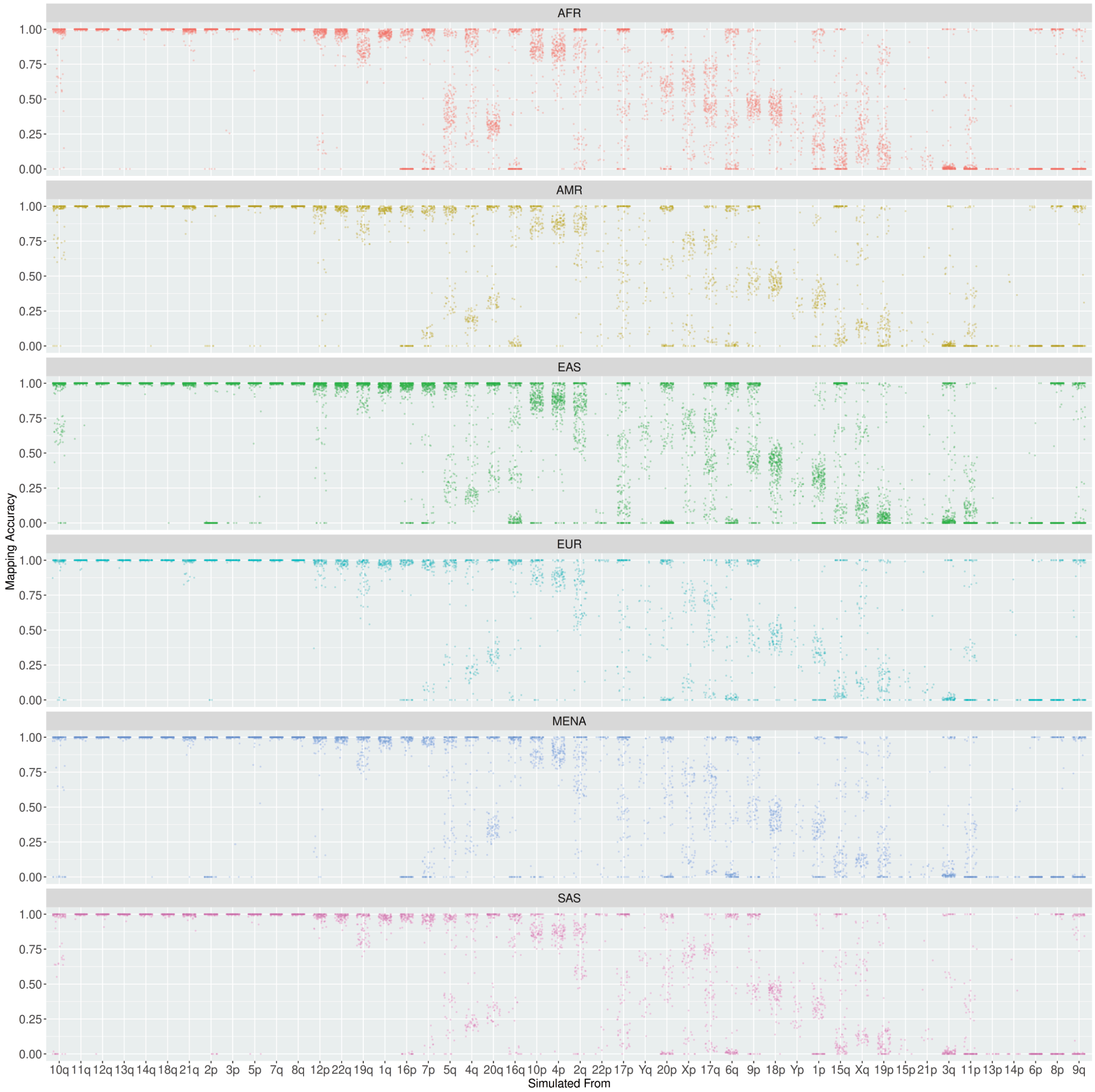


**Supplementary Fig. S129: Subtelomere Simulation stratified by ancestry.** Jitter plot of mapping accuracy to the pangenome reference for simulated telomeric reads (15kb, 95% accuracy, 15x), stratified by sample ancestry. Each point represents one sample.

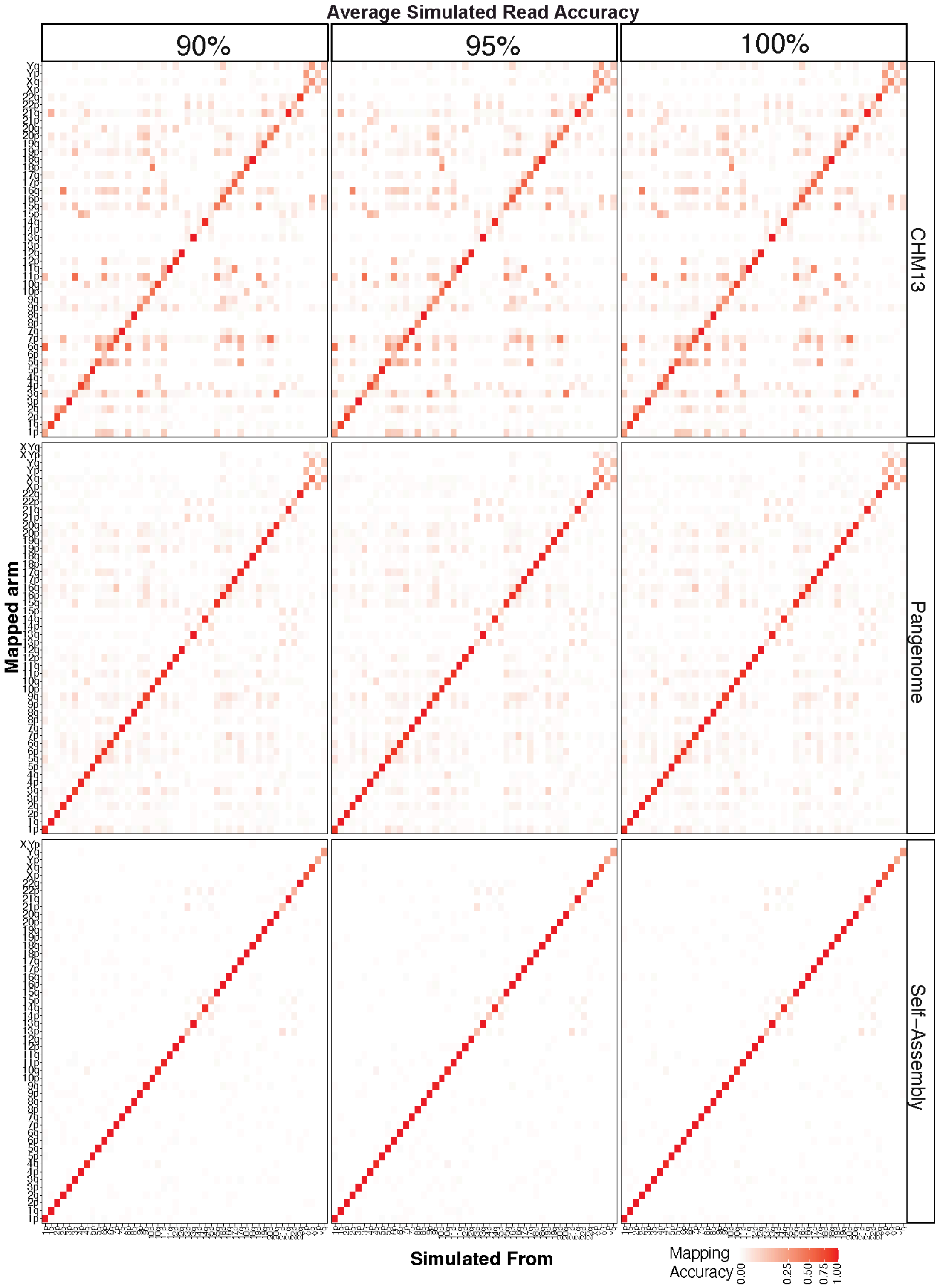


**Supplementary Fig. S130: Subtelomere simulation stratified by read accuracy.** Heatmaps of simulated telomeric read mapping accuracy from each arm to individual chromosomal arms across all three references. Baseline parameters: 20 kb read length, 15x depth, 95% read accuracy. In this panel, read accuracy is varied (90%, 95%, 100%), with all other parameters held at baseline.

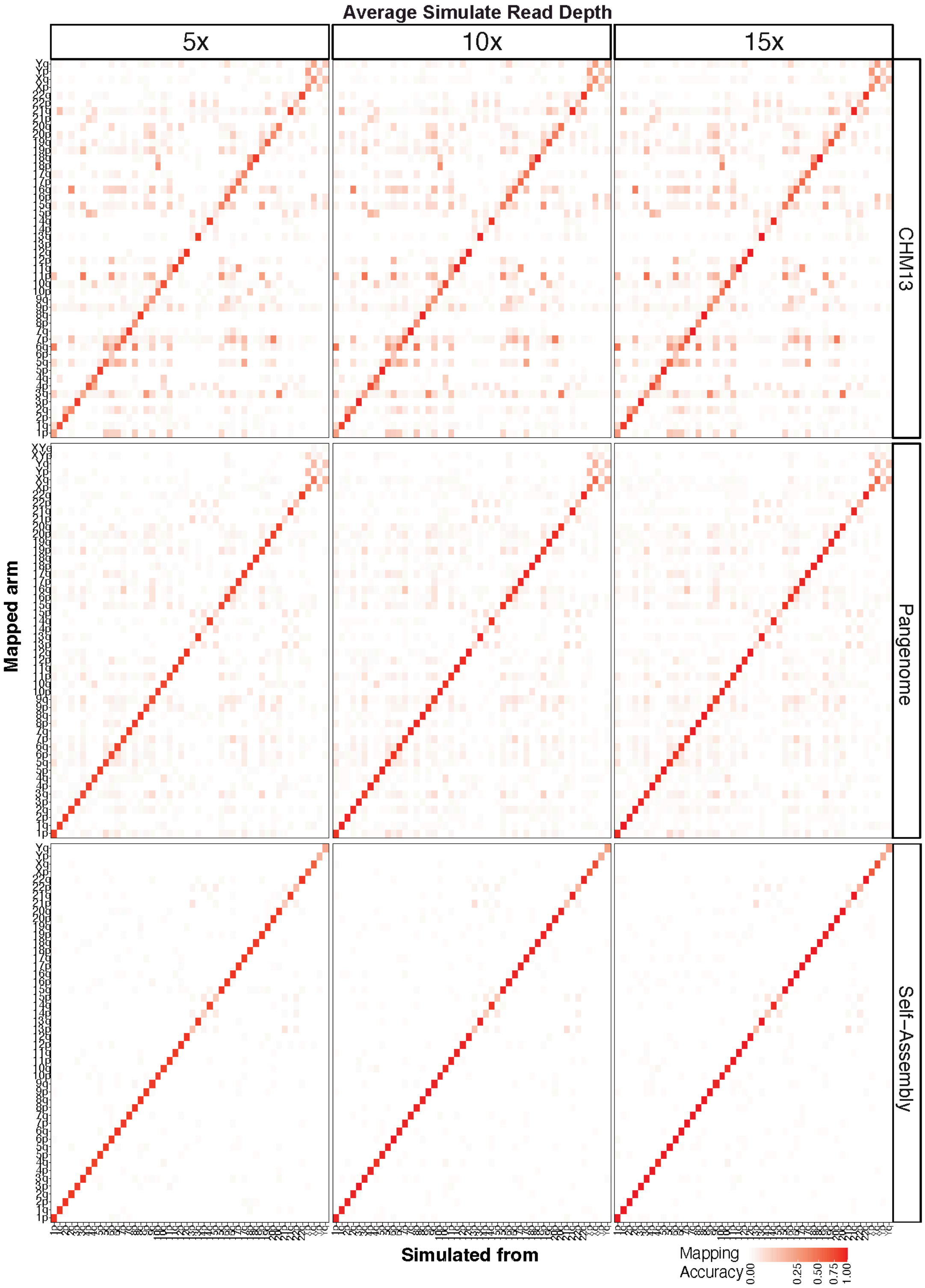


**Supplementary Fig. S131: Subtelomere simulation stratified by sequencing depth.** As Supplementary Fig. S130, but with sequencing depth varied (5x, 10x, 15x). Read length (20 kb) and read accuracy (95%) are held at baseline.

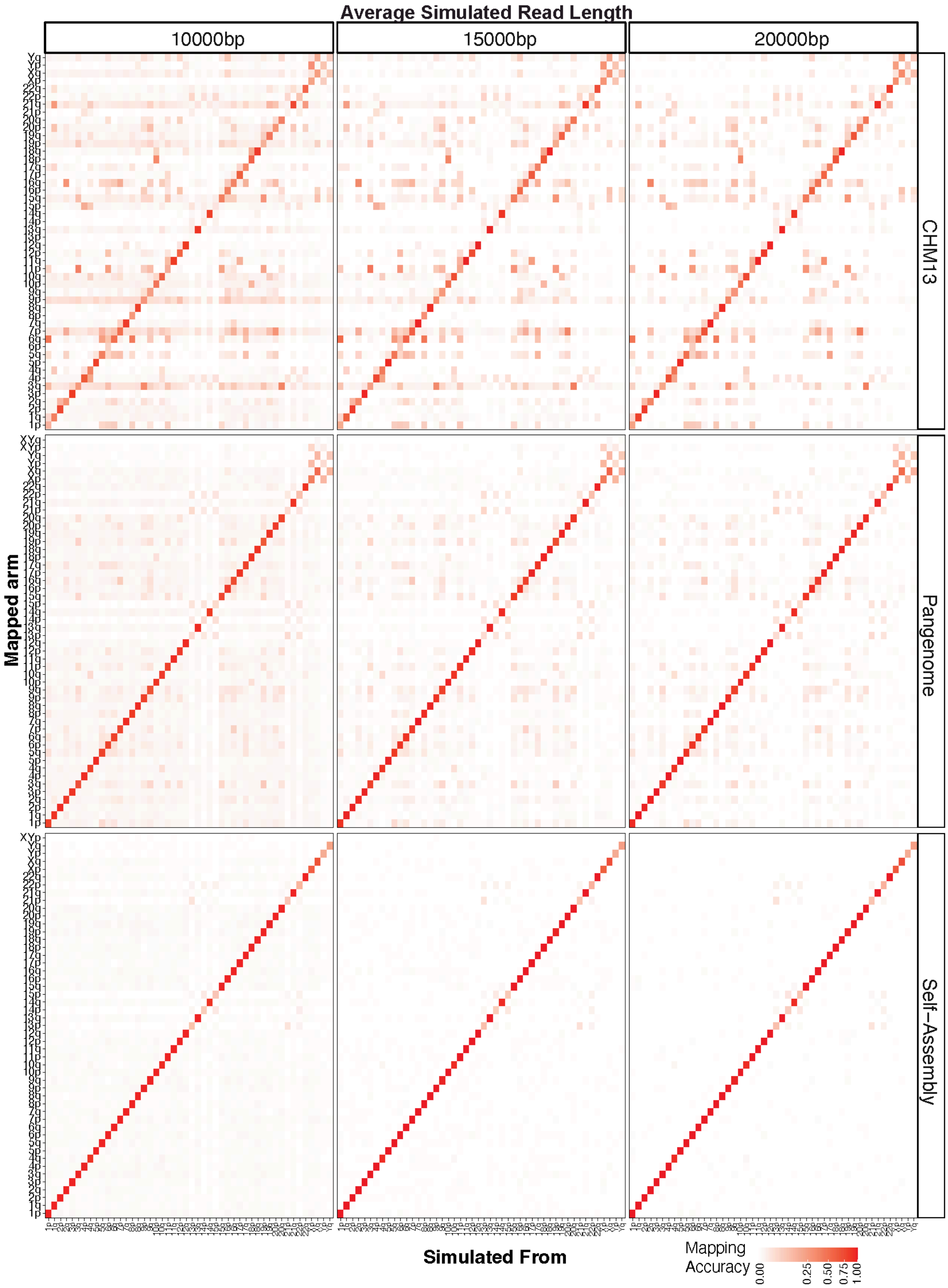


**Supplementary Fig. S132: Subtelomere simulation stratified by read length.** As Supplementary Fig. S130, but with read length varied (10 kb, 15 kb, 20 kb). Sequencing depth (15x) and read accuracy (95%) are held at baseline.

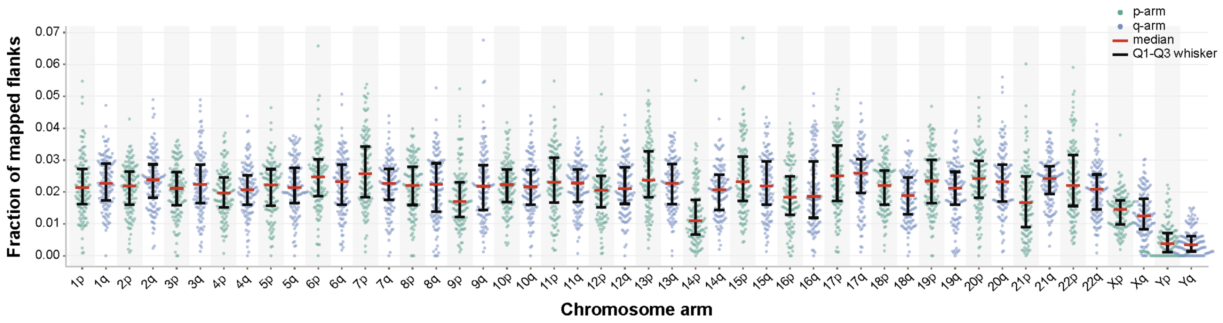


**Supplementary Fig. S133: Chromosome arm mapping fractions are broadly consistent across human samples.** Beeswarm plot showing the fraction of mapped telomeric flank sequences assigned to each human chromosome arm across 128 long-read sequencing samples. Fractions were calculated using all best-scoring mappings that passed the mapping quality filters and are expressed relative to the total number of mapped flanks per sample. Each point represents an individual sample; green points denote p-arms and purple points denote q-arms. Red horizontal lines indicate the median, and black whiskers represent the interquartile range (Q1–Q3). Reads with equally supported best-scoring mappings contributed to all corresponding chromosome arms.
